# A Multi-Omic Phenobank Reveals Axes of Glioblastoma Growth, Invasion, and Therapeutic Vulnerability

**DOI:** 10.1101/2025.03.25.645260

**Authors:** Cecilia Krona, Soumi Kundu, Emil Rosen, Filippa Kruse, Madeleine Skeppås, Haris Babačić, Ida Larsson, Ludmila Elfineh, Maya Jeje Schuan Lü, Maria Escriva Conde, Ramy Elgendy, Zankruti Dave, Milena Doroszko, Katrin Rut-Halldorsdottir, Xiaofang Cao, Rashmi Ramachandra, Karl Holmberg Olausson, Mats Nilsson, Joachim Weischenfeldt, Johan Wikström, Maria Pernemalm, Anders Sundström, Irem Uppman, Hitesh Bhagavanbhai Mangukiya, Sven Nelander

## Abstract

**Background:** Glioblastoma (GBM) invasion is clinically decisive but difficult to model systematically. Existing patient-derived xenograft (PDX) resources rarely couple reproducible *in vivo* invasion phenotypes with matched multi-omic profiles at scale, limiting mechanistic insight and phenotype-informed therapeutic hypotheses.

**Methods:** We established the **HGCC Phenobank**, comprising 65 patient-derived GBM stem-like cultures with matched multi-omic profiling and orthotopic engraftment in 449 mice. Blinded histopathology quantified ten invasion traits per case. These phenotypes were integrated with RNA sequencing, DNA methylation, and mass-spectrometry-based proteomics. Multi-Omic Factor Analysis (MOFA) identified latent molecular programs. Phenotype-specific RNA signatures were matched to LINCS drug-perturbation profiles and validated in 3D gliomasphere and *ex vivo* brain-slice assays.

**Results:** Two dominant, reproducible invasion modes emerged across models: *diffuse parenchymal infiltration* and *perivascular/condensed growth*. Proneural cultures formed more aggressive tumors in immunodeficient mice, and mouse survival showed a modest correlation with patient survival in matched cases (Pearson 0.1832, 0.045). MOFA identified 15 latent factors; **Factor 1**, enriched for *ASCL1/OLIG1/OLIG2* programs and associated with *TP53/DCHS2/WNK2* alterations, was linked to increased tumor formation, diffuse invasion, and shorter mouse survival, and stratified GBM patients in TCGA and in our matched patient cohort. Drug-signature matching separated mechanisms targeting diffuse versus perivascular invasion. Experimental validation confirmed phenotype-selective sensitivities, and inhibitors PIK-75 and buparlisib suppressed invasion dynamics across representative models in 3D and brain-slice assays.

**Conclusions:** The HGCC Phenobank provides the first openly available PDX resource that systematically links GBM invasion phenotypes to multi-omic programs and therapeutic predictions. This framework enables reproducible model selection, mechanistic dissection of invasion modes, and phenotype-guided therapeutic discovery.

**Key Points:**

- Diffuse and perivascular invasion define orthogonal GBM axes
- ASCL1/OLIG factor links initiation, diffuse growth, and survival
- Phenotype-matched drugs validated; PIK-75 and buparlisib curb invasion dynamics

**Importance of the Study:** Glioblastoma invasion varies substantially between patients, yet existing patient-derived xenograft resources rarely combine reproducible *in vivo* phenotyping with matched multi-omic profiling at scale. The HGCC Phenobank addresses this gap with standardized, blinded scoring of ten invasion traits across 449 orthotopic xenografts from 65 molecularly characterized GBM stem-like cultures, integrated with transcriptomic, methylomic, and proteomic data. We identify two dominant, reproducible invasion modes and a cross-modal neurodevelopmental program, the ASCL1/OLIG1/2-associated Factor 1, that links tumor initiation, diffuse growth, and survival in mice, and stratifies GBM patients in TCGA and in our matched patient cohort. In a spatially resolved xenograft section, Factor 1 signal localizes to the invasive tumor periphery. By matching phenotype-specific RNA signatures to drug-induced transcriptional responses, we show that invasion phenotypes nominate selective vulnerabilities, exemplified by PIK-75. This openly shared resource enables reproducible model selection, mechanistic dissection of invasion programs, and phenotype-guided therapeutic discovery.

## Introduction

Glioblastoma (GBM) remains a lethal disease, with median survival of 15–17 months despite multimodal therapy^1–3^. A defining clinical challenge is the early, infiltrative spread of tumor cells throughout the brain, which limits resectability and predisposes to diffuse or multifocal recurrence^4^. GBM exhibits extensive genetic, epigenetic, and transcriptional heterogeneity^5–7^, and within tissue, tumor cells occupy distinct microenvironmental niches reflected in classical secondary Scherer structures: perivascular invasion, perineuronal satellitosis, and white matter spread^8,9^. These invasive patterns are prominent in diffuse or multifocal presentations and are associated with challenging clinical courses^10^. Together, these considerations underscore the need for scalable *in vivo* models that systematically connect GBM invasion phenotypes to underlying molecular programs and actionable vulnerabilities.

Orthotopic patient-derived xenograft (PDX) models remain the foundation of experimental neuro-oncology: they retain key histopathological and molecular features of patient tumors and enable mechanistic and therapeutic studies in a brain context^7,11,12^. Their fidelity is enhanced when patient-derived glioma stem-like cells are maintained under serum-free, neural stem cell-like conditions^13,14^. However, existing PDX panels rarely capture the full spectrum of patient-to-patient variation in *in vivo* growth and invasion. Cohorts are often modest, phenotyping is seldom blinded or standardized, and emphasis is typically placed on tumor take rate or survival rather than on quantitative invasion traits. As a result, conclusions about mechanisms of diffuse infiltration or perivascular co-option are highly dependent on model selection, and few datasets link *in vivo* invasion phenotypes to matched transcriptomic, DNA-methylation, and proteomic profiles at patient-aligned scale. Furthermore, the ad hoc nature of generating multiple, small grafting cohorts complicates comparisons across studies.

To address these limitations, we established the HGCC Phenobank: a standardized panel of 65 GBM stem-like cultures within the Human Glioblastoma Cell Culture (HGCC) resource^7,15^, with matched multi-omic profiling and orthotopic engraftment in 449 mice. Each culture was engineered to express firefly luciferase and GFP to support longitudinal tracking in live animals and explants. Tumors were scored by blinded observers across ten invasion traits and paired with matched transcriptomic, DNA-methylation, and proteomic profiles. Unlike prior PDX panels (e.g., Vaubel et al.) or anatomical atlases such as the IVY Glioblastoma Atlas^16,17^, the HGCC Phenobank uniquely integrates quantitative in vivo invasion phenotyping with multi-omic data across a broad, patient-aligned cohort.

Using this resource, we identify two dominant invasion modes, diffuse parenchymal infiltration and perivascular/condensed growth, and show that proneural xenografts are more tumorigenic in immunodeficient mice than classical or mesenchymal cultures. Mouse survival shows a modest but significant correlation with survival in corresponding patients (Pearson 0.1832, 0.045). Integrative modeling with Multi-Omic Factor Analysis (MOFA) reveals 15 latent factors capturing coordinated variation across omics layers. Factor 1, enriched for *ASCL1/OLIG1/OLIG2* programs and associated with alterations in *TP53*, *DCHS2*, and *WNK2*, links elevated tumor formation, diffuse invasion, and shortened mouse survival, and stratifies outcomes in TCGA and matched patient cohorts. To translate phenotypes into therapeutic hypotheses, we match phenotype-specific RNA signatures to drug-induced transcriptional perturbations (TargetTranslator), predicting mechanisms aligned along diffuse versus perivascular axes. We validate selective vulnerabilities in 3D gliomasphere invasion and *ex vivo* brain-slice assays; notably, the multi-target PI3K/CDK/TAL1 inhibitor PIK-75 reduces cell velocity and displacement across representative models.

Together, the HGCC Phenobank provides a scalable, phenotype-anchored map linking GBM invasion behavior to latent molecular programs, clinical outcomes, and phenotype-guided therapeutic predictions. This resource enables reproducible model selection and accelerates the mechanistic and translational study of GBM invasion.

## Materials and Methods

### Experimental models

Patient-derived GBM cultures (PDCs) were established from surgical specimens collected with informed consent (ethical permit 2007/353) and maintained within the HGCC biobank^7,15^. Tissue dissociation was followed by adherent culture on laminin-coated Primaria plates (Corning) under serum-free neural stem cell conditions according to standardized HGCC protocols. Cell identity was confirmed by STR profiling (Eurofins Genomics). All PDCs were engineered to express GFP and firefly luciferase using the lentiviral vector pBMN(CMV-copGFP-Luc2-Puro) (Addgene #80389). After puromycin selection, 100,000 cells were stereotactically injected into the striatum of immunodeficient mice to generate patient-derived cell culture xenografts (PDCXs). The project used NOD-scid, NSG, NOG, and NMRI-nude mice as institutional availability and animal-welfare practices evolved over the course of the study; strain information for each injection is provided in Supplementary Table ST1. Because strain-specific survival data were incomplete, including infection-related losses in one strain, strain was not used for formal survival comparisons. Representative histological comparisons across strain contexts are shown in Supplementary Figure SF6. Tumor development was monitored by bioluminescence imaging for up to 40 weeks. Detailed procedures are provided in the Supplementary Methods.

### Case identity reconciliation

Case identities were validated prior to downstream analysis by combining cross-platform transcriptomic matching and STR-based evidence. Direct cross-platform identity auditing was possible for 77 models with both RNA-seq and legacy Affymetrix mRNA profiles available. In most cases, the expected identity was confirmed by RNA best match, while a subset of models underwent identifier harmonization based on concordant transcriptomic and STR evidence before downstream analysis. Multi-omic profiling was performed on cultures prior to orthotopic transplantation. After identifier reconciliation and duplicate collapsing, the identity-reconciled RNA-seq cohort comprised 65 unique HGCC lines. Passage number for each injected culture is provided in Supplementary Table ST1. As an additional check of culture–xenograft coherence, pseudobulk RNA profiles from parental cultures and matched xenograft-derived mouse-brain samples were compared by PCA and sample-sample correlation (**Supplementary Figure SF8**).

### Immunohistochemical analyses

Mouse brains were fixed in 4% phosphate-buffered formaldehyde (Histolab, Sweden) for 2–7 days, divided coronally into five tissue blocks, dehydrated, and embedded in paraffin using an automated processor (TPC15 DUO, Medite Medizintecknik). Im-munohistochemistry was performed on 3 μm sections using antibodies against human-specific nuclear NuMA (Abcam ab97585), cytoplasmic STEM121 (Takara Y40410), and Ki67 (clone MIB-1, Dako M7240). Further details are provided in the Supplementary Methods.

### Image acquisition, tumor initiation scoring, invasion phenotyping, and survival analysis

Procedures for bioluminescence imaging, histopathological scoring of ten prespecified invasion traits, hierarchical estimation of PDC-specific phenotypes, and survival analysis are described in the Supplementary Methods.

### Library preparation and RNA sequencing

Total RNA was extracted using Direct-zol RNA MiniPrep (Zymo Research). Libraries were prepared from 500 ng RNA using the TruSeq Stranded mRNA kit (Illumina #20020595) with polyA selection. Library quality was assessed using the Fragment Analyzer (Agilent) with the DNF-910 dsDNA kit. Sequencing was performed on a NovaSeq 6000 system (Illumina) using paired-end 50 bp reads (SP flowcell v1.5 chemistry), yielding at least 650 million read pairs per flowcell with 85% bases at Q30. Additional details are provided in the Supplementary Methods.

### RNA-seq normalization

Normalization was performed in R v3.3 using edgeR 3.16 with TMM normalization^18^. Genes were transformed using, and genes with mean normalized log count < were excluded.

### Expression-based subtyping

PDCs were assigned to proneural, classical, or mesenchymal subtypes using single-sample GSEA (ssGSEA) with the Wang et al. glioma-intrinsic subtype signatures ^19^ (**Supplementary Table ST2**). For each subtype signature, ssGSEA scores were Z-transformed across HGCC cultures, and each culture was assigned the subtype with the highest Z-transformed score.

**ssGSEA.** Enrichment scores were computed using gseapy 0.10. Neftel et al. cellular-state meta-modules (MES1, MES2, AC, OPC, NPC1, NPC2, G1/S, and G2/M) were scored using the same ssGSEA framework^20^ (**Supplementary Table ST2**). Scores were regressed against phenotypic components using linear regression (statsmodels 0.12). FDR was controlled using Benjamini–Hochberg within each gene set collection (neural cell type, GBM subtype, chromosome location) at.

### Proteomics (sample preparation and LC-MS/MS)

Mass-spectrometry-based proteomics, including cell lysis, in-solution digestion, TMT labeling, high-resolution isoelectric focusing, LC–MS/MS acquisition, and protein identification, were performed as described in the Supplementary Methods.

### Multi-omic factor analysis

Multi-Omics Factor Analysis (MOFA)^21^ was performed using MOFA2 v1.12.1 in R v4.3.3. The input comprised preprocessed matrices of ssGSEA scores, DNA methylation, RNA-seq, and proteomic data across 79 HGCC model-level entries, leveraging MOFA’s tolerance for missing data and inclusion of partially overlapping multi-omic measurements. Because MOFA integrates partially overlapping modalities, its input set differs from the RNA-seq cohort and from the histopathologically scored subsets. Missing values were handled using MOFA’s built-in probabilistic framework. Samples were stratified by Factor 1 score, and survival associations were evaluated using Kaplan–Meier curves (survival v3.7-0; survminer v0.5.0) and Cox proportional hazards models. MOFA feature weights are provided in Supplementary Table ST3. Additional methods are provided in the Supplement.

### Single-cell Factor 1 and invasivity scoring

To compare Factor 1 with an independent single-cell invasion state, we analyzed the Yu et al. GBM single-cell RNA-seq dataset^22^ (GSE117891) after filtering non-GBM and IDH1-mutant tumors and retaining malignant cells. Invasivity scores were computed from the invasion pseudotime gene signatures described in the glioblastoma hijacking study^9^: Seurat AddModuleScore was applied separately to 19 invasion-correlated and 141 invasion-anticorrelated genes, and the anticorrelated score was subtracted from the correlated score. To isolate within-tumor variation, each cell’s invasivity score was centered by subtracting the mean score of all malignant cells from the same tumor. Factor 1 scores were obtained by projecting MOFA Factor 1 loadings onto gene-wise z-scored single-cell expression values for overlapping genes. The relationship between within-tumor-centered invasivity and Factor 1 was estimated using a Bayesian hierarchical random-slope model with patient-specific intercepts and slopes.

### In situ sequencing

In situ sequencing^23^ was performed on a single coronal tissue section from a U3013MG xenograft using a custom 150-gene panel targeting key GBM cell-state markers across proneural, mesenchymal, and microenvironmental programs (Supplementary Table ST11). Spatial Factor 1 scores were mapped across tissue bins by least-squares fitting of the Factor 1 MOFA loadings to the in situ expression values for overlapping panel genes. Tumor cell density was visualised by summing all human-specific reads per spatial bin.

### TargetTranslator analysis

We applied TargetTranslator^24^ with modifications. For each phenotypic trait, RNA signatures were computed as gene–trait correlations. These signatures were matched to 19,763 LINCS drug-induced transcriptional profiles to produce a score matrix. Singular value decomposition () was used to account for collinearity. Drugs were ranked by the first two principal components of: PC1 (tumor initiation/malignancy) and PC2 (diffuse vs. perivascular invasion). Drug targets were annotated using the Drug Repurposing Hub and tested for enrichment along PC1 and PC2 using Kolmogorov–Smirnov tests (). Only significant targets are shown in Figure 6. Further details are provided in the Supplementary Methods.

### DepMap/PRISM phenotype-signature bridge

To compare HGCC-derived phenotype programs with public pharmacogenomic data, we generated signed RNA signatures for each of the ten xenograft phenotypes by computing Pearson correlations between phenotype scores and HGCC RNA-seq expression across models. Each signature was projected into GBM IDH-wildtype DepMap cell lines by z-scoring expression per gene and calculating a weighted mean using the HGCC gene–phenotype correlation coefficients. Projected phenotype scores were then correlated with PRISM Repurposing log-fold-change values across overlapping GBM lines. Negative correlations indicate that higher projected phenotype score is associated with stronger drug sensitivity, because lower PRISM log-fold-change denotes greater viability loss.

### 2D live-cell proliferation assays

Based on TargetTranslator predictions, ten compounds predicted to inhibit perivascular/condensed growth and eleven predicted to inhibit diffuse growth were tested in U3013MG and U3180MG. An 11-point dilution series was generated for each compound, and cell growth was monitored over 3 days using an IncuCyte S3 live-cell imaging system (Sartorius). Group differences in the response values plotted in Figure 6D were assessed using two-sided Welch’s t-tests across compounds (Group 1, 10; Group 2, 11); compound-level response summaries are provided in Supplementary Table ST4. See Supplementary Methods.

### Sphere invasion assay

Spheroids were generated by seeding 1,000 PDCs per well in PrimeSurface 384-well ultra-low attachment plates (S-BIO #MS-9384UZ). Matrigel and eight candidate compounds were added at 4–6 concentrations (Supplementary Table ST5). Images of spheroid cores and invasive fronts were acquired every 6 hours for seven days using the IncuCyte S3 spheroid module (10 phase/brightfield). Details are in the Supplement.

### Brain slice assay

Ex vivo brain slice cultures were prepared as previously described ^25^. Briefly, brains from xenograft-bearing mice were embedded in low-melting-point agarose and sectioned into 300 μm slices using a Leica VT1200 S vibratome. Slices were transferred to transwell membranes (Corning #3460), and vasculature was visualized with 2 μg/mL Tomato lectin–DyLight 594. Slices were treated with DMSO (0.3%), PIK-75 (0.5–4 μM) or Buparlisib (1–8 μM) and imaged every 0.5–1.7 hours for up to six days using an ImageXpress Micro Confocal system (Molecular Devices). Additional methods are provided in the Supplement.

## Results

### A standardized PDCX cohort with blinded histopathological scoring enables quantitative invasion phenotyping

A major barrier to understanding glioblastoma (GBM) invasion is the lack of scalable, *in vivo* systems that combine reproducible phenotyping with matched multi-omic profiles. To address this, we established the HGCC Phenobank, an orthotopic xenograft cohort designed to capture patient-to-patient variation in GBM growth and invasion under standardized conditions.

The full in vivo resource comprised 449 mice representing 76 HGCC lines. For the core histopathological analyses, we focused on the subset with aggregate mouse tumor-initiation metrics and endpoint pathology available after sample-identity harmonization. Fifty-eight HGCC lines contributed to the aggregate mouse tumor-initiation table, 50 produced detectable tumor cells at endpoint within a 40-week observation window (hereafter “tumor-bearing”; Supplementary Table ST1), and 37 generated tumors of sufficient size and quality for full invasion phenotyping (hereafter “fully scored”). At the endpoint, blinded observers scored ten prespecified histopathological traits (including perivascular co-option, white-matter infiltration, perineuronal satellitosis, and diffuse parenchymal spread) on human-specific NuMA– and STEM121-stained tissue sections (Figure 1A–B). A hierarchical statistical model was used to derive PDC-specific phenotype estimates while accounting for observer and batch effects (Methods).

**Figure 1.**
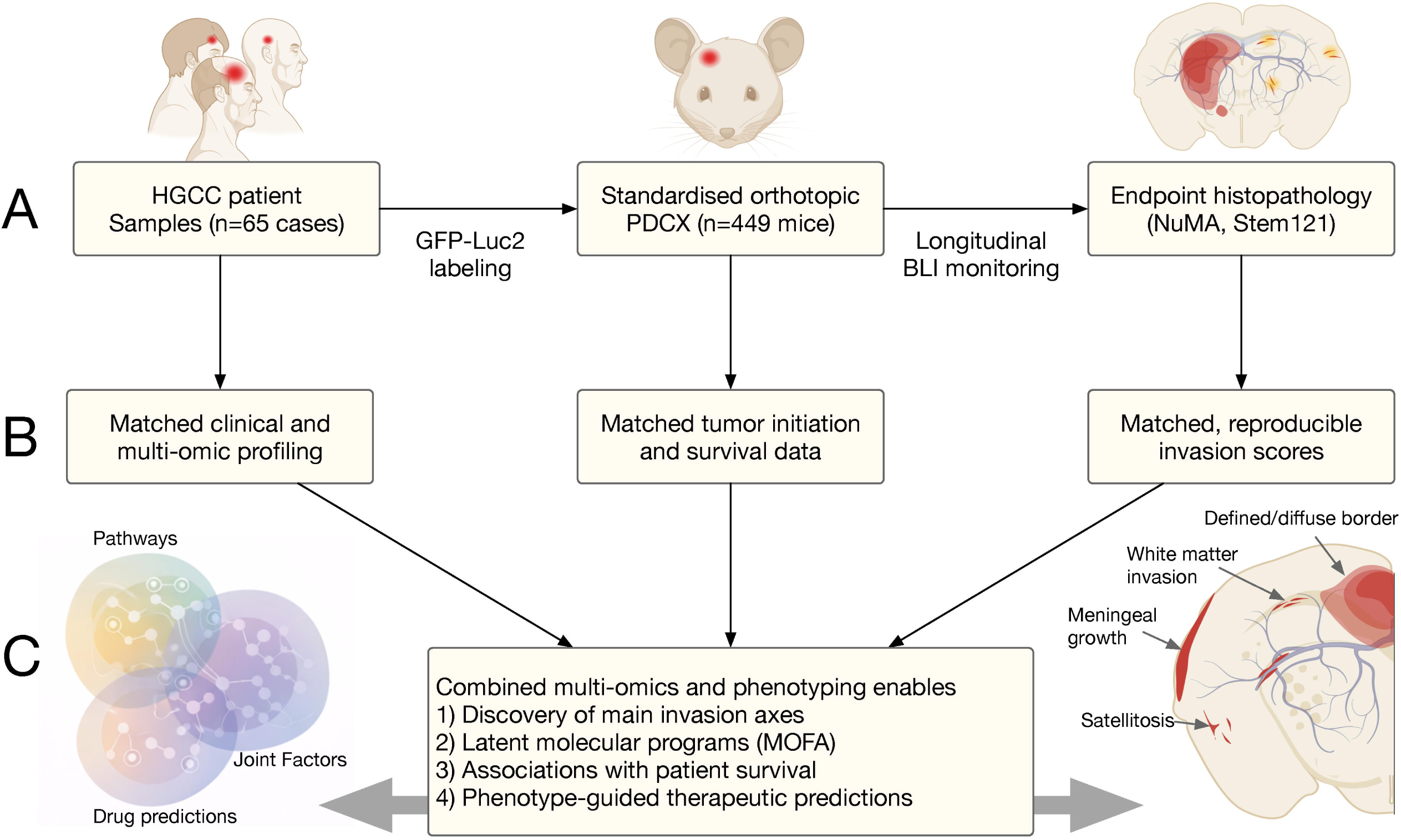
Establishment of the HGCC Phenobank and integrated profiling pipeline. (A) Overview of the HGCC Phenobank workflow. The *in vivo* resource comprised 449 orthotopic xenografts representing 76 HGCC lines. The RNA-profiled molecular cohort comprised 65 unique GBM stem-like cultures with matched multi-omic profiling. Tumor development was monitored longitudinally by bioluminescence imaging, followed by endpoint histopathology (NuMA, STEM121). (B) Each model was paired with matched multi-omic profiling (RNA-seq, DNA methylation, TMT proteomics, ATAC-seq, ssGSEA) and reproducible, blinded scoring of ten invasion traits, including diffuse parenchymal spread, white matter invasion, and perivascular co-option. (C) Integration of multi-omics with *in vivo* phenotyping enables discovery of dominant invasion axes, identification of latent molecular programs (MOFA), mapping of clinically relevant variation, and generation of phenotype-guided therapeutic predictions.

These standardized measurements revealed substantial and reproducible differences in both tumor initiation and invasion behavior between patient-derived models. Importantly, mouse survival was modestly correlated with survival in the corresponding patients (Pearson 0.1832, 0.045; Supplementary Figure SF1B), highlighting the translational relevance of the cohort.

To relate these phenotypes to underlying molecular features, each PDC was paired with RNA-seq, DNA methylation, and mass-spectrometry-based proteomic profiles generated from the corresponding HGCC culture (Figure 1B). This integrated, patient-aligned dataset provided the foundation for linking *in vivo* growth behaviors to multi-omic states and for identifying latent molecular programs associated with GBM invasion.

### Neurodevelopmental programs associate with engraftment and align with patient survival

Tumor engraftment varied broadly across the 50 tumor-bearing HGCC cultures, with some models generating rapidly progressive tumors and others showing delayed or incomplete engraftment (**Figure 2A**). This variation was highly reproducible across mice injected with the same culture, consistent with engraftment potential being an intrinsic property of each model.

**Figure 2.**
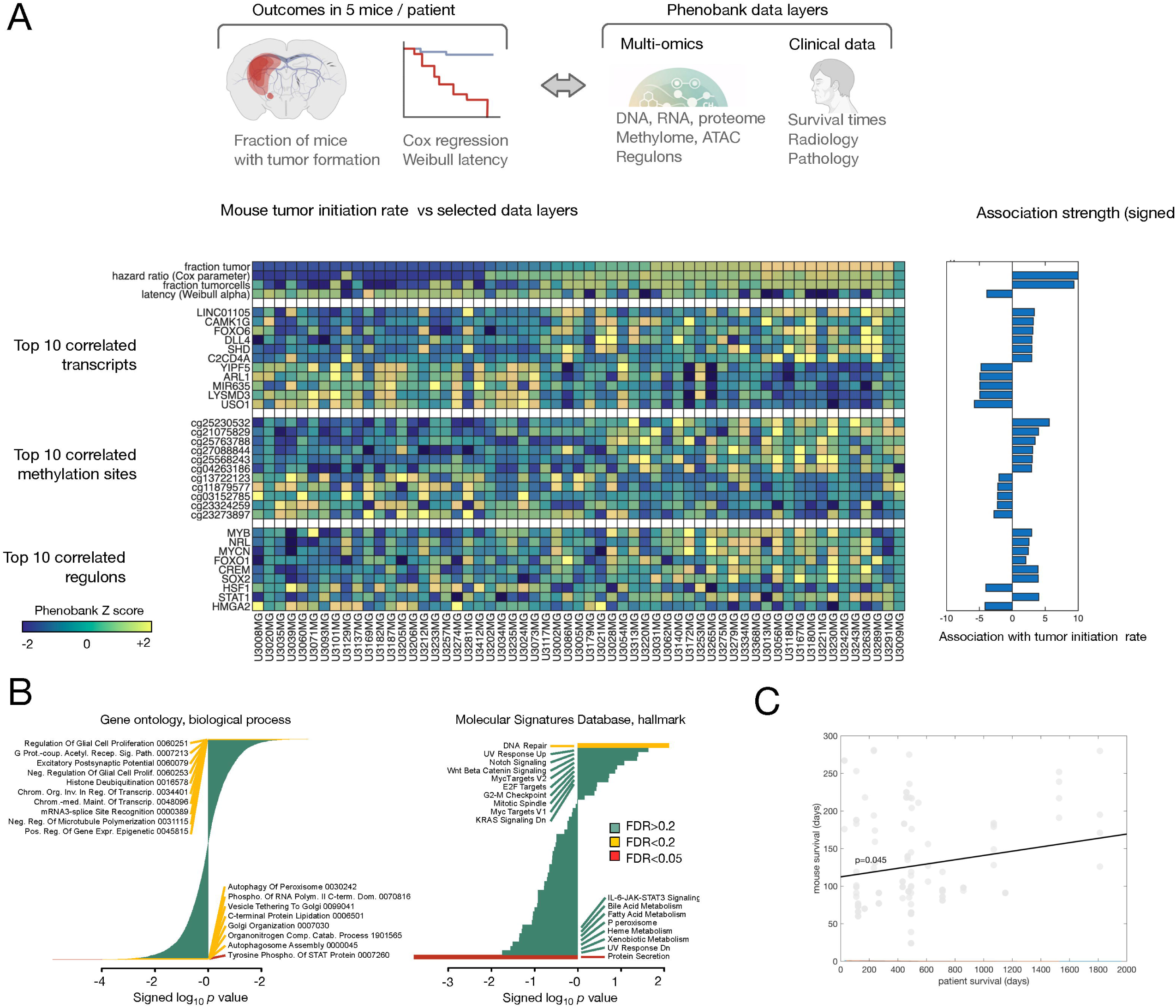
Tumor engraftment associates with neurodevelopmental and stem-like molecular programs. (A) Heatmap showing tumor engraftment frequency in HGCC Phenobank cases, sorted from low (left) to high (right). Epigenetic and transcriptional features linked to tumorigenicity include elevated expression of C2CD4A and LINC01105, and methylation of *TACR3* (cg04263186) and *HTR1B* (cg25763788). (B) ssGSEA-based pathway enrichment scores correlated with tumor engraftment frequency across PDCs, revealing positive associations with NOTCH, Wnt, MYCN, and DNA repair pathways, and negative associations with JAK–STAT signaling and autophagy.

In line with previous characterization of GBM PDCX models^7^, tumors derived from proneural PDCs were associated with shorter survival in mice, while mesenchymal cultures showed less aggressive growth. This pattern likely reflects the strong contribution of tumor-immune interactions (absent in immunodeficient hosts) to mesenchymal GBM aggressiveness.

To explore the pathway and process correlates of tumor engraftment, we examined ssGSEA enrichment scores for curated pathway gene sets and regressed these against tumor-engraftment frequencies. Cultures with high engraftment potential were enriched for neurodevelopmental and proliferative pathways including Notch, Wnt/β-catenin, MYCN-associated transcriptional programs, DNA repair, and synaptic signaling modules (**Figure 2A–B**). Prominent individual markers included *C2CD4A* (NLF1), implicated in calcium-dependent vesicle trafficking and vascular interactions, and the long non-coding RNA *LINC01105*/*SILC1*, a regulator of *SOX11*. High-engraftment cultures also showed increased methylation at *TACR3* (cg04263186) and *HTR1B* (cg25763788), encoding neuroreceptors associated with excitatory signaling. In contrast, low-engraftment cultures were enriched for autophagy, catabolic processes, and JAK–STAT signaling pathways, which may reflect reduced proliferative competence or compensatory stress responses. The significance of these associations was supported by FDR correction; 154 mRNA transcripts were associated with tumor engraftment at FDR 10%, with additional significant associations across methylation, regulon, and pathway data layers (**Figure 2A**).

Initiation-associated survival in xenografted mice modestly but significantly correlated with the survival of the corresponding patients (Pearson 0.1832, 0.045; **Supplementary Figure SF1B**), providing an external clinical anchor for the PDCX-derived traits.

### Two orthogonal invasion axes emerge from standardized phenotyping and are reproducible across mice

Having established molecular correlates of tumor initiation, we next examined whether the histological invasion patterns observed across the HGCC Phenobank were similarly structured. Blinded scoring of ten invasion traits revealed extensive patient-to-patient variation, but within each PDCX model, growth patterns were remarkably consistent across mice. A spectrum of phenotypes was observed including diffuse parenchymal infiltration, white-matter tract invasion, perivascular co-option, and condensed, margin-forming growth (**Figure 3A**).

**Figure 3.**
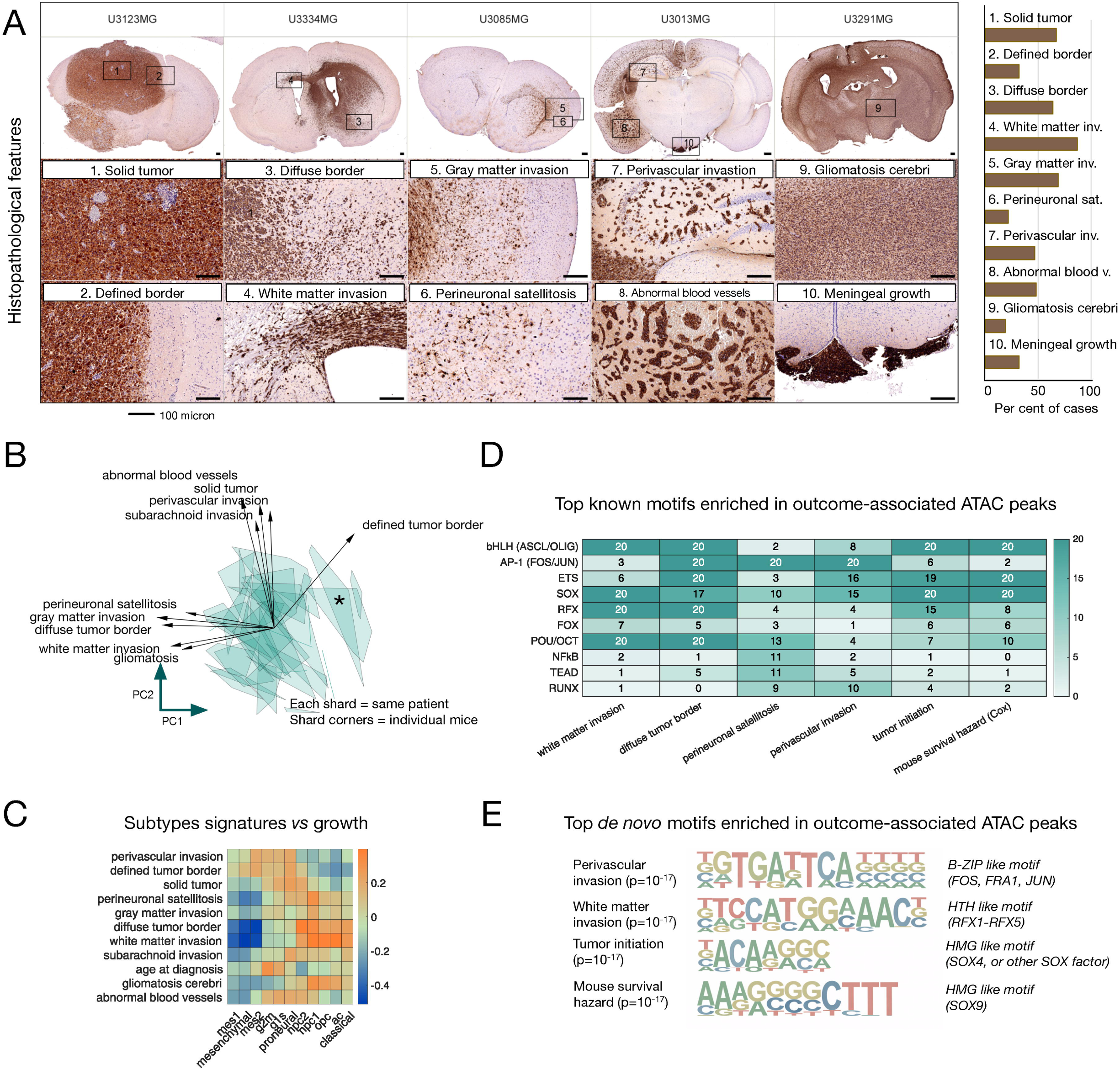
Histological growth patterns define phenotypic axes linked to transcriptional and epigenetic programs. (A) Immunohistochemical images showing xenografts stained for the human cell-specific marker (STEM121). Key phenotypic traits such as perivascular growth, white matter invasion, and diffuse spread are depicted, with a summary of the fraction of cases scored positive for each trait. (B) PCA of morphological scores across 37 fully scored models reveals two major axes: diffuse invasion (PC1) and perivascular/condensed growth (PC2), with gliosarcoma-like cases forming a distinct cluster (*). (C) ssGSEA subtype scores correlated with phenotypes show that mesenchymal tumors align with perivascular invasion (PC2), while proneural/OPC tumors align with diffuse growth (PC1). (D) Known transcription factor motif enrichment analysis from ATAC-seq peaks shows divergence between invasion types and outcome-associated traits. (E) De novo motif enrichment analysis highlights SOX/OLIG-associated motifs for diffuse invasion and AP-1/ETS-associated motifs for perivascular/condensed growth.

To identify major axes of invasion behavior, we performed principal component analysis (PCA) on the ten phenotypic scores across the 37 fully scored models. Two dominant, orthogonal components accounted for the majority of variance (**Figure 3B**). PC1 captured diffuse, parenchyma-infiltrating traits (white-matter invasion, perineuronal satellitosis, and infiltrative edges), whereas PC2 aligned with perivascular and condensed growth patterns. A small number of cases formed a distinct cluster with sarcomatous or spindle-cell histology, consistent with variants similar to gliosarcoma. These phenotypic axes were robust: inter-mouse variability within each PDCX was markedly smaller than inter-model variability (Supplementary Figure SF2B), indicating that invasion mode is an intrinsic, lineage-encoded property rather than a stochastic response to the host environment.

Integration with transcriptomic subtype classifications further supported the biological interpretation of these axes. Proneural/OPC-like models were enriched at the diffuse end of PC1, while mesenchymal-like models showed higher PC2 scores and more pronounced perivascular growth (**Figure 3C**). This aligns with prior evidence that OPC-like states promote infiltrative behavior, whereas mesenchymal programs are associated with vascular remodeling and compact growth.

To investigate underlying regulatory architecture, we next examined transcription factor (TF) motif enrichments in ATAC-seq peaks correlated with PC1 and PC2. Diffuse invasion (PC1) was associated with bHLH lineage regulators including *ASCL1*, *OLIG1/2*, and RFX-family motifs, consistent with neural progenitor-like identity (**Figure 3D,E**). In contrast, perivascular/condensed growth (PC2) showed enrichment for AP-1 (FOS/JUN), ETS, and SOX motifs, which are implicated in vascular interaction, stress response, and extracellular matrix remodeling. Among models with high tumor-propagating potential, SOX, ETS, and TAL1 motifs were also enriched, correlating with reduced mouse survival (**Figure 3D,E**). Full HOMER known and de novo motif outputs are provided in Supplementary Tables ST6 and ST7. Together, these findings demonstrate that the phenotypic axes captured by blinded scoring are underpinned by distinct regulatory programs, providing a mechanistic foundation for latent-factor modeling in the subsequent analyses.

The reproducibility of these invasion modes across mice, their alignment with molecular subtypes, and their association with coherent TF programs suggested that a smaller number of latent molecular programs might integrate across omics layers to shape GBM growth behavior. To identify such cross-modal programs, we next applied Multi-Omic Factor Analysis (MOFA) to the HGCC Phenobank dataset.

### MOFA identifies 15 latent factors that unify multi-omic heterogeneity with growth phenotypes and prognosis

The transcriptional and epigenetic programs associated with diffuse versus perivascular invasion (Figure 3) suggested that a smaller number of coordinated molecular axes might structure GBM behavior across omics layers. To test this, we applied Multi-Omic Factor Analysis (MOFA) to RNA expression, DNA methylation, pathway activity scores, and proteomic profiles across 79 HGCC models, leveraging MOFA’s tolerance for missing data to maximize the integrated sample size across modalities. MOFA learns latent variables that capture shared sources of variation across modalities, enabling us to identify molecular programs that integrate transcriptomic, epigenomic, and protein-level signals.

The resulting model contained 15 factors, each explaining distinct proportions of variance across omics layers (**Figure 4A**). Among these, **Factor 1** emerged as the dominant, cross-modal axis, with strong loadings from RNA, methylation, and proteomic features. Its top molecular contributors included the neurodevelopmental regulators *ASCL1*, *OLIG1/2*, and associated bHLH lineage programs (**Figure 4B**), consistent with the regulatory architecture underlying diffuse invasion (Figure 3D,E).

**Figure 4.**
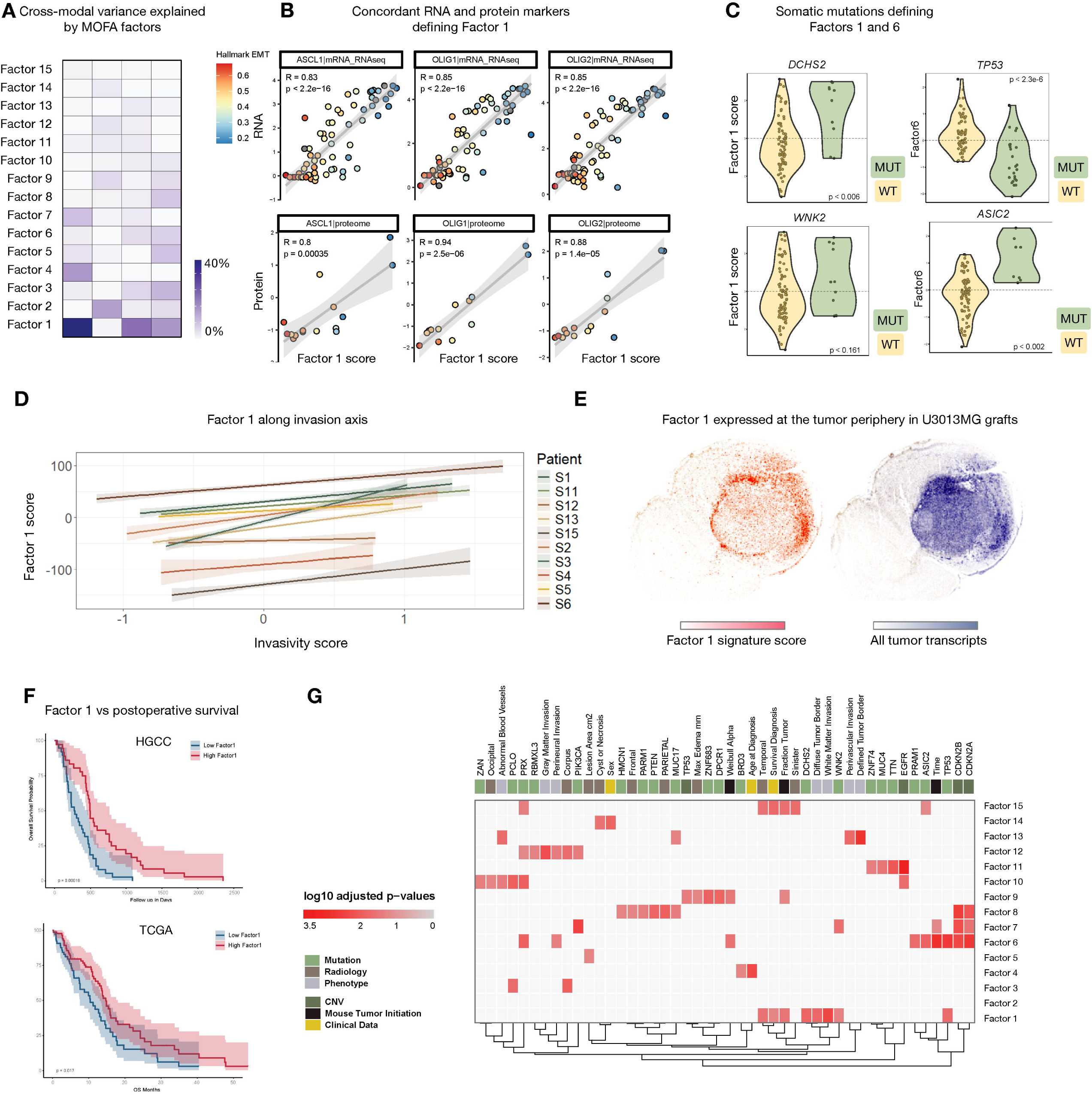
MOFA reveals latent factors that bridge phenotype, prognosis, and molecular state. MOFA was fit across 79 model-level entries with partially overlapping omic measurements. (A) Variance explained across omics layers by the top 15 MOFA factors, with Factor 1 showing the strongest cross-modal signal. (B) Concordant RNA and protein markers defining Factor 1: scatter plots show expression of ASCL1, OLIG1, and OLIG2 at mRNA (top) and protein (bottom) levels against Factor 1 scores, coloured by Hallmark EMT score. Pearson R and p-values are indicated. (C) Somatic mutations associated with Factors 1 and 6: violin plots show Factor scores stratified by mutational status for DCHS2 and WNK2 (Factor 1) and TP53 and ASIC2 (Factor 6). (D) Factor 1 score increases along a within-tumor-centered single-cell invasivity axis in the Yu et al. GBM dataset. Invasivity was scored from published invasion pseudotime signatures by subtracting an invasion-anticorrelated module score from an invasion-correlated module score and centering the result within each tumor. (E) Representative in situ sequencing map from a U3013MG xenograft section showing high Factor 1 signature score at the tumor periphery and all human tumor transcripts marking the tumor mass. (F) Stratification by Factor 1 score shows significant survival differences in HGCC PDCX mice and TCGA GBM patients. (G) Heatmap of associations (log10 adjusted p-values) between MOFA factors and phenotypic, clinical, radiology, mutation, CNV, and mouse tumor-engraftment variables. Factor 1 is linked to diffuse invasion and tumor initiation; additional factors capture diverse molecular–clinical combinations including mutation context and clinical features (see text).

Factor 1 was significantly associated with multiple tumor traits: higher tumor engraftment frequency in mice, increased diffuse parenchymal invasion, and reduced mouse survival (**Figure 4F,G**). In addition, Factor 1 showed enrichment for recurrent alterations in *TP53*, *DCHS2*, and *WNK2*, supporting a link between lineage programs and genomic context. When projected onto TCGA GBM transcriptomes, the Factor 1 signature stratified patient survival (**Figure 4F**), establishing its clinical relevance beyond the HGCC cohort.

Beyond Factor 1, the remaining 14 factors captured diverse combinations of molecular and clinical variables (**Figure 4G**). Factor 6, for example, was associated with mutations in *TP53*, *PRX*, *PRAM1*, and *ASIC2*, and with the Weibull survival shape parameter, suggesting a subgroup with a distinct mutational landscape and hazard dynamics. Other factors, such as Factors 8 and 9, showed associations with *PTEN* copy-number variation and radiological features including frontal placement and edema extent. These associations are hypothesis-generating and illustrate the breadth of molecular–clinical variation captured by the full factor model.

These latent factors provide a compact representation of GBM heterogeneity that bridges molecular state, growth phenotype, and clinical outcome, and form the basis for the spatial and therapeutic analyses that follow.

### Single-cell and spatial mapping link Factor 1 to invasive states

To evaluate whether Factor 1 corresponds to an invasive cell state, we first projected the MOFA Factor 1 loadings into an independent GBM single-cell RNA-seq dataset and compared the resulting cell-level Factor 1 scores with a within-tumor-centered invasivity score derived from published invasion pseudotime signatures (Methods). Factor 1 was positively associated with invasivity within tumors (population-level slope 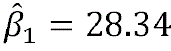, 95% CI 13.45,43.25), although the strength of this coupling varied substantially between patients (between-patient slope SD 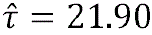 CI 12.42,38.24) (**Figure 4D**; **Supplementary Figure SF7**).

We then projected Factor 1 onto in situ sequencing (ISS; ^23^) data from a coronal section of a U3013MG xenograft (150-gene panel; Methods). Factor 1 signal was enriched at the tumor periphery relative to the tumor core (**Figure 4E**), consistent with Factor 1 marking cells at the active invasion front, where ASCL1/OLIG-driven progenitor programs enable engagement with the brain parenchyma.

These results demonstrate that the latent molecular program identified by MOFA from bulk mul-ti-omic data maps to a spatially distinct, functionally interpretable cell state within tumors. The leading-edge localization of Factor 1 reinforces its mechanistic link to infiltrative behavior and supports phenotype-resolved therapeutic strategies targeting cells at the invasion front.

### Cross-modal coherence of invasion programs revealed by RNA–protein consensus markers

The drug-prediction analysis that follows relies on RNA signatures as proxies for invasion-associated molecular state. For this to be valid, the transcriptional programs linked to invasion phenotypes must be genuinely reflected at the protein level. We therefore asked whether the invasion programs identified by MOFA show consistent signatures across both RNA and protein expression.

Across the HGCC Phenobank, RNA–protein correlations were generally high (median 0.63), with stronger concordance observed among moderately to highly expressed gene products (**Figure 5A**). Importantly, genes whose RNA levels correlated with diffuse or perivascular invasion showed parallel correlations at the protein level, indicating that the phenotypic axes identified earlier reflect coordinated multi-layer regulation rather than transcription-only variation (**Figure 5B**).

**Figure 5.**
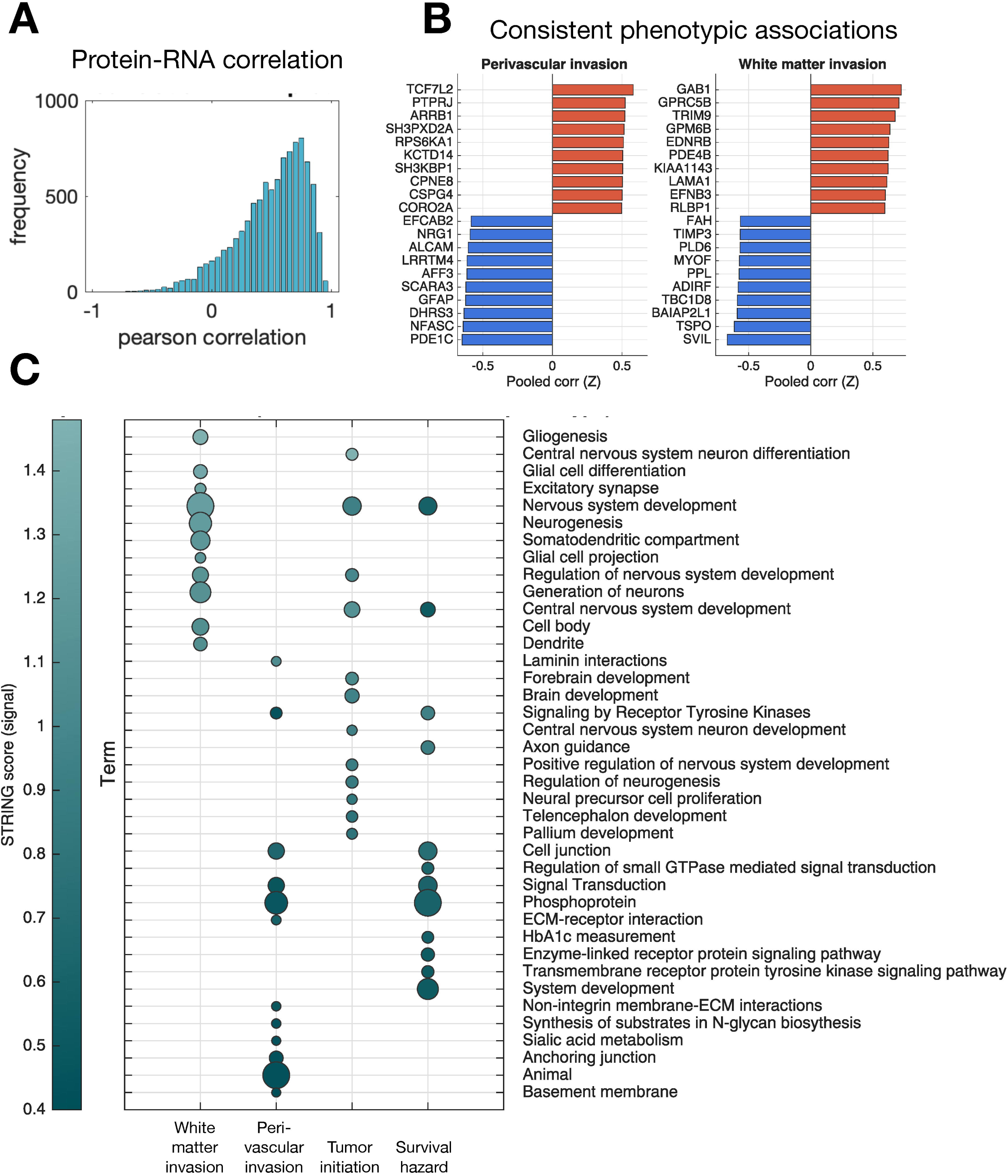
Consensus RNA-protein biomarkers reveal pathways governing GBM growth and survival phenotypes. (A) Global RNA-protein correlation across gene products in HGCC models (left) and dependency of RNA-protein correlation on expression level (right). (B) Phenotype-specific correlation for representative gene products shows alignment of RNA and protein associations. (C) Pathway enrichment scores computed using STRING for sets of RNAs/proteins jointly correlated with white matter invasion, perivascular invasion, survival hazard, and tumor initiation.

To summarize these patterns in a biologically interpretable form, we combined RNA and protein correlation coefficients using Fisher’s Z-transformation and ranked gene products by their consensus association with each phenotype. Pathway analysis of the top-ranked genes revealed enrichment of neurodevelopmental, OPC-like, and DNA-repair programs for diffuse invasion, and AP-1/ETS-associated stress and matrix–interaction pathways for perivascular/condensed growth (**Figure 5C**; Supplementary Table ST8). These pathway-level enrichments closely mirror the regulatory programs captured by the MOFA factor model, reinforcing that the latent factors identified earlier reflect stable, cross-modal biological states.

Together, these consensus RNA–protein markers provide orthogonal validation that the invasion programs observed in vivo correspond to coherent molecular pathways observable at both the transcript and protein levels. This cross-layer coherence supports the use of RNA signatures for phenotype-guided therapeutic prediction in the subsequent analyses.

### Latent invasion programs nominate phenotype-specific drug vulnerabilities

Having established that GBM invasion is structured by reproducible phenotypic axes, encoded by cross-modal molecular programs, and deployed in spatially distinct tumor regions, we next asked whether these molecular programs could be translated into actionable therapeutic predictions. Specifically, we examined whether the transcriptional states underlying diffuse versus perivascular/condensed invasion, as well as the neurodevelopmental Factor 1 program associated with initiation and poor prognosis, could identify compounds capable of selectively suppressing these growth modes.

To accomplish this, we applied TargetTranslator, which matches disease-specific RNA signatures to drug-induced transcriptional responses from the LINCS L1000 compendium. Using RNA signatures for each phenotypic trait and MOFA factor, we computed perturbation scores for 19,763 compounds (**Figure 6A**). Negative scores indicated predicted suppression of a phenotype-linked signature, while positive scores indicated reinforcement.

Principal component analysis of drug perturbation scores revealed two dominant axes (**Supplementary Figure SF4**). The first (drug-PC1) reflected global malignancy, capturing tumor engraftment and survival hazard, consistent with its alignment with Factor 1. The second (drug-PC2) separated compounds predicted to inhibit diffuse invasion from those predicted to inhibit perivascular/condensed growth, mirroring the two histological modes defined earlier. Thus, the same invasion axes observed at the phenotypic, transcriptional, proteomic, and spatial levels are recapitulated in the therapeutic prediction space.

Drug target enrichment along these axes yielded mechanistic insight (**Figure 6C**). Compounds predicted to suppress diffuse invasion were enriched for modulators of histamine and prostaglandin signaling, while inhibitors of PI3K, MAPK, and CDK pathways were selectively associated with suppression of perivascular/condensed growth. Targets linked to proteostasis (proteasome, HSPs) were associated with reduced tumor initiation and improved survival (**Figure 6B**), consistent with the dependence of highly proliferative Factor 1-high states on protein homeostasis.

**Figure 6.**
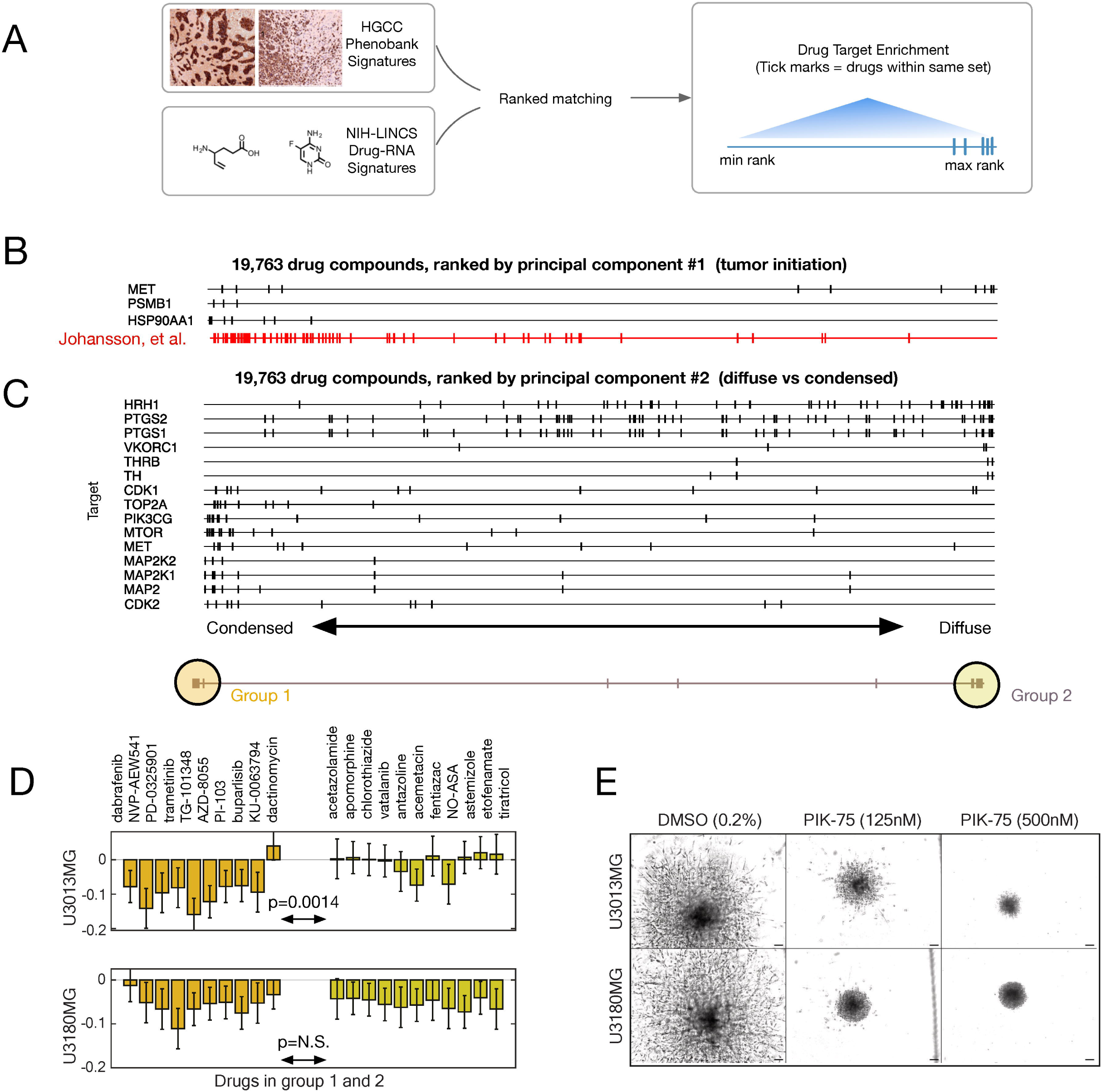
Computational repurposing and validation identifies selective inhibitors of invasion phenotypes. (A) Schematic illustration of the TargetTranslator workflow, linking HGCC-derived RNA signatures to drug perturbation profiles from the L1000 dataset. (B) Drug target enrichment ranked by PC1 (tumor initiation and survival). Tick marks indicate positions of drugs targeting the indicated gene products among 19,763 ranked compounds; targets enriched among top-ranked compounds include MET, PSMB1, and HSP90AA1. Positions of drugs validated in an independent screen (Johansson et al., red) confirm enrichment at the high-malignancy end of PC1. (C) Drug target enrichment ranked by PC2 (diffuse vs condensed invasion), defining Group 1 (condensed-suppressing, left) and Group 2 (diffuse-suppressing, right). Targets enriched in Group 1 include CDK1, CDK2, PIK3CG, and MAPK-pathway kinases; Group 2 is enriched for histamine (HRH1) and prostaglandin (PTGS1/2) targets. (D) Live-cell assay comparing Group 1 and Group 2 compounds in U3013MG (perivascular/condensed) and U3180MG (diffuse) models. Group 1 compounds showed stronger inhibition of U3013MG than Group 2 compounds (two-sided Welch’s *t*-test, 0.0014), whereas the corresponding comparison in U3180MG was not significant. (E) Sphere invasion assay demonstrates that PIK-75 reduces both proliferation and invasion of U3013MG and U3180MG. Scale bar indicates 100 μm.

To functionally validate phenotype-specific predictions, we performed a focused drug screen on two representative models: U3013MG (perivascular/condensed) and U3180MG (diffuse). Twenty-one compounds, pre-selected based on their predicted selectivity for each invasion mode, were tested across 11-point dose ranges. Group 1 compounds showed stronger inhibition of the perivascular model U3013MG than Group 2 compounds (mean response, 8.84 vs. 1.09; two-sided Welch’s *t*-test, *p* 0.0014), whereas the corresponding comparison in the diffuse model U3180MG was not significant (–5.73 vs. –5.41; *p* = 0.73; **Figure 6D**). These results support phenotype-associated differences in drug sensitivity while indicating that the strongest group-level separation in this assay was observed in U3013MG.

As orthogonal support, we asked whether the same xenograft-derived RNA phenotype signatures could be mapped into public GBM pharmacogenomic data. Projection of the HGCC signatures into GBM IDH-wildtype DepMap lines followed by correlation with PRISM drug-response profiles showed that PIK-75 sensitivity was associated with vascular and perivascular/condensed xenograft programs, including perivascular invasion (Pearson *r* 0.41, *p* 0.049) and abnormal blood-vessel signatures (*r* =-0.45, *p* = 0.031; **Supplementary Figure SF5**). By contrast, the PI3K inhibitor buparlisib showed weaker associations across these xenograft signatures. This external bridge supports the prioritization of PIK-75 as a phenotype-linked vulnerability while remaining consistent with its subsequent validation in invasion assays.

We next examined whether compounds could reduce invasion dynamics in 3D contexts. A sphere invasion assay performed over seven days (**Supplementary Figure SF3**) showed that several predicted compounds suppressed radial invasion, with one multi-target inhibitor, PIK-75, emerging as the most broadly effective. PIK-75, an inhibitor of PI3K, CDK1/2, and the transcription factor TAL1, reduced both proliferation and invasion in U3013MG and U3180MG across multiple doses (**Figure 6E**). These effects were consistent with the placement of PIK-75 along drug-PC2 and with TAL1/PI3K-axis enrichment in aggressive, Factor 1-high models.

Finally, in ex vivo brain-slice assays, PIK-75 significantly reduced tumor cell migration and velocity across three HGCC models (**Figure 7A,B,D**; **Supplementary Video 1**), confirming that its inhibitory effects extend to an organotypic environment that preserves vascular architecture, a key element of the perivascular phenotype. Buparlisib similarly reduced cell displacement across models (**Figure 7C,E**; **Supplementary Video 2**).

**Figure 7.**
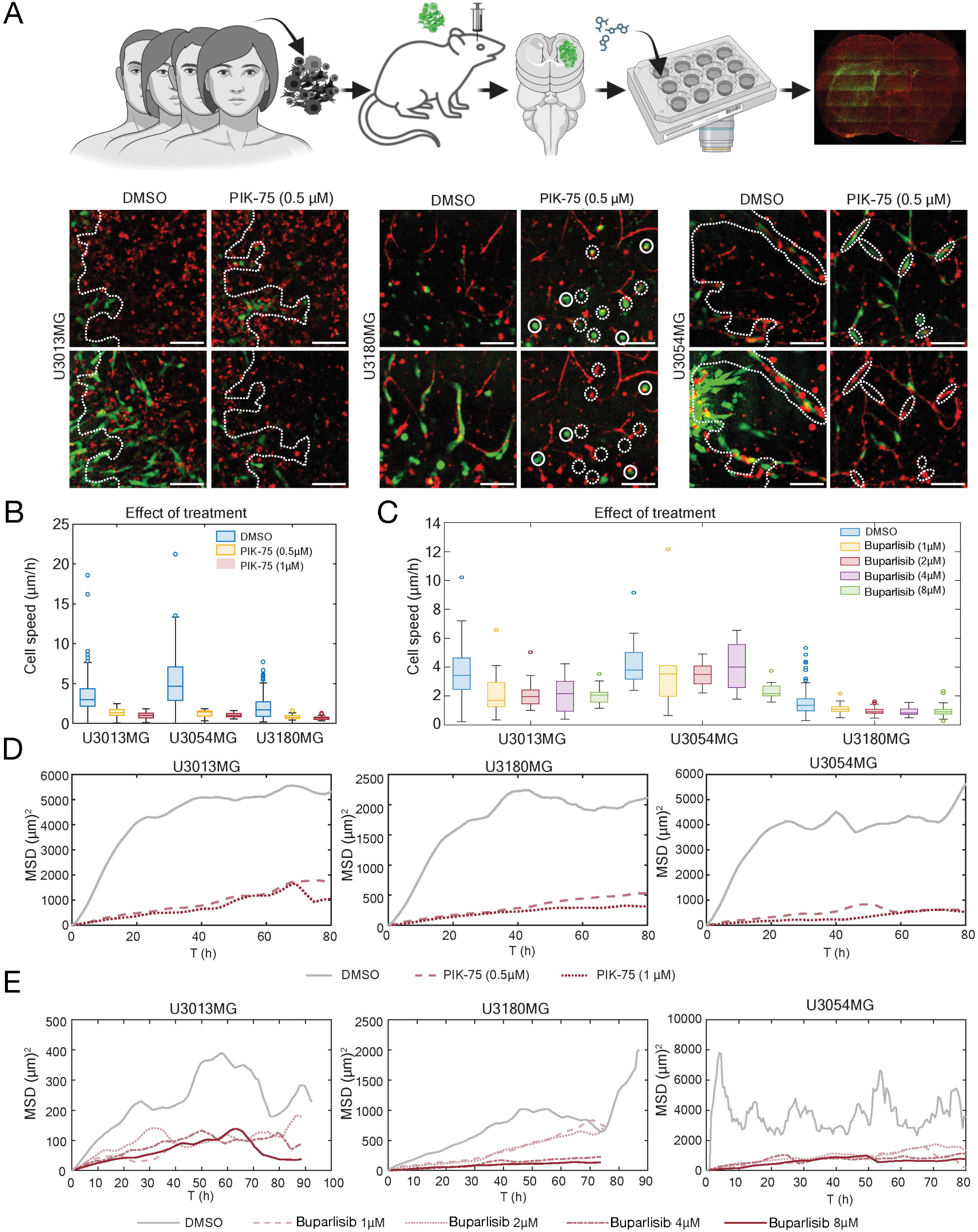
Ex vivo brain-slice validation of invasion inhibition. (A) Workflow for organotypic brain-slice assays (top) and representative confocal images of tumor cell spread (green, GFP) and vasculature (red, tomato lectin) after vehicle (DMSO) or PIK-75 (0.5 μM) treatment in U3013MG, U3180MG, and U3054MG. Scale bar indicates 100 μm. (B) Quantification of cell speed after PIK-75 treatment across three HGCC models. (C) Quantification of cell speed after Buparlisib treatment across three HGCC models. (D) Mean-squared displacement (MSD) over time following vehicle or PIK-75 treatment across three HGCC models. (E) Mean-squared displacement (MSD) over time following vehicle or Buparlisib treatment across three HGCC models. Representative time-lapse videos are provided for PIK-75 (Supplementary Video 1) and Buparlisib (Supplementary Video 2). *** indicates p 0.001.

Together, these results connect phenotype, molecular program, therapeutic prediction, and functional validation in a phenotype-anchored framework. The invasion programs uncovered by the HGCC Phenobank are not only descriptive but mechanistically actionable, revealing phenotype-linked drug vulnerabilities and identifying compounds such as PIK-75 that suppress invasion-associated behaviors across representative models.

## Discussion

The HGCC Phenobank addresses a longstanding gap: few resources systematically link *in vivo* invasion phenotypes to multi-omic programs across a patient-aligned panel. The findings reported here establish that GBM invasion is not simply heterogeneous but structured, and that this structure is molecularly encoded, reproducible, and therapeutically actionable.

A principal insight from this work is that GBM invasion resolves into a structured landscape anchored by two dominant modes: diffuse parenchymal infiltration and perivascular/condensed expansion. These behaviors were strikingly reproducible across mice engrafted with the same culture, emphasizing that invasion mode is a stable, lineage-encoded trait rather than a stochastic response to the host microenvironment. Moreover, these phenotypes aligned with transcriptional subtypes^19^ and reflected distinct regulatory architectures, recapitulating classical secondary Scherer structures^8,9^.

Latent-factor modeling provided a further unifying framework. Using MOFA^21^, we identified 15 molecular programs that organize transcriptomic, proteomic, and epigenomic variation across the cohort. Among these, Factor 1 emerged as the dominant cross-modal axis, enriched for ASCL1, OLIG1/2, and associated neural progenitor and glioma stemness programs^19,26–28^. Factor 1 linked increased tumor-engraftment capacity, diffuse invasion, and reduced survival in xenografted mice; projection of the Factor 1 signature onto TCGA GBM transcriptomes further stratified patient survival, underscoring its biological consistency and clinical relevance.

Importantly, in situ sequencing (ISS) of U3013MG xenograft sections provided spatial confirmation that Factor 1 reflects a bona fide *in vivo* transcriptional state. Factor 1 scores were elevated at the invasive periphery relative to the tumor core, forming a gradient that peaks at the tumor–brain interface. This spatial architecture is consistent with Factor 1 representing a leading-edge, progenitor-like program deployed by cells actively engaging the brain parenchyma rather than a uniform property of all tumor cells. These findings demonstrate that latent programs derived from bulk multi-omics are spatially organized within the tumor microenvironment, reinforcing their functional and mechanistic interpretation.

The between-patient variation in the invasivity–Factor 1 slope (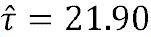) was large relative to the average effect (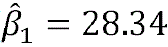), indicating that the coupling is not uniform across tumors but varies substantially between patients. This is consistent with the well-established transcriptional heterogeneity of glioblastoma, and identifying the molecular or clinical features that underlie this patient-to-patient variability represents a direction for future work.

These findings collectively support a model in which GBM behavior emerges from the interplay of lineage-specific programs and epigenetic remodeling rather than from genetic alterations alone. The enrichment of SOX, OLIG, and TAL1 motifs in aggressive cases aligns with prior work implicating chromatin regulation and developmental plasticity in glioma pathogenesis^26,29,30^. These motifs form part of a self-reinforcing PRC2-linked circuit that sustains stem-like, invasion-competent states while suppressing differentiation and adhesion programs such as neuronal maturation and E-cadherin expression^29–31^. MYCN, which is enriched in Factor 1-high, aggressive models and is synthetically lethal with CDK2 inhibition in neuroblastoma^32^, further highlights potential shared vulnerabilities between GBM and other MYCN-driven cancers.

By linking phenotypes to molecular programs, the HGCC Phenobank also enables rational therapeutic prediction. Using TargetTranslator^24^, we scored RNA signatures characteristic of diffuse versus perivascular invasion against drug-induced transcriptional profiles from LINCS. This analysis identified drug classes predicted to selectively suppress invasion modes or reduce malignancy-associated programs. Experimental validation in representative models confirmed phenotype-specific responses: diffuse and perivascular models exhibited preferential sensitivity to distinct compound groups in 2D proliferation assays, 3D sphere invasion assays, and ex vivo brain slices. Among these, PIK-75, a multi-target inhibitor of PI3K, CDK1/2, and TAL1, reduced invasion and proliferation across assays and models, consistent with its ability to disrupt the TAL1–MYCN–PI3K regulatory axis^33–36^. Its limited clinical viability^37,38^ underscores the need for improved derivatives but provides a mechanistic foothold for therapeutic development.

The HGCC Phenobank also reproduces clinically meaningful histopathological traits (including white matter infiltration, perivascular spread, and gliosarcoma-like morphology) that are difficult to model in vitro or using long-passaged cell lines^7,11,14^. The correlation between mouse and patient survival further demonstrates the translational value of these PDCX models. Beyond known drivers, our analyses highlight candidate genes such as DCHS2, PRX, C2CD4A, and TACR3 as potential regulators of GBM growth, vascular association, or neural signaling, suggesting new directions for mechanistic work.

As with all xenograft systems, important limitations remain. The absence of an intact immune microenvironment is particularly relevant for mesenchymal GBM, where macrophage and microglia infiltration shapes both the transcriptional state and the invasive behavior of tumor cells; these interactions cannot be recapitulated in PDCXs. This likely contributes to the weaker phenotypic concordance observed in mesenchymal lines and should be considered when interpreting subtype-specific findings. Incomplete modeling of spatial intratumoral heterogeneity and potential selection biases inherent to stem-like culture conditions are additional constraints. Future extensions incorporating humanized immune systems, organoid invasion models, or patient-matched spatial transcriptomics will help address these gaps. A further limitation is that our multi-omic profiles were derived from cell cultures prior to transplantation. Transcriptional and epigenetic states are likely to shift following engraftment and clonal selection in the mouse brain^39^, and additional programs relevant to invasion may only become apparent in sequenced tumor tissue. Future studies profiling extracted xenograft material at endpoint would complement the cell-culture-derived data presented here. In a matched single-cell RNA-seq pseudobulk comparison, parental culture and xenograft-derived mouse-brain samples retained substantial cell-line coherence, but xenograft-derived samples shifted along an *in vivo*-associated axis and showed greater dispersion (**Supplementary Figure SF8**), consistent with our previous observation that the brain microenvironment expands transcriptional-state diversity. Additionally, although our drug predictions were validated across multiple assays, full preclinical evaluation in pharmacokinetically optimized models is still required before clinical translation.

In summary, the HGCC Phenobank offers a next-generation resource for dissecting GBM heterogeneity in vivo. By anchoring molecular variation to reproducible phenotypic traits, latent regulatory programs, and therapeutic responses, it provides a coherent framework for understanding and targeting the biological drivers of GBM invasion. More broadly, this work illustrates how integrating multi-omics, spatial profiling, and phenotypic assays can provide a scalable blueprint for decoding the complexity of human tumors. HGCC Phenobank models are available to the research community; investigators interested in accessing cultures are encouraged to contact us via www.hgcc.se.

## Acknowledgements

We thank the glioblastoma patients donating tissue and the neurosurgery and neuropathology departments at the Uppsala Akademiska Hospital for their help in collecting and curating tumor samples and prof. Irina Alafuzoff at the Uppsala Akademiska Hospital for pathological evaluation. The authors gratefully acknowledge the HGCC biobank at Uppsala University for providing PDC cultures and Magnus Essand at Uppsala University for generously sharing the GFP-luciferase lentiviral vector with us. We also thank the FoU department at the Uppsala Akademiska Hospital and SciLifeLab Tissue Profiling Facility staff for their help with the digitalization of stained tissue slides. RNA sequencing was performed by the SNP&SEQ Technology Platform in Uppsala. The facility is part of the National Genomics Infrastructure (NGI) Sweden and Science for Life Laboratory. We used Grammarly and Claude (Anthropic) for grammar checks and language style and Codex (openAI) code code audit. The Science for Life Laboratory Drug Discovery and Development platform provided compounds from their drug libraries for the small drug screen. We thank prof. Monica Nister for commenting on the manuscript.

## Author statement

CK developed and benchmarked the mouse model and phenotyping experimental methods, designed and performed drug experiments, coordinated the study, and participated in data interpretation. SK collected mouse data, performed mouse experiments and phenotyping, and established PDCXs. ER developed the phenotype-scoring system, performed statistical analyses and effect-size estimation, and generated ssGSEA summaries used in downstream regressions. AS performed the MOFA analysis and contributed to Figure 4. HB performed mass-spectrometry analysis and delivered proteomics data; XC and MP contributed to proteomics data generation and interpretation. IU contributed substantially to scientific discussions and proposed the Yu et al. single-cell dataset analysis. FK developed the spatial-gradient single-cell analysis and the single-cell Bayesian regression analysis. HBM led the ex vivo brain-slice validation and Figure 7. MS ran the GlioTrace pipeline and summarized the brain-slice analysis results. IL and RE performed transcriptomics. LE performed cell work, including HGCC labeling. RR, KRH, ZD, MD, and KH performed phenotypic analysis; MD designed Figure 3A. JW annotated the MRI images. MJSL and JWei performed additional biostatistical analyses. MEC and MN performed in situ sequencing experiments. SN led the study, designed the overall analysis and figure strategy, and wrote the first draft of the manuscript, later revised by all authors.

## Data availability

Data supporting the findings of this study are available within the article and its supplementary figures, supplementary tables, supplementary methods, and supplementary videos. HGCC Phenobank model-level data and microscopy resources are available through the HGCC Phenobank portal (www.hgcc.se). Additional source data are available from the corresponding author Sven Nelander upon reasonable request, subject to ethical and data-use restrictions. Public datasets and resources used for secondary analyses, including TCGA, LINCS, DepMap/PRISM, and GSE117891, are cited in the relevant Methods sections.

