## Supplementary figures and images for "A Multi-Omic Phenobank Reveals Axes of Glioblastoma Growth, Invasion, and Therapeutic Vulnerability"

### supplementary figure 1

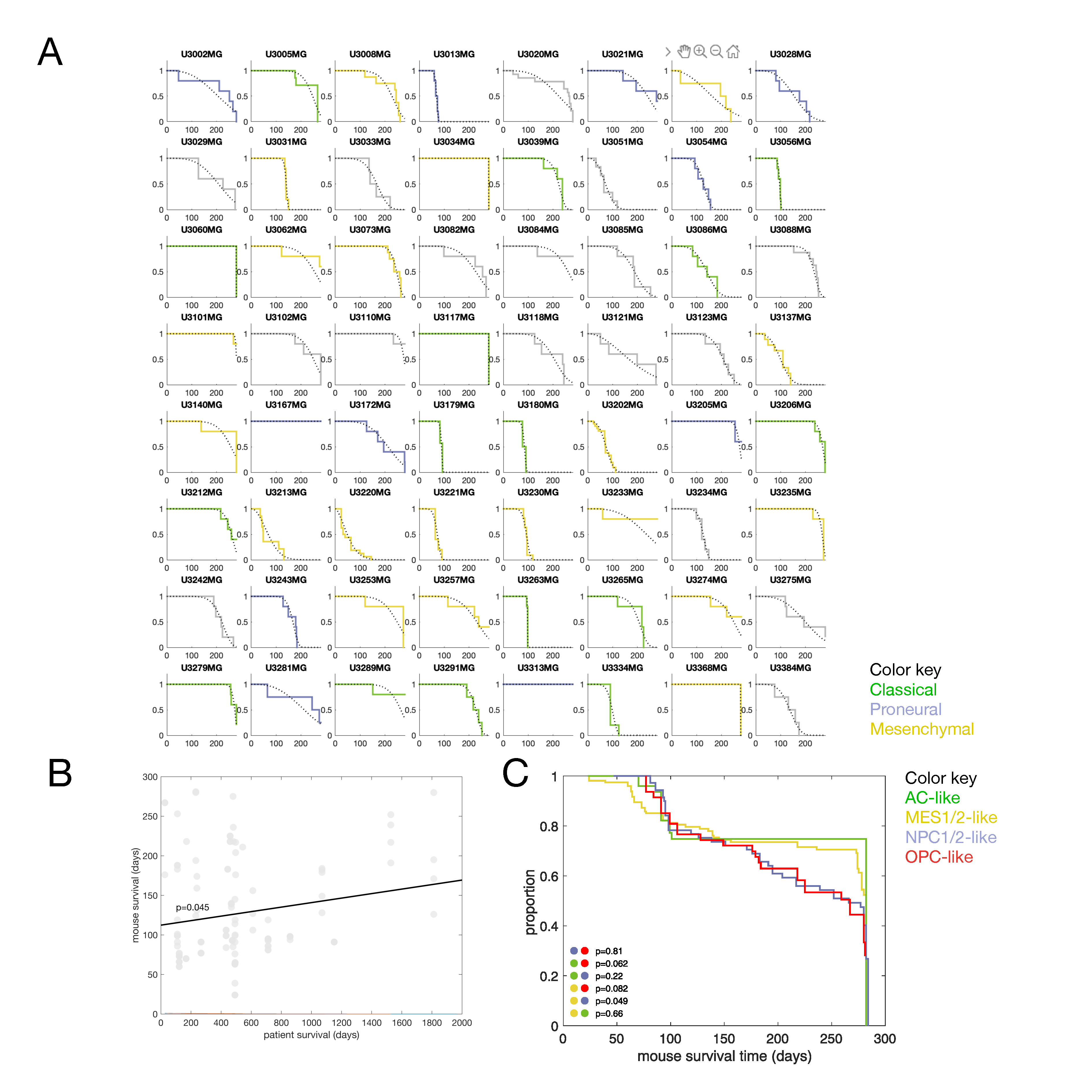

### supplementary figure 2

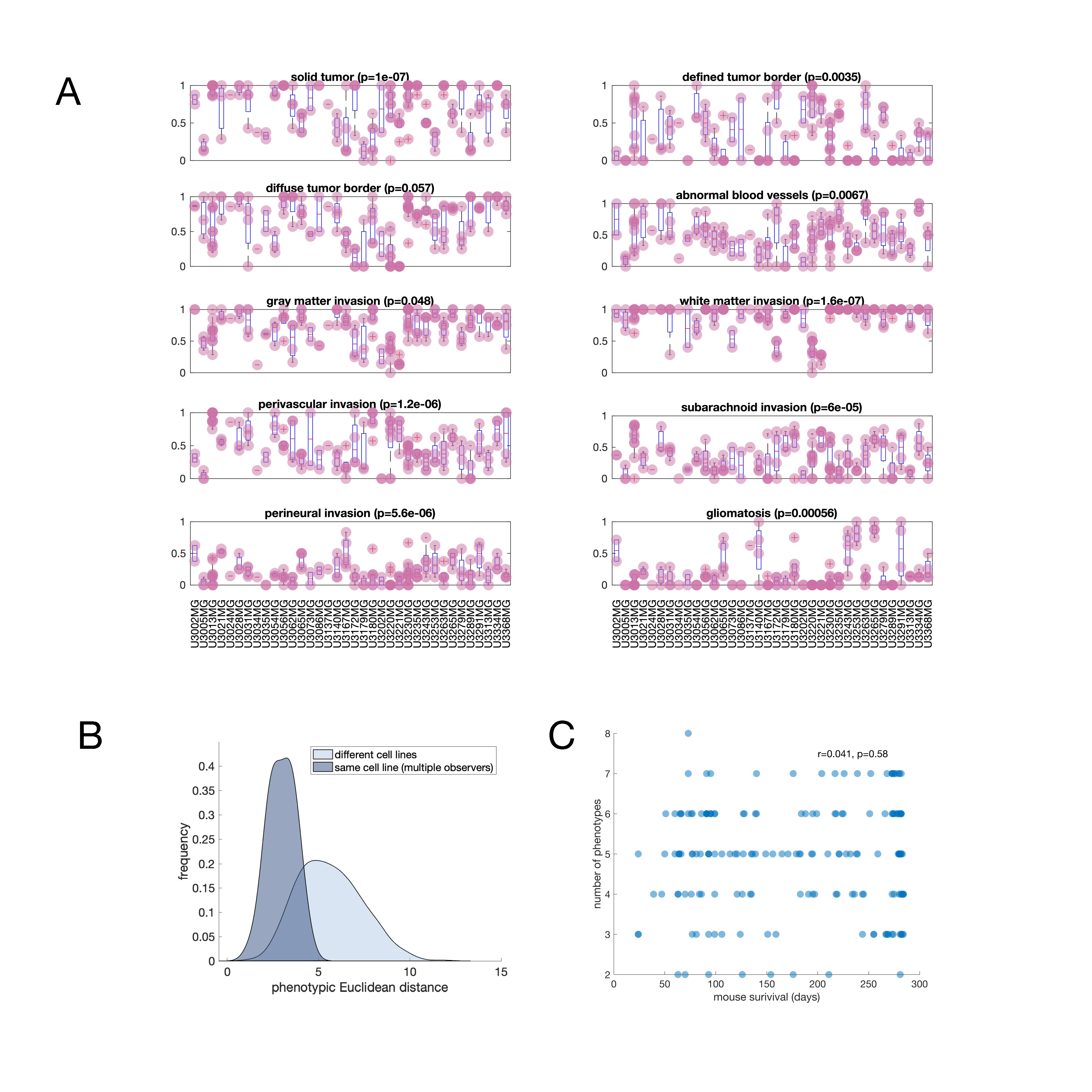

### supplementary figure 3

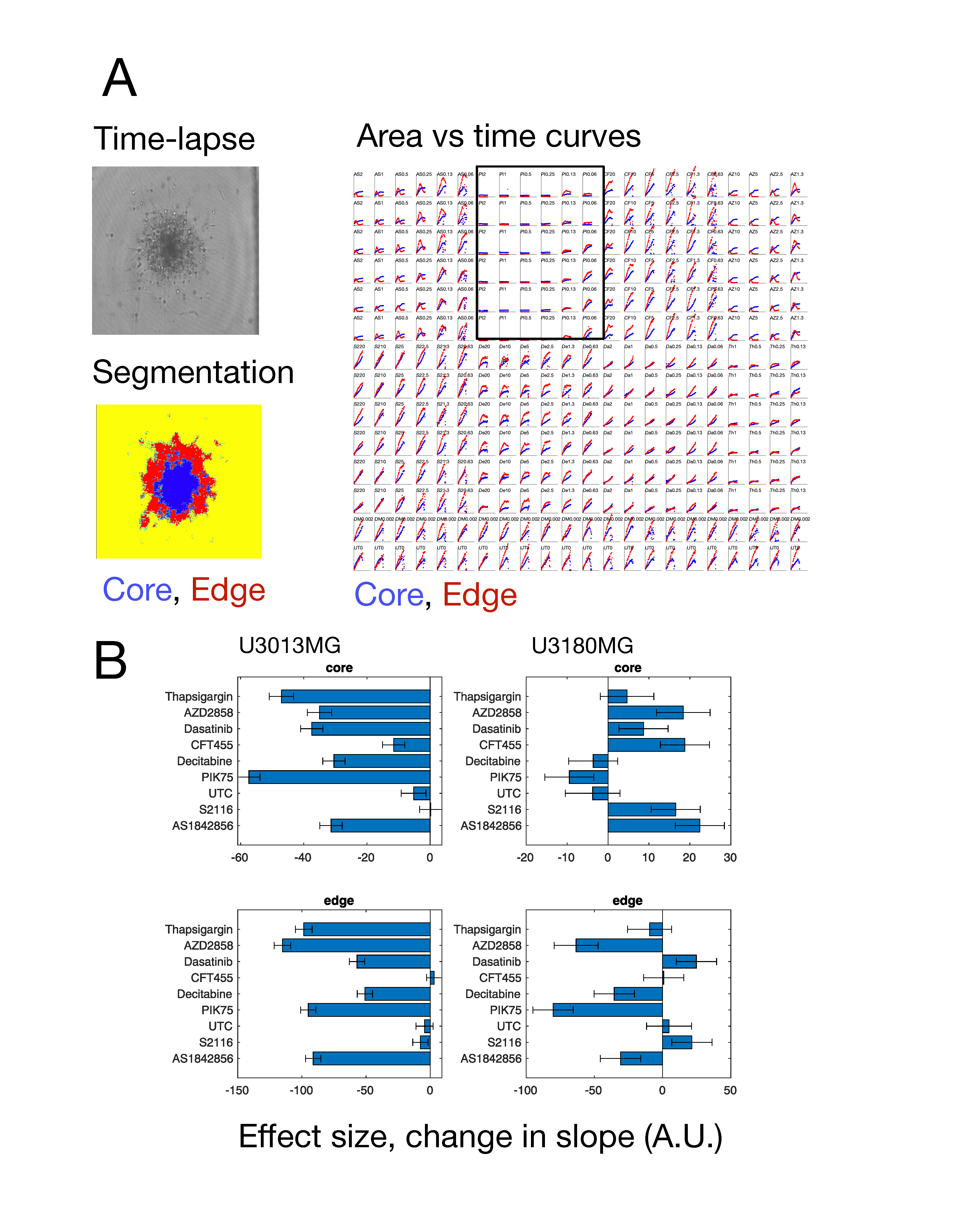

### supplementary figure 4

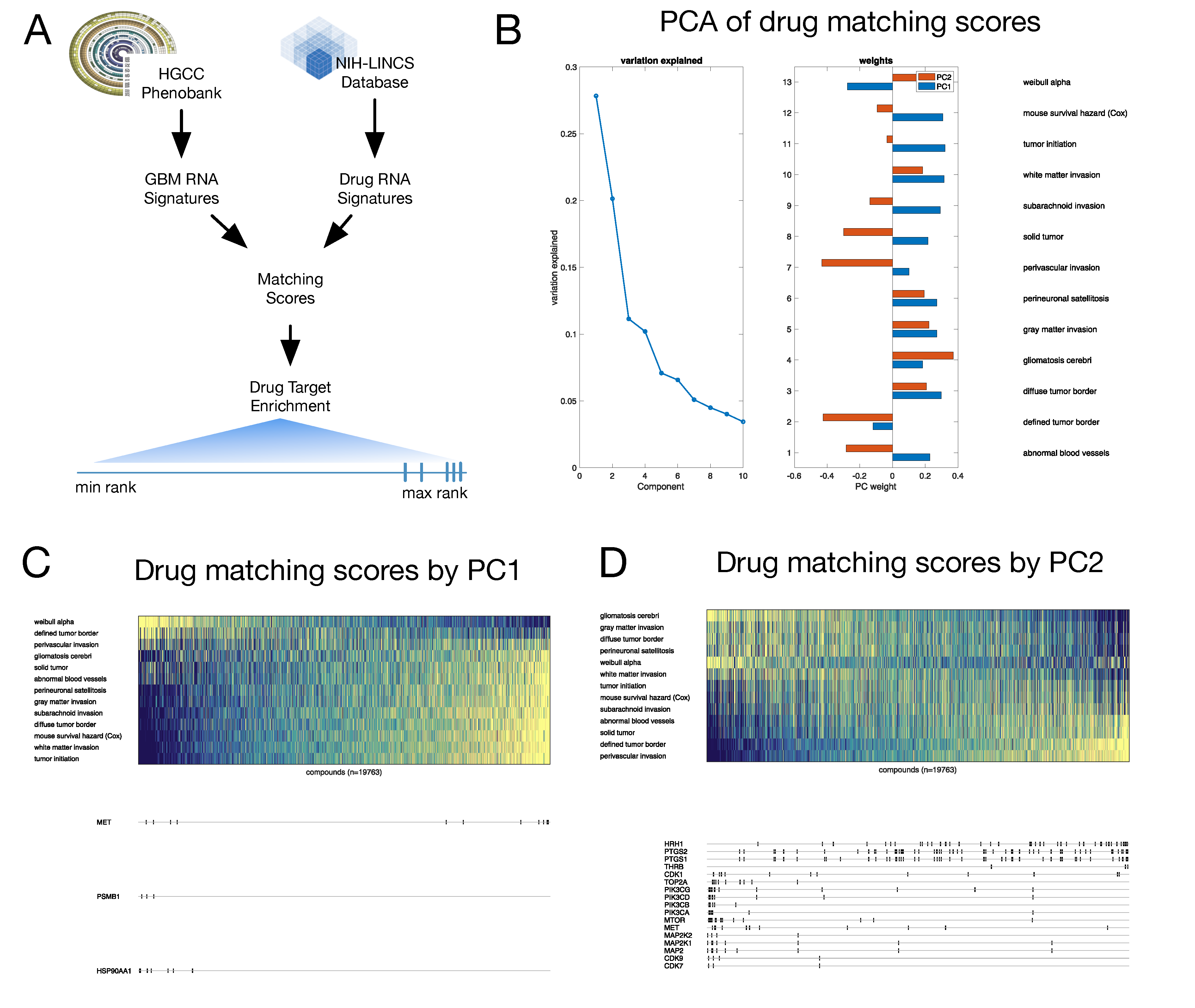

### supplementary figure 5

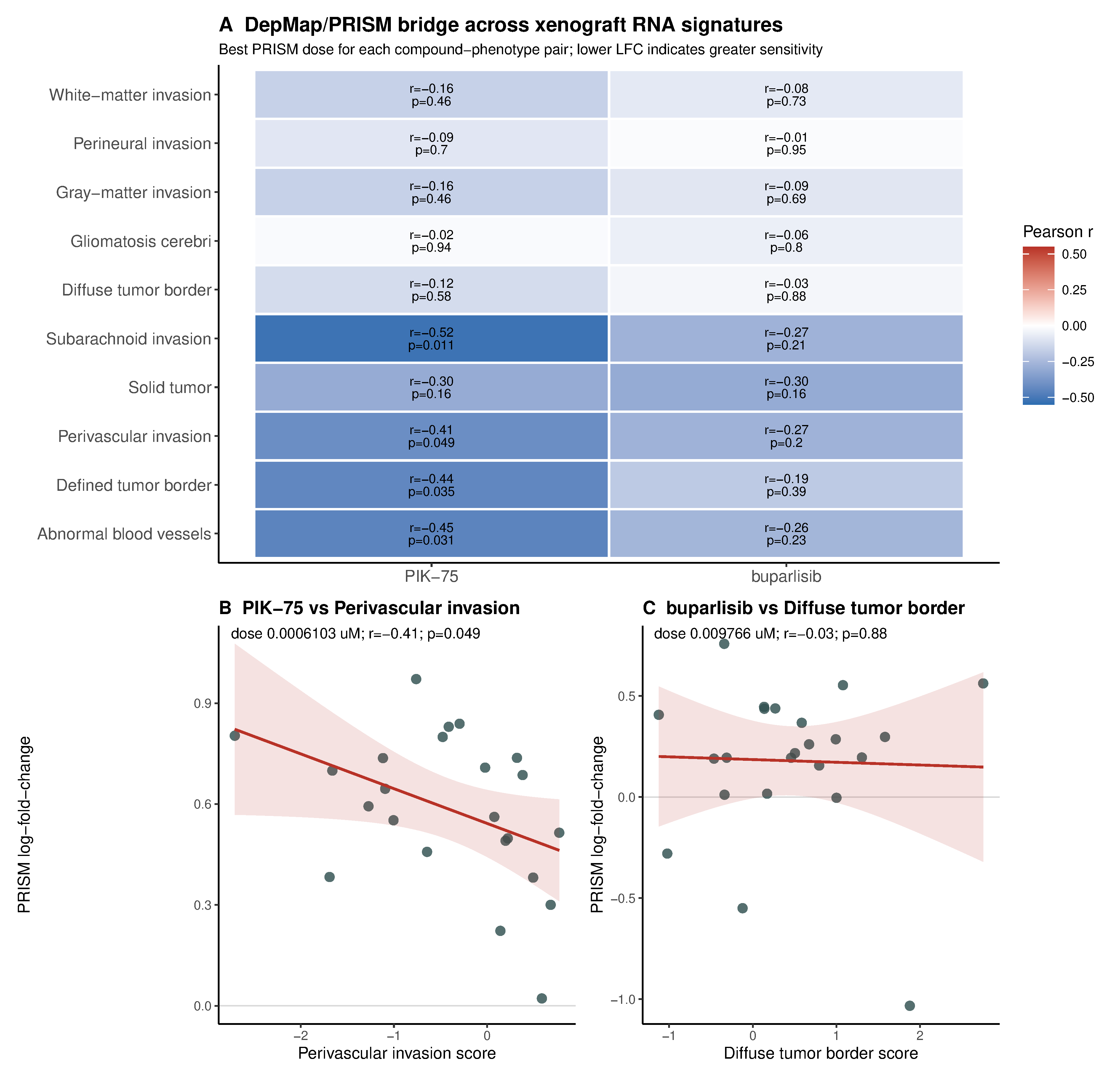

### supplementary figure 6

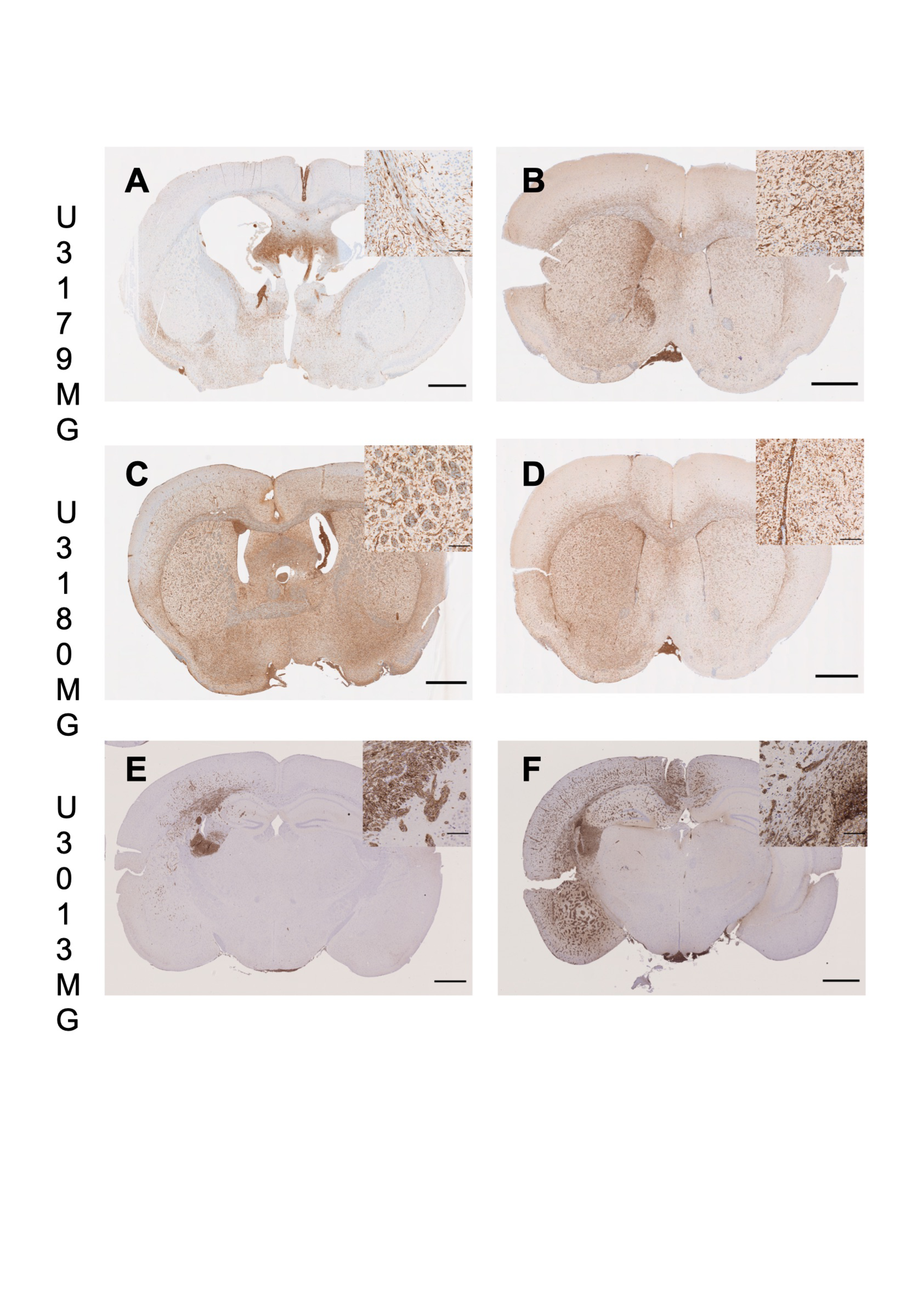

### supplementary figure 7

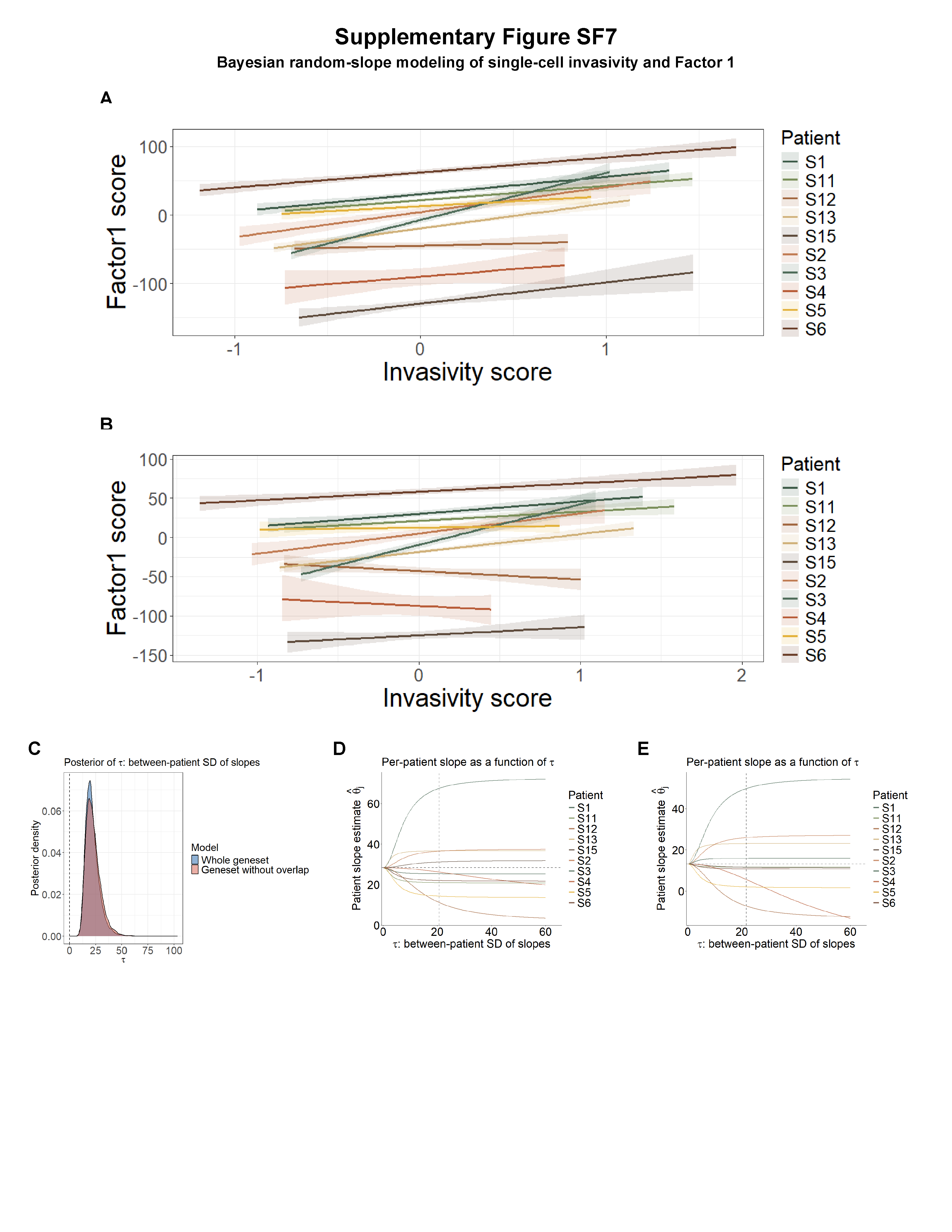

### supplementary figure 8

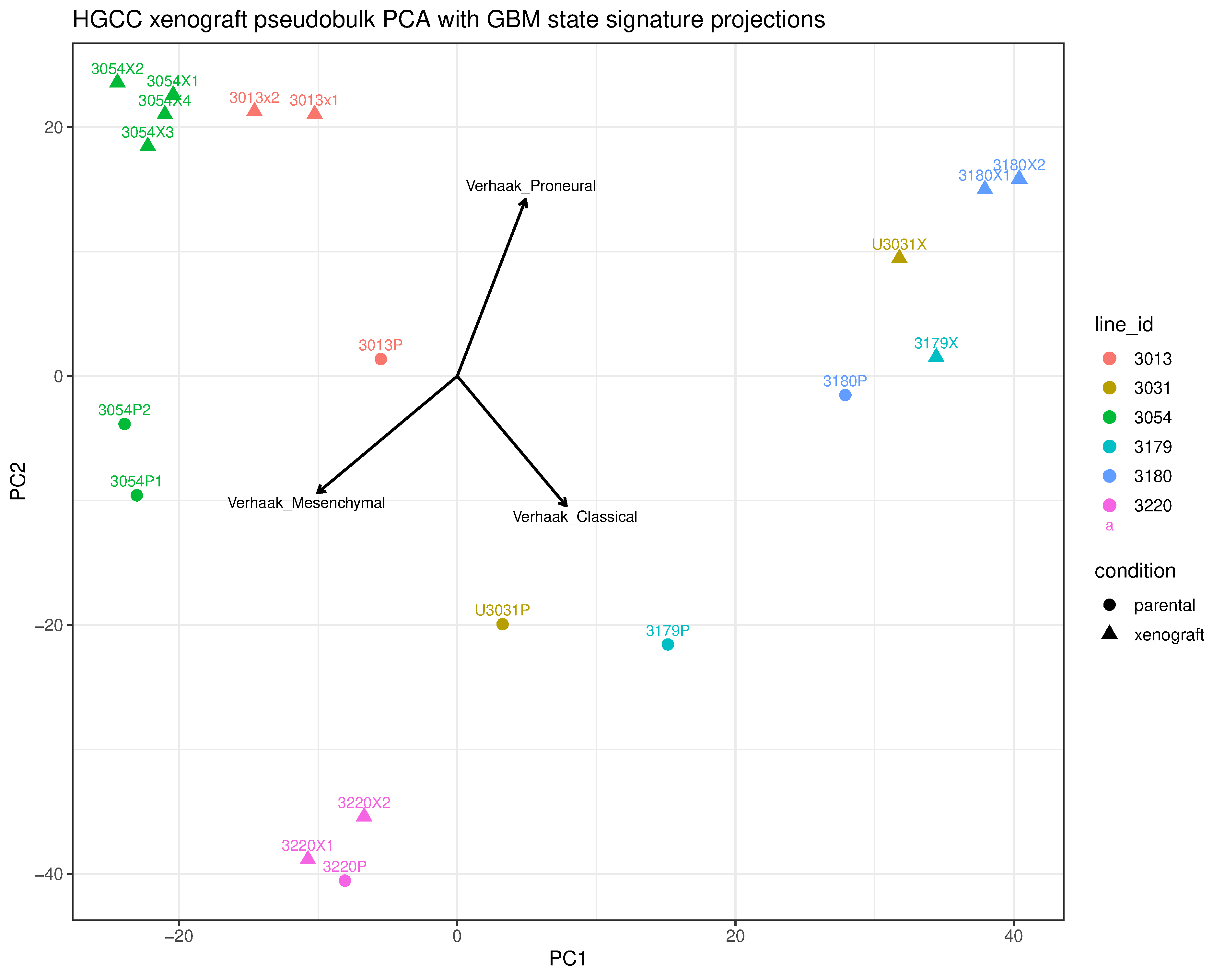
