## Supplementary methods for "A Multi-Omic Phenobank Reveals Axes of Glioblastoma Growth, Invasion, and Therapeutic Vulnerability"

**Materials and Methods**

**HGCC database and cell culture.** PDCs were established from glioblastoma surgical specimens and collected, following informed consent from all subjects included (ethical permit 2007/353), within the HGCC biobank[1,2] using a standardized protocol, in which dissociation of tissue is followed by adherent culture on laminin (Sigma-Aldrich) coated Primaria plates (Corning). Adherent cells were dissociated in Accutase (Gibco) and passaged in serum-free neural stem cell (NSC) media containing an equal mix of DMEM/F12 (1:1) with GlutaMAX (Gibco) and neurobasal media (Gibco) supplemented with 1x B27 (Gibco), 1x N-2 supplement (Gibco), penicillin-streptomycin (Sigma-Aldrich), bFGF (10 ng/mL, Peprotech) and EGF (10 ng/mL, Peprotech) at 37°C with 5% $CO_2$. Cell pellets were submitted to Eurofin Genomics for identification by STR profiling to ensure the identity of each PDC.

**Vectors and viral transduction of GFP-luciferase.** The GFP-luciferase reporter was introduced into the genome of each PDC culture by a lentiviral vector containing GFP-luc2 driven by the CMV promoter. To generate the lentivirus, 293T cells were transfected with plasmids encoding the vesicular stomatitis virus G envelope, gag-pol, and GFP-luc2 (pBMN(CMV-copGFP-Luc2-Puro, Addgene plasmid #80389, a kind gift from Prof. Magnus Essand, Uppsala University). After 24 hours, the conditioned medium was harvested, filtered (0.45 $\mu m$), and ultracentrifuged to produce a high titer virus. Aliquoted virus stock was kept frozen until use. Cell cultures were infected at MOI 1-10. After 24 hours, the virus was removed, and cells were subjected to a 2-week selection for infected cells with 0.5-1.25 $\mu g/ml$ puromycin.

**Orthotopic injection of PDCs into mice.** Animal protocols were reviewed and approved by the Uppsala animal research ethics committee (permits C41/14, 5.8.18-02571-2017, 5.8.18-01070-2019, and 5.8.18-06726-2020). Following dissociation, counting, and washing in PBS, 100,000 PDCs dissolved in 2 $\mu$l PBS were stereotactically injected into the striatum of isofluorane (Baxter Medical AB, Sweden) anesthetized immunodeficient mice of the following strains NOD scid (NOD.CB17-Prkdcscid/J, The Jackson Laboratory, stock #001303 and NOD/MrkBomTac-Prkdcscid, Taconic), NSG (NOD.Cg-Prkdc<scid> Il2rg<tm1Wjl>/SzJ, The Jackson Laboratory, stock #005557), NOG (NOD.Cg-Prkdcscid Il2rgtm1Sug/JicTac, Taconic), and NMRI nude (Rj:NMRI-Foxn1 nu/nu, Janvier Labs) at a median age of 8 weeks (range, 6-14 weeks; 95% CI, 7.6-8.1 weeks). The median passage of the 76 injected cell cultures was 21 (range 13-34 passages; 95% CI, 19.6-22.4). The cells were injected into the striatum at the following coordinates using bregma as reference point 0: AP 0, ML 1.5 (right), DV -3.0. The median number of mice injected with the same PDC culture was 5, with a minimum of 4 and a maximum of 21 injected animals (95% CI, 5.5-7.7 animals). The choice of strain evolved during the project as institutional availability and animal-welfare practices changed, with later experiments transitioning toward NMRI nude mice. Representative histology from matched or comparable strain contexts showed preservation of model-characteristic growth patterns where material was available. Because strain-specific survival data were incomplete, including infection-related losses in one strain, strain was not used for formal survival comparisons. Other injection coordinates were also tested and were not found to affect tumor engraftment capacity (not included in the analyzed subset of this paper). A preoperative subcutaneous injection of the nonsteroidal anti-inflammatory drug Carprofen (5 mg/kg, Orion Pharma Animal Health, Sweden) was administered before surgery and the following day to control postoperative pain. Mice were allowed to recover under supervision and were monitored for up to 40 weeks.

**Bioluminescence imaging.** For bioluminescence imaging, mice were anesthetized by inhalation of 2% isofluorane (Baxter Medical AB) and intraperitoneally injected with 75 mg/kg luciferin (Promega) diluted in PBS. Measurements were taken using a NightOWL *in vivo* imaging system (Berthold Technologies) after 5 minutes of incubation following injection of the substrate. Tumor light output was quantitated using the indiGO software version 2.0.5.0.

**Tissue processing** Animals were sacrificed at the 40-week endpoint or earlier when tumor growth was indicated by luciferase levels increasing by at least 100-fold, development of neurological symptoms, or a bodyweight decrease of at least 10%. Mouse brains were fixed in 4% phosphate-buffered formaldehyde (Histolab, Sweden) for 2-7 days, divided coronally into five pieces, dehydrated, and embedded in paraffin blocks using an automated tissue processor system (TPC15 DUO, Medite Medizintecknik, Germany).

**Immunohistochemical analyses.** Paraffin-embedded mouse brains cut into 3-micron sections were dewaxed using xylene and then rehydrated before hematoxylin and eosin staining, dehydration, and fixation using a xylene based mounting media. In addition, immunohistochemistry using antibodies binding to human-specific nuclear NuMA (ab 97585, Abcam), antibodies binding to human cytoplasmic STEM121 (Y40410, Takara), and anti-Ki67 antibodies (clone MIB-1, M7240 Dako) were also performed. Paraffin sections from xenografted mouse brains were dewaxed using xylene and then rehydrated. Antigen retrieval was performed using Antigen unmasking solution (Vector Laboratories) at 90°C for 20 minutes. Brain sections were then washed using $H_2O$ and incubated with blocking solution 0.3% hydrogen peroxide solution in $H_2O$ followed by washing in tris-buffered saline containing 0.1% tween (TBST) and incubation with primary antibodies diluted in phosphate buffered saline (PBS) overnight at 4°C. After this, they were washed in TBST and incubated with horseradish peroxidase (HRP) conjugated secondary antibodies for 2 hours at room temperature (RT) followed by additional washing steps and incubation with DAB Quanto (Thermo Fisher Scientific). The following secondary antibodies were used: goat anti-mouse IgG HRP conjugated (AP308P, Millipore) and goat anti-rabbit IgG HRP conjugated (AP307P, Millipore). Slides were then dehydrated and mounted using a Mounting medium for light microscopy Pertex® (HistoLab), and stainings were evaluated prior to scanning and processing as outlined below.

**Image Acquisition.** The IHC slides were scanned at the Swedish Science for Life Laboratory (SciLifeLab) Tissue Profiling facility at Uppsala University using an Aperio ScanScope XT Slide Scanner (Aperio Technologies, Vista, CA, USA) with a magnification of 20x and resolution of 0.5$\mu m$. Digital images were acquired in Aperio's SVS format (single-file pyramidal tiled TIFF, with non-standard metadata and compression, http://openslide.org/formats/aperio/) as 24-bit RGB images with dimensions of about 60 000 by 40 000 pixels.

**Determination of tumor initiation.** All IHC slides were manually inspected for the presence of tumor cells, regardless of end-point luciferase signal. Slides with a proliferating bulky tumor or with proliferating invasive tumor cells were considered to have developed an active tumor. Slides with no tumor cells, or a few residual non-proliferating tumor cells, were considered non-tumor initiating.

**Scoring of growth phenotypes.** A selection of 8 individual observers determined upon 10 distinct growth patterns in either a selection of 5 IHC slides (that were not included in the study) or as described in[3]. Each NuMA- and STEM121-stained IHC section with a determined tumor was scored by each observer individually on each phenotype. All observers were blinded, and the order of slides was randomized between observers to prevent eventual drift over time. If an observer deemed that the phenotype was expressed in the tumor, they scored a one. If not, they scored a zero. The scoring was done in two batches, with roughly the same number of mice in each batch.

**Estimation of PDC specific growth phenotypes.** Each phenotype, observer, and tumor triplet were treated as a random Bernoulli distributed variable weighted by: i) the average expression of the phenotype ($\alpha_p$), ii) PDC-specific expression ($\beta_{p,c}$), iii) the expression of the phenotype in that particular tumor ($\gamma_{p,t}$) and iv) the bias the specific expert has for that phenotype ($\rho_{p,e}$). Thus, for score $S_{p,t,e}$, phenotype $p$, PDC $c$, tumor $t$ and expert $e$:

$$S_{p,t,e} \sim \text{Bernoulli}\left(\text{logit}^{-1}(\gamma_{p,t} + \rho_{p,e})\right)$$

with priors:

$$\alpha_p \sim \mathcal{N}(0, 10^2) \quad \text{(Average expression of phenotype } p\text{)}$$

$$\sigma_p \sim \text{HalfNormal}(10) \quad \text{(Scale parameter for phenotype variability)}$$

$$\beta_{p,c} \sim \mathcal{N}(\alpha_p, \sigma_p^2) \quad \text{(PDC-specific expression for phenotype } p\text{)}$$

$$\theta_{p,c} \sim \text{HalfNormal}(10) \quad \text{(Scale parameter for tumor-level variability within PDC } c\text{)}$$

$$\gamma_{p,t} \sim \mathcal{N}(\beta_{p,c}, \theta_{p,c}^2) \quad \text{(Tumor-specific expression of phenotype } p\text{)}$$

$$\rho_{p,e} \sim \mathcal{N}(0, 10^2) \quad \text{(Bias of expert } e \text{ for phenotype } p\text{)}$$

The point estimate for each phenotype and PDC pair was calculated as the expected value of the $\beta_{p,c}$ posteriors. $\theta_{p,c}$ was set to zero for PDCs with only one tumor. Since the scoring was done in two batches, we treated each investigator as a unique person in each batch. Hence, if any investigator changed their bias between the batches, it would be captured in $\rho$.

***In vivo* survival analysis.** PDC-specific survival curves were scored using Cox regression (Matlab fitcox), to assess the log hazard of individual PDCX models compared to baseline. Survival curves were also fitted to the Weibull distribution using wblfit (Matlab).

**Library preparation and RNA sequencing.** Total RNA was extracted using the Direct-zol RNA MiniPrep (Zymo Research) following the manufacturer's instructions and eluted in DNAse/RNAse free water. DNase-treated RNA was evaluated on a Fragment Analyzer Automated CE System (AATi) using a DNF-471 Standard Sensitivity RNA kit (Agilent). Sequencing libraries were prepared from 500ng total RNA using the TruSeq stranded mRNA library preparation kit (Cat# 20020595, Illumina Inc.), including polyA selection. Unique dual indexes (cat# 20022371, Illumina Inc.) were used. The library preparation was performed according to the manufacturers' protocol (#1000000040498). The quality of the libraries was evaluated using the Fragment Analyzer system and a DNF-910 dsDNA kit. The adapter-ligated fragments were quantified by qPCR using the KAPA Biosystems SYBR FAST Universal qPCR Kit and Primer Premix (Roche-Diagnostics) on a CFX384 Touch Real-Time PCR Detection System (Bio-Rad) prior to cluster generation and sequencing. The sequencing libraries were pooled and subjected to cluster generation, and sequencing with paired-end 50 bp read-length run on an SP flowcell on the NovaSeq 6000 system (Illumina Inc.) using the v1.5 sequencing chemistry according to the manufacturer's protocols. A negative control was included in the preparation of libraries and a positive control of 1% of PhiX control library during sequencing. At least 650 million read-pairs per flow cell, with at least 85% of the bases having a base quality score of 30 or higher, was generated.

**RNA sequencing normalization.** Normalization was done in R v3.3, package edgeR 3.16 and TMM normalization[4]. Genes were log-transformed using $log(0.1+\text{normalized counts})$, and genes with mean normalized log count smaller than $log(1.5)$ were removed.

**Expression-based subtyping.** Based on the normalized gene expression data, PDCs were classified as proneural, classical or mesenchymal using single sample gene set enrichment analysis (ssGSEA) with the Wang et al. glioma-intrinsic subtype signatures[5] (**Supplementary Table ST2**). For each subtype signature, ssGSEA scores were Z-transformed across HGCC cultures, and each culture was assigned the subtype with the highest Z-transformed score.

**ssGSEA.** Each PDC was given an enrichment score using the ssGSEA function in gseapy 0.10 for each gene set of interest. Neftel et al. cellular-state meta-modules (MES1, MES2, AC, OPC, NPC1, NPC2, G1/S, and G2/M) were scored with the same ssGSEA framework[6] (**Supplementary Table ST2**). Gene set enrichment scores were regressed against the components using linear regression (statsmodels 0.12). The false discovery rate was controlled using the Benjamini-Hochberg method within each distinct collection of gene sets (neural cell type, GBM subtype, and chromosome location) with $\alpha = 0.05$.

**Tumor-engraftment multi-omic association analysis.** To identify features associated with tumor engraftment and tumor-growth metrics, molecular features were analyzed against model-level phenotypic summaries including tumor-engraftment frequency, fraction of mice with macroscopic tumor, fraction of mice with detected tumor cells, number of mice with macroscopic tumor, Weibull latency parameters, and Cox-regression log-hazard estimates. Continuous variables were compared using univariate linear regression, and binary or fractional engraftment summaries were modeled using the corresponding model-level aggregate values. Association statistics were signed according to the direction of the regression coefficient and visualized as signed $\log_{10} p$-values. Gene-set collections were tested separately, and false-discovery rates were controlled within each collection using the Benjamini-Hochberg method.

**Single-cell Factor 1–invasivity regression.** To test whether Factor 1 tracks an independent single-cell invasion state, we analyzed the Yu et al. GBM single-cell RNA-seq dataset (GSE117891) after excluding non-GBM and IDH1-mutant tumors and retaining malignant cells. Invasivity scores were computed from the invasion pseudotime signatures described by Venkataramani et al.[7]. Seurat AddModuleScore was applied separately to 19 invasion-correlated genes and 141 invasion-anticorrelated genes, and the anticorrelated score was subtracted from the correlated score. Scores were centered within each tumor by subtracting the mean invasivity score across malignant cells from the same tumor. Factor 1 scores were computed by projecting MOFA Factor 1 loadings onto the single-cell data: expression values for genes shared between the loading table and the Seurat object were z-scored across cells, multiplied by their Factor 1 loading weights, and summed per cell. Before model fitting, both Factor 1 scores and centered invasivity scores were residualized against nCount_RNA to reduce sequencing-depth effects.

The relationship between Factor 1 and invasivity was estimated using a Bayesian hierarchical random-slope model:

$$\text{F1}_{ij} = \beta_0 + \beta_1 S_{ij} + u_{0j} + u_{1j} S_{ij} + \varepsilon_{ij},$$

where $i$ indexes cells, $j$ indexes patients, and $S_{ij}$ is the within-tumor-centered invasivity score. Patient-specific intercepts and slopes were modeled as correlated random effects. Models were fit with `brms` using four Hamiltonian Monte Carlo chains, 4,000 iterations per chain, 1,000 warmup iterations, and `adapt_delta`=0.95. Weakly informative priors were used: $\beta_0, \beta_1 \sim \mathcal{N}(0, 200^2)$, standard deviations $\sim$ half-Student-$t(3, 0, 50)$, and an LKJ(2) prior for the random-effect correlation matrix. The same model was refit after removing the 12 genes shared between the invasivity signature and Factor 1 loading set to evaluate sensitivity to direct gene overlap. Model diagnostics and posterior summaries are shown in Supplementary Figure SF7.

**Parental culture–xenograft pseudobulk RNA comparison.** To assess whether parental HGCC cultures and matched xenograft-derived mouse-brain samples remained coherent in RNA space, we analyzed a private single-cell RNA-seq bundle comprising 19 selected 10x Genomics filtered feature-barcode matrices from parental and xenograft-derived HGCC cultures. The analysis included U3013MG, U3031MG, U3054MG, U3179MG, U3180MG, and U3220MG; one misidentified sample was excluded, and U3065MG-labeled runs were reassigned to the second U3054MG branch based on the documented sample mix-up. For each run, raw counts were summed gene-wise across all cells to generate a sample-level pseudobulk expression vector. Genes were filtered to retain those with counts per million greater than 1 in at least three samples, converted to $\log_2(\text{CPM} + 1)$, and the 2,000 most variable genes were z-scored across samples before principal component analysis with `prcomp` in R. In the 19-sample analysis, PC1 explained 27.5% and PC2 explained 24.1% of the variance. Sample-sample similarity was summarized by Pearson correlation between log-CPM pseudobulk profiles. Median correlation was higher for pairs from the same cell line than for pairs from different cell lines (0.905 versus 0.814), while the condition effect was smaller (same-condition median 0.832 versus different-condition median 0.819). Verhaak proneural, classical, and mesenchymal subtype directions were overlaid on the PCA to aid biological interpretation of the dominant axes. PCA and nearest-neighbor analyses were also inspected using the top 1,000 variable genes and all filtered genes, with no qualitative change in sample-level structure.

**Mass-spectrometry-based proteomics.** The samples were prepared and analyzed following the HiRIEF LC-MS/MS protocol, as previously described[8]. Brief description is provided below.

**Cell lysis and in-solution digestion.** The PDCs were lysed in 200 $\mu$l SDS-lysis buffer (containing 4% (w/v) SDS, 50 mM HEPES pH 7.6, and 1 mM dithiothreitol) using 1:4-10 of sample to buffer ratio. Afterwards, the cells were heated at 95°C for 5 min while shaking on a pre-warmed block and sonicated to dissolve the pellet and disrupt the remaining DNA. The lysate was then centrifugated at 14,000×g for 15 min, and the supernatant was removed. The protein concentration in the lysate was determined by Bio-Rad DC Assay, and equal amounts of each sample were subjected to in-solution digestion. Briefly, the cell pellet was denatured at 95°C for 5 min, followed by reduction with dithiothreitol and alkylation with chloroacetamide at end concentrations of 5 mM and 10 mM, respectively. Lys-C was added at a 1:50 (w/w) ratio, and digestion was performed at 37°C, for six hours. The samples were further digested by trypsin at a 1:50 (w/w) ratio with 37°C overnight incubation. After LysC/trypsin digestion, 1% of each peptide sample was aliquoted for 15 min gradient LC-MS/MS runs to check for protease activity by the samples' miscleavage rate. **TMT-labelling.** Before labeling, equal amounts of peptide samples were pH-adjusted using TEAB, pH 8.5. The resulting peptide mixtures were labeled with Thermo Scientific isobaric Tandem Mass Tags (TMT). The three biological replicates of the PDCs were distributed across three TMT-16-plex sets, with 30 $\mu g$ protein from each replicate labeled with a randomly assigned TMT channel in a different set, baring previously allocated channel(s). The internal standards were made of sample pools. Labeling efficiency was determined by LC-MS/MS before pooling of samples. Subsequently, sample clean-up was performed by solid phase extraction (SPE strata-X-C, Phenomenex). The labeling scheme is provided in Supplementary Table ST9.

**High resolution isoelectric focusing (HiRIEF).** After sample clean-up, the sample pool was subjected to peptide IEF-IPG (isoelectric focusing by immobilized pH gradient) in pI range 3-10 (total of 480 $\mu g$ per set, per strip). The freeze-dried peptide sample was dissolved in 250 $\mu l$ rehydration solution containing 8 M urea and allowed to adsorb to the gel strip by swelling overnight. The 24 cm linear gradient IPG (Immobilized PH Gradient) strip (GE Healthcare) was incubated overnight in 8 M rehydration solution containing 1% IPG pharmalyte pH 3-10 (GE Healthcare). After focusing, the peptides were passively eluted into 72 contiguous fractions with MilliQ water / 35% acetonitrile / 35% acetonitrile and 0.1% formic acid, using an in-house constructed IPG extractor robotics (GE Healthcare Biosciences AB, prototype instrument) into a 96-well plate (V-bottom, Greiner product #651201). The resulting fractions were concatenated in 40 fractions, then dried, frozen, and kept at -20°C until LC-MS/MS analysis.

**LC-MS/MS analysis.** Online LC-MS/MS was performed using a Dionex UltiMate™ 3000 RSLCnano System coupled to a Q-Exactive HF mass spectrometer (Thermo Scientific). Each plate well was dissolved in 20 $\mu$l solvent A, and 10 $\mu$l were injected. Samples were trapped on a C18 guard-desalting column (Acclaim PepMap 100, 75 $\mu$m x 2 cm, nanoViper, C18, 5 $\mu$m, 100 Å), and separated on a 50 cm long C18 column (Easy spray PepMap RSLC, C18, 2 $\mu$m, 100 Å, 75 $\mu$m x 50 cm). The nano capillary solvent A was 94.9% water, 5% DMSO, 0.1% formic acid; and solvent B was 4.9% water, 5% DMSO, 90% acetonitrile, 0.1% formic acid. At a constant flow of 0.25 $\mu$l min−1, the curved gradient went from 2% B up to 40% B in each fraction, followed by a steep increase to 100% B in 5 min and subsequent re-equilibration with 2% B. Some of the fractions were pooled and analyzed together; details on pooling and gradient length per fraction are available in Supplementary Table ST10. FTMS master scans with 60,000 resolution (and mass range 300-1700 m/z) were followed by data-dependent MS/MS (30,000 resolution) on the top 5 ions using higher energy collision dissociation (HCD) at 30% normalized collision energy. Precursors were isolated with a 2 m/z window. Automatic gain control (AGC) targets were 16 for MS1 and 15 for MS2, with minimum AGC target of 13. Maximum injection times were 100 ms for MS1 and 100 ms for MS2. The entire duty cycle lasted 2.5 s. Dynamic exclusion was used with 30.0 s duration. Precursors with unassigned charge state or charge state 1, 7, 8, or >8 were excluded.

**Protein identification.** Raw MS/MS files were converted to mzML format using msconvert from the ProteoWizard tool suite[9]. Spectra were then searched in the Galaxy framework using tools from the Galaxy-P project[10,11], including MSGF+[12] (v2020.03.14) and Percolator[13] (v3.04.0), where eight subsequent HiRIEF search result fractions were grouped for Percolator target/decoy analysis. Peptide and PSM (Peptide Spectrum Matches) FDR (False discovery rate) were recalculated after merging the percolator groups of eight search results into one result per TMT set. The reference database used was the human protein subset of ENSEMBL104. Quantification of isobaric reporter ions was done using OpenMS project's IsobaricAnalyzer[14] (v2.5.0). Quantification on reporter ions in MS2 was for both protein and peptide level quantification based on median of PSM ratios, limited to PSMs mapping only to one protein and with an FDR q-value < 0.01. FDR for protein level identities was calculated using the -log10 of best-peptide q-value as a score. The search settings included enzymatic cleavage of proteins to peptides using trypsin limited to fully tryptic peptides. Carbamidomethylation of cysteine was specified as a fixed modification. The minimum peptide length was specified to be six amino acids. Variable modification was oxidation of methionine.

**Multi-omic factor analysis.** Multi-Omics Factor Analysis (MOFA)[15] was performed using the MOFA2 package v1.12.1 in R v4.3.3. The input data consisted of preprocessed matrices for ssGSEA scored gene sets, DNA methylation, RNA-seq, and proteomics data. The model was trained using standard parameters to identify latent factors capturing shared and unique sources of variation across omics layers. Covariates from different modalities were incorporated into the analysis to enhance the interpretability of the latent factors identified by MOFA. The first MOFA-derived factor was projected on The Cancer Genome Atlas (TCGA)[16] dataset to investigate survival differences. Patient samples were stratified into high and low groups based on their Factor 1 scores, and Kaplan-Meier survival analysis was conducted using the survival v3.7-0 and survminer v0.5.0 R packages. Cox proportional hazards models were used to assess the association between Factor 1 and overall survival.

**Target Translator Algorithm.** We applied the method described in[17], incorporating the following adaptations. For each phenotypic trait (e.g., white matter invasion), we derived an RNA signature by computing the correlation of each gene's expression with the corresponding trait. These signatures were then scored according to[17], generating a matrix $X$ of match scores, where rows corresponded to phenotypic traits and columns to 19,763 drug-induced transcriptional signatures from the NIH-LINCS database. To account for collinearity in the drug response profiles, we performed singular value decomposition (SVD), expressing the matrix as $X = USV^T$. We ranked drugs based on the first two principal components (PC1 and PC2) extracted from the matrix $V$. Component analysis revealed that PC1 primarily reflected tumor initiation, while PC2 distinguished drugs targeting perivascular versus diffuse invasion (Supplementary Figure SF4). Next, we mapped the ranked drugs to the Drug Repurposing Hub database and identified their respective molecular targets. To assess the enrichment of specific drug targets along PC1 and PC2, we performed a Kolmogorov-Smirnov test, applying a significance threshold of $p < 0.001$. Only targets meeting this criterion were included in Figure 6.

**2D live-cell image proliferation analysis assay.** Based on the Target Translator predictions, ten compounds inhibiting targets associated with bulky, perivascular growth and eleven compounds against targets associated with diffuse growth were evaluated in the representative U3013MG and U3180MG PDC lines. Cells were seeded in NSC medium using a Viaflo 96/384 electronic multichannel pipette (Integra) at a density of 1000 cells per well in 384-well microplates (Nunc, #142761) pre-coated with laminin. The compounds were added the following day with a D300e digital dispenser (Tecan) in an 11-point dose dilution series, with 3-fold dilution steps, starting from 30 $\mu$M. Phase contrast images were acquired every hour over the next three days using an IncuCyte S3 live cell imaging system (Sartorius). Cell confluency was normalized to the starting density for each well, and linear regression was used to determine the growth rate of treated cells relative to DMSO controls.

**Sphere invasion assay.** A miniaturized 384-well plate format was implemented with 1000 PDCs per well seeded in PrimeSurface 3D Culture Spheroid Ultra-Low attachment plates (S-BIO, #MS-9384UZ) in 40 $\mu$l NSC media supplemented with B27 and N-2 (NSC++) using a Viaflo 96/384 device. Three days later, another 40 $\mu$l NSC++ media and Matrigel (Corning, #354234) at a final concentration of 4-5 mg/ml was added. Images of the sphere core and the invading edge population were acquired every sixth hour over seven days using the spheroid module of an IncuCyte S3 live cell imaging system with 10X phase and brightfield imaging. The eight compounds were also added on day three using the D300e digital dispenser with six replicates each at 4-6 different concentrations in a two-fold dilution range **(Supplementary Table ST5)**. The total concentration of DMSO was normalized to 0.2% for all wells, and treatments were evaluated against DMSO vehicle controls (0.2%) and untreated controls (UTC). A custom made analysis pipeline was created for analysis of the core and edge populations

**(Supplementary Figure SF3)**.

**Brain slice assay.** A brain slice culture experiment was carried out following the experimental procedure we described recently[18]. In brief, patient-derived cell culture xenografts were generated with three HGCC lines (U3013MG, U3180MG, and U3054MG) tagged with Green Fluorescent Protein (GFP) and Luciferase (pBMN(CMV-copGFP-Luc2-Puro), Addgene plasmid #80389) as mentioned above. Live tumor-bearing brains, at optimal luciferase signal, were removed and placed in ice-cold HBSS medium (Gibco, #24020117). Individual brains were embedded in a square plastic mold with a low-melting agarose-HBSS medium. Next, 300 $\mu$m thick brain slices were sectioned using a Leica vibratome (Leica VT 1200 S), with the cutting speed adjusted between 20-200 $\mu$m/second. These slices were then transferred onto transwell membrane in a 12-well plate (Corning, #3460) containing brain slice culture medium (NSC, 2.5 mM HEPES, 10 mM glucose). To visualize the vasculature, culture medium was supplemented with 2 $\mu$g/ml Tomato lectin-DyLight 594 conjugate. Excess medium around the slices were removed to facilitate an air-liquid interface for optimal culture conditions. The culture slices were then incubated at 37°C in a 5% $CO_2$ incubator. The following day, the treatments with DMSO (0.3%) as a control and PIK-75 inhibitor at concentrations of 0.5, 1, 2, and 4 $\mu$M or Buparlisib at concentrations of 1, 2, 4, and 8 $\mu$M were added to the slice culture during a media change. Immediately after treatment, the slice culture plate was transferred to the ImageXpress Micro Confocal High-Content Imaging System (IXMC) (Molecular Devices). Time-lapse live images of whole brain slices were acquired at intervals of less than 2 hours for up to 6 days and saved as maximum projection images. The brain slice culture medium, including treatments, was changed every 48 hours. The time-lapse TIFF images were processed and stitched using a montage and overlay journal in MetaXpress 6.5 software (Molecular devices). The TIFF stacks were then registered for frame-to-frame coherence using custom MATLAB (The MathWorks) scripts to facilitate image analysis.

Regions of interest (ROIs) in the tumor periphery with lower cell density suitable for single-cell tracking were manually selected to create stacks of 500x500xT, where T was the length of the time-lapse. To automatically identify cells within each frame of the ROI stack, we applied Gaussian filtering and blob detection by convolution with a Logarithm of Gaussian filter corresponding to average cell size.

Putative cell coordinates were pruned using binary versions of the original image, manipulated with morphological operations to ensure that cell coordinates corresponded to cell bodies, not protrusions, and that each cell had only one coordinate. Cells were tracked throughout the time-lapse using a Kalman filter-based framework. Cell coordinates passed to the tracking framework were connected across frames to create cell trajectories. For each frame, extrapolated coordinates from the previous frame were matched to coordinates in the current frame by solving a linear assignment problem. Cell speed (microns per hour) was averaged across the whole lifetime of the track for each cell and then averaged for all cells detected in a ROI. As a complementary measure of movement, an adjusted mean of absolute differences was calculated according to equation 1, where g is the green channel in stack G of size WxWxT. The absolute difference between two consecutive frames was normalized by frame size and mean pixel intensity in the two frames, to create the ratio of pixel movement between the two frames, adjusted for possible increase in signal between the frames. All algorithms for cell identification, tracking, and subsequent analysis were written in MATLAB (Image Processing Toolbox, Computer Vision Toolbox, Version 23.2, Release 2023b, The MathWorks, Inc., Natick, Massachusetts, United States).

$$adMAD(i) = \frac{|g(:,:,t+1) - g(:,:,t)|}{g(:,:,t+1) + g(:,:,t)}$$

### Supplementary Figures

**Figure SF1: Survival trends for HGCC cultures grafted in mice.** (A) Survival curves for individual HGCC cultures, fitted to Weibull distributions (dotted lines). Colors indicate transcriptional subtypes according to Wang et al. (signatures listed in **Supplementary Table ST2**). (B) Correlation between survival times in mice and the corresponding patient survival times for tumor-forming HGCC cultures (Pearson $r = 0.1832$, $p = 0.045$, $n = 17$ matched pairs). (C) Survival curves grouped by cellular state signatures according to Neftel et al. (signatures listed in **Supplementary Table ST2**).

**Figure SF2: Robustness of phenotyping scores.** (A) ANOVA tests were performed to evaluate whether observed phenotypic differences among HGCC cultures significantly exceed the expected variability due to biological or technical factors. (B) Distribution of phenotypic distances between mice injected with the same HGCC culture (dark distribution) compared to mice injected with different HGCC cultures (light distribution). The distributions were significantly different (K–S two-sample test, $p = 0.004$). (C) Correlation analysis between the number of phenotypes observed and survival time shows no dependency, confirming robustness of the scoring approach.

**Figure SF3: PIK-75 as an inhibitor of core and edge cell expansion in gliomaspheres.** (A) Sphere cultures from HGCC models invading through an extracellular matrix were evaluated using time-lapse microscopy over seven days following treatment with eight different compounds. Compounds were tested at 4–6 concentrations with six technical replicates per concentration; treatments were compared with DMSO vehicle controls and untreated controls (UTC). The selected PIK-75 plots show the 0.06–2 µM dose range. Images were segmented into core and edge areas using a semantic segmentation neural network (U-Net). (B) Growth curves were generated, and rates of change were fitted and analyzed using a linear model to assess drug efficacy.

**Figure SF4: TargetTranslator scoring of drugs and molecular targets.** (A) Overview of the computational approach: RNA signatures derived from the HGCC Phenobank were correlated with NIH-LINCS L1000 data, yielding correlation scores aggregated across cell lines (hits shown in blue). (B) Principal component analysis (PCA) identified two dominant components. The first component (PC1) was associated primarily with tumor initiation and survival hazard, whereas the second component (PC2) distinguished drugs predicted to target perivascular versus diffuse invasion signatures. Drug targets were identified using the Drug Repurposing Hub database, and significantly skewed distributions of drug-target rankings were assessed by a Kolmogorov–Smirnov (K–S) test. (C) Enriched drug targets for PC1. (D) Enriched drug targets for PC2. Targets shown meet a significance threshold of $p < 0.001$.

**Figure SF5: DepMap/PRISM bridge supports PIK-75 sensitivity in vascular and perivascular xenograft RNA-signature states.** (A) Pearson correlations between projected HGCC xenograft RNA-signature scores and PRISM log-fold-change values across GBM IDH-wildtype DepMap lines for PIK-75 and buparlisib. For each compound–phenotype pair, the best-correlating PRISM dose is shown; negative correlations indicate that higher projected phenotype score is associated with greater drug sensitivity. (B) Example scatter plot showing the association between the perivascular-invasion signature and PIK-75 PRISM response. (C) Example weak/non-correlating comparison for the diffuse tumor border signature and buparlisib.

**Figure SF6: Comparison of tumor growth patterns across immunodeficient mouse strains.** Representative Stem121-stained sections show GBM PDC xenografts grown in different immunodeficient strain contexts. U3179MG injected into NMRI-nude (A) and NOG (B), U3180MG injected into NMRI-nude (C) and NOG (D), and U3013MG injected into NOD-scid (E) and NMRI-nude (F). U3179MG and U3180MG show diffuse growth in the brain parenchyma, whereas U3013MG shows dense growth with vascular infiltration. Scale bars indicate 1 mm in whole sections and 100 µm in insets.

**Figure SF7: Bayesian random-slope modeling of single-cell invasivity and Factor 1.** (A) Posterior median fits relating within-tumor-centered invasivity to Factor 1 score in the Yu et al. GBM single-cell dataset using the full invasivity and Factor 1 scoring gene sets. (B) Corresponding fits after removing the 12 genes overlapping between the invasivity signature and the Factor 1 loading set. (C) Posterior distributions of the between-patient slope standard deviation $\tau$, shown for the whole and no-overlap gene sets. (D,E) Shrinkage curves showing patient-specific slope estimates as a function of $\tau$ for the whole gene set (D) and no-overlap gene set (E).

**Figure SF8: Pseudobulk RNA comparison of parental cultures and matched xenograft-derived mouse-brain samples.** Principal component analysis of pseudobulk profiles generated from single-cell RNA-seq data for parental HGCC cultures and xenograft-derived mouse-brain samples. Points are colored by HGCC line and shaped by growth condition. Verhaak subtype projection vectors are overlaid to support interpretation of the PCA axes: negative PC1 aligns with mesenchymal-like programs, whereas positive PC1 aligns with proneural/classical, neurodevelopmental-like programs. PC2 separates parental in vitro cultures from xenograft-derived in vivo samples. Same-line parental and xenograft-derived samples remain relatively close in RNA space, while xenograft-derived samples show greater dispersion.

#### Supplementary Videos

**Supplementary Video 1.** Representative ex vivo brain-slice time-lapse movie following PIK-75 treatment (`Video1_PIK75.mp4`).

**Supplementary Video 2.** Representative ex vivo brain-slice time-lapse movie following buparlisib treatment (`Video2_Buparlisib.mp4`).

[6] C. Neftel, J. Laffy, M. G. Filbin, T. Hara, M. E. Shore, G. J. Rahme, A. R. Richman, D. Silverbush, M. L. Shaw, C. M. Hebert, J. Dewitt, S. Gritsch, E. M. Perez, L. N. Gonzalez Castro, X. Lan, N. Druck, C. Rodman, D. Dionne, A. Kaplan, M. S. Bertalan, J. Small, K. Pelton, S. Becker, D. Bonal, Q. D. Nguyen, R. L. Servis, J. M. Fung, R. Mylvaganam, L. Mayr, J. Gojo, C. Haber-

ler, R. Geyeregger, T. Czech, I. Slavc, B. V. Nahed, W. T. Curry, B. S. Carter, H. Wakimoto, P. K. Brastianos, T. T. Batchelor, A. Stemmer-Rachamimov, M. Martinez-Lage, M. P. Frosch, I. Stamenkovic, N. Riggi, E. Rheinbay, M. Monje, O. Rozenblatt-Rosen, D. P. Cahill, A. P. Patel, T. Hunter, I. M. Verma, K. L. Ligon, D. N. Louis, A. Regev, B. E. Bernstein, I. Tirosh, and M. L. Suva. An Integrative Model of Cellular States, Plasticity, and Genetics for Glioblastoma. *Cell*, 178 (4):835–849, Aug 2019.
