## Supplementary material for "A Multi-Omic Phenobank Reveals Axes of Glioblastoma Growth, Invasion, and Therapeutic Vulnerability": supp table 6

### Motifs found for: abnormal blood vessels

### Homer Known Motif Enrichment Results (homerpos1)

[Homer *de novo* Motif Results](#)
[Gene Ontology Enrichment Results](#)
[Known Motif Enrichment Results (txt file)](#)
Total Target Sequences = 244, Total Background Sequences = 96240

| Rank | Motif | Name | P-value | log P-pvalue | q-value (Benjamini) | # Target Sequences with Motif | % of Targets Sequences with Motif | # Background Sequences with Motif | % of Background Sequences with Motif | Motif File | SVG |
|---|---|---|---|---|---|---|---|---|---|---|---|
| 1 | 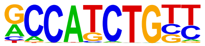 | NeuroD1(bHLH)/Islet-NeuroD1-ChIP-Seq(GSE30298)/Homer | 1e-9 | -2.116e+01 | 0.0000 | 47.0 | 19.26% | 6929.3 | 7.20% | motif file (matrix) | svg |
| 2 | 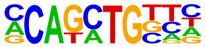 | Twist2(bHLH)/Myoblast-Twist2.Ty1-ChIP-Seq(GSE127998)/Homer | 1e-8 | -1.995e+01 | 0.0000 | 80.0 | 32.79% | 16483.8 | 17.14% | motif file (matrix) | svg |
| 3 | 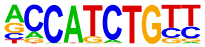 | NeuroG2(bHLH)/Fibroblast-NeuroG2-ChIP-Seq(GSE75910)/Homer | 1e-8 | -1.973e+01 | 0.0000 | 73.0 | 29.92% | 14439.5 | 15.01% | motif file (matrix) | svg |
| 4 | | Atoh1(bHLH)/Cerebellum-Atoh1-ChIP-Seq(GSE22111)/Homer | 1e-8 | -1.948e+01 | 0.0000 | 55.0 | 22.54% | 9399.5 | 9.77% | motif file (matrix) | svg |

| | | | | | | | | | | |
|---|---|---|---|---|---|---|---|---|---|---|
| | 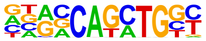 | | | | | | | | | |
| 5 | Sox3(HMG)/NPC-Sox3-ChIP-Seq(GSE33059)/Homer 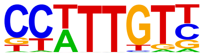 | 1e-7 | -1.821e+01 | 0.0000 | 87.0 | 35.66% | 19334.4 | 20.10% | motif file (matrix) | svg |
| 6 | Atoh7(bHLH)/Retina-Atoh7-CutnRun(GSE156756)/Homer 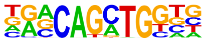 | 1e-7 | -1.755e+01 | 0.0000 | 42.0 | 17.21% | 6494.1 | 6.75% | motif file (matrix) | svg |
| 7 | Ap4(bHLH)/AML-Tfap4-ChIP-Seq(GSE45738)/Homer 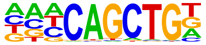 | 1e-7 | -1.716e+01 | 0.0000 | 58.0 | 23.77% | 10899.0 | 11.33% | motif file (matrix) | svg |
| 8 | Brn1(POU,Homeobox)/NPC-Brn1-ChIP-Seq(GSE35496)/Homer 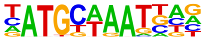 | 1e-7 | -1.649e+01 | 0.0000 | 30.0 | 12.30% | 3861.2 | 4.01% | motif file (matrix) | svg |
| 9 | BHLHA15(bHLH)/NIH3T3-BHLHB8.HA-ChIP-Seq(GSE119782)/Homer 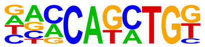 | 1e-7 | -1.613e+01 | 0.0000 | 63.0 | 25.82% | 12694.2 | 13.20% | motif file (matrix) | svg |
| 10 | TCF4(bHLH)/SHSY5Y-TCF4-ChIP-Seq(GSE96915)/Homer 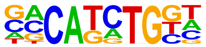 | 1e-6 | -1.578e+01 | 0.0000 | 68.0 | 27.87% | 14323.7 | 14.89% | motif file (matrix) | svg |
| 11 | Olig2(bHLH)/Neuron-Olig2-ChIP-Seq(GSE30882)/Homer | 1e-6 | -1.434e+01 | 0.0000 | 81.0 | 33.20% | 19035.9 | 19.79% | motif file (matrix) | svg |

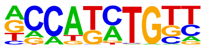

| | | | | | | | | | | | |
|---|---|---|---|---|---|---|---|---|---|---|---|
| 12 | 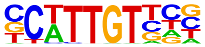 | Sox10(HMG)/SciaticNerve-Sox3-ChIP-Seq(GSE35132)/Homer | 1e-6 | -1.427e+01 | 0.0000 | 76.0 | 31.15% | 17445.3 | 18.14% | motif file (matrix) | svg |
| 13 | 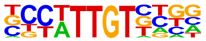 | Sox21(HMG)/ESC-SOX21-ChIP-Seq(GSE110505)/Homer | 1e-6 | -1.422e+01 | 0.0000 | 81.0 | 33.20% | 19091.3 | 19.85% | motif file (matrix) | svg |
| 14 | 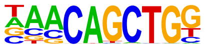 | Tcf21(bHLH)/ArterySmoothMuscle-Tcf21-ChIP-Seq(GSE61369)/Homer | 1e-6 | -1.418e+01 | 0.0000 | 45.0 | 18.44% | 8175.5 | 8.50% | motif file (matrix) | svg |
| 15 | 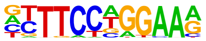 | STAT4(Stat)/CD4-Stat4-ChIP-Seq(GSE22104)/Homer | 1e-5 | -1.290e+01 | 0.0001 | 48.0 | 19.67% | 9431.4 | 9.81% | motif file (matrix) | svg |
| 16 | 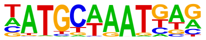 | Oct6(POU,Homeobox)/NPC-Pou3f1-ChIP-Seq(GSE35496)/Homer | 1e-5 | -1.166e+01 | 0.0003 | 31.0 | 12.70% | 5153.8 | 5.36% | motif file (matrix) | svg |
| 17 | 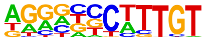 | Sox9(HMG)/Limb-SOX9-ChIP-Seq(GSE73225)/Homer | 1e-4 | -1.097e+01 | 0.0005 | 41.0 | 16.80% | 8093.3 | 8.42% | motif file (matrix) | svg |
| 18 | | SOX1(HMG)/NPC-SOX1-ChIP-Seq(GSE138215)/Homer | 1e-4 | -1.082e+01 | 0.0005 | 91.0 | 37.30% | 24246.7 | 25.21% | motif file (matrix) | svg |

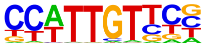

| | | | | | | | | | | | |
|---|---|---|---|---|---|---|---|---|---|---|---|
| 19 | 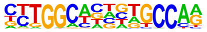 | NF1(CTF)/LNCAP-NF1-ChIP-Seq(Unpublished)/Homer | 1e-4 | -1.075e+01 | 0.0005 | 21.0 | 8.61% | 2912.7 | 3.03% | motif file (matrix) | svg |
| 20 | 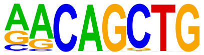 | MyoG(bHLH)/C2C12-MyoG-ChIP-Seq(GSE36024)/Homer | 1e-4 | -1.020e+01 | 0.0009 | 43.0 | 17.62% | 8956.5 | 9.31% | motif file (matrix) | svg |
| 21 | 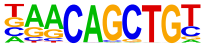 | Myf5(bHLH)/GM-Myf5-ChIP-Seq(GSE24852)/Homer | 1e-4 | -1.013e+01 | 0.0009 | 33.0 | 13.52% | 6129.6 | 6.37% | motif file (matrix) | svg |
| 22 | 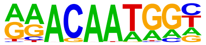 | Sox15(HMG)/CPA-Sox15-ChIP-Seq(GSE62909)/Homer | 1e-4 | -9.985e+00 | 0.0010 | 53.0 | 21.72% | 12081.9 | 12.56% | motif file (matrix) | svg |
| 23 | 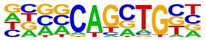 | Ascl1(bHLH)/NeuralTubes-Ascl1-ChIP-Seq(GSE55840)/Homer | 1e-4 | -9.763e+00 | 0.0012 | 57.0 | 23.36% | 13445.1 | 13.98% | motif file (matrix) | svg |
| 24 | 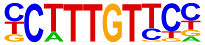 | Sox4(HMG)/proB-Sox4-ChIP-Seq(GSE50066)/Homer | 1e-3 | -8.910e+00 | 0.0027 | 40.0 | 16.39% | 8578.2 | 8.92% | motif file (matrix) | svg |
| 25 | | Oct4(POU,Homeobox)/mES-Oct4-ChIP-Seq(GSE11431)/Homer | 1e-3 | -8.894e+00 | 0.0027 | 32.0 | 13.11% | 6260.4 | 6.51% | motif file (matrix) | svg |

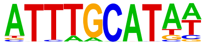

| | | | | | | | | | | | |
|---|---|---|---|---|---|---|---|---|---|---|---|
| 26 | ERG(ETS)/VCaP-ERG-ChIP-Seq(GSE14097)/Homer | 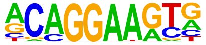 | 1e-3 | -8.847e+00 | 0.0027 | 57.0 | 23.36% | 13905.8 | 14.46% | motif file (matrix) | svg |
| 27 | Sox17(HMG)/Endoderm-Sox17-ChIP-Seq(GSE61475)/Homer | 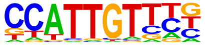 | 1e-3 | -8.573e+00 | 0.0033 | 37.0 | 15.16% | 7824.3 | 8.14% | motif file (matrix) | svg |
| 28 | Zic(Zf)/Cerebellum-ZIC1.2-ChIP-Seq(GSE60731)/Homer | 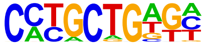 | 1e-3 | -8.421e+00 | 0.0037 | 32.0 | 13.11% | 6428.5 | 6.68% | motif file (matrix) | svg |
| 29 | SCL(bHLH)/HPC7-Scl-ChIP-Seq(GSE13511)/Homer | 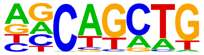 | 1e-3 | -8.384e+00 | 0.0037 | 137.0 | 56.15% | 43037.0 | 44.75% | motif file (matrix) | svg |
| 30 | OCT4-SOX2-TCF-NANOG(POU,Homeobox,HMG)/mES-Oct4-ChIP-Seq(GSE11431)/Homer | 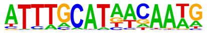 | 1e-3 | -8.369e+00 | 0.0037 | 16.0 | 6.56% | 2241.9 | 2.33% | motif file (matrix) | svg |
| 31 | Rfx5(HTH)/GM12878-Rfx5-ChIP-Seq(GSE31477)/Homer | 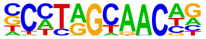 | 1e-3 | -8.368e+00 | 0.0037 | 17.0 | 6.97% | 2475.6 | 2.57% | motif file (matrix) | svg |
| 32 | MyoD(bHLH)/Myotube-MyoD-ChIP-Seq(GSE21614)/Homer | | 1e-3 | -7.881e+00 | 0.0056 | 31.0 | 12.70% | 6341.4 | 6.59% | motif file (matrix) | svg |

| | | | | | | | | | | | |
|---|---|---|---|---|---|---|---|---|---|---|---|
| | 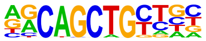 | | | | | | | | | | |
| 33 | 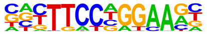 | Stat3+il21(Stat)/CD4-Stat3-ChIP-Seq(GSE19198)/Homer | 1e-3 | -7.821e+00 | 0.0057 | 29.0 | 11.89% | 5790.5 | 6.02% | motif file (matrix) | svg |
| 34 | 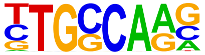 | NF1-halfsite(CTF)/LNCaP-NF1-ChIP-Seq(Unpublished)/Homer | 1e-3 | -7.706e+00 | 0.0062 | 63.0 | 25.82% | 16531.7 | 17.19% | motif file (matrix) | svg |
| 35 | 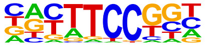 | Fli1(ETS)/CD8-FLI-ChIP-Seq(GSE20898)/Homer | 1e-3 | -7.508e+00 | 0.0074 | 37.0 | 15.16% | 8273.9 | 8.60% | motif file (matrix) | svg |
| 36 | 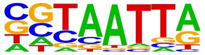 | DLX5(Homeobox)/BasalGanglia-Dlx5-ChIP-seq(GSE124936)/Homer | 1e-3 | -7.476e+00 | 0.0074 | 42.0 | 17.21% | 9829.1 | 10.22% | motif file (matrix) | svg |
| 37 | 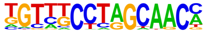 | Rfx6(HTH)/Min6b1-Rfx6.HA-ChIP-Seq(GSE62844)/Homer | 1e-3 | -7.403e+00 | 0.0078 | 37.0 | 15.16% | 8320.5 | 8.65% | motif file (matrix) | svg |
| 38 | 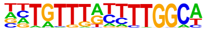 | NF1:FOXA1(CTF,Forkhead)/LNCAP-FOXA1-ChIP-Seq(GSE27824)/Homer | 1e-3 | -7.383e+00 | 0.0078 | 7.0 | 2.87% | 558.0 | 0.58% | motif file (matrix) | svg |
| 39 | | Tlx?(NR)/NPC-H3K4me1-ChIP-Seq(GSE16256)/Homer | 1e-3 | -7.331e+00 | 0.0079 | 19.0 | 7.79% | 3222.7 | 3.35% | motif file (matrix) | svg |

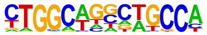

| | | | | | | | | | | | |
|---|---|---|---|---|---|---|---|---|---|---|---|
| 40 | 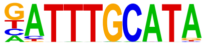 | Oct11(POU,Homeobox)/NCIH1048-POU2F3-ChIP-seq(GSE115123)/Homer | 1e-3 | -7.313e+00 | 0.0079 | 22.0 | 9.02% | 4019.1 | 4.18% | motif file (matrix) | svg |
| 41 | 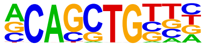 | Ptf1a(bHLH)/Panc1-Ptf1a-ChIP-Seq(GSE47459)/Homer | 1e-3 | -6.944e+00 | 0.0111 | 88.0 | 36.07% | 25798.0 | 26.82% | motif file (matrix) | svg |
| 42 | 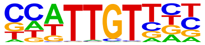 | Sox6(HMG)/Myotubes-Sox6-ChIP-Seq(GSE32627)/Homer | 1e-2 | -6.729e+00 | 0.0134 | 66.0 | 27.05% | 18190.2 | 18.91% | motif file (matrix) | svg |
| 43 | 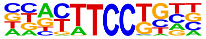 | Etv2(ETS)/ES-ER71-ChIP-Seq(GSE59402)/Homer | 1e-2 | -6.528e+00 | 0.0160 | 34.0 | 13.93% | 7806.5 | 8.12% | motif file (matrix) | svg |
| 44 | 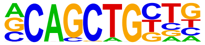 | Tcf12(bHLH)/GM12878-Tcf12-ChIP-Seq(GSE32465)/Homer | 1e-2 | -6.413e+00 | 0.0176 | 35.0 | 14.34% | 8170.7 | 8.50% | motif file (matrix) | svg |
| 45 | 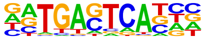 | Fosl2(bZIP)/3T3L1-Fosl2-ChIP-Seq(GSE56872)/Homer | 1e-2 | -6.341e+00 | 0.0185 | 15.0 | 6.15% | 2463.2 | 2.56% | motif file (matrix) | svg |
| 46 | | Ets1-distal(ETS)/CD4+-PolII-ChIP-Seq(Barski_et_al.)/Homer | 1e-2 | -6.322e+00 | 0.0185 | 15.0 | 6.15% | 2468.9 | 2.57% | motif file (matrix) | svg |

| | | | | | | | | | | | |
|---|---|---|---|---|---|---|---|---|---|---|---|
| | 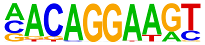 | | | | | | | | | | |
| 47 | 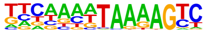 | MYNN(Zf)/HEK293-MYNN.eGFP-ChIP-Seq(Encode)/Homer | 1e-2 | -6.266e+00 | 0.0191 | 16.0 | 6.56% | 2739.6 | 2.85% | motif file (matrix) | svg |
| 48 | 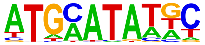 | Pit1(Homeobox)/GCrat-Pit1-ChIP-Seq(GSE58009)/Homer | 1e-2 | -6.144e+00 | 0.0211 | 50.0 | 20.49% | 13155.9 | 13.68% | motif file (matrix) | svg |
| 49 | 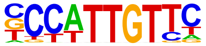 | Sox2(HMG)/mES-Sox2-ChIP-Seq(GSE11431)/Homer | 1e-2 | -6.138e+00 | 0.0211 | 38.0 | 15.57% | 9252.3 | 9.62% | motif file (matrix) | svg |
| 50 | 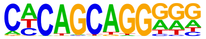 | Unknown-ESC-element(?)/mES-Nanog-ChIP-Seq(GSE11724)/Homer | 1e-2 | -6.132e+00 | 0.0211 | 22.0 | 9.02% | 4415.1 | 4.59% | motif file (matrix) | svg |
| 51 | 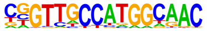 | RFX(HTH)/K562-RFX3-ChIP-Seq(SRA012198)/Homer | 1e-2 | -6.092e+00 | 0.0211 | 5.0 | 2.05% | 354.7 | 0.37% | motif file (matrix) | svg |
| 52 | 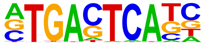 | AP-1(bZIP)/ThioMac-PU.1-ChIP-Seq(GSE21512)/Homer | 1e-2 | -6.059e+00 | 0.0212 | 30.0 | 12.30% | 6802.5 | 7.07% | motif file (matrix) | svg |
| 53 | | Zfp281(Zf)/ES-Zfp281-ChIP-Seq(GSE81042)/Homer | 1e-2 | -6.047e+00 | 0.0212 | 7.0 | 2.87% | 705.5 | 0.73% | motif file (matrix) | svg |

| | | | | | | | | | | | |
|---|---|---|---|---|---|---|---|---|---|---|---|
| | 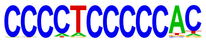 | | | | | | | | | | |
| 54 | 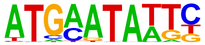 | Brn2(POU,Homeobox)/NPC-Brn2-ChIP-Seq(GSE35496)/Homer | 1e-2 | -5.993e+00 | 0.0218 | 9.0 | 3.69% | 1120.5 | 1.17% | motif file (matrix) | svg |
| 55 | 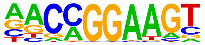 | ETV1(ETS)/GIST48-ETV1-ChIP-Seq(GSE22441)/Homer | 1e-2 | -5.959e+00 | 0.0222 | 43.0 | 17.62% | 10962.1 | 11.40% | motif file (matrix) | svg |
| 56 | 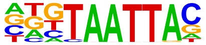 | Dlx3(Homeobox)/Kerainocytes-Dlx3-ChIP-Seq(GSE89884)/Homer | 1e-2 | -5.954e+00 | 0.0222 | 35.0 | 14.34% | 8401.6 | 8.74% | motif file (matrix) | svg |
| 57 | 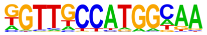 | X-box(HTH)/NPC-H3K4me1-ChIP-Seq(GSE16256)/Homer | 1e-2 | -5.901e+00 | 0.0227 | 7.0 | 2.87% | 724.7 | 0.75% | motif file (matrix) | svg |
| 58 | 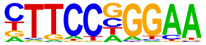 | Stat3(Stat)/mES-Stat3-ChIP-Seq(GSE11431)/Homer | 1e-2 | -5.729e+00 | 0.0265 | 19.0 | 7.79% | 3714.7 | 3.86% | motif file (matrix) | svg |
| 59 | 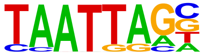 | Lhx2(Homeobox)/HFSC-Lhx2-ChIP-Seq(GSE48068)/Homer | 1e-2 | -5.649e+00 | 0.0282 | 49.0 | 20.08% | 13142.3 | 13.66% | motif file (matrix) | svg |
| 60 | | ETS1(ETS)/Jurkat-ETS1-ChIP-Seq(GSE17954)/Homer | 1e-2 | -5.630e+00 | 0.0282 | 34.0 | 13.93% | 8256.6 | 8.59% | motif file (matrix) | svg |

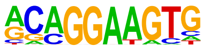

| | | | | | | | | | | | |
|---|---|---|---|---|---|---|---|---|---|---|---|
| 61 | 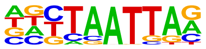 | Lhx1(Homeobox)/EmbryoCarcinoma-Lhx1-ChIP-Seq(GSE70957)/Homer | 1e-2 | -5.610e+00 | 0.0283 | 53.0 | 21.72% | 14520.1 | 15.10% | motif file (matrix) | svg |
| 62 | 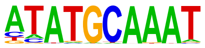 | Oct2(POU,Homeobox)/Bcell-Oct2-ChIP-Seq(GSE21512)/Homer | 1e-2 | -5.594e+00 | 0.0283 | 20.0 | 8.20% | 4044.3 | 4.21% | motif file (matrix) | svg |
| 63 | 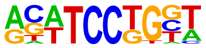 | SPDEF(ETS)/VCaP-SPDEF-ChIP-Seq(SRA014231)/Homer | 1e-2 | -5.593e+00 | 0.0283 | 35.0 | 14.34% | 8593.1 | 8.93% | motif file (matrix) | svg |
| 64 | 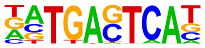 | BATF(bZIP)/Th17-BATF-ChIP-Seq(GSE39756)/Homer | 1e-2 | -5.414e+00 | 0.0328 | 26.0 | 10.66% | 5884.7 | 6.12% | motif file (matrix) | svg |
| 65 | 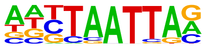 | Lhx3(Homeobox)/Neuron-Lhx3-ChIP-Seq(GSE31456)/Homer | 1e-2 | -5.403e+00 | 0.0328 | 71.0 | 29.10% | 20953.5 | 21.79% | motif file (matrix) | svg |
| 66 | 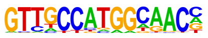 | Rfx2(HTH)/LoVo-RFX2-ChIP-Seq(GSE49402)/Homer | 1e-2 | -5.255e+00 | 0.0374 | 5.0 | 2.05% | 432.8 | 0.45% | motif file (matrix) | svg |
| 67 | | Mesp1(bHLH)/ESC-Mesp1-ChIP-Seq(GSE165102)/Homer | 1e-2 | -5.118e+00 | 0.0422 | 24.0 | 9.84% | 5414.5 | 5.63% | motif file (matrix) | svg |

| | | | | | | | | | | | |
|---|---|---|---|---|---|---|---|---|---|---|---|
| 68 | 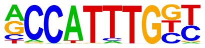 | Fra1(bZIP)/BT549-Fra1-ChIP-Seq(GSE46166)/Homer | 1e-2 | -5.106e+00 | 0.0422 | 22.0 | 9.02% | 4820.2 | 5.01% | motif file (matrix) | svg |
| 69 | 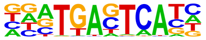 | Rfx1(HTH)/NPC-H3K4me1-ChIP-Seq(GSE16256)/Homer | 1e-2 | -5.034e+00 | 0.0446 | 9.0 | 3.69% | 1301.6 | 1.35% | motif file (matrix) | svg |
| 70 | 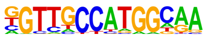 | Atf7(bZIP)/3T3L1-Atf7-ChIP-Seq(GSE56872)/Homer | 1e-2 | -5.023e+00 | 0.0446 | 21.0 | 8.61% | 4559.1 | 4.74% | motif file (matrix) | svg |
| 71 | 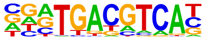 | JunB(bZIP)/DendriticCells-Junb-ChIP-Seq(GSE36099)/Homer | 1e-2 | -5.023e+00 | 0.0446 | 22.0 | 9.02% | 4856.3 | 5.05% | motif file (matrix) | svg |
| 72 | 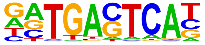 | Zic2(Zf)/ESC-Zic2-ChIP-Seq(SRP197560)/Homer | 1e-2 | -4.954e+00 | 0.0463 | 14.0 | 5.74% | 2596.0 | 2.70% | motif file (matrix) | svg |
| 73 | 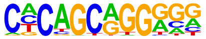 | Atf3(bZIP)/GBM-ATF3-ChIP-Seq(GSE33912)/Homer | 1e-2 | -4.705e+00 | 0.0585 | 25.0 | 10.25% | 5918.2 | 6.15% | motif file (matrix) | svg |
| 74 | 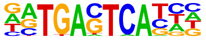 | ZFX(Zf)/mES-Zfx-ChIP-Seq(GSE11431)/Homer | 1e-2 | -4.632e+00 | 0.0621 | 36.0 | 14.75% | 9486.3 | 9.86% | motif file (matrix) | svg |

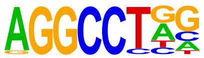

### Motifs found for: defined tumor border

### Homer Known Motif Enrichment Results (homerpos2)

[Homer *de novo* Motif Results](#)
[Gene Ontology Enrichment Results](#)
[Known Motif Enrichment Results (txt file)](#)
Total Target Sequences = 86, Total Background Sequences = 98018

| Rank | Motif | Name | P-value | log P-pvalue | q-value (Benjamini) | # Target Sequences with Motif | % of Targets Sequences with Motif | # Background Sequences with Motif | % of Background Sequences with Motif | Motif File | SVG |
|---|---|---|---|---|---|---|---|---|---|---|---|
| 1 | 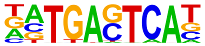 | BATF(bZIP)/Th17-BATF-ChIP-Seq(GSE39756)/Homer | 1e-5 | -1.216e+01 | 0.0025 | 17.0 | 19.77% | 5508.6 | 5.62% | motif file (matrix) | svg |
| 2 | 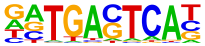 | JunB(bZIP)/DendriticCells-Junb-ChIP-Seq(GSE36099)/Homer | 1e-5 | -1.172e+01 | 0.0025 | 15.0 | 17.44% | 4494.6 | 4.58% | motif file (matrix) | svg |
| 3 | 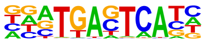 | Fra1(bZIP)/BT549-Fra1-ChIP-Seq(GSE46166)/Homer | 1e-5 | -1.168e+01 | 0.0025 | 15.0 | 17.44% | 4510.4 | 4.60% | motif file (matrix) | svg |
| 4 | | CEBP:AP1(bZIP)/ThioMac-CEBPb-ChIP-Seq(GSE21512)/Homer | 1e-4 | -9.547e+00 | 0.0084 | 19.0 | 22.09% | 8173.9 | 8.33% | motif file (matrix) | svg |

| | | | | | | | | | | | |
|---|---|---|---|---|---|---|---|---|---|---|---|
| | 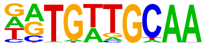 | | | | | | | | | | |
| 5 | 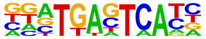 | Fos(bZIP)/TSC-Fos-ChIP-Seq(GSE110950)/Homer | 1e-4 | -9.466e+00 | 0.0084 | 14.0 | 16.28% | 4827.4 | 4.92% | motif file (matrix) | svg |
| 6 | 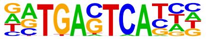 | Atf3(bZIP)/GBM-ATF3-ChIP-Seq(GSE33912)/Homer | 1e-4 | -9.339e+00 | 0.0084 | 15.0 | 17.44% | 5528.4 | 5.64% | motif file (matrix) | svg |
| 7 | 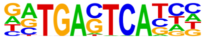 | Jun-AP1(bZIP)/K562-cJun-ChIP-Seq(GSE31477)/Homer | 1e-3 | -7.498e+00 | 0.0374 | 7.0 | 8.14% | 1609.5 | 1.64% | motif file (matrix) | svg |
| 8 | 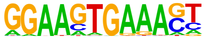 | PU.1:IRF8(ETS:IRF)/pDC-Irf8-ChIP-Seq(GSE66899)/Homer | 1e-2 | -5.718e+00 | 0.1939 | 6.0 | 6.98% | 1640.6 | 1.67% | motif file (matrix) | svg |
| 9 | 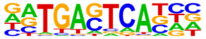 | Fosl2(bZIP)/3T3L1-Fosl2-ChIP-Seq(GSE56872)/Homer | 1e-2 | -5.520e+00 | 0.2101 | 7.0 | 8.14% | 2283.5 | 2.33% | motif file (matrix) | svg |
| 10 | 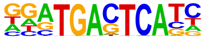 | Fra2(bZIP)/Striatum-Fra2-ChIP-Seq(GSE43429)/Homer | 1e-2 | -5.369e+00 | 0.2198 | 9.0 | 10.47% | 3638.1 | 3.71% | motif file (matrix) | svg |
| 11 | | Bach1(bZIP)/K562-Bach1-ChIP-Seq(GSE31477)/Homer | 1e-2 | -5.181e+00 | 0.2411 | 3.0 | 3.49% | 406.3 | 0.41% | motif file (matrix) | svg |

| | | | | | | | | | | |
|---|---|---|---|---|---|---|---|---|---|---|
| | 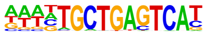 | | | | | | | | | |
| 12 | AP-1(bZIP)/ThioMac-PU.1-ChIP-Seq(GSE21512)/Homer | 1e-2 | -4.896e+00 | 0.2940 | 12.0 | 13.95% | 6161.4 | 6.28% | motif file (matrix) | svg |
| | 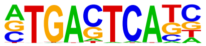 | | | | | | | | | |
| 13 | NF1:FOXA1(CTF,Forkhead)/LNCAP-FOXA1-ChIP-Seq(GSE27824)/Homer | 1e-2 | -4.829e+00 | 0.2940 | 3.0 | 3.49% | 462.8 | 0.47% | motif file (matrix) | svg |
| | 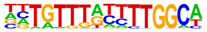 | | | | | | | | | |
| 14 | RFX(HTH)/K562-RFX3-ChIP-Seq(SRA012198)/Homer | 1e-2 | -4.748e+00 | 0.2940 | 3.0 | 3.49% | 476.3 | 0.49% | motif file (matrix) | svg |
| | 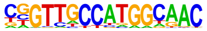 | | | | | | | | | |

### Motifs found for: diffuse tumor border

### Homer Known Motif Enrichment Results (homerpos3)

[Homer *de novo* Motif Results](#)
[Gene Ontology Enrichment Results](#)
[Known Motif Enrichment Results (txt file)](#)
Total Target Sequences = 709, Total Background Sequences = 96186

| Rank | Motif | Name | P-value | log P-pvalue | q-value (Benjamini) | # Target Sequences with Motif | % of Targets Sequences with Motif | # Background Sequences with Motif | % of Background Sequences with Motif | Motif File | SVG |
|---|---|---|---|---|---|---|---|---|---|---|---|
| 1 | 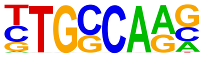 | NF1-halfsite(CTF)/LNCaP-NF1-ChIP-Seq(Unpublished)/Homer | 1e-27 | -6.260e+01 | 0.0000 | 250.0 | 35.26% | 17313.5 | 18.00% | motif file (matrix) | svg |
| 2 | 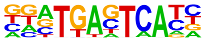 | Fos(bZIP)/TSC-Fos-ChIP-Seq(GSE110950)/Homer | 1e-16 | -3.883e+01 | 0.0000 | 95.0 | 13.40% | 4864.1 | 5.06% | motif file (matrix) | svg |
| 3 | 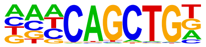 | Ap4(bHLH)/AML-Tfap4-ChIP-Seq(GSE45738)/Homer | 1e-16 | -3.835e+01 | 0.0000 | 166.0 | 23.41% | 11537.6 | 11.99% | motif file (matrix) | svg |
| 4 | | Oct6(POU,Homeobox)/NPC-Pou3f1-ChIP-Seq(GSE35496)/Homer | 1e-15 | -3.523e+01 | 0.0000 | 89.0 | 12.55% | 4645.0 | 4.83% | motif file (matrix) | svg |

| | | | | | | | | | | | |
|---|---|---|---|---|---|---|---|---|---|---|---|
| | 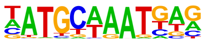 | | | | | | | | | | |
| 5 | 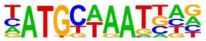 | Brn1(POU,Homeobox)/NPC-Brn1-ChIP-Seq(GSE35496)/Homer | 1e-15 | -3.513e+01 | 0.0000 | 74.0 | 10.44% | 3434.8 | 3.57% | motif file (matrix) | svg |
| 6 | 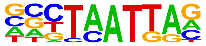 | Emx2(Homeobox)/Cortex-Emx2-ChIP-Seq(GSE183130)/Homer | 1e-14 | -3.271e+01 | 0.0000 | 171.0 | 24.12% | 12806.9 | 13.31% | motif file (matrix) | svg |
| 7 | 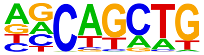 | SCL(bHLH)/HPC7-Scl-ChIP-Seq(GSE13511)/Homer | 1e-12 | -2.844e+01 | 0.0000 | 425.0 | 59.94% | 44720.4 | 46.48% | motif file (matrix) | svg |
| 8 | 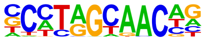 | Rfx5(HTH)/GM12878-Rfx5-ChIP-Seq(GSE31477)/Homer | 1e-12 | -2.813e+01 | 0.0000 | 58.0 | 8.18% | 2661.0 | 2.77% | motif file (matrix) | svg |
| 9 | 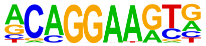 | ERG(ETS)/VCaP-ERG-ChIP-Seq(GSE14097)/Homer | 1e-11 | -2.745e+01 | 0.0000 | 180.0 | 25.39% | 14592.7 | 15.17% | motif file (matrix) | svg |
| 10 | 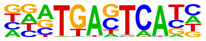 | Fra1(bZIP)/BT549-Fra1-ChIP-Seq(GSE46166)/Homer | 1e-11 | -2.738e+01 | 0.0000 | 80.0 | 11.28% | 4543.5 | 4.72% | motif file (matrix) | svg |
| 11 | | Tcf12(bHLH)/GM12878-Tcf12-ChIP-Seq(GSE32465)/Homer | 1e-11 | -2.709e+01 | 0.0000 | 125.0 | 17.63% | 8840.1 | 9.19% | motif file (matrix) | svg |

| | | | | | | | | | | | |
|---|---|---|---|---|---|---|---|---|---|---|---|
| | 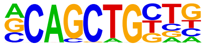 | | | | | | | | | | |
| 12 | 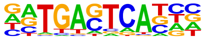 | Fosl2(bZIP)/3T3L1-Fosl2-ChIP-Seq(GSE56872)/Homer | 1e-11 | -2.648e+01 | 0.0000 | 53.0 | 7.48% | 2386.7 | 2.48% | motif file (matrix) | svg |
| 13 | 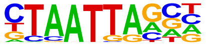 | Gsx2(Homeobox)/LGE-Gsx2.Flag-ChIP-Seq(GSE162589)/Homer | 1e-11 | -2.648e+01 | 0.0000 | 185.0 | 26.09% | 15313.6 | 15.92% | motif file (matrix) | svg |
| 14 | 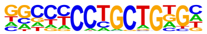 | Zic3(Zf)/mES-Zic3-ChIP-Seq(GSE37889)/Homer | 1e-11 | -2.608e+01 | 0.0000 | 75.0 | 10.58% | 4224.8 | 4.39% | motif file (matrix) | svg |
| 15 | 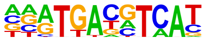 | CREB5(bZIP)/LNCaP-CREB5.V5-ChIP-Seq(GSE137775)/Homer | 1e-11 | -2.593e+01 | 0.0000 | 66.0 | 9.31% | 3467.4 | 3.60% | motif file (matrix) | svg |
| 16 | 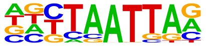 | Lhx1(Homeobox)/EmbryoCarcinoma-Lhx1-ChIP-Seq(GSE70957)/Homer | 1e-11 | -2.580e+01 | 0.0000 | 164.0 | 23.13% | 13119.8 | 13.64% | motif file (matrix) | svg |
| 17 | 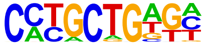 | Zic(Zf)/Cerebellum-ZIC1.2-ChIP-Seq(GSE60731)/Homer | 1e-11 | -2.538e+01 | 0.0000 | 101.0 | 14.25% | 6690.8 | 6.95% | motif file (matrix) | svg |
| 18 | | Atf3(bZIP)/GBM-ATF3-ChIP-Seq(GSE33912)/Homer | 1e-11 | -2.534e+01 | 0.0000 | 89.0 | 12.55% | 5559.8 | 5.78% | motif file (matrix) | svg |

| | | | | | | | | | | | |
|---|---|---|---|---|---|---|---|---|---|---|---|
| | 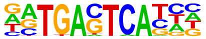 | | | | | | | | | | |
| 19 | 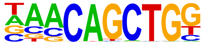 | Tcf21(bHLH)/ArterySmoothMuscle-Tcf21-ChIP-Seq(GSE61369)/Homer | 1e-10 | -2.530e+01 | 0.0000 | 123.0 | 17.35% | 8880.2 | 9.23% | motif file (matrix) | svg |
| 20 | 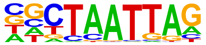 | LHX9(Homeobox)/Hct116-LHX9.V5-ChIP-Seq(GSE116822)/Homer | 1e-10 | -2.527e+01 | 0.0000 | 186.0 | 26.23% | 15641.7 | 16.26% | motif file (matrix) | svg |
| 21 | 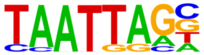 | Lhx2(Homeobox)/HFSC-Lhx2-ChIP-Seq(GSE48068)/Homer | 1e-10 | -2.525e+01 | 0.0000 | 152.0 | 21.44% | 11915.5 | 12.39% | motif file (matrix) | svg |
| 22 | 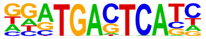 | Fra2(bZIP)/Striatum-Fra2-ChIP-Seq(GSE43429)/Homer | 1e-10 | -2.508e+01 | 0.0000 | 68.0 | 9.59% | 3708.0 | 3.85% | motif file (matrix) | svg |
| 23 | 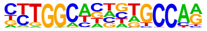 | NF1(CTF)/LNCAP-NF1-ChIP-Seq(Unpublished)/Homer | 1e-10 | -2.306e+01 | 0.0000 | 61.0 | 8.60% | 3285.8 | 3.42% | motif file (matrix) | svg |
| 24 | 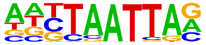 | Lhx3(Homeobox)/Neuron-Lhx3-ChIP-Seq(GSE31456)/Homer | 1e-9 | -2.293e+01 | 0.0000 | 212.0 | 29.90% | 19077.5 | 19.83% | motif file (matrix) | svg |
| 25 | | MyoD(bHLH)/Myotube-MyoD-ChIP-Seq(GSE21614)/Homer | 1e-9 | -2.277e+01 | 0.0000 | 100.0 | 14.10% | 6915.1 | 7.19% | motif file (matrix) | svg |

| | | | | | | | | | | | |
|---|---|---|---|---|---|---|---|---|---|---|---|
| | 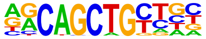 | | | | | | | | | | |
| 26 | | X-box(HTH)/NPC-H3K4me1-ChIP-Seq(GSE16256)/Homer | 1e-9 | -2.230e+01 | 0.0000 | 28.0 | 3.95% | 872.7 | 0.91% | motif file (matrix) | svg |
| | 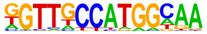 | | | | | | | | | | |
| 27 | | ETV1(ETS)/GIST48-ETV1-ChIP-Seq(GSE22441)/Homer | 1e-9 | -2.225e+01 | 0.0000 | 145.0 | 20.45% | 11657.2 | 12.12% | motif file (matrix) | svg |
| | 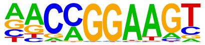 | | | | | | | | | | |
| 28 | | ETS1(ETS)/Jurkat-ETS1-ChIP-Seq(GSE17954)/Homer | 1e-9 | -2.134e+01 | 0.0000 | 115.0 | 16.22% | 8630.0 | 8.97% | motif file (matrix) | svg |
| | 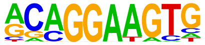 | | | | | | | | | | |
| 29 | | Tlx?(NR)/NPC-H3K4me1-ChIP-Seq(GSE16256)/Homer | 1e-8 | -2.046e+01 | 0.0000 | 64.0 | 9.03% | 3785.6 | 3.93% | motif file (matrix) | svg |
| | 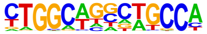 | | | | | | | | | | |
| 30 | | BATF(bZIP)/Th17-BATF-ChIP-Seq(GSE39756)/Homer | 1e-8 | -2.044e+01 | 0.0000 | 82.0 | 11.57% | 5456.1 | 5.67% | motif file (matrix) | svg |
| | 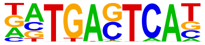 | | | | | | | | | | |
| 31 | | Jun-AP1(bZIP)/K562-cJun-ChIP-Seq(GSE31477)/Homer | 1e-8 | -2.010e+01 | 0.0000 | 38.0 | 5.36% | 1663.1 | 1.73% | motif file (matrix) | svg |
| | 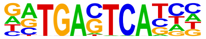 | | | | | | | | | | |
| 32 | | JunB(bZIP)/DendriticCells-Junb-ChIP-Seq(GSE36099)/Homer | 1e-8 | -1.969e+01 | 0.0000 | 71.0 | 10.01% | 4505.0 | 4.68% | motif file (matrix) | svg |

| | | | | | | | | | | | |
|---|---|---|---|---|---|---|---|---|---|---|---|
| | 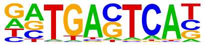 | | | | | | | | | | |
| 33 | 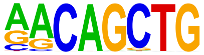 | MyoG(bHLH)/C2C12-MyoG-ChIP-Seq(GSE36024)/Homer | 1e-8 | -1.946e+01 | 0.0000 | 121.0 | 17.07% | 9554.5 | 9.93% | motif file (matrix) | svg |
| 34 | 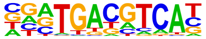 | Atf2(bZIP)/3T3L1-Atf2-ChIP-Seq(GSE56872)/Homer | 1e-8 | -1.935e+01 | 0.0000 | 51.0 | 7.19% | 2756.1 | 2.86% | motif file (matrix) | svg |
| 35 | 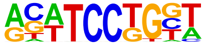 | SPDEF(ETS)/VCaP-SPDEF-ChIP-Seq(SRA014231)/Homer | 1e-8 | -1.915e+01 | 0.0000 | 114.0 | 16.08% | 8867.0 | 9.22% | motif file (matrix) | svg |
| 36 | 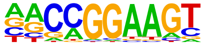 | GABPA(ETS)/Jurkat-GABPa-ChIP-Seq(GSE17954)/Homer | 1e-8 | -1.876e+01 | 0.0000 | 97.0 | 13.68% | 7165.5 | 7.45% | motif file (matrix) | svg |
| 37 | 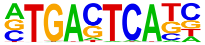 | AP-1(bZIP)/ThioMac-PU.1-ChIP-Seq(GSE21512)/Homer | 1e-8 | -1.858e+01 | 0.0000 | 89.0 | 12.55% | 6383.3 | 6.63% | motif file (matrix) | svg |
| 38 | 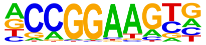 | ETV4(ETS)/HepG2-ETV4-ChIP-Seq(ENCODE)/Homer | 1e-7 | -1.831e+01 | 0.0000 | 110.0 | 15.51% | 8582.5 | 8.92% | motif file (matrix) | svg |
| 39 | | Oct11(POU,Homeobox)/NCIH1048-POU2F3-ChIP-seq(GSE115123)/Homer | 1e-7 | -1.771e+01 | 0.0000 | 58.0 | 8.18% | 3524.9 | 3.66% | motif file (matrix) | svg |

| | | | | | | | | | | | |
|---|---|---|---|---|---|---|---|---|---|---|---|
| | 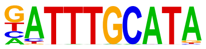 | | | | | | | | | | |
| 40 | 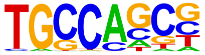 | HIC1(Zf)/Treg-ZBTB29-ChIP-Seq(GSE99889)/Homer | 1e-7 | -1.750e+01 | 0.0000 | 218.0 | 30.75% | 21043.7 | 21.87% | motif file (matrix) | svg |
| 41 | 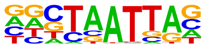 | En1(Homeobox)/SUM149-EN1-ChIP-Seq(GSE120957)/Homer | 1e-7 | -1.745e+01 | 0.0000 | 216.0 | 30.47% | 20815.3 | 21.64% | motif file (matrix) | svg |
| 42 | 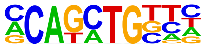 | Twist2(bHLH)/Myoblast-Twist2.Ty1-ChIP-Seq(GSE127998)/Homer | 1e-7 | -1.742e+01 | 0.0000 | 186.0 | 26.23% | 17257.9 | 17.94% | motif file (matrix) | svg |
| 43 | 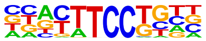 | Etv2(ETS)/ES-ER71-ChIP-Seq(GSE59402)/Homer | 1e-7 | -1.725e+01 | 0.0000 | 104.0 | 14.67% | 8125.0 | 8.45% | motif file (matrix) | svg |
| 44 | 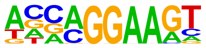 | EHF(ETS)/LoVo-EHF-ChIP-Seq(GSE49402)/Homer | 1e-7 | -1.708e+01 | 0.0000 | 142.0 | 20.03% | 12285.4 | 12.77% | motif file (matrix) | svg |
| 45 | 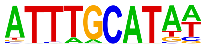 | Oct4(POU,Homeobox)/mES-Oct4-ChIP-Seq(GSE11431)/Homer | 1e-7 | -1.691e+01 | 0.0000 | 78.0 | 11.00% | 5519.8 | 5.74% | motif file (matrix) | svg |
| 46 | | Zic2(Zf)/ESC-Zic2-ChIP-Seq(SRP197560)/Homer | 1e-7 | -1.675e+01 | 0.0000 | 52.0 | 7.33% | 3083.5 | 3.21% | motif file (matrix) | svg |

| | | | | | | | | | | | |
|---|---|---|---|---|---|---|---|---|---|---|---|
| | 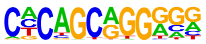 | | | | | | | | | | |
| 47 | 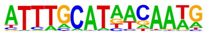 | OCT4-SOX2-TCF-NANOG(POU,Homeobox,HMG)/mES-Oct4-ChIP-Seq(GSE11431)/Homer | 1e-7 | -1.656e+01 | 0.0000 | 38.0 | 5.36% | 1907.1 | 1.98% | motif file (matrix) | svg |
| 48 | 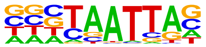 | DLX2(Homeobox)/BasalGanglia-Dlx2-ChIP-seq(GSE124936)/Homer | 1e-7 | -1.645e+01 | 0.0000 | 185.0 | 26.09% | 17372.7 | 18.06% | motif file (matrix) | svg |
| 49 | 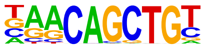 | Myf5(bHLH)/GM-Myf5-ChIP-Seq(GSE24852)/Homer | 1e-6 | -1.605e+01 | 0.0000 | 85.0 | 11.99% | 6346.8 | 6.60% | motif file (matrix) | svg |
| 50 | 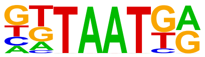 | Nkx6.1(Homeobox)/Islet-Nkx6.1-ChIP-Seq(GSE40975)/Homer | 1e-6 | -1.562e+01 | 0.0000 | 286.0 | 40.34% | 30007.3 | 31.19% | motif file (matrix) | svg |
| 51 | 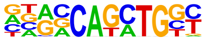 | Atoh1(bHLH)/Cerebellum-Atoh1-ChIP-Seq(GSE22111)/Homer | 1e-6 | -1.559e+01 | 0.0000 | 121.0 | 17.07% | 10253.2 | 10.66% | motif file (matrix) | svg |
| 52 | 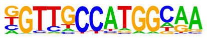 | Rfx1(HTH)/NPC-H3K4me1-ChIP-Seq(GSE16256)/Homer | 1e-6 | -1.543e+01 | 0.0000 | 32.0 | 4.51% | 1514.4 | 1.57% | motif file (matrix) | svg |
| 53 | | Fli1(ETS)/CD8-FLI-ChIP-Seq(GSE20898)/Homer | 1e-6 | -1.537e+01 | 0.0000 | 108.0 | 15.23% | 8877.2 | 9.23% | motif file (matrix) | svg |

| | | | | | | | | | | | |
|---|---|---|---|---|---|---|---|---|---|---|---|
| | 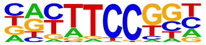 | | | | | | | | | | |
| 54 | 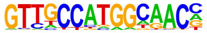 | Rfx2(HTH)/LoVo-RFX2-ChIP-Seq(GSE49402)/Homer | 1e-6 | -1.507e+01 | 0.0000 | 18.0 | 2.54% | 553.2 | 0.57% | motif file (matrix) | svg |
| 55 | 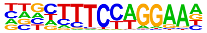 | Bcl6(Zf)/Liver-Bcl6-ChIP-Seq(GSE31578)/Homer | 1e-6 | -1.474e+01 | 0.0000 | 141.0 | 19.89% | 12671.6 | 13.17% | motif file (matrix) | svg |
| 56 | 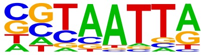 | DLX5(Homeobox)/BasalGanglia-Dlx5-ChIP-seq(GSE124936)/Homer | 1e-6 | -1.470e+01 | 0.0000 | 107.0 | 15.09% | 8895.5 | 9.25% | motif file (matrix) | svg |
| 57 | 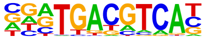 | Atf7(bZIP)/3T3L1-Atf7-ChIP-Seq(GSE56872)/Homer | 1e-6 | -1.430e+01 | 0.0000 | 66.0 | 9.31% | 4690.7 | 4.88% | motif file (matrix) | svg |
| 58 | 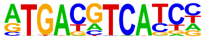 | c-Jun-CRE(bZIP)/K562-cJun-ChIP-Seq(GSE31477)/Homer | 1e-6 | -1.394e+01 | 0.0000 | 47.0 | 6.63% | 2923.6 | 3.04% | motif file (matrix) | svg |
| 59 | 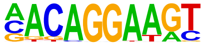 | Ets1-distal(ETS)/CD4+-PolII-ChIP-Seq(Barski_et_al.)/Homer | 1e-6 | -1.393e+01 | 0.0000 | 42.0 | 5.92% | 2476.0 | 2.57% | motif file (matrix) | svg |
| 60 | | STAT4(Stat)/CD4-Stat4-ChIP-Seq(GSE22104)/Homer | 1e-5 | -1.341e+01 | 0.0000 | 109.0 | 15.37% | 9368.8 | 9.74% | motif file (matrix) | svg |

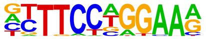

| | | | | | | | | | | | |
|---|---|---|---|---|---|---|---|---|---|---|---|
| 61 | 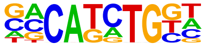 | TCF4(bHLH)/SHSY5Y-TCF4-ChIP-Seq(GSE96915)/Homer | 1e-5 | -1.317e+01 | 0.0000 | 158.0 | 22.28% | 15017.9 | 15.61% | motif file (matrix) | svg |
| 62 | 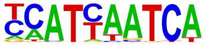 | Pdx1(Homeobox)/Islet-Pdx1-ChIP-Seq(SRA008281)/Homer | 1e-5 | -1.298e+01 | 0.0000 | 113.0 | 15.94% | 9902.3 | 10.29% | motif file (matrix) | svg |
| 63 | 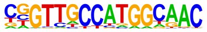 | RFX(HTH)/K562-RFX3-ChIP-Seq(SRA012198)/Homer | 1e-5 | -1.283e+01 | 0.0000 | 15.0 | 2.12% | 460.7 | 0.48% | motif file (matrix) | svg |
| 64 | 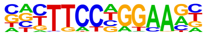 | Stat3+il21(Stat)/CD4-Stat3-ChIP-Seq(GSE19198)/Homer | 1e-5 | -1.282e+01 | 0.0000 | 77.0 | 10.86% | 6035.6 | 6.27% | motif file (matrix) | svg |
| 65 | 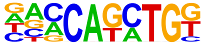 | BHLHA15(bHLH)/NIH3T3-BHLHB8.HA-ChIP-Seq(GSE119782)/Homer | 1e-5 | -1.281e+01 | 0.0000 | 144.0 | 20.31% | 13474.2 | 14.01% | motif file (matrix) | svg |
| 66 | 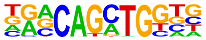 | Atoh7(bHLH)/Retina-Atoh7-CutnRun(GSE156756)/Homer | 1e-5 | -1.271e+01 | 0.0000 | 88.0 | 12.41% | 7219.1 | 7.50% | motif file (matrix) | svg |
| 67 | | JunD(bZIP)/K562-JunD-ChIP-Seq/Homer | 1e-5 | -1.268e+01 | 0.0000 | 15.0 | 2.12% | 466.8 | 0.49% | motif file (matrix) | svg |

| | | | | | | | | | | | |
|---|---|---|---|---|---|---|---|---|---|---|---|
| | 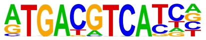 | | | | | | | | | | |
| 68 | 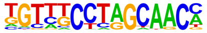 | Rfx6(HTH)/Min6b1-Rfx6.HA-ChIP-Seq(GSE62844)/Homer | 1e-5 | -1.252e+01 | 0.0000 | 104.0 | 14.67% | 8998.2 | 9.35% | motif file (matrix) | svg |
| 69 | 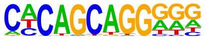 | Unknown-ESC-element(?)/mES-Nanog-ChIP-Seq(GSE11724)/Homer | 1e-5 | -1.244e+01 | 0.0000 | 65.0 | 9.17% | 4861.1 | 5.05% | motif file (matrix) | svg |
| 70 | 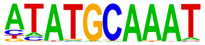 | Oct2(POU,Homeobox)/Bcell-Oct2-ChIP-Seq(GSE21512)/Homer | 1e-5 | -1.235e+01 | 0.0000 | 52.0 | 7.33% | 3583.1 | 3.72% | motif file (matrix) | svg |
| 71 | 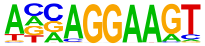 | ELF3(ETS)/PDAC-ELF3-ChIP-Seq(GSE64557)/Homer | 1e-5 | -1.226e+01 | 0.0000 | 84.0 | 11.85% | 6873.9 | 7.14% | motif file (matrix) | svg |
| 72 | 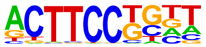 | Elf4(ETS)/BMDM-Elf4-ChIP-Seq(GSE88699)/Homer | 1e-5 | -1.222e+01 | 0.0000 | 101.0 | 14.25% | 8730.0 | 9.07% | motif file (matrix) | svg |
| 73 | 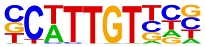 | Sox10(HMG)/SciaticNerve-Sox3-ChIP-Seq(GSE35132)/Homer | 1e-5 | -1.199e+01 | 0.0000 | 167.0 | 23.55% | 16405.5 | 17.05% | motif file (matrix) | svg |
| 74 | | Dlx3(Homeobox)/Kerainocytes-Dlx3-ChIP-Seq(GSE89884)/Homer | 1e-5 | -1.181e+01 | 0.0000 | 90.0 | 12.69% | 7607.4 | 7.91% | motif file (matrix) | svg |

| | | | | | | | | | | | |
|---|---|---|---|---|---|---|---|---|---|---|---|
| | 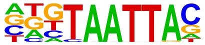 | | | | | | | | | | |
| 75 | | Ascl1(bHLH)/NeuralTubes-Ascl1-ChIP-Seq(GSE55840)/Homer | 1e-5 | -1.175e+01 | 0.0000 | 153.0 | 21.58% | 14806.6 | 15.39% | motif file (matrix) | svg |
| | 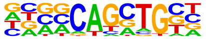 | | | | | | | | | | |
| 76 | | Egr1(Zf)/K562-Egr1-ChIP-Seq(GSE32465)/Homer | 1e-5 | -1.155e+01 | 0.0001 | 47.0 | 6.63% | 3204.2 | 3.33% | motif file (matrix) | svg |
| | 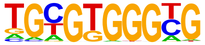 | | | | | | | | | | |
| 77 | | Stat3(Stat)/mES-Stat3-ChIP-Seq(GSE11431)/Homer | 1e-4 | -1.143e+01 | 0.0001 | 56.0 | 7.90% | 4105.3 | 4.27% | motif file (matrix) | svg |
| | 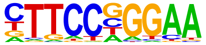 | | | | | | | | | | |
| 78 | | Hoxc10(Homeobox)/EB-Hoxc10.iFlag-ChIP-Seq(GSE142377)/Homer | 1e-4 | -1.131e+01 | 0.0001 | 149.0 | 21.02% | 14456.6 | 15.03% | motif file (matrix) | svg |
| | 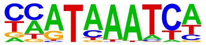 | | | | | | | | | | |
| 79 | | Six2(Homeobox)/NephronProgenitor-Six2-ChIP-Seq(GSE39837)/Homer | 1e-4 | -1.110e+01 | 0.0001 | 104.0 | 14.67% | 9310.0 | 9.68% | motif file (matrix) | svg |
| | 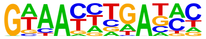 | | | | | | | | | | |
| 80 | | EWS:ERG-fusion(ETS)/CADO_ES1-EWS:ERG-ChIP-Seq(SRA014231)/Homer | 1e-4 | -1.099e+01 | 0.0001 | 86.0 | 12.13% | 7336.0 | 7.63% | motif file (matrix) | svg |
| | 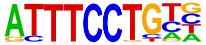 | | | | | | | | | | |
| 81 | | NeuroG2(bHLH)/Fibroblast-NeuroG2-ChIP-Seq(GSE75910)/Homer | 1e-4 | -1.062e+01 | 0.0001 | 153.0 | 21.58% | 15126.0 | 15.72% | motif file (matrix) | svg |

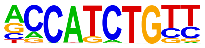

| | | | | | | | | | | | |
|---|---|---|---|---|---|---|---|---|---|---|---|
| 82 | 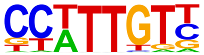 | Sox3(HMG)/NPC-Sox3-ChIP-Seq(GSE33059)/Homer | 1e-4 | -1.002e+01 | 0.0003 | 175.0 | 24.68% | 17970.9 | 18.68% | motif file (matrix) | svg |
| 83 | 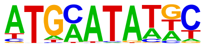 | Pit1(Homeobox)/GCrat-Pit1-ChIP-Seq(GSE58009)/Homer | 1e-4 | -9.977e+00 | 0.0003 | 118.0 | 16.64% | 11185.6 | 11.63% | motif file (matrix) | svg |
| 84 | 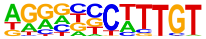 | Sox9(HMG)/Limb-SOX9-ChIP-Seq(GSE73225)/Homer | 1e-4 | -9.894e+00 | 0.0003 | 88.0 | 12.41% | 7790.2 | 8.10% | motif file (matrix) | svg |
| 85 | 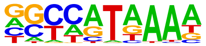 | Hoxd11(Homeobox)/ChickenMSG-Hoxd11.Flag-ChIP-Seq(GSE86088)/Homer | 1e-4 | -9.729e+00 | 0.0003 | 250.0 | 35.26% | 27455.7 | 28.54% | motif file (matrix) | svg |
| 86 | 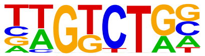 | Smad3(MAD)/NPC-Smad3-ChIP-Seq(GSE36673)/Homer | 1e-4 | -9.643e+00 | 0.0004 | 248.0 | 34.98% | 27233.9 | 28.31% | motif file (matrix) | svg |
| 87 | 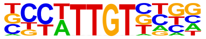 | Sox21(HMG)/ESC-SOX21-ChIP-Seq(GSE110505)/Homer | 1e-4 | -9.436e+00 | 0.0004 | 172.0 | 24.26% | 17798.9 | 18.50% | motif file (matrix) | svg |
| 88 | | Mef2d(MADS)/Retina-Mef2d-ChIP-Seq(GSE61391)/Homer | 1e-4 | -9.398e+00 | 0.0004 | 31.0 | 4.37% | 1957.4 | 2.03% | motif file (matrix) | svg |

| | | | | | | | | | | | |
|---|---|---|---|---|---|---|---|---|---|---|---|
| | 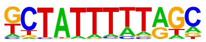 | | | | | | | | | | |
| 89 | 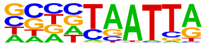 | DLX1(Homeobox)/BasalGanglia-Dlx1-ChIP-seq(GSE124936)/Homer | 1e-4 | -9.320e+00 | 0.0005 | 153.0 | 21.58% | 15526.5 | 16.14% | motif file (matrix) | svg |
| 90 | 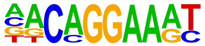 | EWS:FLI1-fusion(ETS)/SK_N_MC-EWS:FLI1-ChIP-Seq(SRA014231)/Homer | 1e-4 | -9.240e+00 | 0.0005 | 60.0 | 8.46% | 4876.8 | 5.07% | motif file (matrix) | svg |
| 91 | 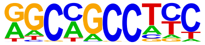 | ZNF91(Zf)/HEK-ZNF91.HA-ChIP-Seq(GSE162571)/Homer | 1e-3 | -9.015e+00 | 0.0006 | 73.0 | 10.30% | 6323.4 | 6.57% | motif file (matrix) | svg |
| 92 | 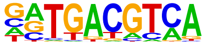 | Atf1(bZIP)/K562-ATF1-ChIP-Seq(GSE31477)/Homer | 1e-3 | -9.010e+00 | 0.0006 | 75.0 | 10.58% | 6544.6 | 6.80% | motif file (matrix) | svg |
| 93 | 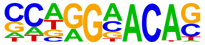 | AR-halfsite(NR)/LNCaP-AR-ChIP-Seq(GSE27824)/Homer | 1e-3 | -9.005e+00 | 0.0006 | 309.0 | 43.58% | 35408.2 | 36.80% | motif file (matrix) | svg |
| 94 | 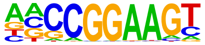 | ELF1(ETS)/Jurkat-ELF1-ChIP-Seq(SRA014231)/Homer | 1e-3 | -8.991e+00 | 0.0006 | 44.0 | 6.21% | 3264.6 | 3.39% | motif file (matrix) | svg |
| 95 | | MafA(bZIP)/Islet-MafA-ChIP-Seq(GSE30298)/Homer | 1e-3 | -8.513e+00 | 0.0010 | 76.0 | 10.72% | 6765.1 | 7.03% | motif file (matrix) | svg |

| | | | | | | | | | | | |
|---|---|---|---|---|---|---|---|---|---|---|---|
| | 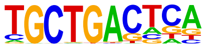 | | | | | | | | | | |
| 96 | 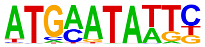 | Brn2(POU,Homeobox)/NPC-Brn2-ChIP-Seq(GSE35496)/Homer | 1e-3 | -8.494e+00 | 0.0010 | 20.0 | 2.82% | 1076.1 | 1.12% | motif file (matrix) | svg |
| 97 | 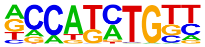 | Olig2(bHLH)/Neuron-Olig2-ChIP-Seq(GSE30882)/Homer | 1e-3 | -8.314e+00 | 0.0012 | 182.0 | 25.67% | 19431.4 | 20.20% | motif file (matrix) | svg |
| 98 | 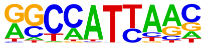 | Nanog(Homeobox)/mES-Nanog-ChIP-Seq(GSE11724)/Homer | 1e-3 | -7.875e+00 | 0.0018 | 378.0 | 53.31% | 45147.6 | 46.93% | motif file (matrix) | svg |
| 99 | 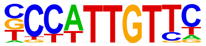 | Sox2(HMG)/mES-Sox2-ChIP-Seq(GSE11431)/Homer | 1e-3 | -7.812e+00 | 0.0019 | 90.0 | 12.69% | 8526.3 | 8.86% | motif file (matrix) | svg |
| 100 | 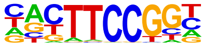 | Elk1(ETS)/Hela-Elk1-ChIP-Seq(GSE31477)/Homer | 1e-3 | -7.713e+00 | 0.0021 | 44.0 | 6.21% | 3469.8 | 3.61% | motif file (matrix) | svg |
| 101 | 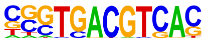 | CRE(bZIP)/Promoter/Homer | 1e-3 | -7.569e+00 | 0.0024 | 19.0 | 2.68% | 1072.7 | 1.11% | motif file (matrix) | svg |
| 102 | | Hoxc9(Homeobox)/Ainv15-Hoxc9-ChIP-Seq(GSE21812)/Homer | 1e-3 | -7.545e+00 | 0.0024 | 59.0 | 8.32% | 5102.4 | 5.30% | motif file (matrix) | svg |

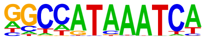

| | | | | | | | | | | | |
|---|---|---|---|---|---|---|---|---|---|---|---|
| 103 | 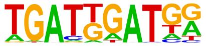 | HOXA1(Homeobox)/mES-Hoxa1-ChIP-Seq(SRP084292)/Homer | 1e-3 | -7.410e+00 | 0.0028 | 32.0 | 4.51% | 2300.9 | 2.39% | motif file (matrix) | svg |
| 104 | 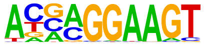 | ELF5(ETS)/T47D-ELF5-ChIP-Seq(GSE30407)/Homer | 1e-3 | -7.368e+00 | 0.0029 | 72.0 | 10.16% | 6587.7 | 6.85% | motif file (matrix) | svg |
| 105 | 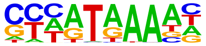 | Hoxa13(Homeobox)/ChickenMSG-Hoxa13.Flag-ChIP-Seq(GSE86088)/Homer | 1e-3 | -7.233e+00 | 0.0032 | 258.0 | 36.39% | 29558.1 | 30.72% | motif file (matrix) | svg |
| 106 | 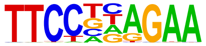 | STAT6(Stat)/Macrophage-Stat6-ChIP-Seq(GSE38377)/Homer | 1e-2 | -6.737e+00 | 0.0053 | 59.0 | 8.32% | 5281.5 | 5.49% | motif file (matrix) | svg |
| 107 | 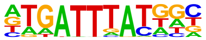 | Hoxd9(Homeobox)/EB-Hoxd9.HA-ChIP-Seq(GSE142377)/Homer | 1e-2 | -6.562e+00 | 0.0062 | 44.0 | 6.21% | 3679.1 | 3.82% | motif file (matrix) | svg |
| 108 | 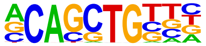 | Ptf1a(bHLH)/Panc1-Ptf1a-ChIP-Seq(GSE47459)/Homer | 1e-2 | -6.526e+00 | 0.0064 | 242.0 | 34.13% | 27827.4 | 28.92% | motif file (matrix) | svg |
| 109 | | PU.1(ETS)/ThioMac-PU.1-ChIP-Seq(GSE21512)/Homer | 1e-2 | -6.457e+00 | 0.0068 | 48.0 | 6.77% | 4132.4 | 4.30% | motif file (matrix) | svg |

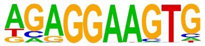

| | | | | | | | | | | | |
|---|---|---|---|---|---|---|---|---|---|---|---|
| 110 | 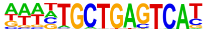 | Bach1(bZIP)/K562-Bach1-ChIP-Seq(GSE31477)/Homer | 1e-2 | -6.425e+00 | 0.0070 | 10.0 | 1.41% | 431.1 | 0.45% | motif file (matrix) | svg |
| 111 | 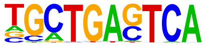 | Bach2(bZIP)/OCILy7-Bach2-ChIP-Seq(GSE44420)/Homer | 1e-2 | -6.105e+00 | 0.0095 | 21.0 | 2.96% | 1407.2 | 1.46% | motif file (matrix) | svg |
| 112 | 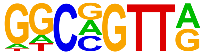 | MYB(HTH)/ERMYB-Myb-ChIPSeq(GSE22095)/Homer | 1e-2 | -6.057e+00 | 0.0099 | 144.0 | 20.31% | 15604.4 | 16.22% | motif file (matrix) | svg |
| 113 | 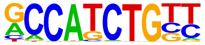 | NeuroD1(bHLH)/Islet-NeuroD1-ChIP-Seq(GSE30298)/Homer | 1e-2 | -6.038e+00 | 0.0100 | 77.0 | 10.86% | 7518.1 | 7.81% | motif file (matrix) | svg |
| 114 | 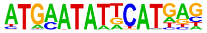 | OCT:OCT(POU,Homeobox)/NPC-Brn1-ChIP-Seq(GSE35496)/Homer | 1e-2 | -5.947e+00 | 0.0108 | 4.0 | 0.56% | 76.4 | 0.08% | motif file (matrix) | svg |
| 115 | 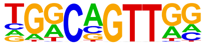 | AMYB(HTH)/Testes-AMYB-ChIP-Seq(GSE44588)/Homer | 1e-2 | -5.869e+00 | 0.0116 | 129.0 | 18.19% | 13827.3 | 14.37% | motif file (matrix) | svg |
| 116 | | RORgt(NR)/EL4-RORgt.Flag-ChIP-Seq(GSE56019)/Homer | 1e-2 | -5.843e+00 | 0.0117 | 16.0 | 2.26% | 974.2 | 1.01% | motif file (matrix) | svg |

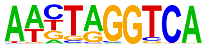

| | | | | | | | | | | | |
|---|---|---|---|---|---|---|---|---|---|---|---|
| 117 | 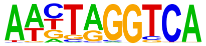 | RORgt(NR)/EL4-RORgt.Flag-ChIP-Seq(GSE56019)/Homer | 1e-2 | -5.843e+00 | 0.0117 | 16.0 | 2.26% | 974.2 | 1.01% | motif file (matrix) | svg |
| 118 | 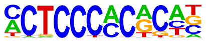 | WT1(Zf)/Kidney-WT1-ChIP-Seq(GSE90016)/Homer | 1e-2 | -5.832e+00 | 0.0117 | 41.0 | 5.78% | 3502.4 | 3.64% | motif file (matrix) | svg |
| 119 | 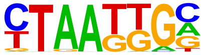 | Isl1(Homeobox)/Neuron-Isl1-ChIP-Seq(GSE31456)/Homer | 1e-2 | -5.800e+00 | 0.0120 | 185.0 | 26.09% | 20869.0 | 21.69% | motif file (matrix) | svg |
| 120 | 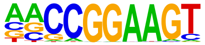 | ETS(ETS)/Promoter/Homer | 1e-2 | -5.794e+00 | 0.0120 | 28.0 | 3.95% | 2140.7 | 2.23% | motif file (matrix) | svg |
| 121 | 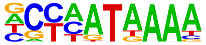 | Hoxd13(Homeobox)/ChickenMSG-Hoxd13.Flag-ChIP-Seq(GSE86088)/Homer | 1e-2 | -5.760e+00 | 0.0123 | 170.0 | 23.98% | 18988.2 | 19.74% | motif file (matrix) | svg |
| 122 | 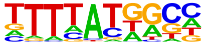 | Hoxa11(Homeobox)/ChickenMSG-Hoxa11.Flag-ChIP-Seq(GSE86088)/Homer | 1e-2 | -5.635e+00 | 0.0138 | 235.0 | 33.15% | 27378.8 | 28.46% | motif file (matrix) | svg |
| 123 | | STAT1(Stat)/HelaS3-STAT1-ChIP-Seq(GSE12782)/Homer | 1e-2 | -5.625e+00 | 0.0138 | 33.0 | 4.65% | 2687.8 | 2.79% | motif file (matrix) | svg |

| | | | | | | | | | | | |
|---|---|---|---|---|---|---|---|---|---|---|---|
| | 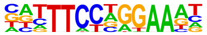 | | | | | | | | | | |
| 124 | 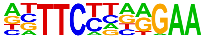 | STAT6(Stat)/CD4-Stat6-ChIP-Seq(GSE22104)/Homer | 1e-2 | -5.357e+00 | 0.0179 | 57.0 | 8.04% | 5399.1 | 5.61% | motif file (matrix) | svg |
| 125 | 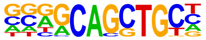 | Ascl2(bHLH)/ESC-Ascl2-ChIP-Seq(GSE97712)/Homer | 1e-2 | -5.343e+00 | 0.0181 | 110.0 | 15.51% | 11705.2 | 12.17% | motif file (matrix) | svg |
| 126 | 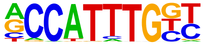 | Mesp1(bHLH)/ESC-Mesp1-ChIP-Seq(GSE165102)/Homer | 1e-2 | -5.102e+00 | 0.0228 | 56.0 | 7.90% | 5354.6 | 5.57% | motif file (matrix) | svg |
| 127 | 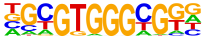 | Egr2(Zf)/Thymocytes-Egr2-ChIP-Seq(GSE34254)/Homer | 1e-2 | -5.068e+00 | 0.0234 | 11.0 | 1.55% | 608.1 | 0.63% | motif file (matrix) | svg |
| 128 | 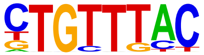 | Foxo1(Forkhead)/RAW-Foxo1-ChIP-Seq(Fan_et_al.)/Homer | 1e-2 | -5.019e+00 | 0.0244 | 174.0 | 24.54% | 19855.8 | 20.64% | motif file (matrix) | svg |
| 129 | 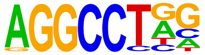 | ZFX(Zf)/mES-Zfx-ChIP-Seq(GSE11431)/Homer | 1e-2 | -4.882e+00 | 0.0277 | 101.0 | 14.25% | 10787.6 | 11.21% | motif file (matrix) | svg |
| 130 | | Arnt:Ahr(bHLH)/MCF7-Arnt-ChIP-Seq(Lo_et_al.)/Homer | 1e-2 | -4.882e+00 | 0.0277 | 47.0 | 6.63% | 4385.9 | 4.56% | motif file (matrix) | svg |

| | | | | | | | | | | | |
|---|---|---|---|---|---|---|---|---|---|---|---|
| | 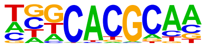 | | | | | | | | | | |
| 131 | 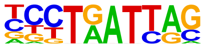 | Lhx6/Neurons-Lhx6-ChIP-seq(GSE85704)/Homer | 1e-2 | -4.852e+00 | 0.0282 | 120.0 | 16.93% | 13141.0 | 13.66% | motif file (matrix) | svg |
| 132 | 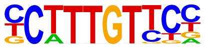 | Sox4(HMG)/proB-Sox4-ChIP-Seq(GSE50066)/Homer | 1e-2 | -4.703e+00 | 0.0324 | 79.0 | 11.14% | 8188.6 | 8.51% | motif file (matrix) | svg |
| 133 | 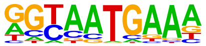 | Hoxa10(Homeobox)/ChickenMSG-Hoxa10.Flag-ChIP-Seq(GSE86088)/Homer | 1e-2 | -4.652e+00 | 0.0339 | 63.0 | 8.89% | 6302.7 | 6.55% | motif file (matrix) | svg |
| 134 | 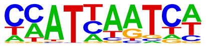 | Hoxc6(Homeobox)/EB-Hoxc6.iFlag-ChIP-Seq(GSE142377)/Homer | 1e-2 | -4.651e+00 | 0.0339 | 275.0 | 38.79% | 33194.3 | 34.50% | motif file (matrix) | svg |
| 135 | 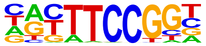 | Elk4(ETS)/Hela-Elk4-ChIP-Seq(GSE31477)/Homer | 1e-2 | -4.621e+00 | 0.0344 | 37.0 | 5.22% | 3326.4 | 3.46% | motif file (matrix) | svg |

### Motifs found for: gliomatosis cerebri

### Homer Known Motif Enrichment Results (homerpos4)

[Homer *de novo* Motif Results](#)
[Gene Ontology Enrichment Results](#)
[Known Motif Enrichment Results (txt file)](#)
Total Target Sequences = 55, Total Background Sequences = 97595

| Rank | Motif | Name | P-value | log P-pvalue | q-value (Benjamini) | # Target Sequences with Motif | % of Targets Sequences with Motif | # Background Sequences with Motif | % of Background Sequences with Motif | Motif File | SVG |
|---|---|---|---|---|---|---|---|---|---|---|---|
| 1 | 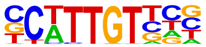 | Sox10(HMG)/SciaticNerve-Sox3-ChIP-Seq(GSE35132)/Homer | 1e-8 | -1.900e+01 | 0.0000 | 28.0 | 50.91% | 16278.8 | 16.68% | motif file (matrix) | svg |
| 2 | 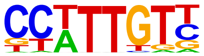 | Sox3(HMG)/NPC-Sox3-ChIP-Seq(GSE33059)/Homer | 1e-7 | -1.732e+01 | 0.0000 | 28.0 | 50.91% | 17528.1 | 17.96% | motif file (matrix) | svg |
| 3 | 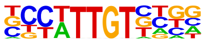 | Sox21(HMG)/ESC-SOX21-ChIP-Seq(GSE110505)/Homer | 1e-7 | -1.718e+01 | 0.0000 | 28.0 | 50.91% | 17641.5 | 18.07% | motif file (matrix) | svg |
| 4 | | SOX1(HMG)/NPC-SOX1-ChIP-Seq(GSE138215)/Homer | 1e-6 | -1.572e+01 | 0.0000 | 31.0 | 56.36% | 22842.3 | 23.40% | motif file (matrix) | svg |

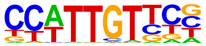

| | | | | | | | | | | |
|---|---|---|---|---|---|---|---|---|---|---|
| 5 | Sox6(HMG)/Myotubes-Sox6-ChIP-Seq(GSE32627)/Homer 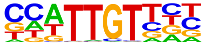 | 1e-5 | -1.255e+01 | 0.0003 | 24.0 | 43.64% | 16611.2 | 17.02% | motif file (matrix) | svg |
| 6 | X-box(HTH)/NPC-H3K4me1-ChIP-Seq(GSE16256)/Homer 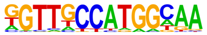 | 1e-5 | -1.156e+01 | 0.0008 | 6.0 | 10.91% | 863.2 | 0.88% | motif file (matrix) | svg |
| 7 | TCF4(bHLH)/SHSY5Y-TCF4-ChIP-Seq(GSE96915)/Homer 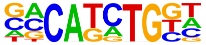 | 1e-5 | -1.153e+01 | 0.0008 | 22.0 | 40.00% | 15074.7 | 15.44% | motif file (matrix) | svg |
| 8 | Tcf21(bHLH)/ArterySmoothMuscle-Tcf21-ChIP-Seq(GSE61369)/Homer 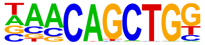 | 1e-4 | -1.058e+01 | 0.0015 | 16.0 | 29.09% | 9034.9 | 9.26% | motif file (matrix) | svg |
| 9 | Twist2(bHLH)/Myoblast-Twist2.Ty1-ChIP-Seq(GSE127998)/Homer 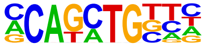 | 1e-4 | -1.050e+01 | 0.0015 | 23.0 | 41.82% | 17354.4 | 17.78% | motif file (matrix) | svg |
| 10 | Ap4(bHLH)/AML-Tfap4-ChIP-Seq(GSE45738)/Homer 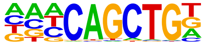 | 1e-4 | -1.041e+01 | 0.0015 | 18.0 | 32.73% | 11358.6 | 11.64% | motif file (matrix) | svg |
| 11 | Sox7(HMG)/ESC-Sox7-ChIP-Seq(GSE133899)/Homer | 1e-3 | -9.112e+00 | 0.0047 | 8.0 | 14.55% | 2634.2 | 2.70% | motif file (matrix) | svg |

| | | | | | | | | | | | |
|---|---|---|---|---|---|---|---|---|---|---|---|
| | 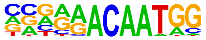 | | | | | | | | | | |
| 12 | 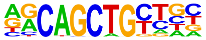 | MyoD(bHLH)/Myotube-MyoD-ChIP-Seq(GSE21614)/Homer | 1e-3 | -8.981e+00 | 0.0049 | 13.0 | 23.64% | 7099.5 | 7.27% | motif file (matrix) | svg |
| 13 | 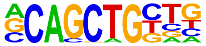 | Tcf12(bHLH)/GM12878-Tcf12-ChIP-Seq(GSE32465)/Homer | 1e-3 | -8.717e+00 | 0.0059 | 15.0 | 27.27% | 9425.9 | 9.66% | motif file (matrix) | svg |
| 14 | 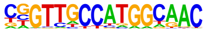 | RFX(HTH)/K562-RFX3-ChIP-Seq(SRA012198)/Homer | 1e-3 | -8.150e+00 | 0.0097 | 4.0 | 7.27% | 558.6 | 0.57% | motif file (matrix) | svg |
| 15 | 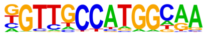 | Rfx1(HTH)/NPC-H3K4me1-ChIP-Seq(GSE16256)/Homer | 1e-3 | -8.096e+00 | 0.0097 | 6.0 | 10.91% | 1623.6 | 1.66% | motif file (matrix) | svg |
| 16 | 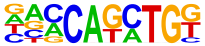 | BHLHA15(bHLH)/NIH3T3-BHLHB8.HA-ChIP-Seq(GSE119782)/Homer | 1e-3 | -8.053e+00 | 0.0097 | 18.0 | 32.73% | 13610.7 | 13.94% | motif file (matrix) | svg |
| 17 | 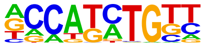 | Olig2(bHLH)/Neuron-Olig2-ChIP-Seq(GSE30882)/Homer | 1e-3 | -7.878e+00 | 0.0105 | 22.0 | 40.00% | 19078.5 | 19.55% | motif file (matrix) | svg |
| 18 | | Rfx2(HTH)/LoVo-RFX2-ChIP-Seq(GSE49402)/Homer | 1e-3 | -7.624e+00 | 0.0128 | 4.0 | 7.27% | 642.2 | 0.66% | motif file (matrix) | svg |

| | | | | | | | | | | | |
|---|---|---|---|---|---|---|---|---|---|---|---|
| | 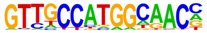 | | | | | | | | | | |
| 19 | 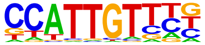 | Sox17(HMG)/Endoderm-Sox17-ChIP-Seq(GSE61475)/Homer | 1e-3 | -7.427e+00 | 0.0148 | 12.0 | 21.82% | 7239.9 | 7.42% | motif file (matrix) | svg |
| 20 | 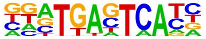 | Fos(bZIP)/TSC-Fos-ChIP-Seq(GSE110950)/Homer | 1e-3 | -7.388e+00 | 0.0148 | 9.0 | 16.36% | 4289.1 | 4.39% | motif file (matrix) | svg |
| 21 | 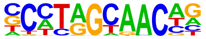 | Rfx5(HTH)/GM12878-Rfx5-ChIP-Seq(GSE31477)/Homer | 1e-3 | -7.189e+00 | 0.0170 | 7.0 | 12.73% | 2682.2 | 2.75% | motif file (matrix) | svg |
| 22 | 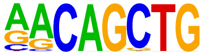 | MyoG(bHLH)/C2C12-MyoG-ChIP-Seq(GSE36024)/Homer | 1e-3 | -7.028e+00 | 0.0190 | 14.0 | 25.45% | 9839.5 | 10.08% | motif file (matrix) | svg |
| 23 | 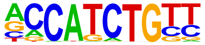 | NeuroG2(bHLH)/Fibroblast-NeuroG2-ChIP-Seq(GSE75910)/Homer | 1e-2 | -6.878e+00 | 0.0211 | 18.0 | 32.73% | 14987.2 | 15.35% | motif file (matrix) | svg |
| 24 | 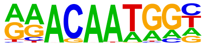 | Sox15(HMG)/CPA-Sox15-ChIP-Seq(GSE62909)/Homer | 1e-2 | -6.849e+00 | 0.0211 | 15.0 | 27.27% | 11221.9 | 11.50% | motif file (matrix) | svg |
| 25 | | Brn1(POU,Homeobox)/NPC-Brn1-ChIP-Seq(GSE35496)/Homer | 1e-2 | -6.654e+00 | 0.0243 | 7.0 | 12.73% | 2942.8 | 3.01% | motif file (matrix) | svg |

| | | | | | | | | | | | |
|---|---|---|---|---|---|---|---|---|---|---|---|
| | 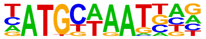 | | | | | | | | | | |
| 26 | 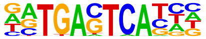 | Atf3(bZIP)/GBM-ATF3-ChIP-Seq(GSE33912)/Homer | 1e-2 | -6.494e+00 | 0.0275 | 9.0 | 16.36% | 4871.1 | 4.99% | motif file (matrix) | svg |
| 27 | 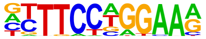 | STAT4(Stat)/CD4-Stat4-ChIP-Seq(GSE22104)/Homer | 1e-2 | -6.445e+00 | 0.0278 | 13.0 | 23.64% | 9255.9 | 9.48% | motif file (matrix) | svg |
| 28 | 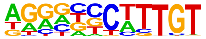 | Sox9(HMG)/Limb-SOX9-ChIP-Seq(GSE73225)/Homer | 1e-2 | -6.424e+00 | 0.0278 | 12.0 | 21.82% | 8121.4 | 8.32% | motif file (matrix) | svg |
| 29 | 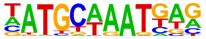 | Oct6(POU,Homeobox)/NPC-Pou3f1-ChIP-Seq(GSE35496)/Homer | 1e-2 | -6.303e+00 | 0.0298 | 8.0 | 14.55% | 4037.4 | 4.14% | motif file (matrix) | svg |
| 30 | 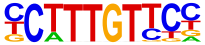 | Sox4(HMG)/proB-Sox4-ChIP-Seq(GSE50066)/Homer | 1e-2 | -6.183e+00 | 0.0325 | 12.0 | 21.82% | 8356.9 | 8.56% | motif file (matrix) | svg |
| 31 | 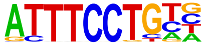 | EWS:ERG-fusion(ETS)/CADO_ES1-EWS:ERG-ChIP-Seq(SRA014231)/Homer | 1e-2 | -6.168e+00 | 0.0325 | 11.0 | 20.00% | 7237.7 | 7.41% | motif file (matrix) | svg |
| 32 | | Nanog(Homeobox)/mES-Nanog-ChIP-Seq(GSE11724)/Homer | 1e-2 | -5.890e+00 | 0.0408 | 35.0 | 63.64% | 43050.0 | 44.10% | motif file (matrix) | svg |

| | | | | | | | | | | | |
|---|---|---|---|---|---|---|---|---|---|---|---|
| | 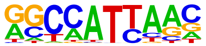 | | | | | | | | | | |
| 33 | 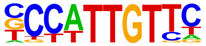 | Sox2(HMG)/mES-Sox2-ChIP-Seq(GSE11431)/Homer | 1e-2 | -5.877e+00 | 0.0408 | 12.0 | 21.82% | 8667.4 | 8.88% | motif file (matrix) | svg |
| 34 | 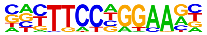 | Stat3+il21(Stat)/CD4-Stat3-ChIP-Seq(GSE19198)/Homer | 1e-2 | -5.778e+00 | 0.0430 | 10.0 | 18.18% | 6487.8 | 6.65% | motif file (matrix) | svg |
| 35 | 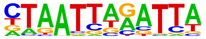 | Mixl1(Homeobox)/EpiBlast-Mixl1-ChIP-seq(GSE161164)/Homer | 1e-2 | -5.747e+00 | 0.0430 | 9.0 | 16.36% | 5439.1 | 5.57% | motif file (matrix) | svg |
| 36 | 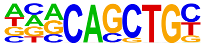 | E2A(bHLH)/proBcell-E2A-ChIP-Seq(GSE21978)/Homer | 1e-2 | -5.696e+00 | 0.0441 | 19.0 | 34.55% | 18004.1 | 18.44% | motif file (matrix) | svg |
| 37 | 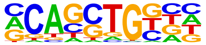 | HEB(bHLH)/mES-Heb-ChIP-Seq(GSE53233)/Homer | 1e-2 | -5.695e+00 | 0.0441 | 22.0 | 40.00% | 22337.3 | 22.88% | motif file (matrix) | svg |
| 38 | 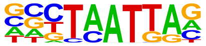 | Emx2(Homeobox)/Cortex-Emx2-ChIP-Seq(GSE183130)/Homer | 1e-2 | -5.482e+00 | 0.0517 | 14.0 | 25.45% | 11584.4 | 11.87% | motif file (matrix) | svg |
| 39 | | KLF6(Zf)/PDAC-KLF6-ChIP-Seq(GSE64557)/Homer | 1e-2 | -5.466e+00 | 0.0517 | 13.0 | 23.64% | 10342.6 | 10.60% | motif file (matrix) | svg |

| | | | | | | | | | | | |
|---|---|---|---|---|---|---|---|---|---|---|---|
| | 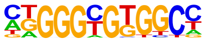 | | | | | | | | | | |
| 40 | | NFAT(RHD)/Jurkat-NFATC1-ChIP-Seq(Jolma_et_al.)/Homer | 1e-2 | -5.449e+00 | 0.0517 | 12.0 | 21.82% | 9133.5 | 9.36% | motif file (matrix) | svg |
| | 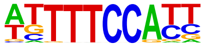 | | | | | | | | | | |
| 41 | | Lhx2(Homeobox)/HFSC-Lhx2-ChIP-Seq(GSE48068)/Homer | 1e-2 | -5.437e+00 | 0.0517 | 13.0 | 23.64% | 10377.7 | 10.63% | motif file (matrix) | svg |
| | 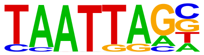 | | | | | | | | | | |
| 42 | | NFkB-p65-Rel(RHD)/ThioMac-LPS-Expression(GSE23622)/Homer | 1e-2 | -5.433e+00 | 0.0517 | 3.0 | 5.45% | 581.0 | 0.60% | motif file (matrix) | svg |
| | 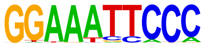 | | | | | | | | | | |
| 43 | | BATF(bZIP)/Th17-BATF-ChIP-Seq(GSE39756)/Homer | 1e-2 | -5.206e+00 | 0.0602 | 8.0 | 14.55% | 4834.8 | 4.95% | motif file (matrix) | svg |
| | 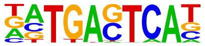 | | | | | | | | | | |
| 44 | | NFATC2(RHD)/Islets-NFATC2-ChIP-Seq(GSE158496)/Homer | 1e-2 | -5.203e+00 | 0.0602 | 21.0 | 38.18% | 21717.0 | 22.25% | motif file (matrix) | svg |
| | 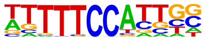 | | | | | | | | | | |
| 45 | | Ptf1a(bHLH)/Panc1-Ptf1a-ChIP-Seq(GSE47459)/Homer | 1e-2 | -5.194e+00 | 0.0602 | 26.0 | 47.27% | 29418.1 | 30.14% | motif file (matrix) | svg |
| | 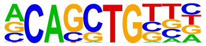 | | | | | | | | | | |
| 46 | | ETS1(ETS)/Jurkat-ETS1-ChIP-Seq(GSE17954)/Homer | 1e-2 | -5.193e+00 | 0.0602 | 12.0 | 21.82% | 9430.7 | 9.66% | motif file (matrix) | svg |

| | | | | | | | | | | | |
|---|---|---|---|---|---|---|---|---|---|---|---|
| | 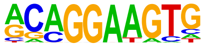 | | | | | | | | | | |
| 47 | 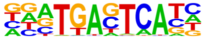 | Fra1(bZIP)/BT549-Fra1-ChIP-Seq(GSE46166)/Homer | 1e-2 | -5.121e+00 | 0.0602 | 7.0 | 12.73% | 3880.3 | 3.98% | motif file (matrix) | svg |
| 48 | 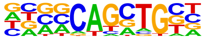 | Ascl1(bHLH)/NeuralTubes-Ascl1-ChIP-Seq(GSE55840)/Homer | 1e-2 | -5.047e+00 | 0.0632 | 17.0 | 30.91% | 16223.9 | 16.62% | motif file (matrix) | svg |
| 49 | 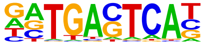 | JunB(bZIP)/DendriticCells-Junb-ChIP-Seq(GSE36099)/Homer | 1e-2 | -4.986e+00 | 0.0658 | 7.0 | 12.73% | 3980.1 | 4.08% | motif file (matrix) | svg |
| 50 | 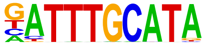 | Oct11(POU,Homeobox)/NCIH1048-POU2F3-ChIP-seq(GSE115123)/Homer | 1e-2 | -4.897e+00 | 0.0705 | 6.0 | 10.91% | 3067.1 | 3.14% | motif file (matrix) | svg |
| 51 | 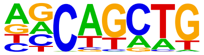 | SCL(bHLH)/HPC7-Scl-ChIP-Seq(GSE13511)/Homer | 1e-2 | -4.837e+00 | 0.0734 | 34.0 | 61.82% | 43634.2 | 44.70% | motif file (matrix) | svg |
| 52 | 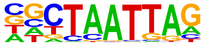 | LHX9(Homeobox)/Hct116-LHX9.V5-ChIP-Seq(GSE116822)/Homer | 1e-2 | -4.817e+00 | 0.0734 | 15.0 | 27.27% | 13829.9 | 14.17% | motif file (matrix) | svg |
| 53 | | Lhx1(Homeobox)/EmbryoCarcinoma-Lhx1-ChIP-Seq(GSE70957)/Homer | 1e-2 | -4.664e+00 | 0.0840 | 13.0 | 23.64% | 11386.7 | 11.67% | motif file (matrix) | svg |

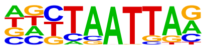

| | | | | | | | | | | |
|---|---|---|---|---|---|---|---|---|---|---|
| 54 | Brachyury(T-box)/Mesoendoderm-Brachyury-ChIP-exo(GSE54963)/Homer | 1e-2 | -4.620e+00 | 0.0861 | 5.0 | 9.09% | 2320.8 | 2.38% | motif file (matrix) | svg |

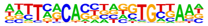

### Motifs found for: gray matter invasion

### Homer Known Motif Enrichment Results (homerpos5)

Homer *de novo* Motif Results
Gene Ontology Enrichment Results
Known Motif Enrichment Results (txt file)
Total Target Sequences = 66, Total Background Sequences = 97852

| Rank | Motif | Name | P-value | log P-pvalue | q-value (Benjamini) | # Target Sequences with Motif | % of Targets Sequences with Motif | # Background Sequences with Motif | % of Background Sequences with Motif | Motif File | SVG |
|---|---|---|---|---|---|---|---|---|---|---|---|
| 1 | 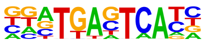 | Fos(bZIP)/TSC-Fos-ChIP-Seq(GSE110950)/Homer | 1e-5 | -1.195e+01 | 0.0031 | 14.0 | 21.21% | 5107.9 | 5.22% | motif file (matrix) | svg |
| 2 | 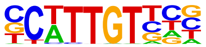 | Sox10(HMG)/SciaticNerve-Sox3-ChIP-Seq(GSE35132)/Homer | 1e-4 | -1.143e+01 | 0.0031 | 26.0 | 39.39% | 16460.5 | 16.81% | motif file (matrix) | svg |
| 3 | 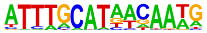 | OCT4-SOX2-TCF-NANOG(POU,Homeobox,HMG)/mES-Oct4-ChIP-Seq(GSE11431)/Homer | 1e-4 | -1.120e+01 | 0.0031 | 9.0 | 13.64% | 2139.8 | 2.19% | motif file (matrix) | svg |
| 4 | | Rfx2(HTH)/LoVo-RFX2-ChIP-Seq(GSE49402)/Homer | 1e-4 | -1.075e+01 | 0.0031 | 5.0 | 7.58% | 489.9 | 0.50% | motif file (matrix) | svg |

| | | | | | | | | | | |
|---|---|---|---|---|---|---|---|---|---|---|
| | 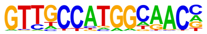 | | | | | | | | | |
| 5 | Oct4(POU,Homeobox)/mES-Oct4-ChIP-Seq(GSE11431)/Homer 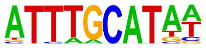 | 1e-3 | -9.078e+00 | 0.0108 | 13.0 | 19.70% | 5792.0 | 5.91% | motif file (matrix) | svg |
| 6 | Sox21(HMG)/ESC-SOX21-ChIP-Seq(GSE110505)/Homer 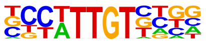 | 1e-3 | -9.047e+00 | 0.0108 | 25.0 | 37.88% | 17684.9 | 18.06% | motif file (matrix) | svg |
| 7 | Brn1(POU,Homeobox)/NPC-Brn1-ChIP-Seq(GSE35496)/Homer 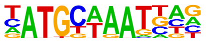 | 1e-3 | -8.869e+00 | 0.0108 | 10.0 | 15.15% | 3589.5 | 3.67% | motif file (matrix) | svg |
| 8 | Zic(Zf)/Cerebellum-ZIC1.2-ChIP-Seq(GSE60731)/Homer 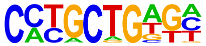 | 1e-3 | -8.744e+00 | 0.0108 | 14.0 | 21.21% | 6845.4 | 6.99% | motif file (matrix) | svg |
| 9 | Lhx2(Homeobox)/HFSC-Lhx2-ChIP-Seq(GSE48068)/Homer 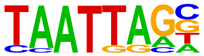 | 1e-3 | -8.741e+00 | 0.0108 | 20.0 | 30.30% | 12584.7 | 12.85% | motif file (matrix) | svg |
| 10 | X-box(HTH)/NPC-H3K4me1-ChIP-Seq(GSE16256)/Homer 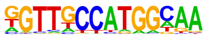 | 1e-3 | -8.645e+00 | 0.0108 | 5.0 | 7.58% | 766.2 | 0.78% | motif file (matrix) | svg |
| 11 | Oct11(POU,Homeobox)/NCIH1048-POU2F3-ChIP-seq(GSE115123)/Homer | 1e-3 | -8.632e+00 | 0.0108 | 10.0 | 15.15% | 3696.2 | 3.77% | motif file (matrix) | svg |

| | | | | | | | | | | | |
|---|---|---|---|---|---|---|---|---|---|---|---|
| | 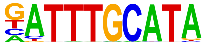 | | | | | | | | | | |
| 12 | 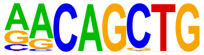 | MyoG(bHLH)/C2C12-MyoG-ChIP-Seq(GSE36024)/Homer | 1e-3 | -8.594e+00 | 0.0108 | 16.0 | 24.24% | 8762.4 | 8.95% | motif file (matrix) | svg |
| 13 | 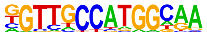 | Rfx1(HTH)/NPC-H3K4me1-ChIP-Seq(GSE16256)/Homer | 1e-3 | -7.865e+00 | 0.0139 | 6.0 | 9.09% | 1408.1 | 1.44% | motif file (matrix) | svg |
| 14 | 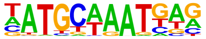 | Oct6(POU,Homeobox)/NPC-Pou3f1-ChIP-Seq(GSE35496)/Homer | 1e-3 | -7.846e+00 | 0.0139 | 11.0 | 16.67% | 4872.2 | 4.98% | motif file (matrix) | svg |
| 15 | 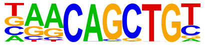 | Myf5(bHLH)/GM-Myf5-ChIP-Seq(GSE24852)/Homer | 1e-3 | -7.813e+00 | 0.0139 | 12.0 | 18.18% | 5724.7 | 5.85% | motif file (matrix) | svg |
| 16 | 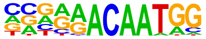 | Sox7(HMG)/ESC-Sox7-ChIP-Seq(GSE133899)/Homer | 1e-3 | -7.693e+00 | 0.0139 | 8.0 | 12.12% | 2695.4 | 2.75% | motif file (matrix) | svg |
| 17 | 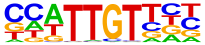 | Sox6(HMG)/Myotubes-Sox6-ChIP-Seq(GSE32627)/Homer | 1e-3 | -7.633e+00 | 0.0139 | 23.0 | 34.85% | 16992.7 | 17.35% | motif file (matrix) | svg |
| 18 | | LHX9(Homeobox)/Hct116-LHX9.V5-ChIP-Seq(GSE116822)/Homer | 1e-3 | -7.067e+00 | 0.0224 | 22.0 | 33.33% | 16521.8 | 16.87% | motif file (matrix) | svg |

| | | | | | | | | | | | |
|---|---|---|---|---|---|---|---|---|---|---|---|
| | 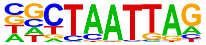 | | | | | | | | | | |
| 19 | 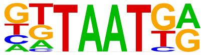 | Nkx6.1(Homeobox)/Islet-Nkx6.1-ChIP-Seq(GSE40975)/Homer | 1e-3 | -6.970e+00 | 0.0233 | 34.0 | 51.52% | 31627.7 | 32.30% | motif file (matrix) | svg |
| 20 | 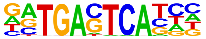 | Jun-AP1(bZIP)/K562-cJun-ChIP-Seq(GSE31477)/Homer | 1e-2 | -6.836e+00 | 0.0253 | 6.0 | 9.09% | 1717.1 | 1.75% | motif file (matrix) | svg |
| 21 | 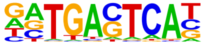 | JunB(bZIP)/DendriticCells-Junb-ChIP-Seq(GSE36099)/Homer | 1e-2 | -6.816e+00 | 0.0253 | 10.0 | 15.15% | 4665.6 | 4.76% | motif file (matrix) | svg |
| 22 | 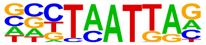 | Emx2(Homeobox)/Cortex-Emx2-ChIP-Seq(GSE183130)/Homer | 1e-2 | -6.778e+00 | 0.0253 | 19.0 | 28.79% | 13503.1 | 13.79% | motif file (matrix) | svg |
| 23 | 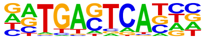 | Fosl2(bZIP)/3T3L1-Fosl2-ChIP-Seq(GSE56872)/Homer | 1e-2 | -6.621e+00 | 0.0273 | 7.0 | 10.61% | 2456.9 | 2.51% | motif file (matrix) | svg |
| 24 | 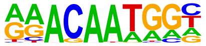 | Sox15(HMG)/CPA-Sox15-ChIP-Seq(GSE62909)/Homer | 1e-2 | -6.573e+00 | 0.0275 | 17.0 | 25.76% | 11571.4 | 11.82% | motif file (matrix) | svg |
| 25 | | TEAD3(TEA)/HepG2-TEAD3-ChIP-Seq(Encode)/Homer | 1e-2 | -6.493e+00 | 0.0286 | 18.0 | 27.27% | 12730.6 | 13.00% | motif file (matrix) | svg |

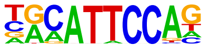

| | | | | | | | | | | | |
|---|---|---|---|---|---|---|---|---|---|---|---|
| 26 | 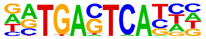 | Atf3(bZIP)/GBM-ATF3-ChIP-Seq(GSE33912)/Homer | 1e-2 | -6.437e+00 | 0.0291 | 11.0 | 16.67% | 5786.6 | 5.91% | motif file (matrix) | svg |
| 27 | 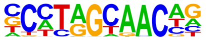 | Rfx5(HTH)/GM12878-Rfx5-ChIP-Seq(GSE31477)/Homer | 1e-2 | -6.330e+00 | 0.0311 | 7.0 | 10.61% | 2585.8 | 2.64% | motif file (matrix) | svg |
| 28 | 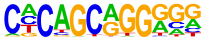 | Zic2(Zf)/ESC-Zic2-ChIP-Seq(SRP197560)/Homer | 1e-2 | -6.318e+00 | 0.0311 | 8.0 | 12.12% | 3340.2 | 3.41% | motif file (matrix) | svg |
| 29 | 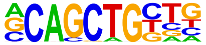 | Tcf12(bHLH)/GM12878-Tcf12-ChIP-Seq(GSE32465)/Homer | 1e-2 | -6.045e+00 | 0.0386 | 13.0 | 19.70% | 7993.3 | 8.16% | motif file (matrix) | svg |
| 30 | 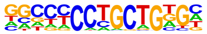 | Zic3(Zf)/mES-Zic3-ChIP-Seq(GSE37889)/Homer | 1e-2 | -6.009e+00 | 0.0387 | 9.0 | 13.64% | 4335.4 | 4.43% | motif file (matrix) | svg |
| 31 | 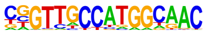 | RFX(HTH)/K562-RFX3-ChIP-Seq(SRA012198)/Homer | 1e-2 | -5.792e+00 | 0.0465 | 3.0 | 4.55% | 425.0 | 0.43% | motif file (matrix) | svg |
| 32 | | En1(Homeobox)/SUM149-EN1-ChIP-Seq(GSE120957)/Homer | 1e-2 | -5.743e+00 | 0.0473 | 25.0 | 37.88% | 21919.8 | 22.39% | motif file (matrix) | svg |

| | | | | | | | | | | | |
|---|---|---|---|---|---|---|---|---|---|---|---|
| 33 | 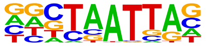 | STAT1(Stat)/HelaS3-STAT1-ChIP-Seq(GSE12782)/Homer | 1e-2 | -5.682e+00 | 0.0487 | 7.0 | 10.61% | 2905.9 | 2.97% | motif file (matrix) | svg |
| 34 | 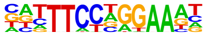 | Ap4(bHLH)/AML-Tfap4-ChIP-Seq(GSE45738)/Homer | 1e-2 | -5.640e+00 | 0.0493 | 15.0 | 22.73% | 10451.0 | 10.67% | motif file (matrix) | svg |
| 35 | 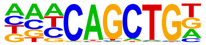 | Fra1(bZIP)/BT549-Fra1-ChIP-Seq(GSE46166)/Homer | 1e-2 | -5.465e+00 | 0.0571 | 9.0 | 13.64% | 4707.6 | 4.81% | motif file (matrix) | svg |
| 36 | 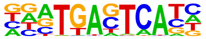 | Twist2(bHLH)/Myoblast-Twist2.Ty1-ChIP-Seq(GSE127998)/Homer | 1e-2 | -5.459e+00 | 0.0571 | 20.0 | 30.30% | 16286.3 | 16.63% | motif file (matrix) | svg |
| 37 | 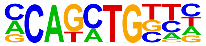 | NFkB-p65-Rel(RHD)/ThioMac-LPS-Expression(GSE23622)/Homer | 1e-2 | -5.341e+00 | 0.0611 | 3.0 | 4.55% | 500.8 | 0.51% | motif file (matrix) | svg |
| 38 | 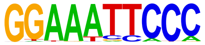 | Atf4(bZIP)/MEF-Atf4-ChIP-Seq(GSE35681)/Homer | 1e-2 | -5.267e+00 | 0.0641 | 7.0 | 10.61% | 3136.3 | 3.20% | motif file (matrix) | svg |
| 39 | 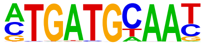 | Tcf21(bHLH)/ArterySmoothMuscle-Tcf21-ChIP-Seq(GSE61369)/Homer | 1e-2 | -5.052e+00 | 0.0774 | 12.0 | 18.18% | 7944.2 | 8.11% | motif file (matrix) | svg |

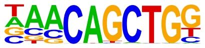

| | | | | | | | | | | | |
|---|---|---|---|---|---|---|---|---|---|---|---|
| 40 | 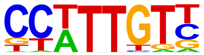 | Sox3(HMG)/NPC-Sox3-ChIP-Seq(GSE33059)/Homer | 1e-2 | -4.956e+00 | 0.0831 | 21.0 | 31.82% | 18226.5 | 18.61% | motif file (matrix) | svg |
| 41 | 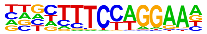 | Bcl6(Zf)/Liver-Bcl6-ChIP-Seq(GSE31578)/Homer | 1e-2 | -4.919e+00 | 0.0841 | 17.0 | 25.76% | 13549.0 | 13.84% | motif file (matrix) | svg |
| 42 | 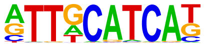 | Chop(bZIP)/MEF-Chop-ChIP-Seq(GSE35681)/Homer | 1e-2 | -4.859e+00 | 0.0872 | 6.0 | 9.09% | 2571.1 | 2.63% | motif file (matrix) | svg |
| 43 | 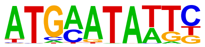 | Brn2(POU,Homeobox)/NPC-Brn2-ChIP-Seq(GSE35496)/Homer | 1e-2 | -4.835e+00 | 0.0872 | 4.0 | 6.06% | 1161.2 | 1.19% | motif file (matrix) | svg |
| 44 | 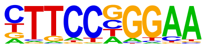 | Stat3(Stat)/mES-Stat3-ChIP-Seq(GSE11431)/Homer | 1e-2 | -4.744e+00 | 0.0933 | 8.0 | 12.12% | 4346.0 | 4.44% | motif file (matrix) | svg |
| 45 | 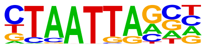 | Gsx2(Homeobox)/LGE-Gsx2.Flag-ChIP-Seq(GSE162589)/Homer | 1e-2 | -4.743e+00 | 0.0933 | 19.0 | 28.79% | 16146.2 | 16.49% | motif file (matrix) | svg |
| 46 | | MyoD(bHLH)/Myotube-MyoD-ChIP-Seq(GSE21614)/Homer | 1e-2 | -4.699e+00 | 0.0934 | 10.0 | 15.15% | 6280.3 | 6.41% | motif file (matrix) | svg |

| | | | | | | | | | | |
|---|---|---|---|---|---|---|---|---|---|---|
| 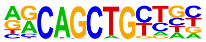 | | | | | | | | | | |
| 47 | Sox4(HMG)/proB-Sox4-ChIP-Seq(GSE50066)/Homer | 1e-2 | -4.689e+00 | 0.0934 | 12.0 | 18.18% | 8329.6 | 8.51% | motif file (matrix) | svg |
| 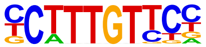 | | | | | | | | | | |
| 48 | Ascl2(bHLH)/ESC-Ascl2-ChIP-Seq(GSE97712)/Homer | 1e-2 | -4.665e+00 | 0.0934 | 14.0 | 21.21% | 10508.6 | 10.73% | motif file (matrix) | svg |
| 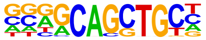 | | | | | | | | | | |

### Motifs found for: perineuronal satellitosis

### Homer Known Motif Enrichment Results (homerpos6)

Homer *de novo* Motif Results
Gene Ontology Enrichment Results
Known Motif Enrichment Results (txt file)
Total Target Sequences = 188, Total Background Sequences = 97404

| Rank | Motif | Name | P-value | log P-pvalue | q-value (Benjamini) | # Target Sequences with Motif | % of Targets Sequences with Motif | # Background Sequences with Motif | % of Background Sequences with Motif | Motif File | SVG |
|---|---|---|---|---|---|---|---|---|---|---|---|
| 1 | 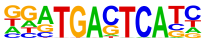 | Fra2(bZIP)/Striatum-Fra2-ChIP-Seq(GSE43429)/Homer | 1e-25 | -5.890e+01 | 0.0000 | 48.0 | 25.53% | 3642.9 | 3.74% | motif file (matrix) | svg |
| 2 | 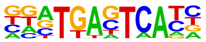 | Fos(bZIP)/TSC-Fos-ChIP-Seq(GSE110950)/Homer | 1e-25 | -5.852e+01 | 0.0000 | 54.0 | 28.72% | 4874.6 | 5.00% | motif file (matrix) | svg |
| 3 | 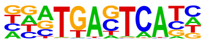 | Fra1(bZIP)/BT549-Fra1-ChIP-Seq(GSE46166)/Homer | 1e-25 | -5.803e+01 | 0.0000 | 52.0 | 27.66% | 4507.8 | 4.62% | motif file (matrix) | svg |
| 4 | | JunB(bZIP)/DendriticCells-Junb-ChIP-Seq(GSE36099)/Homer | 1e-22 | -5.288e+01 | 0.0000 | 49.0 | 26.06% | 4406.3 | 4.52% | motif file (matrix) | svg |

| | | | | | | | | | | | |
|---|---|---|---|---|---|---|---|---|---|---|---|
| | 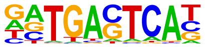 | | | | | | | | | | |
| 5 | 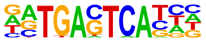 | Atf3(bZIP)/GBM-ATF3-ChIP-Seq(GSE33912)/Homer | 1e-22 | -5.141e+01 | 0.0000 | 53.0 | 28.19% | 5448.6 | 5.59% | motif file (matrix) | svg |
| 6 | 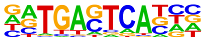 | Fosl2(bZIP)/3T3L1-Fosl2-ChIP-Seq(GSE56872)/Homer | 1e-21 | -5.060e+01 | 0.0000 | 36.0 | 19.15% | 2203.3 | 2.26% | motif file (matrix) | svg |
| 7 | 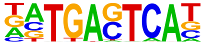 | BATF(bZIP)/Th17-BATF-ChIP-Seq(GSE39756)/Homer | 1e-21 | -4.837e+01 | 0.0000 | 51.0 | 27.13% | 5358.7 | 5.50% | motif file (matrix) | svg |
| 8 | 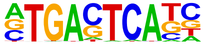 | AP-1(bZIP)/ThioMac-PU.1-ChIP-Seq(GSE21512)/Homer | 1e-20 | -4.731e+01 | 0.0000 | 54.0 | 28.72% | 6215.8 | 6.38% | motif file (matrix) | svg |
| 9 | 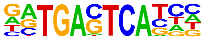 | Jun-AP1(bZIP)/K562-cJun-ChIP-Seq(GSE31477)/Homer | 1e-19 | -4.529e+01 | 0.0000 | 29.0 | 15.43% | 1490.9 | 1.53% | motif file (matrix) | svg |
| 10 | 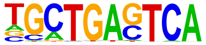 | Bach2(bZIP)/OCILy7-Bach2-ChIP-Seq(GSE44420)/Homer | 1e-8 | -1.978e+01 | 0.0000 | 16.0 | 8.51% | 1210.0 | 1.24% | motif file (matrix) | svg |
| 11 | | Oct4(POU,Homeobox)/mES-Oct4-ChIP-Seq(GSE11431)/Homer | 1e-5 | -1.345e+01 | 0.0001 | 31.0 | 16.49% | 6278.3 | 6.44% | motif file (matrix) | svg |

| | | | | | | | | | | | |
|---|---|---|---|---|---|---|---|---|---|---|---|
| | 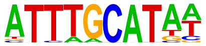 | | | | | | | | | | |
| 12 | 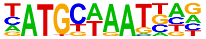 | Brn1(POU,Homeobox)/NPC-Brn1-ChIP-Seq(GSE35496)/Homer | 1e-5 | -1.311e+01 | 0.0001 | 23.0 | 12.23% | 3877.1 | 3.98% | motif file (matrix) | svg |
| 13 | 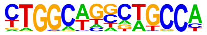 | Tlx?(NR)/NPC-H3K4me1-ChIP-Seq(GSE16256)/Homer | 1e-5 | -1.161e+01 | 0.0003 | 19.0 | 10.11% | 3066.0 | 3.15% | motif file (matrix) | svg |
| 14 | 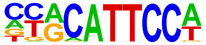 | TEAD1(TEAD)/HepG2-TEAD1-ChIP-Seq(Encode)/Homer | 1e-4 | -1.131e+01 | 0.0004 | 40.0 | 21.28% | 10261.6 | 10.53% | motif file (matrix) | svg |
| 15 | 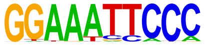 | NFkB-p65-Rel(RHD)/ThioMac-LPS-Expression(GSE23622)/Homer | 1e-4 | -1.054e+01 | 0.0008 | 7.0 | 3.72% | 437.8 | 0.45% | motif file (matrix) | svg |
| 16 | 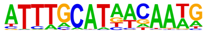 | OCT4-SOX2-TCF-NANOG(POU,Homeobox,HMG)/mES-Oct4-ChIP-Seq(GSE11431)/Homer | 1e-4 | -1.018e+01 | 0.0011 | 16.0 | 8.51% | 2531.8 | 2.60% | motif file (matrix) | svg |
| 17 | 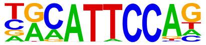 | TEAD3(TEA)/HepG2-TEAD3-ChIP-Seq(Encode)/Homer | 1e-4 | -9.702e+00 | 0.0017 | 43.0 | 22.87% | 12206.8 | 12.52% | motif file (matrix) | svg |
| 18 | | Sox3(HMG)/NPC-Sox3-ChIP-Seq(GSE33059)/Homer | 1e-3 | -8.900e+00 | 0.0036 | 58.0 | 30.85% | 18996.7 | 19.49% | motif file (matrix) | svg |

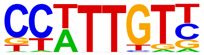

| | | | | | | | | | | | |
|---|---|---|---|---|---|---|---|---|---|---|---|
| 19 | 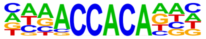 | RUNX2(Runt)/PCa-RUNX2-ChIP-Seq(GSE33889)/Homer | 1e-3 | -8.884e+00 | 0.0036 | 29.0 | 15.43% | 7211.6 | 7.40% | motif file (matrix) | svg |
| 20 | 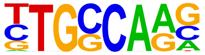 | NF1-halfsite(CTF)/LNCaP-NF1-ChIP-Seq(Unpublished)/Homer | 1e-3 | -8.827e+00 | 0.0036 | 51.0 | 27.13% | 16021.5 | 16.43% | motif file (matrix) | svg |
| 21 | 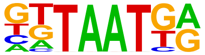 | Nkx6.1(Homeobox)/Islet-Nkx6.1-ChIP-Seq(GSE40975)/Homer | 1e-3 | -8.825e+00 | 0.0036 | 88.0 | 46.81% | 32928.3 | 33.78% | motif file (matrix) | svg |
| 22 | 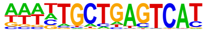 | NFE2L2(bZIP)/HepG2-NFE2L2-ChIP-Seq(Encode)/Homer | 1e-3 | -8.748e+00 | 0.0036 | 6.0 | 3.19% | 408.7 | 0.42% | motif file (matrix) | svg |
| 23 | 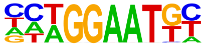 | TEAD(TEA)/Fibroblast-PU.1-ChIP-Seq(Unpublished)/Homer | 1e-3 | -8.736e+00 | 0.0036 | 30.0 | 15.96% | 7644.6 | 7.84% | motif file (matrix) | svg |
| 24 | 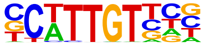 | Sox10(HMG)/SciaticNerve-Sox3-ChIP-Seq(GSE35132)/Homer | 1e-3 | -8.207e+00 | 0.0054 | 53.0 | 28.19% | 17290.5 | 17.74% | motif file (matrix) | svg |
| 25 | | Sox21(HMG)/ESC-SOX21-ChIP-Seq(GSE110505)/Homer | 1e-3 | -7.934e+00 | 0.0068 | 56.0 | 29.79% | 18797.6 | 19.28% | motif file (matrix) | svg |

| | | | | | | | | | | | |
|---|---|---|---|---|---|---|---|---|---|---|---|
| | 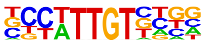 | | | | | | | | | | |
| 26 | 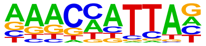 | Barx1(Homeobox)/Stomach-Barx1.3xFlag-ChIP-Seq(GSE69483)/Homer | 1e-3 | -7.925e+00 | 0.0068 | 25.0 | 13.30% | 6155.5 | 6.31% | motif file (matrix) | svg |
| 27 | 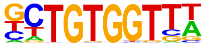 | RUNX-AML(Runt)/CD4+-PolII-ChIP-Seq(Barski_et_al.)/Homer | 1e-3 | -7.873e+00 | 0.0068 | 25.0 | 13.30% | 6177.5 | 6.34% | motif file (matrix) | svg |
| 28 | 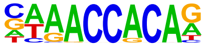 | RUNX(Runt)/HPC7-Runx1-ChIP-Seq(GSE22178)/Homer | 1e-3 | -7.854e+00 | 0.0068 | 24.0 | 12.77% | 5823.0 | 5.97% | motif file (matrix) | svg |
| 29 | 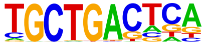 | MafA(bZIP)/Islet-MafA-ChIP-Seq(GSE30298)/Homer | 1e-3 | -7.362e+00 | 0.0103 | 24.0 | 12.77% | 6030.5 | 6.19% | motif file (matrix) | svg |
| 30 | 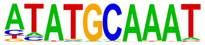 | Oct2(POU,Homeobox)/Bcell-Oct2-ChIP-Seq(GSE21512)/Homer | 1e-3 | -7.241e+00 | 0.0113 | 19.0 | 10.11% | 4299.3 | 4.41% | motif file (matrix) | svg |
| 31 | 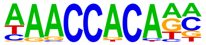 | RUNX1(Runt)/Jurkat-RUNX1-ChIP-Seq(GSE29180)/Homer | 1e-3 | -6.949e+00 | 0.0146 | 32.0 | 17.02% | 9307.7 | 9.55% | motif file (matrix) | svg |
| 32 | | Oct6(POU,Homeobox)/NPC-Pou3f1-ChIP-Seq(GSE35496)/Homer | 1e-2 | -6.486e+00 | 0.0225 | 21.0 | 11.17% | 5312.1 | 5.45% | motif file (matrix) | svg |

| | | | | | | | | | | | |
|---|---|---|---|---|---|---|---|---|---|---|---|
| | 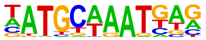 | | | | | | | | | | |
| 33 | 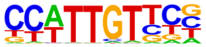 | SOX1(HMG)/NPC-SOX1-ChIP-Seq(GSE138215)/Homer | 1e-2 | -6.376e+00 | 0.0243 | 65.0 | 34.57% | 24163.6 | 24.79% | motif file (matrix) | svg |
| 34 | 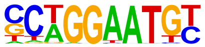 | TEAD2(TEA)/Py2T-Tead2-ChIP-Seq(GSE55709)/Homer | 1e-2 | -6.300e+00 | 0.0255 | 21.0 | 11.17% | 5395.1 | 5.53% | motif file (matrix) | svg |
| 35 | 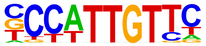 | Sox2(HMG)/mES-Sox2-ChIP-Seq(GSE11431)/Homer | 1e-2 | -6.126e+00 | 0.0295 | 30.0 | 15.96% | 8981.6 | 9.21% | motif file (matrix) | svg |
| 36 | 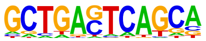 | MafK(bZIP)/C2C12-MafK-ChIP-Seq(GSE36030)/Homer | 1e-2 | -6.061e+00 | 0.0306 | 9.0 | 4.79% | 1463.9 | 1.50% | motif file (matrix) | svg |
| 37 | 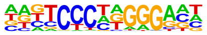 | EBF2(EBF)/BrownAdipose-EBF2-ChIP-Seq(GSE97114)/Homer | 1e-2 | -5.740e+00 | 0.0410 | 24.0 | 12.77% | 6814.8 | 6.99% | motif file (matrix) | svg |
| 38 | 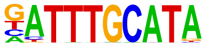 | Oct11(POU,Homeobox)/NCIH1048-POU2F3-ChIP-seq(GSE115123)/Homer | 1e-2 | -5.713e+00 | 0.0410 | 17.0 | 9.04% | 4190.4 | 4.30% | motif file (matrix) | svg |
| 39 | | Stat3+il21(Stat)/CD4-Stat3-ChIP-Seq(GSE19198)/Homer | 1e-2 | -5.589e+00 | 0.0452 | 21.0 | 11.17% | 5733.6 | 5.88% | motif file (matrix) | svg |

| | | | | | | | | | | | |
|---|---|---|---|---|---|---|---|---|---|---|---|
| | 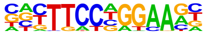 | | | | | | | | | | |
| 40 | 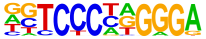 | EBF(EBF)/proBcell-EBF-ChIP-Seq(GSE21978)/Homer | 1e-2 | -5.378e+00 | 0.0545 | 8.0 | 4.26% | 1331.5 | 1.37% | motif file (matrix) | svg |
| 41 | 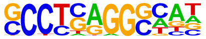 | AP-2gamma(AP2)/MCF7-TFAP2C-ChIP-Seq(GSE21234)/Homer | 1e-2 | -5.154e+00 | 0.0665 | 26.0 | 13.83% | 7954.7 | 8.16% | motif file (matrix) | svg |
| 42 | 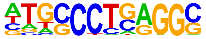 | AP-2alpha(AP2)/Hela-AP2alpha-ChIP-Seq(GSE31477)/Homer | 1e-2 | -5.036e+00 | 0.0730 | 21.0 | 11.17% | 6024.4 | 6.18% | motif file (matrix) | svg |
| 43 | 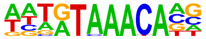 | FoxL2(Forkhead)/Ovary-FoxL2-ChIP-Seq(GSE60858)/Homer | 1e-2 | -4.973e+00 | 0.0760 | 31.0 | 16.49% | 10155.0 | 10.42% | motif file (matrix) | svg |
| 44 | 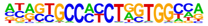 | CTCF(Zf)/CD4+-CTCF-ChIP-Seq(Barski_et_al.)/Homer | 1e-2 | -4.932e+00 | 0.0774 | 5.0 | 2.66% | 616.4 | 0.63% | motif file (matrix) | svg |
| 45 | 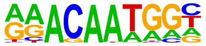 | Sox15(HMG)/CPA-Sox15-ChIP-Seq(GSE62909)/Homer | 1e-2 | -4.919e+00 | 0.0774 | 35.0 | 18.62% | 11908.7 | 12.22% | motif file (matrix) | svg |

### Motifs found for: perivascular invasion

### Homer Known Motif Enrichment Results (homerpos7)

Homer *de novo* Motif Results
Gene Ontology Enrichment Results
Known Motif Enrichment Results (txt file)
Total Target Sequences = 190, Total Background Sequences = 98238

| Rank | Motif | Name | P-value | log P-pvalue | q-value (Benjamini) | # Target Sequences with Motif | % of Targets Sequences with Motif | # Background Sequences with Motif | % of Background Sequences with Motif | Motif File | SVG |
|---|---|---|---|---|---|---|---|---|---|---|---|
| 1 | 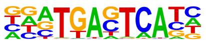 | Fra1(bZIP)/BT549-Fra1-ChIP-Seq(GSE46166)/Homer | 1e-16 | -3.803e+01 | 0.0000 | 45.0 | 23.68% | 5285.2 | 5.38% | motif file (matrix) | svg |
| 2 | 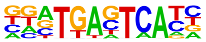 | Fos(bZIP)/TSC-Fos-ChIP-Seq(GSE110950)/Homer | 1e-16 | -3.790e+01 | 0.0000 | 47.0 | 24.74% | 5802.3 | 5.90% | motif file (matrix) | svg |
| 3 | 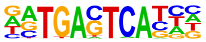 | Atf3(bZIP)/GBM-ATF3-ChIP-Seq(GSE33912)/Homer | 1e-16 | -3.769e+01 | 0.0000 | 49.0 | 25.79% | 6349.8 | 6.46% | motif file (matrix) | svg |
| 4 | | Fra2(bZIP)/Striatum-Fra2-ChIP-Seq(GSE43429)/Homer | 1e-15 | -3.644e+01 | 0.0000 | 39.0 | 20.53% | 4088.3 | 4.16% | motif file (matrix) | svg |

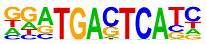

| | | | | | | | | | | | |
|---|---|---|---|---|---|---|---|---|---|---|---|
| 5 | 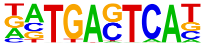 | BATF(bZIP)/Th17-BATF-ChIP-Seq(GSE39756)/Homer | 1e-14 | -3.451e+01 | 0.0000 | 47.0 | 24.74% | 6346.6 | 6.46% | motif file (matrix) | svg |
| 6 | 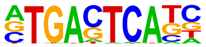 | AP-1(bZIP)/ThioMac-PU.1-ChIP-Seq(GSE21512)/Homer | 1e-14 | -3.316e+01 | 0.0000 | 49.0 | 25.79% | 7133.3 | 7.26% | motif file (matrix) | svg |
| 7 | 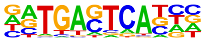 | Fosl2(bZIP)/3T3L1-Fosl2-ChIP-Seq(GSE56872)/Homer | 1e-14 | -3.265e+01 | 0.0000 | 29.0 | 15.26% | 2418.4 | 2.46% | motif file (matrix) | svg |
| 8 | 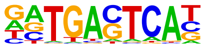 | JunB(bZIP)/DendriticCells-Junb-ChIP-Seq(GSE36099)/Homer | 1e-14 | -3.245e+01 | 0.0000 | 41.0 | 21.58% | 5122.9 | 5.21% | motif file (matrix) | svg |
| 9 | 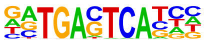 | Jun-AP1(bZIP)/K562-cJun-ChIP-Seq(GSE31477)/Homer | 1e-12 | -2.919e+01 | 0.0000 | 23.0 | 12.11% | 1636.0 | 1.66% | motif file (matrix) | svg |
| 10 | 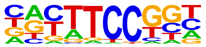 | Fli1(ETS)/CD8-FLI-ChIP-Seq(GSE20898)/Homer | 1e-6 | -1.552e+01 | 0.0000 | 37.0 | 19.47% | 7613.6 | 7.75% | motif file (matrix) | svg |
| 11 | | Sox3(HMG)/NPC-Sox3-ChIP-Seq(GSE33059)/Homer | 1e-6 | -1.460e+01 | 0.0000 | 74.0 | 38.95% | 22423.3 | 22.82% | motif file (matrix) | svg |

| | Logo | Motif | P-value | log P-value | q-value | # Target | % Target | # Background | % Background | | |
|---|---|---|---|---|---|---|---|---|---|---|---|
| | 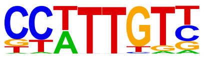 | | | | | | | | | | |
| 12 | 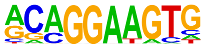 | ETS1(ETS)/Jurkat-ETS1-ChIP-Seq(GSE17954)/Homer | 1e-6 | -1.430e+01 | 0.0000 | 36.0 | 18.95% | 7664.9 | 7.80% | motif file (matrix) | svg |
| 13 | 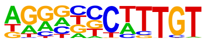 | Sox9(HMG)/Limb-SOX9-ChIP-Seq(GSE73225)/Homer | 1e-5 | -1.342e+01 | 0.0001 | 38.0 | 20.00% | 8659.0 | 8.81% | motif file (matrix) | svg |
| 14 | 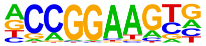 | ETV4(ETS)/HepG2-ETV4-ChIP-Seq(ENCODE)/Homer | 1e-5 | -1.257e+01 | 0.0001 | 31.0 | 16.32% | 6533.2 | 6.65% | motif file (matrix) | svg |
| 15 | 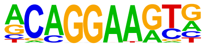 | ERG(ETS)/VCaP-ERG-ChIP-Seq(GSE14097)/Homer | 1e-5 | -1.185e+01 | 0.0002 | 48.0 | 25.26% | 13070.2 | 13.30% | motif file (matrix) | svg |
| 16 | 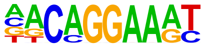 | EWS:FLI1-fusion(ETS)/SK_N_MC-EWS:FLI1-ChIP-Seq(SRA014231)/Homer | 1e-4 | -1.122e+01 | 0.0004 | 23.0 | 12.11% | 4344.7 | 4.42% | motif file (matrix) | svg |
| 17 | 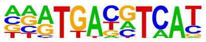 | CREB5(bZIP)/LNCaP-CREB5.V5-ChIP-Seq(GSE137775)/Homer | 1e-4 | -1.039e+01 | 0.0009 | 23.0 | 12.11% | 4583.3 | 4.66% | motif file (matrix) | svg |
| 18 | | Etv2(ETS)/ES-ER71-ChIP-Seq(GSE59402)/Homer | 1e-4 | -1.025e+01 | 0.0009 | 31.0 | 16.32% | 7357.2 | 7.49% | motif file (matrix) | svg |

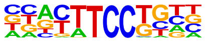

| | | | | | | | | | | | |
|---|---|---|---|---|---|---|---|---|---|---|---|
| 19 | 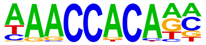 | RUNX1(Runt)/Jurkat-RUNX1-ChIP-Seq(GSE29180)/Homer | 1e-4 | -1.008e+01 | 0.0010 | 39.0 | 20.53% | 10412.8 | 10.60% | motif file (matrix) | svg |
| 20 | 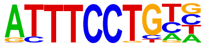 | EWS:ERG-fusion(ETS)/CADO_ES1-EWS:ERG-ChIP-Seq(SRA014231)/Homer | 1e-4 | -9.849e+00 | 0.0012 | 32.0 | 16.84% | 7882.4 | 8.02% | motif file (matrix) | svg |
| 21 | 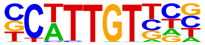 | Sox10(HMG)/SciaticNerve-Sox3-ChIP-Seq(GSE35132)/Homer | 1e-4 | -9.507e+00 | 0.0017 | 61.0 | 32.11% | 19836.0 | 20.18% | motif file (matrix) | svg |
| 22 | 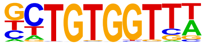 | RUNX-AML(Runt)/CD4+-PolII-ChIP-Seq(Barski_et_al.)/Homer | 1e-4 | -9.493e+00 | 0.0017 | 29.0 | 15.26% | 6940.9 | 7.06% | motif file (matrix) | svg |
| 23 | 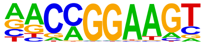 | ETV1(ETS)/GIST48-ETV1-ChIP-Seq(GSE22441)/Homer | 1e-4 | -9.410e+00 | 0.0017 | 37.0 | 19.47% | 9960.6 | 10.13% | motif file (matrix) | svg |
| 24 | 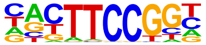 | Elk1(ETS)/Hela-Elk1-ChIP-Seq(GSE31477)/Homer | 1e-4 | -9.330e+00 | 0.0017 | 15.0 | 7.89% | 2429.9 | 2.47% | motif file (matrix) | svg |
| 25 | | RUNX2(Runt)/PCa-RUNX2-ChIP-Seq(GSE33889)/Homer | 1e-3 | -8.856e+00 | 0.0027 | 32.0 | 16.84% | 8316.0 | 8.46% | motif file (matrix) | svg |

| | | | | | | | | | | | |
|---|---|---|---|---|---|---|---|---|---|---|---|
| | 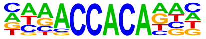 | | | | | | | | | | |
| 26 | | Sox21(HMG)/ESC-SOX21-ChIP-Seq(GSE110505)/Homer | 1e-3 | -8.720e+00 | 0.0030 | 65.0 | 34.21% | 22163.9 | 22.55% | motif file (matrix) | svg |
| | 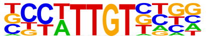 | | | | | | | | | | |
| 27 | | RUNX(Runt)/HPC7-Runx1-ChIP-Seq(GSE22178)/Homer | 1e-3 | -8.586e+00 | 0.0033 | 27.0 | 14.21% | 6587.6 | 6.70% | motif file (matrix) | svg |
| | 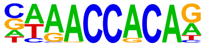 | | | | | | | | | | |
| 28 | | Sox6(HMG)/Myotubes-Sox6-ChIP-Seq(GSE32627)/Homer | 1e-3 | -8.243e+00 | 0.0044 | 63.0 | 33.16% | 21620.8 | 22.00% | motif file (matrix) | svg |
| | 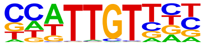 | | | | | | | | | | |
| 29 | | Atf2(bZIP)/3T3L1-Atf2-ChIP-Seq(GSE56872)/Homer | 1e-3 | -8.212e+00 | 0.0044 | 15.0 | 7.89% | 2696.6 | 2.74% | motif file (matrix) | svg |
| | 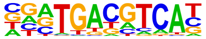 | | | | | | | | | | |
| 30 | | EHF(ETS)/LoVo-EHF-ChIP-Seq(GSE49402)/Homer | 1e-3 | -7.996e+00 | 0.0053 | 40.0 | 21.05% | 11902.3 | 12.11% | motif file (matrix) | svg |
| | 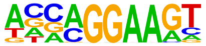 | | | | | | | | | | |
| 31 | | Atoh7(bHLH)/Retina-Atoh7-CutnRun(GSE156756)/Homer | 1e-3 | -7.911e+00 | 0.0056 | 22.0 | 11.58% | 5077.5 | 5.17% | motif file (matrix) | svg |
| | 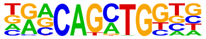 | | | | | | | | | | |
| 32 | | Atf7(bZIP)/3T3L1-Atf7-ChIP-Seq(GSE56872)/Homer | 1e-3 | -7.148e+00 | 0.0116 | 19.0 | 10.00% | 4321.6 | 4.40% | motif file (matrix) | svg |

| | | | | | | | | | | | |
|---|---|---|---|---|---|---|---|---|---|---|---|
| | 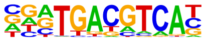 | | | | | | | | | | |
| 33 | 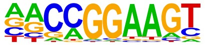 | GABPA(ETS)/Jurkat-GABPa-ChIP-Seq(GSE17954)/Homer | 1e-2 | -6.762e+00 | 0.0165 | 23.0 | 12.11% | 5911.0 | 6.01% | motif file (matrix) | svg |
| 34 | 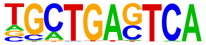 | Bach2(bZIP)/OCILy7-Bach2-ChIP-Seq(GSE44420)/Homer | 1e-2 | -6.710e+00 | 0.0169 | 9.0 | 4.74% | 1325.2 | 1.35% | motif file (matrix) | svg |
| 35 | 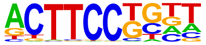 | Elf4(ETS)/BMDM-Elf4-ChIP-Seq(GSE88699)/Homer | 1e-2 | -6.563e+00 | 0.0190 | 27.0 | 14.21% | 7524.3 | 7.66% | motif file (matrix) | svg |
| 36 | 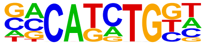 | TCF4(bHLH)/SHSY5Y-TCF4-ChIP-Seq(GSE96915)/Homer | 1e-2 | -6.541e+00 | 0.0190 | 40.0 | 21.05% | 12822.0 | 13.05% | motif file (matrix) | svg |
| 37 | 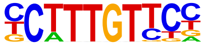 | Sox4(HMG)/proB-Sox4-ChIP-Seq(GSE50066)/Homer | 1e-2 | -6.409e+00 | 0.0210 | 32.0 | 16.84% | 9594.9 | 9.76% | motif file (matrix) | svg |
| 38 | 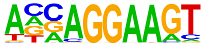 | ELF3(ETS)/PDAC-ELF3-ChIP-Seq(GSE64557)/Homer | 1e-2 | -6.346e+00 | 0.0218 | 25.0 | 13.16% | 6865.0 | 6.99% | motif file (matrix) | svg |
| 39 | | NeuroD1(bHLH)/Islet-NeuroD1-ChIP-Seq(GSE30298)/Homer | 1e-2 | -6.237e+00 | 0.0237 | 22.0 | 11.58% | 5780.3 | 5.88% | motif file (matrix) | svg |

| | | | | | | | | | | | |
|---|---|---|---|---|---|---|---|---|---|---|---|
| | 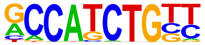 | | | | | | | | | | |
| 40 | 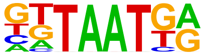 | Nkx6.1(Homeobox)/Islet-Nkx6.1-ChIP-Seq(GSE40975)/Homer | 1e-2 | -6.180e+00 | 0.0244 | 98.0 | 51.58% | 40298.3 | 41.00% | motif file (matrix) | svg |
| 41 | 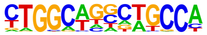 | Tlx?(NR)/NPC-H3K4me1-ChIP-Seq(GSE16256)/Homer | 1e-2 | -6.077e+00 | 0.0264 | 12.0 | 6.32% | 2355.6 | 2.40% | motif file (matrix) | svg |
| 42 | 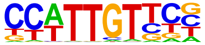 | SOX1(HMG)/NPC-SOX1-ChIP-Seq(GSE138215)/Homer | 1e-2 | -6.052e+00 | 0.0264 | 72.0 | 37.89% | 27688.2 | 28.17% | motif file (matrix) | svg |
| 43 | 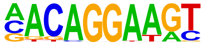 | Ets1-distal(ETS)/CD4+-PolII-ChIP-Seq(Barski_et_al.)/Homer | 1e-2 | -5.992e+00 | 0.0274 | 12.0 | 6.32% | 2380.0 | 2.42% | motif file (matrix) | svg |
| 44 | 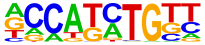 | Olig2(bHLH)/Neuron-Olig2-ChIP-Seq(GSE30882)/Homer | 1e-2 | -5.893e+00 | 0.0296 | 53.0 | 27.89% | 19019.8 | 19.35% | motif file (matrix) | svg |
| 45 | 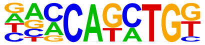 | BHLHA15(bHLH)/NIH3T3-BHLHB8.HA-ChIP-Seq(GSE119782)/Homer | 1e-2 | -5.890e+00 | 0.0296 | 35.0 | 18.42% | 11163.8 | 11.36% | motif file (matrix) | svg |
| 46 | | MyoD(bHLH)/Myotube-MyoD-ChIP-Seq(GSE21614)/Homer | 1e-2 | -5.827e+00 | 0.0302 | 18.0 | 9.47% | 4491.4 | 4.57% | motif file (matrix) | svg |

| | | | | | | | | | | | |
|---|---|---|---|---|---|---|---|---|---|---|---|
| 47 | 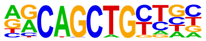 | Tcf21(bHLH)/ArterySmoothMuscle-Tcf21-ChIP-Seq(GSE61369)/Homer | 1e-2 | -5.815e+00 | 0.0302 | 23.0 | 12.11% | 6368.2 | 6.48% | motif file (matrix) | svg |
| 48 | 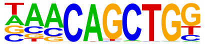 | Atoh1(bHLH)/Cerebellum-Atoh1-ChIP-Seq(GSE22111)/Homer | 1e-2 | -5.454e+00 | 0.0421 | 26.0 | 13.68% | 7748.6 | 7.88% | motif file (matrix) | svg |
| 49 | 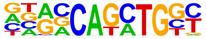 | EBF2(EBF)/BrownAdipose-EBF2-ChIP-Seq(GSE97114)/Homer | 1e-2 | -5.365e+00 | 0.0451 | 20.0 | 10.53% | 5449.7 | 5.55% | motif file (matrix) | svg |
| 50 | 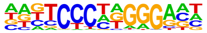 | Lhx1(Homeobox)/EmbryoCarcinoma-Lhx1-ChIP-Seq(GSE70957)/Homer | 1e-2 | -5.353e+00 | 0.0451 | 50.0 | 26.32% | 18143.5 | 18.46% | motif file (matrix) | svg |
| 51 | 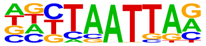 | NF1(CTF)/LNCAP-NF1-ChIP-Seq(Unpublished)/Homer | 1e-2 | -5.310e+00 | 0.0458 | 11.0 | 5.79% | 2266.6 | 2.31% | motif file (matrix) | svg |
| 52 | 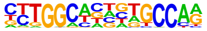 | NeuroG2(bHLH)/Fibroblast-NeuroG2-ChIP-Seq(GSE75910)/Homer | 1e-2 | -5.303e+00 | 0.0458 | 39.0 | 20.53% | 13304.0 | 13.54% | motif file (matrix) | svg |
| 53 | 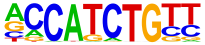 | Hoxc6(Homeobox)/EB-Hoxc6.iFlag-ChIP-Seq(GSE142377)/Homer | 1e-2 | -5.242e+00 | 0.0471 | 106.0 | 55.79% | 45470.1 | 46.27% | motif file (matrix) | svg |

| | | | | | | | | | | | |
|---|---|---|---|---|---|---|---|---|---|---|---|
| | 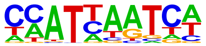 | | | | | | | | | | |
| 54 | 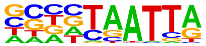 | DLX1(Homeobox)/BasalGanglia-Dlx1-ChIP-seq(GSE124936)/Homer | 1e-2 | -5.201e+00 | 0.0482 | 55.0 | 28.95% | 20569.8 | 20.93% | motif file (matrix) | svg |
| 55 | 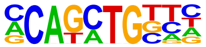 | Twist2(bHLH)/Myoblast-Twist2.Ty1-ChIP-Seq(GSE127998)/Homer | 1e-2 | -5.133e+00 | 0.0506 | 42.0 | 22.11% | 14758.1 | 15.02% | motif file (matrix) | svg |
| 56 | 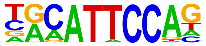 | TEAD3(TEA)/HepG2-TEAD3-ChIP-Seq(Encode)/Homer | 1e-2 | -5.113e+00 | 0.0507 | 39.0 | 20.53% | 13456.2 | 13.69% | motif file (matrix) | svg |
| 57 | 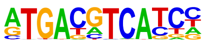 | c-Jun-CRE(bZIP)/K562-cJun-ChIP-Seq(GSE31477)/Homer | 1e-2 | -5.043e+00 | 0.0534 | 13.0 | 6.84% | 3031.5 | 3.08% | motif file (matrix) | svg |
| 58 | 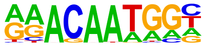 | Sox15(HMG)/CPA-Sox15-ChIP-Seq(GSE62909)/Homer | 1e-2 | -4.879e+00 | 0.0619 | 39.0 | 20.53% | 13651.0 | 13.89% | motif file (matrix) | svg |
| 59 | 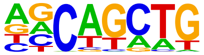 | SCL(bHLH)/HPC7-Scl-ChIP-Seq(GSE13511)/Homer | 1e-2 | -4.825e+00 | 0.0642 | 92.0 | 48.42% | 38867.9 | 39.55% | motif file (matrix) | svg |
| 60 | | EBF(EBF)/proBcell-EBF-ChIP-Seq(GSE21978)/Homer | 1e-2 | -4.818e+00 | 0.0642 | 6.0 | 3.16% | 889.4 | 0.90% | motif file (matrix) | svg |

| | | | | | | | | | | | |
|---|---|---|---|---|---|---|---|---|---|---|---|
| | 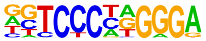 | | | | | | | | | | |
| 61 | 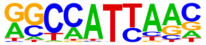 | Nanog(Homeobox)/mES-Nanog-ChIP-Seq(GSE11724)/Homer | 1e-2 | -4.762e+00 | 0.0662 | 115.0 | 60.53% | 50761.3 | 51.65% | motif file (matrix) | svg |
| 62 | 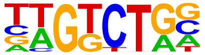 | Smad3(MAD)/NPC-Smad3-ChIP-Seq(GSE36673)/Homer | 1e-2 | -4.735e+00 | 0.0669 | 64.0 | 33.68% | 25271.1 | 25.71% | motif file (matrix) | svg |
| 63 | 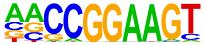 | ETS(ETS)/Promoter/Homer | 1e-2 | -4.704e+00 | 0.0679 | 8.0 | 4.21% | 1493.8 | 1.52% | motif file (matrix) | svg |
| 64 | 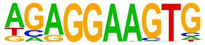 | PU.1(ETS)/ThioMac-PU.1-ChIP-Seq(GSE21512)/Homer | 1e-2 | -4.627e+00 | 0.0722 | 15.0 | 7.89% | 3926.0 | 3.99% | motif file (matrix) | svg |

### Motifs found for: solid tumor

### Homer Known Motif Enrichment Results (homerpos8)

Homer *de novo* Motif Results
Gene Ontology Enrichment Results
Known Motif Enrichment Results (txt file)
Total Target Sequences = 209, Total Background Sequences = 96676

| Rank | Motif | Name | P-value | log P-pvalue | q-value (Benjamini) | # Target Sequences with Motif | % of Targets Sequences with Motif | # Background Sequences with Motif | % of Background Sequences with Motif | Motif File | SVG |
|---|---|---|---|---|---|---|---|---|---|---|---|
| 1 | 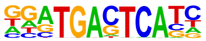 | Fra2(bZIP)/Striatum-Fra2-ChIP-Seq(GSE43429)/Homer | 1e-8 | -2.014e+01 | 0.0000 | 30.0 | 14.35% | 3876.7 | 4.01% | motif file (matrix) | svg |
| 2 | 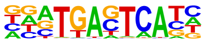 | Fra1(bZIP)/BT549-Fra1-ChIP-Seq(GSE46166)/Homer | 1e-8 | -1.912e+01 | 0.0000 | 33.0 | 15.79% | 4814.9 | 4.98% | motif file (matrix) | svg |
| 3 | 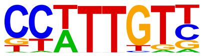 | Sox3(HMG)/NPC-Sox3-ChIP-Seq(GSE33059)/Homer | 1e-8 | -1.872e+01 | 0.0000 | 79.0 | 37.80% | 19857.9 | 20.52% | motif file (matrix) | svg |
| 4 | | Atf3(bZIP)/GBM-ATF3-ChIP-Seq(GSE33912)/Homer | 1e-7 | -1.760e+01 | 0.0000 | 36.0 | 17.22% | 5953.6 | 6.15% | motif file (matrix) | svg |

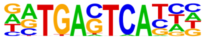

| | | | | | | | | | | | |
|---|---|---|---|---|---|---|---|---|---|---|---|
| 5 | 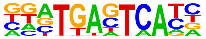 | Fos(bZIP)/TSC-Fos-ChIP-Seq(GSE110950)/ Homer | 1e-7 | -1.711e+01 | 0.0000 | 33.0 | 15.79% | 5239.0 | 5.41% | motif file (matrix) | svg |
| 6 | 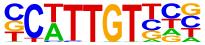 | Sox10(HMG)/SciaticNerve-Sox3-ChIP-Seq(GSE35132)/Homer | 1e-7 | -1.694e+01 | 0.0000 | 72.0 | 34.45% | 18025.9 | 18.63% | motif file (matrix) | svg |
| 7 | 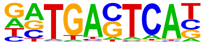 | JunB(bZIP)/DendriticCells-Junb-ChIP-Seq(GSE36099)/Homer | 1e-7 | -1.650e+01 | 0.0000 | 31.0 | 14.83% | 4834.2 | 5.00% | motif file (matrix) | svg |
| 8 | 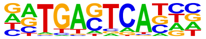 | Fosl2(bZIP)/3T3L1-Fosl2-ChIP-Seq(GSE56872)/Homer | 1e-7 | -1.649e+01 | 0.0000 | 21.0 | 10.05% | 2379.7 | 2.46% | motif file (matrix) | svg |
| 9 | 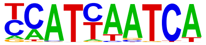 | Pdx1(Homeobox)/Islet-Pdx1-ChIP-Seq(SRA008281)/Homer | 1e-6 | -1.482e+01 | 0.0000 | 53.0 | 25.36% | 12139.8 | 12.55% | motif file (matrix) | svg |
| 10 | 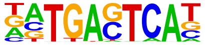 | BATF(bZIP)/Th17-BATF-ChIP-Seq(GSE39756)/ Homer | 1e-6 | -1.433e+01 | 0.0000 | 33.0 | 15.79% | 5919.9 | 6.12% | motif file (matrix) | svg |
| 11 | | Oct6(POU,Homeobox)/NPC-Pou3f1-ChIP-Seq(GSE35496)/Homer | 1e-6 | -1.386e+01 | 0.0000 | 31.0 | 14.83% | 5465.2 | 5.65% | motif file (matrix) | svg |

| | | | | | | | | | | | |
|---|---|---|---|---|---|---|---|---|---|---|---|
| | 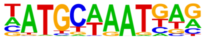 | | | | | | | | | | |
| 12 | 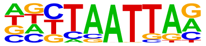 | Lhx1(Homeobox)/EmbryoCarcinoma-Lhx1-ChIP-Seq(GSE70957)/Homer | 1e-5 | -1.375e+01 | 0.0000 | 63.0 | 30.14% | 16144.6 | 16.69% | motif file (matrix) | svg |
| 13 | 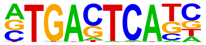 | AP-1(bZIP)/ThioMac-PU.1-ChIP-Seq(GSE21512)/Homer | 1e-5 | -1.348e+01 | 0.0001 | 35.0 | 16.75% | 6758.2 | 6.99% | motif file (matrix) | svg |
| 14 | 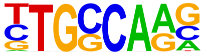 | NF1-halfsite(CTF)/LNCaP-NF1-ChIP-Seq(Unpublished)/Homer | 1e-5 | -1.343e+01 | 0.0001 | 64.0 | 30.62% | 16667.2 | 17.23% | motif file (matrix) | svg |
| 15 | 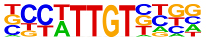 | Sox21(HMG)/ESC-SOX21-ChIP-Seq(GSE110505)/Homer | 1e-5 | -1.332e+01 | 0.0001 | 71.0 | 33.97% | 19357.1 | 20.01% | motif file (matrix) | svg |
| 16 | 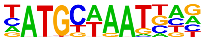 | Brn1(POU,Homeobox)/NPC-Brn1-ChIP-Seq(GSE35496)/Homer | 1e-5 | -1.287e+01 | 0.0001 | 25.0 | 11.96% | 4030.4 | 4.17% | motif file (matrix) | svg |
| 17 | 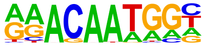 | Sox15(HMG)/CPA-Sox15-ChIP-Seq(GSE62909)/Homer | 1e-5 | -1.244e+01 | 0.0001 | 51.0 | 24.40% | 12419.2 | 12.84% | motif file (matrix) | svg |
| 18 | | Myf5(bHLH)/GM-Myf5-ChIP-Seq(GSE24852)/Homer | 1e-5 | -1.179e+01 | 0.0002 | 30.0 | 14.35% | 5748.7 | 5.94% | motif file (matrix) | svg |

| | | | | | | | | | | | |
|---|---|---|---|---|---|---|---|---|---|---|---|
| | 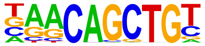 | | | | | | | | | | |
| 19 | 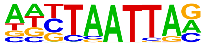 | Lhx3(Homeobox)/Neuron-Lhx3-ChIP-Seq(GSE31456)/Homer | 1e-5 | -1.179e+01 | 0.0002 | 78.0 | 37.32% | 22945.4 | 23.72% | motif file (matrix) | svg |
| 20 | 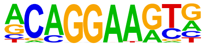 | ERG(ETS)/VCaP-ERG-ChIP-Seq(GSE14097)/Homer | 1e-4 | -1.120e+01 | 0.0003 | 53.0 | 25.36% | 13701.6 | 14.16% | motif file (matrix) | svg |
| 21 | 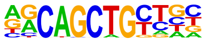 | MyoD(bHLH)/Myotube-MyoD-ChIP-Seq(GSE21614)/Homer | 1e-4 | -1.115e+01 | 0.0003 | 30.0 | 14.35% | 5946.5 | 6.15% | motif file (matrix) | svg |
| 22 | 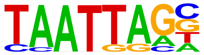 | Lhx2(Homeobox)/HFSC-Lhx2-ChIP-Seq(GSE48068)/Homer | 1e-4 | -1.067e+01 | 0.0005 | 54.0 | 25.84% | 14331.9 | 14.81% | motif file (matrix) | svg |
| 23 | 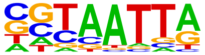 | DLX5(Homeobox)/BasalGanglia-Dlx5-ChIP-seq(GSE124936)/Homer | 1e-4 | -1.055e+01 | 0.0005 | 44.0 | 21.05% | 10793.9 | 11.16% | motif file (matrix) | svg |
| 24 | 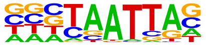 | DLX2(Homeobox)/BasalGanglia-Dlx2-ChIP-seq(GSE124936)/Homer | 1e-4 | -1.054e+01 | 0.0005 | 71.0 | 33.97% | 20917.6 | 21.62% | motif file (matrix) | svg |
| 25 | | SOX1(HMG)/NPC-SOX1-ChIP-Seq(GSE138215)/Homer | 1e-4 | -1.045e+01 | 0.0005 | 80.0 | 38.28% | 24594.8 | 25.42% | motif file (matrix) | svg |

| | | | | | | | | | | | |
|---|---|---|---|---|---|---|---|---|---|---|---|
| | 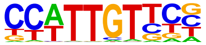 | | | | | | | | | | |
| 26 | 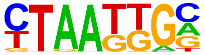 | Isl1(Homeobox)/Neuron-Isl1-ChIP-Seq(GSE31456)/Homer | 1e-4 | -1.044e+01 | 0.0005 | 77.0 | 36.84% | 23378.8 | 24.16% | motif file (matrix) | svg |
| 27 | 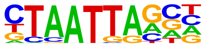 | Gsx2(Homeobox)/LGE-Gsx2.Flag-ChIP-Seq(GSE162589)/Homer | 1e-4 | -9.305e+00 | 0.0016 | 62.0 | 29.67% | 18137.3 | 18.75% | motif file (matrix) | svg |
| 28 | 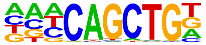 | Ap4(bHLH)/AML-Tfap4-ChIP-Seq(GSE45738)/Homer | 1e-4 | -9.243e+00 | 0.0016 | 41.0 | 19.62% | 10340.4 | 10.69% | motif file (matrix) | svg |
| 29 | 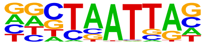 | En1(Homeobox)/SUM149-EN1-ChIP-Seq(GSE120957)/Homer | 1e-3 | -9.053e+00 | 0.0019 | 77.0 | 36.84% | 24316.7 | 25.13% | motif file (matrix) | svg |
| 30 | 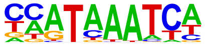 | Hoxc10(Homeobox)/EB-Hoxc10.iFlag-ChIP-Seq(GSE142377)/Homer | 1e-3 | -8.971e+00 | 0.0020 | 60.0 | 28.71% | 17565.2 | 18.15% | motif file (matrix) | svg |
| 31 | 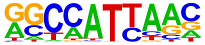 | Nanog(Homeobox)/mES-Nanog-ChIP-Seq(GSE11724)/Homer | 1e-3 | -8.955e+00 | 0.0020 | 131.0 | 62.68% | 48243.1 | 49.86% | motif file (matrix) | svg |
| 32 | | Hoxd9(Homeobox)/EB-Hoxd9.HA-ChIP-Seq(GSE142377)/Homer | 1e-3 | -8.782e+00 | 0.0023 | 24.0 | 11.48% | 4860.4 | 5.02% | motif file (matrix) | svg |

| | | | | | | | | | | | |
|---|---|---|---|---|---|---|---|---|---|---|---|
| | 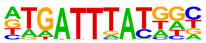 | | | | | | | | | | |
| 33 | | Sox6(HMG)/Myotubes-Sox6-ChIP-Seq(GSE32627)/Homer | 1e-3 | -8.735e+00 | 0.0023 | 62.0 | 29.67% | 18497.4 | 19.12% | motif file (matrix) | svg |
| | 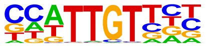 | | | | | | | | | | |
| 34 | | Sox9(HMG)/Limb-SOX9-ChIP-Seq(GSE73225)/Homer | 1e-3 | -8.688e+00 | 0.0023 | 34.0 | 16.27% | 8158.3 | 8.43% | motif file (matrix) | svg |
| | 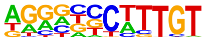 | | | | | | | | | | |
| 35 | | Hoxd11(Homeobox)/ChickenMSG-Hoxd11.Flag-ChIP-Seq(GSE86088)/Homer | 1e-3 | -8.434e+00 | 0.0029 | 93.0 | 44.50% | 31533.1 | 32.59% | motif file (matrix) | svg |
| | 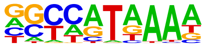 | | | | | | | | | | |
| 36 | | Smad3(MAD)/NPC-Smad3-ChIP-Seq(GSE36673)/Homer | 1e-3 | -8.229e+00 | 0.0035 | 82.0 | 39.23% | 27009.4 | 27.92% | motif file (matrix) | svg |
| | 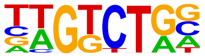 | | | | | | | | | | |
| 37 | | Dlx3(Homeobox)/Kerainocytes-Dlx3-ChIP-Seq(GSE89884)/Homer | 1e-3 | -8.151e+00 | 0.0037 | 37.0 | 17.70% | 9450.3 | 9.77% | motif file (matrix) | svg |
| | 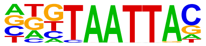 | | | | | | | | | | |
| 38 | | Jun-AP1(bZIP)/K562-cJun-ChIP-Seq(GSE31477)/Homer | 1e-3 | -8.139e+00 | 0.0037 | 12.0 | 5.74% | 1652.7 | 1.71% | motif file (matrix) | svg |
| | 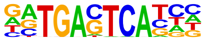 | | | | | | | | | | |
| 39 | | Sox2(HMG)/mES-Sox2-ChIP-Seq(GSE11431)/Homer | 1e-3 | -7.997e+00 | 0.0041 | 37.0 | 17.70% | 9526.2 | 9.85% | motif file (matrix) | svg |

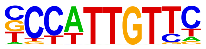

| | | | | | | | | | | | |
|---|---|---|---|---|---|---|---|---|---|---|---|
| 40 | 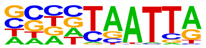 | DLX1(Homeobox)/BasalGanglia-Dlx1-ChIP-seq(GSE124936)/Homer | 1e-3 | -7.820e+00 | 0.0047 | 61.0 | 29.19% | 18714.9 | 19.34% | motif file (matrix) | svg |
| 41 | 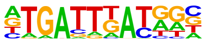 | PBX2(Homeobox)/K562-PBX2-ChIP-Seq(Encode)/Homer | 1e-3 | -7.531e+00 | 0.0062 | 38.0 | 18.18% | 10124.3 | 10.46% | motif file (matrix) | svg |
| 42 | 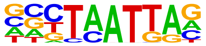 | Emx2(Homeobox)/Cortex-Emx2-ChIP-Seq(GSE183130)/Homer | 1e-3 | -7.529e+00 | 0.0062 | 51.0 | 24.40% | 14992.4 | 15.50% | motif file (matrix) | svg |
| 43 | 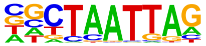 | LHX9(Homeobox)/Hct116-LHX9.V5-ChIP-Seq(GSE116822)/Homer | 1e-3 | -7.383e+00 | 0.0068 | 60.0 | 28.71% | 18624.0 | 19.25% | motif file (matrix) | svg |
| 44 | 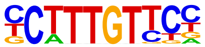 | Sox4(HMG)/proB-Sox4-ChIP-Seq(GSE50066)/Homer | 1e-3 | -7.352e+00 | 0.0069 | 34.0 | 16.27% | 8783.0 | 9.08% | motif file (matrix) | svg |
| 45 | 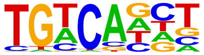 | Tgif2(Homeobox)/mES-Tgif2-ChIP-Seq(GSE55404)/Homer | 1e-3 | -7.340e+00 | 0.0069 | 101.0 | 48.33% | 35976.2 | 37.18% | motif file (matrix) | svg |
| 46 | | ETV4(ETS)/HepG2-ETV4-ChIP-Seq(ENCODE)/Homer | 1e-3 | -7.124e+00 | 0.0083 | 30.0 | 14.35% | 7488.1 | 7.74% | motif file (matrix) | svg |

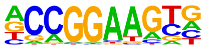

| | | | | | | | | | | | |
|---|---|---|---|---|---|---|---|---|---|---|---|
| 47 | 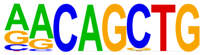 | MyoG(bHLH)/C2C12-MyoG-ChIP-Seq(GSE36024)/Homer | 1e-2 | -6.825e+00 | 0.0109 | 32.0 | 15.31% | 8338.9 | 8.62% | motif file (matrix) | svg |
| 48 | 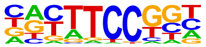 | Fli1(ETS)/CD8-FLI-ChIP-Seq(GSE20898)/Homer | 1e-2 | -6.701e+00 | 0.0121 | 31.0 | 14.83% | 8046.3 | 8.32% | motif file (matrix) | svg |
| 49 | 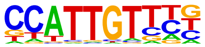 | Sox17(HMG)/Endoderm-Sox17-ChIP-Seq(GSE61475)/Homer | 1e-2 | -6.693e+00 | 0.0121 | 31.0 | 14.83% | 8051.0 | 8.32% | motif file (matrix) | svg |
| 50 | 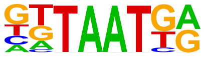 | Nkx6.1(Homeobox)/Islet-Nkx6.1-ChIP-Seq(GSE40975)/Homer | 1e-2 | -6.620e+00 | 0.0126 | 98.0 | 46.89% | 35322.6 | 36.51% | motif file (matrix) | svg |
| 51 | 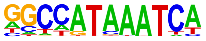 | Hoxc9(Homeobox)/Ainv15-Hoxc9-ChIP-Seq(GSE21812)/Homer | 1e-2 | -6.383e+00 | 0.0156 | 26.0 | 12.44% | 6452.8 | 6.67% | motif file (matrix) | svg |
| 52 | 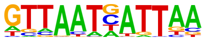 | HNF1b(Homeobox)/PDAC-HNF1B-ChIP-Seq(GSE64557)/Homer | 1e-2 | -6.374e+00 | 0.0156 | 12.0 | 5.74% | 2029.6 | 2.10% | motif file (matrix) | svg |
| 53 | | Oct4(POU,Homeobox)/mES-Oct4-ChIP-Seq(GSE11431)/Homer | 1e-2 | -6.293e+00 | 0.0165 | 26.0 | 12.44% | 6494.6 | 6.71% | motif file (matrix) | svg |

| | | | | | | | | | | | |
|---|---|---|---|---|---|---|---|---|---|---|---|
| | 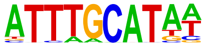 | | | | | | | | | | |
| 54 | 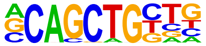 | Tcf12(bHLH)/GM12878-Tcf12-ChIP-Seq(GSE32465)/Homer | 1e-2 | -6.293e+00 | 0.0165 | 29.0 | 13.88% | 7545.5 | 7.80% | motif file (matrix) | svg |
| 55 | 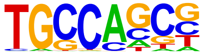 | HIC1(Zf)/Treg-ZBTB29-ChIP-Seq(GSE99889)/Homer | 1e-2 | -6.051e+00 | 0.0202 | 58.0 | 27.75% | 18831.3 | 19.46% | motif file (matrix) | svg |
| 56 | 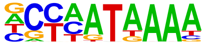 | Hoxd13(Homeobox)/ChickenMSG-Hoxd13.Flag-ChIP-Seq(GSE86088)/Homer | 1e-2 | -5.961e+00 | 0.0217 | 66.0 | 31.58% | 22194.8 | 22.94% | motif file (matrix) | svg |
| 57 | 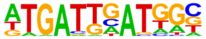 | HOXA3(Homeobox)/mEmbryo-Hoxa3-ChIP-Seq(E-MTAB-8607)/Homer | 1e-2 | -5.878e+00 | 0.0232 | 10.0 | 4.78% | 1604.0 | 1.66% | motif file (matrix) | svg |
| 58 | 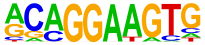 | ETS1(ETS)/Jurkat-ETS1-ChIP-Seq(GSE17954)/Homer | 1e-2 | -5.873e+00 | 0.0232 | 30.0 | 14.35% | 8127.5 | 8.40% | motif file (matrix) | svg |
| 59 | 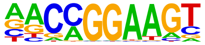 | ETV1(ETS)/GIST48-ETV1-ChIP-Seq(GSE22441)/Homer | 1e-2 | -5.802e+00 | 0.0242 | 37.0 | 17.70% | 10761.3 | 11.12% | motif file (matrix) | svg |
| 60 | | Rfx2(HTH)/LoVo-RFX2-ChIP-Seq(GSE49402)/Homer | 1e-2 | -5.524e+00 | 0.0314 | 5.0 | 2.39% | 476.8 | 0.49% | motif file (matrix) | svg |

| | | | | | | | | | | | |
|---|---|---|---|---|---|---|---|---|---|---|---|
| | 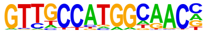 | | | | | | | | | | |
| 61 | 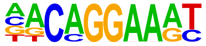 | EWS:FLI1-fusion(ETS)/SK_N_MC-EWS:FLI1-ChIP-Seq(SRA014231)/Homer | 1e-2 | -5.502e+00 | 0.0316 | 19.0 | 9.09% | 4470.0 | 4.62% | motif file (matrix) | svg |
| 62 | 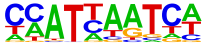 | Hoxc6(Homeobox)/EB-Hoxc6.iFlag-ChIP-Seq(GSE142377)/Homer | 1e-2 | -5.422e+00 | 0.0336 | 104.0 | 49.76% | 39244.7 | 40.56% | motif file (matrix) | svg |
| 63 | 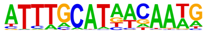 | OCT4-SOX2-TCF-NANOG(POU,Homeobox,HMG)/mES-Oct4-ChIP-Seq(GSE11431)/Homer | 1e-2 | -5.414e+00 | 0.0336 | 13.0 | 6.22% | 2587.2 | 2.67% | motif file (matrix) | svg |
| 64 | 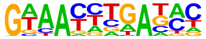 | Six2(Homeobox)/NephronProgenitor-Six2-ChIP-Seq(GSE39837)/Homer | 1e-2 | -5.411e+00 | 0.0336 | 35.0 | 16.75% | 10260.0 | 10.60% | motif file (matrix) | svg |
| 65 | 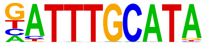 | Oct11(POU,Homeobox)/NCIH1048-POU2F3-ChIP-seq(GSE115123)/Homer | 1e-2 | -5.401e+00 | 0.0336 | 18.0 | 8.61% | 4180.8 | 4.32% | motif file (matrix) | svg |
| 66 | 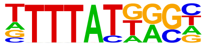 | Hoxc13(Homeobox)/EB-Hoxc13.HA-ChIP-Seq(GSE142377)/Homer | 1e-2 | -5.280e+00 | 0.0364 | 56.0 | 26.79% | 18665.4 | 19.29% | motif file (matrix) | svg |
| 67 | | Etv2(ETS)/ES-ER71-ChIP-Seq(GSE59402)/Homer | 1e-2 | -5.277e+00 | 0.0364 | 28.0 | 13.40% | 7737.9 | 8.00% | motif file (matrix) | svg |

| | | | | | | | | | | | |
|---|---|---|---|---|---|---|---|---|---|---|---|
| 68 | 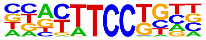 | Tlx?(NR)/NPC-H3K4me1-ChIP-Seq(GSE16256)/Homer | 1e-2 | -5.062e+00 | 0.0440 | 15.0 | 7.18% | 3335.1 | 3.45% | motif file (matrix) | svg |
| 69 | 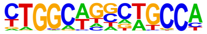 | IRF:BATF(IRF:bZIP)/pDC-Irf8-ChIP-Seq(GSE66899)/Homer | 1e-2 | -5.013e+00 | 0.0455 | 8.0 | 3.83% | 1265.0 | 1.31% | motif file (matrix) | svg |
| 70 | 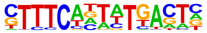 | NF1:FOXA1(CTF,Forkhead)/LNCAP-FOXA1-ChIP-Seq(GSE27824)/Homer | 1e-2 | -4.973e+00 | 0.0467 | 5.0 | 2.39% | 544.2 | 0.56% | motif file (matrix) | svg |
| 71 | 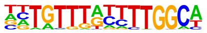 | Tcf7(HMG)/GM12878-TCF7-ChIP-Seq(Encode)/Homer | 1e-2 | -4.960e+00 | 0.0467 | 16.0 | 7.66% | 3701.1 | 3.83% | motif file (matrix) | svg |
| 72 | 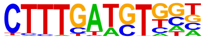 | X-box(HTH)/NPC-H3K4me1-ChIP-Seq(GSE16256)/Homer | 1e-2 | -4.818e+00 | 0.0530 | 6.0 | 2.87% | 795.8 | 0.82% | motif file (matrix) | svg |
| 73 | 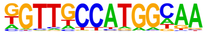 | Smad2(MAD)/ES-SMAD2-ChIP-Seq(GSE29422)/Homer | 1e-2 | -4.772e+00 | 0.0547 | 39.0 | 18.66% | 12257.9 | 12.67% | motif file (matrix) | svg |
| 74 | 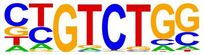 | GABPA(ETS)/Jurkat-GABPa-ChIP-Seq(GSE17954)/Homer | 1e-2 | -4.673e+00 | 0.0596 | 24.0 | 11.48% | 6628.6 | 6.85% | motif file (matrix) | svg |

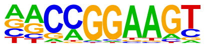

| | | | | | | | | | | | |
|---|---|---|---|---|---|---|---|---|---|---|---|
| 75 | 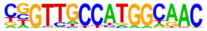 | RFX(HTH)/K562-RFX3-ChIP-Seq(SRA012198)/Homer | 1e-2 | -4.641e+00 | 0.0607 | 4.0 | 1.91% | 379.7 | 0.39% | motif file (matrix) | svg |

#### Motifs found for: subarachnoid invasion

### Homer Known Motif Enrichment Results (homerpos9)

[Homer *de novo* Motif Results]
[Gene Ontology Enrichment Results]
[Known Motif Enrichment Results (txt file)]
Total Target Sequences = 914, Total Background Sequences = 95597

| Rank | Motif | Name | P-value | log P-pvalue | q-value (Benjamini) | # Target Sequences with Motif | % of Targets Sequences with Motif | # Background Sequences with Motif | % of Background Sequences with Motif | Motif File | SVG |
|---|---|---|---|---|---|---|---|---|---|---|---|
| 1 | 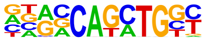 | Atoh1(bHLH)/Cerebellum-Atoh1-ChIP-Seq(GSE22111)/Homer | 1e-45 | -1.042e+02 | 0.0000 | 237.0 | 25.93% | 9158.2 | 9.58% | motif file (matrix) | svg |
| 2 | 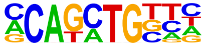 | Twist2(bHLH)/Myoblast-Twist2.Ty1-ChIP-Seq(GSE127998)/Homer | 1e-42 | -9.735e+01 | 0.0000 | 325.0 | 35.56% | 15990.7 | 16.72% | motif file (matrix) | svg |
| 3 | 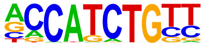 | NeuroG2(bHLH)/Fibroblast-NeuroG2-ChIP-Seq(GSE75910)/Homer | 1e-38 | -8.964e+01 | 0.0000 | 294.0 | 32.17% | 14184.3 | 14.83% | motif file (matrix) | svg |
| 4 | | BHLHA15(bHLH)/NIH3T3-BHLHB8.HA-ChIP-Seq(GSE119782)/Homer | 1e-38 | -8.912e+01 | 0.0000 | 269.0 | 29.43% | 12346.2 | 12.91% | motif file (matrix) | svg |

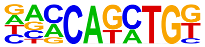

| | | | | | | | | | | | |
|---|---|---|---|---|---|---|---|---|---|---|---|
| 5 | 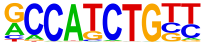 | NeuroD1(bHLH)/Islet-NeuroD1-ChIP-Seq(GSE30298)/Homer | 1e-38 | -8.857e+01 | 0.0000 | 187.0 | 20.46% | 6767.2 | 7.08% | motif file (matrix) | svg |
| 6 | 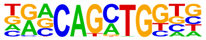 | Atoh7(bHLH)/Retina-Atoh7-CutnRun(GSE156756)/Homer | 1e-37 | -8.579e+01 | 0.0000 | 177.0 | 19.37% | 6286.7 | 6.57% | motif file (matrix) | svg |
| 7 | 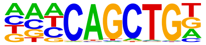 | Ap4(bHLH)/AML-Tfap4-ChIP-Seq(GSE45738)/Homer | 1e-34 | -7.941e+01 | 0.0000 | 232.0 | 25.38% | 10340.1 | 10.81% | motif file (matrix) | svg |
| 8 | 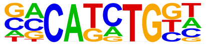 | TCF4(bHLH)/SHSY5Y-TCF4-ChIP-Seq(GSE96915)/Homer | 1e-33 | -7.655e+01 | 0.0000 | 277.0 | 30.31% | 13910.7 | 14.54% | motif file (matrix) | svg |
| 9 | 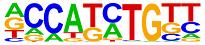 | Olig2(bHLH)/Neuron-Olig2-ChIP-Seq(GSE30882)/Homer | 1e-30 | -7.119e+01 | 0.0000 | 332.0 | 36.32% | 18840.2 | 19.70% | motif file (matrix) | svg |
| 10 | 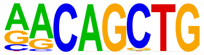 | MyoG(bHLH)/C2C12-MyoG-ChIP-Seq(GSE36024)/Homer | 1e-30 | -6.941e+01 | 0.0000 | 196.0 | 21.44% | 8484.1 | 8.87% | motif file (matrix) | svg |
| 11 | | Sox10(HMG)/SciaticNerve-Sox3-ChIP-Seq(GSE35132)/Homer | 1e-30 | -6.934e+01 | 0.0000 | 319.0 | 34.90% | 17935.4 | 18.75% | motif file (matrix) | svg |

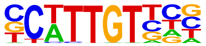

| | | | | | | | | | | | |
|---|---|---|---|---|---|---|---|---|---|---|---|
| 12 | 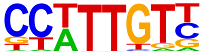 | Sox3(HMG)/NPC-Sox3-ChIP-Seq(GSE33059)/ Homer | 1e-29 | -6.841e+01 | 0.0000 | 338.0 | 36.98% | 19628.0 | 20.52% | motif file (matrix) | svg |
| 13 | 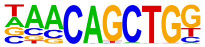 | Tcf21(bHLH)/ArterySmoothMuscle-Tcf21-ChIP-Seq(GSE61369)/Homer | 1e-29 | -6.791e+01 | 0.0000 | 186.0 | 20.35% | 7894.1 | 8.25% | motif file (matrix) | svg |
| 14 | 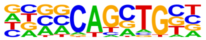 | Ascl1(bHLH)/NeuralTubes-Ascl1-ChIP-Seq(GSE55840)/Homer | 1e-26 | -6.194e+01 | 0.0000 | 248.0 | 27.13% | 12906.0 | 13.49% | motif file (matrix) | svg |
| 15 | 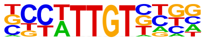 | Sox21(HMG)/ESC-SOX21-ChIP-Seq(GSE110505)/Homer | 1e-26 | -6.135e+01 | 0.0000 | 329.0 | 36.00% | 19609.4 | 20.50% | motif file (matrix) | svg |
| 16 | 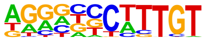 | Sox9(HMG)/Limb-SOX9-ChIP-Seq(GSE73225)/ Homer | 1e-25 | -5.876e+01 | 0.0000 | 181.0 | 19.80% | 8174.9 | 8.55% | motif file (matrix) | svg |
| 17 | 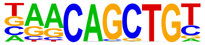 | Myf5(bHLH)/GM-Myf5-ChIP-Seq(GSE24852)/ Homer | 1e-25 | -5.784e+01 | 0.0000 | 145.0 | 15.86% | 5787.9 | 6.05% | motif file (matrix) | svg |
| 18 | | MyoD(bHLH)/Myotube-MyoD-ChIP-Seq(GSE21614)/Homer | 1e-24 | -5.597e+01 | 0.0000 | 147.0 | 16.08% | 6034.0 | 6.31% | motif file (matrix) | svg |

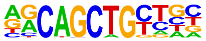

| | | | | | | | | | | | |
|---|---|---|---|---|---|---|---|---|---|---|---|
| 19 | 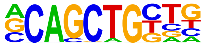 | Tcf12(bHLH)/GM12878-Tcf12-ChIP-Seq(GSE32465)/Homer | 1e-24 | -5.555e+01 | 0.0000 | 171.0 | 18.71% | 7701.8 | 8.05% | motif file (matrix) | svg |
| 20 | 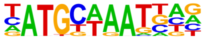 | Brn1(POU,Homeobox)/NPC-Brn1-ChIP-Seq(GSE35496)/Homer | 1e-23 | -5.432e+01 | 0.0000 | 114.0 | 12.47% | 4036.8 | 4.22% | motif file (matrix) | svg |
| 21 | 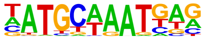 | Oct6(POU,Homeobox)/NPC-Pou3f1-ChIP-Seq(GSE35496)/Homer | 1e-22 | -5.195e+01 | 0.0000 | 133.0 | 14.55% | 5363.6 | 5.61% | motif file (matrix) | svg |
| 22 | 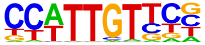 | SOX1(HMG)/NPC-SOX1-ChIP-Seq(GSE138215)/Homer | 1e-21 | -5.023e+01 | 0.0000 | 374.0 | 40.92% | 24963.0 | 26.10% | motif file (matrix) | svg |
| 23 | 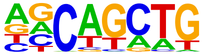 | SCL(bHLH)/HPC7-Scl-ChIP-Seq(GSE13511)/Homer | 1e-20 | -4.813e+01 | 0.0000 | 547.0 | 59.85% | 42232.9 | 44.16% | motif file (matrix) | svg |
| 24 | 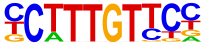 | Sox4(HMG)/proB-Sox4-ChIP-Seq(GSE50066)/Homer | 1e-20 | -4.658e+01 | 0.0000 | 177.0 | 19.37% | 8843.8 | 9.25% | motif file (matrix) | svg |
| 25 | | NF1(CTF)/LNCAP-NF1-ChIP-Seq(Unpublished)/Homer | 1e-20 | -4.627e+01 | 0.0000 | 87.0 | 9.52% | 2842.9 | 2.97% | motif file (matrix) | svg |

| | | | | | | | | | | | |
|---|---|---|---|---|---|---|---|---|---|---|---|
| | 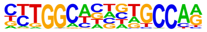 | | | | | | | | | | |
| 26 | 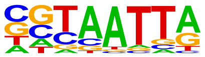 | DLX5(Homeobox)/BasalGanglia-Dlx5-ChIP-seq(GSE124936)/Homer | 1e-18 | -4.336e+01 | 0.0000 | 194.0 | 21.23% | 10433.3 | 10.91% | motif file (matrix) | svg |
| 27 | 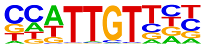 | Sox6(HMG)/Myotubes-Sox6-ChIP-Seq(GSE32627)/Homer | 1e-18 | -4.294e+01 | 0.0000 | 296.0 | 32.39% | 18945.5 | 19.81% | motif file (matrix) | svg |
| 28 | 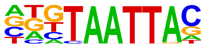 | Dlx3(Homeobox)/Kerainocytes-Dlx3-ChIP-Seq(GSE89884)/Homer | 1e-18 | -4.256e+01 | 0.0000 | 175.0 | 19.15% | 9050.7 | 9.46% | motif file (matrix) | svg |
| 29 | 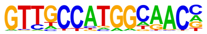 | Rfx2(HTH)/LoVo-RFX2-ChIP-Seq(GSE49402)/Homer | 1e-17 | -4.133e+01 | 0.0000 | 33.0 | 3.61% | 453.4 | 0.47% | motif file (matrix) | svg |
| 30 | 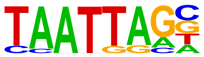 | Lhx2(Homeobox)/HFSC-Lhx2-ChIP-Seq(GSE48068)/Homer | 1e-17 | -4.116e+01 | 0.0000 | 233.0 | 25.49% | 13800.2 | 14.43% | motif file (matrix) | svg |
| 31 | 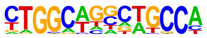 | Tlx?(NR)/NPC-H3K4me1-ChIP-Seq(GSE16256)/Homer | 1e-17 | -4.016e+01 | 0.0000 | 87.0 | 9.52% | 3143.7 | 3.29% | motif file (matrix) | svg |
| 32 | | Lhx1(Homeobox)/EmbryoCarcinoma-Lhx1-ChIP-Seq(GSE70957)/Homer | 1e-17 | -3.981e+01 | 0.0000 | 251.0 | 27.46% | 15469.7 | 16.17% | motif file (matrix) | svg |

| | | | | | | | | | | | |
|---|---|---|---|---|---|---|---|---|---|---|---|
| 33 | 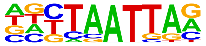 | En1(Homeobox)/SUM149-EN1-ChIP-Seq(GSE120957)/Homer | 1e-17 | -3.951e+01 | 0.0000 | 338.0 | 36.98% | 23187.2 | 24.24% | motif file (matrix) | svg |
| 34 | 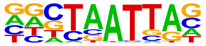 | DLX1(Homeobox)/BasalGanglia-Dlx1-ChIP-seq(GSE124936)/Homer | 1e-17 | -3.929e+01 | 0.0000 | 278.0 | 30.42% | 17856.5 | 18.67% | motif file (matrix) | svg |
| 35 | 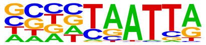 | Lhx3(Homeobox)/Neuron-Lhx3-ChIP-Seq(GSE31456)/Homer | 1e-16 | -3.841e+01 | 0.0000 | 322.0 | 35.23% | 21894.3 | 22.89% | motif file (matrix) | svg |
| 36 | 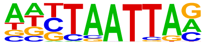 | RFX(HTH)/K562-RFX3-ChIP-Seq(SRA012198)/Homer | 1e-16 | -3.724e+01 | 0.0000 | 29.0 | 3.17% | 387.6 | 0.41% | motif file (matrix) | svg |
| 37 | 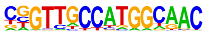 | Nkx6.1(Homeobox)/Islet-Nkx6.1-ChIP-Seq(GSE40975)/Homer | 1e-16 | -3.710e+01 | 0.0000 | 444.0 | 48.58% | 33672.4 | 35.21% | motif file (matrix) | svg |
| 38 | 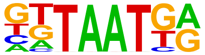 | NF1-halfsite(CTF)/LNCaP-NF1-ChIP-Seq(Unpublished)/Homer | 1e-16 | -3.696e+01 | 0.0000 | 256.0 | 28.01% | 16262.8 | 17.00% | motif file (matrix) | svg |
| 39 | 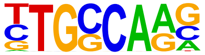 | Gsx2(Homeobox)/LGE-Gsx2.Flag-ChIP-Seq(GSE162589)/Homer | 1e-15 | -3.611e+01 | 0.0000 | 267.0 | 29.21% | 17330.6 | 18.12% | motif file (matrix) | svg |

| | | | | | | | | | | | |
|---|---|---|---|---|---|---|---|---|---|---|---|
| | 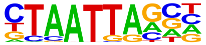 | | | | | | | | | | |
| 40 | 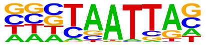 | DLX2(Homeobox)/BasalGanglia-Dlx2-ChIP-seq(GSE124936)/Homer | 1e-15 | -3.573e+01 | 0.0000 | 298.0 | 32.60% | 20128.5 | 21.05% | motif file (matrix) | svg |
| 41 | 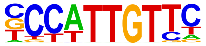 | Sox2(HMG)/mES-Sox2-ChIP-Seq(GSE11431)/Homer | 1e-15 | -3.498e+01 | 0.0000 | 171.0 | 18.71% | 9486.5 | 9.92% | motif file (matrix) | svg |
| 42 | 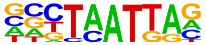 | Emx2(Homeobox)/Cortex-Emx2-ChIP-Seq(GSE183130)/Homer | 1e-15 | -3.486e+01 | 0.0000 | 230.0 | 25.16% | 14319.2 | 14.97% | motif file (matrix) | svg |
| 43 | 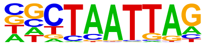 | LHX9(Homeobox)/Hct116-LHX9.V5-ChIP-Seq(GSE116822)/Homer | 1e-15 | -3.482e+01 | 0.0000 | 271.0 | 29.65% | 17862.5 | 18.68% | motif file (matrix) | svg |
| 44 | 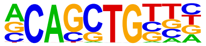 | Ptf1a(bHLH)/Panc1-Ptf1a-ChIP-Seq(GSE47459)/Homer | 1e-14 | -3.334e+01 | 0.0000 | 347.0 | 37.96% | 24998.2 | 26.14% | motif file (matrix) | svg |
| 45 | 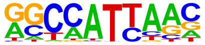 | Nanog(Homeobox)/mES-Nanog-ChIP-Seq(GSE11724)/Homer | 1e-14 | -3.277e+01 | 0.0000 | 565.0 | 61.82% | 46909.6 | 49.05% | motif file (matrix) | svg |
| 46 | | Sox17(HMG)/Endoderm-Sox17-ChIP-Seq(GSE61475)/Homer | 1e-13 | -3.157e+01 | 0.0000 | 147.0 | 16.08% | 7973.2 | 8.34% | motif file (matrix) | svg |

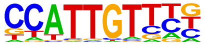

| | | | | | | | | | | | |
|---|---|---|---|---|---|---|---|---|---|---|---|
| 47 | 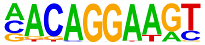 | Ets1-distal(ETS)/CD4+-PolII-ChIP-Seq(Barski_et_al.)/Homer | 1e-12 | -2.991e+01 | 0.0000 | 65.0 | 7.11% | 2363.4 | 2.47% | motif file (matrix) | svg |
| 48 | 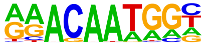 | Sox15(HMG)/CPA-Sox15-ChIP-Seq(GSE62909)/Homer | 1e-12 | -2.922e+01 | 0.0000 | 197.0 | 21.55% | 12279.1 | 12.84% | motif file (matrix) | svg |
| 49 | 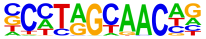 | Rfx5(HTH)/GM12878-Rfx5-ChIP-Seq(GSE31477)/Homer | 1e-12 | -2.838e+01 | 0.0000 | 68.0 | 7.44% | 2633.8 | 2.75% | motif file (matrix) | svg |
| 50 | 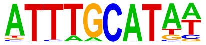 | Oct4(POU,Homeobox)/mES-Oct4-ChIP-Seq(GSE11431)/Homer | 1e-12 | -2.825e+01 | 0.0000 | 121.0 | 13.24% | 6326.7 | 6.62% | motif file (matrix) | svg |
| 51 | 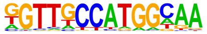 | X-box(HTH)/NPC-H3K4me1-ChIP-Seq(GSE16256)/Homer | 1e-11 | -2.757e+01 | 0.0000 | 34.0 | 3.72% | 789.1 | 0.83% | motif file (matrix) | svg |
| 52 | 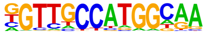 | Rfx1(HTH)/NPC-H3K4me1-ChIP-Seq(GSE16256)/Homer | 1e-11 | -2.618e+01 | 0.0000 | 46.0 | 5.03% | 1448.0 | 1.51% | motif file (matrix) | svg |
| 53 | | Lhx6/Neurons-Lhx6-ChIP-seq(GSE85704)/Homer | 1e-10 | -2.434e+01 | 0.0000 | 215.0 | 23.52% | 14531.0 | 15.19% | motif file (matrix) | svg |

| | | | | | | | | | | | |
|---|---|---|---|---|---|---|---|---|---|---|---|
| | 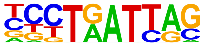 | | | | | | | | | | |
| 54 | 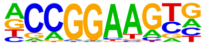 | ETV4(ETS)/HepG2-ETV4-ChIP-Seq(ENCODE)/Homer | 1e-10 | -2.406e+01 | 0.0000 | 132.0 | 14.44% | 7618.1 | 7.97% | motif file (matrix) | svg |
| 55 | 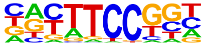 | Fli1(ETS)/CD8-FLI-ChIP-Seq(GSE20898)/Homer | 1e-10 | -2.348e+01 | 0.0000 | 139.0 | 15.21% | 8247.8 | 8.62% | motif file (matrix) | svg |
| 56 | 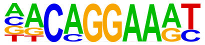 | EWS:FLI1-fusion(ETS)/SK_N_MC-EWS:FLI1-ChIP-Seq(SRA014231)/Homer | 1e-9 | -2.281e+01 | 0.0000 | 91.0 | 9.96% | 4617.8 | 4.83% | motif file (matrix) | svg |
| 57 | 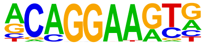 | ERG(ETS)/VCaP-ERG-ChIP-Seq(GSE14097)/Homer | 1e-9 | -2.235e+01 | 0.0000 | 204.0 | 22.32% | 13887.8 | 14.52% | motif file (matrix) | svg |
| 58 | 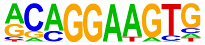 | ETS1(ETS)/Jurkat-ETS1-ChIP-Seq(GSE17954)/Homer | 1e-9 | -2.218e+01 | 0.0000 | 136.0 | 14.88% | 8167.2 | 8.54% | motif file (matrix) | svg |
| 59 | 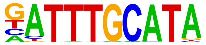 | Oct11(POU,Homeobox)/NCIH1048-POU2F3-ChIP-seq(GSE115123)/Homer | 1e-9 | -2.136e+01 | 0.0000 | 82.0 | 8.97% | 4094.5 | 4.28% | motif file (matrix) | svg |
| 60 | | Isl1(Homeobox)/Neuron-Isl1-ChIP-Seq(GSE31456)/Homer | 1e-8 | -2.063e+01 | 0.0000 | 294.0 | 32.17% | 22404.3 | 23.43% | motif file (matrix) | svg |

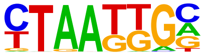

| | | | | | | | | | | | |
|---|---|---|---|---|---|---|---|---|---|---|---|
| 61 | 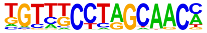 | Rfx6(HTH)/Min6b1-Rfx6.HA-ChIP-Seq(GSE62844)/Homer | 1e-8 | -2.056e+01 | 0.0000 | 134.0 | 14.66% | 8214.5 | 8.59% | motif file (matrix) | svg |
| 62 | 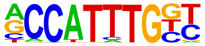 | Mesp1(bHLH)/ESC-Mesp1-ChIP-Seq(GSE165102)/Homer | 1e-8 | -2.042e+01 | 0.0000 | 106.0 | 11.60% | 5995.5 | 6.27% | motif file (matrix) | svg |
| 63 | 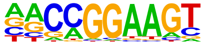 | GABPA(ETS)/Jurkat-GABPa-ChIP-Seq(GSE17954)/Homer | 1e-8 | -1.991e+01 | 0.0000 | 112.0 | 12.25% | 6525.4 | 6.82% | motif file (matrix) | svg |
| 64 | 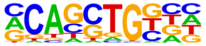 | HEB(bHLH)/mES-Heb-ChIP-Seq(GSE53233)/Homer | 1e-8 | -1.976e+01 | 0.0000 | 240.0 | 26.26% | 17573.1 | 18.37% | motif file (matrix) | svg |
| 65 | 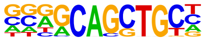 | Ascl2(bHLH)/ESC-Ascl2-ChIP-Seq(GSE97712)/Homer | 1e-8 | -1.897e+01 | 0.0000 | 154.0 | 16.85% | 10109.2 | 10.57% | motif file (matrix) | svg |
| 66 | 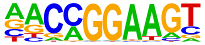 | ETV1(ETS)/GIST48-ETV1-ChIP-Seq(GSE22441)/Homer | 1e-7 | -1.841e+01 | 0.0000 | 160.0 | 17.51% | 10710.9 | 11.20% | motif file (matrix) | svg |
| 67 | | | 1e-7 | -1.819e+01 | 0.0000 | 56.0 | 6.13% | 2529.0 | 2.64% | motif file (matrix) | svg |

| | | | | | | | | | | | |
|---|---|---|---|---|---|---|---|---|---|---|---|
| | 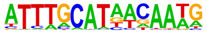 | OCT4-SOX2-TCF-NANOG(POU,Homeobox,HMG)/mES-Oct4-ChIP-Seq(GSE11431)/Homer | | | | | | | | | |
| 68 | 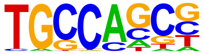 | HIC1(Zf)/Treg-ZBTB29-ChIP-Seq(GSE99889)/Homer | 1e-6 | -1.534e+01 | 0.0000 | 237.0 | 25.93% | 18224.6 | 19.06% | motif file (matrix) | svg |
| 69 | 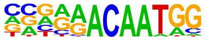 | Sox7(HMG)/ESC-Sox7-ChIP-Seq(GSE133899)/Homer | 1e-6 | -1.502e+01 | 0.0000 | 56.0 | 6.13% | 2790.6 | 2.92% | motif file (matrix) | svg |
| 70 | 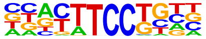 | Etv2(ETS)/ES-ER71-ChIP-Seq(GSE59402)/Homer | 1e-6 | -1.490e+01 | 0.0000 | 119.0 | 13.02% | 7780.5 | 8.14% | motif file (matrix) | svg |
| 71 | 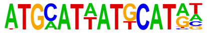 | OCT:OCT-short(POU,Homeobox)/NPC-OCT6-ChIP-Seq(GSE43916)/Homer | 1e-6 | -1.458e+01 | 0.0000 | 135.0 | 14.77% | 9195.7 | 9.61% | motif file (matrix) | svg |
| 72 | 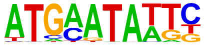 | Brn2(POU,Homeobox)/NPC-Brn2-ChIP-Seq(GSE35496)/Homer | 1e-6 | -1.445e+01 | 0.0000 | 32.0 | 3.50% | 1218.2 | 1.27% | motif file (matrix) | svg |
| 73 | 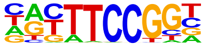 | Elk4(ETS)/Hela-Elk4-ChIP-Seq(GSE31477)/Homer | 1e-6 | -1.420e+01 | 0.0000 | 57.0 | 6.24% | 2939.6 | 3.07% | motif file (matrix) | svg |
| 74 | | Pit1(Homeobox)/GCrat-Pit1-ChIP-Seq(GSE58009)/Homer | 1e-6 | -1.395e+01 | 0.0000 | 175.0 | 19.15% | 12842.2 | 13.43% | motif file (matrix) | svg |

| | | | | | | | | | | | |
|---|---|---|---|---|---|---|---|---|---|---|---|
| | 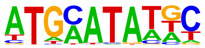 | | | | | | | | | | |
| 75 | 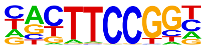 | Elk1(ETS)/Hela-Elk1-ChIP-Seq(GSE31477)/Homer | 1e-6 | -1.385e+01 | 0.0000 | 57.0 | 6.24% | 2973.2 | 3.11% | motif file (matrix) | svg |
| 76 | 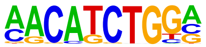 | ZBTB18(Zf)/HEK293-ZBTB18.GFP-ChIP-Seq(GSE58341)/Homer | 1e-5 | -1.338e+01 | 0.0000 | 74.0 | 8.10% | 4320.8 | 4.52% | motif file (matrix) | svg |
| 77 | 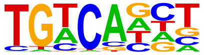 | Tgif2(Homeobox)/mES-Tgif2-ChIP-Seq(GSE55404)/Homer | 1e-5 | -1.233e+01 | 0.0000 | 396.0 | 43.33% | 34553.5 | 36.13% | motif file (matrix) | svg |
| 78 | 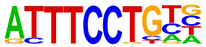 | EWS:ERG-fusion(ETS)/CADO_ES1-EWS:ERG-ChIP-Seq(SRA014231)/Homer | 1e-5 | -1.216e+01 | 0.0000 | 111.0 | 12.14% | 7550.0 | 7.89% | motif file (matrix) | svg |
| 79 | 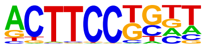 | Elf4(ETS)/BMDM-Elf4-ChIP-Seq(GSE88699)/Homer | 1e-5 | -1.206e+01 | 0.0000 | 117.0 | 12.80% | 8084.9 | 8.45% | motif file (matrix) | svg |
| 80 | 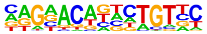 | PR(NR)/T47D-PR-ChIP-Seq(GSE31130)/Homer | 1e-5 | -1.193e+01 | 0.0000 | 252.0 | 27.57% | 20494.6 | 21.43% | motif file (matrix) | svg |
| 81 | | Hoxc6(Homeobox)/EB-Hoxc6.iFlag-ChIP-Seq(GSE142377)/Homer | 1e-5 | -1.156e+01 | 0.0001 | 422.0 | 46.17% | 37451.5 | 39.16% | motif file (matrix) | svg |

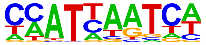

| | | | | | | | | | | | |
|---|---|---|---|---|---|---|---|---|---|---|---|
| 82 | 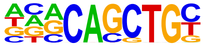 | E2A(bHLH)/proBcell-E2A-ChIP-Seq(GSE21978)/Homer | 1e-4 | -1.096e+01 | 0.0001 | 180.0 | 19.69% | 13969.7 | 14.61% | motif file (matrix) | svg |
| 83 | 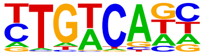 | Tgif1(Homeobox)/mES-Tgif1-ChIP-Seq(GSE55404)/Homer | 1e-4 | -1.086e+01 | 0.0001 | 374.0 | 40.92% | 32824.3 | 34.32% | motif file (matrix) | svg |
| 84 | 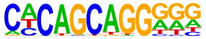 | Unknown-ESC-element(?)/mES-Nanog-ChIP-Seq(GSE11724)/Homer | 1e-4 | -1.066e+01 | 0.0001 | 68.0 | 7.44% | 4195.1 | 4.39% | motif file (matrix) | svg |
| 85 | 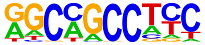 | ZNF91(Zf)/HEK-ZNF91.HA-ChIP-Seq(GSE162571)/Homer | 1e-4 | -1.065e+01 | 0.0001 | 80.0 | 8.75% | 5180.6 | 5.42% | motif file (matrix) | svg |
| 86 | 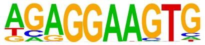 | PU.1(ETS)/ThioMac-PU.1-ChIP-Seq(GSE21512)/Homer | 1e-4 | -1.026e+01 | 0.0002 | 64.0 | 7.00% | 3927.6 | 4.11% | motif file (matrix) | svg |
| 87 | 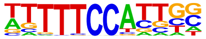 | NFATC2(RHD)/Islets-NFATC2-ChIP-Seq(GSE158496)/Homer | 1e-4 | -9.922e+00 | 0.0003 | 269.0 | 29.43% | 22730.7 | 23.77% | motif file (matrix) | svg |
| 88 | | Oct2(POU,Homeobox)/Bcell-Oct2-ChIP-Seq(GSE21512)/Homer | 1e-3 | -9.209e+00 | 0.0005 | 66.0 | 7.22% | 4241.1 | 4.43% | motif file (matrix) | svg |

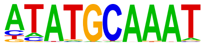

| | | | | | | | | | | | |
|---|---|---|---|---|---|---|---|---|---|---|---|
| 89 | 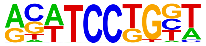 | SPDEF(ETS)/VCaP-SPDEF-ChIP-Seq(SRA014231)/Homer | 1e-3 | -8.788e+00 | 0.0008 | 114.0 | 12.47% | 8468.2 | 8.85% | motif file (matrix) | svg |
| 90 | 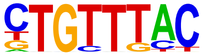 | Foxo1(Forkhead)/RAW-Foxo1-ChIP-Seq(Fan_et_al.)/Homer | 1e-3 | -8.631e+00 | 0.0009 | 239.0 | 26.15% | 20233.6 | 21.16% | motif file (matrix) | svg |
| 91 | 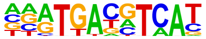 | CREB5(bZIP)/LNCaP-CREB5.V5-ChIP-Seq(GSE137775)/Homer | 1e-3 | -8.631e+00 | 0.0009 | 60.0 | 6.56% | 3834.0 | 4.01% | motif file (matrix) | svg |
| 92 | 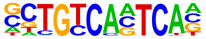 | Pknox1(Homeobox)/ES-Prep1-ChIP-Seq(GSE63282)/Homer | 1e-3 | -8.598e+00 | 0.0009 | 32.0 | 3.50% | 1649.2 | 1.72% | motif file (matrix) | svg |
| 93 | 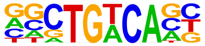 | Meis1(Homeobox)/MastCells-Meis1-ChIP-Seq(GSE48085)/Homer | 1e-3 | -8.148e+00 | 0.0015 | 186.0 | 20.35% | 15298.8 | 16.00% | motif file (matrix) | svg |
| 94 | 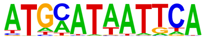 | Pit1+1bp(Homeobox)/GCrat-Pit1-ChIP-Seq(GSE58009)/Homer | 1e-3 | -8.133e+00 | 0.0015 | 70.0 | 7.66% | 4750.5 | 4.97% | motif file (matrix) | svg |
| 95 | | Pdx1(Homeobox)/Islet-Pdx1-ChIP-Seq(SRA008281)/Homer | 1e-3 | -8.027e+00 | 0.0016 | 145.0 | 15.86% | 11482.6 | 12.01% | motif file (matrix) | svg |

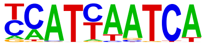

| | | | | | | | | | | | |
|---|---|---|---|---|---|---|---|---|---|---|---|
| 96 | 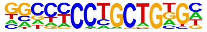 | Zic3(Zf)/mES-Zic3-ChIP-Seq(GSE37889)/Homer | 1e-3 | -7.823e+00 | 0.0020 | 54.0 | 5.91% | 3462.4 | 3.62% | motif file (matrix) | svg |
| 97 | 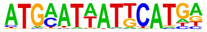 | OCT:OCT(POU,Homeobox,IR1)/NPC-Brn2-ChIP-Seq(GSE35496)/Homer | 1e-3 | -7.725e+00 | 0.0021 | 8.0 | 0.88% | 182.7 | 0.19% | motif file (matrix) | svg |
| 98 | 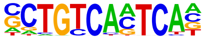 | Pbx3(Homeobox)/GM12878-PBX3-ChIP-Seq(GSE32465)/Homer | 1e-3 | -7.452e+00 | 0.0028 | 32.0 | 3.50% | 1765.4 | 1.85% | motif file (matrix) | svg |
| 99 | 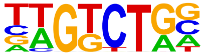 | Smad3(MAD)/NPC-Smad3-ChIP-Seq(GSE36673)/Homer | 1e-3 | -7.032e+00 | 0.0042 | 297.0 | 32.49% | 26533.1 | 27.74% | motif file (matrix) | svg |
| 100 | 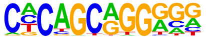 | Zic2(Zf)/ESC-Zic2-ChIP-Seq(SRP197560)/Homer | 1e-2 | -6.867e+00 | 0.0049 | 42.0 | 4.60% | 2622.7 | 2.74% | motif file (matrix) | svg |
| 101 | 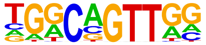 | AMYB(HTH)/Testes-AMYB-ChIP-Seq(GSE44588)/Homer | 1e-2 | -6.821e+00 | 0.0051 | 169.0 | 18.49% | 14098.0 | 14.74% | motif file (matrix) | svg |
| 102 | | HLF(bZIP)/HSC-HLF.Flag-ChIP-Seq(GSE69817)/Homer | 1e-2 | -6.677e+00 | 0.0058 | 125.0 | 13.68% | 9997.6 | 10.45% | motif file (matrix) | svg |

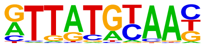

| | | | | | | | | | | | |
|---|---|---|---|---|---|---|---|---|---|---|---|
| 103 | 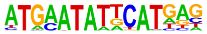 | OCT:OCT(POU,Homeobox)/NPC-Brn1-ChIP-Seq(GSE35496)/Homer | 1e-2 | -6.608e+00 | 0.0062 | 5.0 | 0.55% | 83.0 | 0.09% | motif file (matrix) | svg |
| 104 | 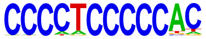 | Zfp281(Zf)/ES-Zfp281-ChIP-Seq(GSE81042)/Homer | 1e-2 | -6.405e+00 | 0.0075 | 16.0 | 1.75% | 709.0 | 0.74% | motif file (matrix) | svg |
| 105 | 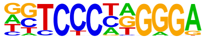 | EBF(EBF)/proBcell-EBF-ChIP-Seq(GSE21978)/Homer | 1e-2 | -6.330e+00 | 0.0080 | 22.0 | 2.41% | 1135.7 | 1.19% | motif file (matrix) | svg |
| 106 | 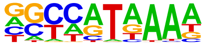 | Hoxd11(Homeobox)/ChickenMSG-Hoxd11.Flag-ChIP-Seq(GSE86088)/Homer | 1e-2 | -6.323e+00 | 0.0080 | 332.0 | 36.32% | 30356.3 | 31.74% | motif file (matrix) | svg |
| 107 | 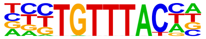 | FOXP1(Forkhead)/H9-FOXP1-ChIP-Seq(GSE31006)/Homer | 1e-2 | -6.248e+00 | 0.0085 | 62.0 | 6.78% | 4409.6 | 4.61% | motif file (matrix) | svg |
| 108 | 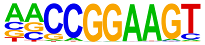 | ETS(ETS)/Promoter/Homer | 1e-2 | -6.021e+00 | 0.0106 | 31.0 | 3.39% | 1855.5 | 1.94% | motif file (matrix) | svg |
| 109 | | Hoxd10(Homeobox)/ChickenMSG-Hoxd10.Flag-ChIP-Seq(GSE86088)/Homer | 1e-2 | -6.002e+00 | 0.0107 | 165.0 | 18.05% | 13990.6 | 14.63% | motif file (matrix) | svg |

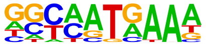

| | | | | | | | | | | | |
|---|---|---|---|---|---|---|---|---|---|---|---|
| 110 | 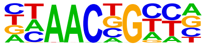 | BMYB(HTH)/Hela-BMYB-ChIP-Seq(GSE27030)/Homer | 1e-2 | -5.869e+00 | 0.0121 | 166.0 | 18.16% | 14133.7 | 14.78% | motif file (matrix) | svg |
| 111 | 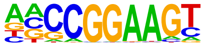 | ELF1(ETS)/Jurkat-ELF1-ChIP-Seq(SRA014231)/Homer | 1e-2 | -5.847e+00 | 0.0123 | 42.0 | 4.60% | 2775.4 | 2.90% | motif file (matrix) | svg |
| 112 | 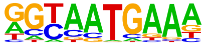 | Hoxa10(Homeobox)/ChickenMSG-Hoxa10.Flag-ChIP-Seq(GSE86088)/Homer | 1e-2 | -5.767e+00 | 0.0132 | 89.0 | 9.74% | 6927.9 | 7.24% | motif file (matrix) | svg |
| 113 | 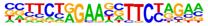 | HRE(HSF)/HepG2-HSF1-ChIP-Seq(GSE31477)/Homer | 1e-2 | -5.683e+00 | 0.0142 | 19.0 | 2.08% | 977.8 | 1.02% | motif file (matrix) | svg |
| 114 | 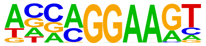 | EHF(ETS)/LoVo-EHF-ChIP-Seq(GSE49402)/Homer | 1e-2 | -5.671e+00 | 0.0143 | 144.0 | 15.75% | 12096.2 | 12.65% | motif file (matrix) | svg |
| 115 | 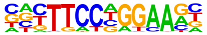 | Stat3+il21(Stat)/CD4-Stat3-ChIP-Seq(GSE19198)/Homer | 1e-2 | -5.586e+00 | 0.0154 | 79.0 | 8.64% | 6064.8 | 6.34% | motif file (matrix) | svg |
| 116 | | Zic(Zf)/Cerebellum-ZIC1.2-ChIP-Seq(GSE60731)/Homer | 1e-2 | -5.507e+00 | 0.0165 | 80.0 | 8.75% | 6174.8 | 6.46% | motif file (matrix) | svg |

| | | | | | | | | | | | |
|---|---|---|---|---|---|---|---|---|---|---|---|
| | 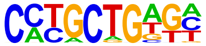 | | | | | | | | | | |
| 117 | 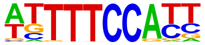 | NFAT(RHD)/Jurkat-NFATC1-ChIP-Seq(Jolma_et_al.)/Homer | 1e-2 | -5.285e+00 | 0.0204 | 119.0 | 13.02% | 9853.4 | 10.30% | motif file (matrix) | svg |
| 118 | 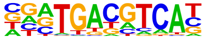 | Atf2(bZIP)/3T3L1-Atf2-ChIP-Seq(GSE56872)/Homer | 1e-2 | -5.247e+00 | 0.0211 | 40.0 | 4.38% | 2706.0 | 2.83% | motif file (matrix) | svg |
| 119 | 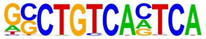 | PBX1(Homeobox)/MCF7-PBX1-ChIP-Seq(GSE28007)/Homer | 1e-2 | -5.182e+00 | 0.0223 | 12.0 | 1.31% | 527.8 | 0.55% | motif file (matrix) | svg |
| 120 | 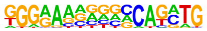 | RBPJ:Ebox(?,bHLH)/Panc1-Rbpj1-ChIP-Seq(GSE47459)/Homer | 1e-2 | -5.076e+00 | 0.0246 | 36.0 | 3.94% | 2398.3 | 2.51% | motif file (matrix) | svg |
| 121 | 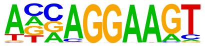 | ELF3(ETS)/PDAC-ELF3-ChIP-Seq(GSE64557)/Homer | 1e-2 | -4.904e+00 | 0.0289 | 85.0 | 9.30% | 6792.8 | 7.10% | motif file (matrix) | svg |
| 122 | 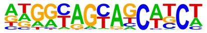 | PRDM9(Zf)/Testis-DMC1-ChIP-Seq(GSE35498)/Homer | 1e-2 | -4.876e+00 | 0.0295 | 51.0 | 5.58% | 3725.7 | 3.90% | motif file (matrix) | svg |
| 123 | | EBF2(EBF)/BrownAdipose-EBF2-ChIP-Seq(GSE97114)/Homer | 1e-2 | -4.652e+00 | 0.0366 | 79.0 | 8.64% | 6310.0 | 6.60% | motif file (matrix) | svg |

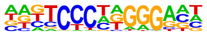

### Motifs found for: white matter invasion

### Homer Known Motif Enrichment Results (homerpos10)

[Homer *de novo* Motif Results](#)
[Gene Ontology Enrichment Results](#)
[Known Motif Enrichment Results (txt file)](#)
Total Target Sequences = 598, Total Background Sequences = 96724

| Rank | Motif | Name | P-value | log P-pvalue | q-value (Benjamini) | # Target Sequences with Motif | % of Targets Sequences with Motif | # Background Sequences with Motif | % of Background Sequences with Motif | Motif File | SVG |
|---|---|---|---|---|---|---|---|---|---|---|---|
| 1 | 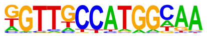 | X-box(HTH)/NPC-H3K4me1-ChIP-Seq(GSE16256)/Homer | 1e-39 | -9.146e+01 | 0.0000 | 55.0 | 9.20% | 735.1 | 0.76% | motif file (matrix) | svg |
| 2 | 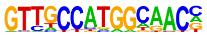 | Rfx2(HTH)/LoVo-RFX2-ChIP-Seq(GSE49402)/Homer | 1e-36 | -8.356e+01 | 0.0000 | 46.0 | 7.69% | 522.5 | 0.54% | motif file (matrix) | svg |
| 3 | 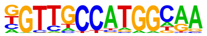 | Rfx1(HTH)/NPC-H3K4me1-ChIP-Seq(GSE16256)/Homer | 1e-35 | -8.275e+01 | 0.0000 | 67.0 | 11.20% | 1438.9 | 1.49% | motif file (matrix) | svg |
| 4 | | RFX(HTH)/K562-RFX3-ChIP-Seq(SRA012198)/Homer | 1e-33 | -7.733e+01 | 0.0000 | 41.0 | 6.86% | 433.8 | 0.45% | motif file (matrix) | svg |

| | | | | | | | | | | | |
|---|---|---|---|---|---|---|---|---|---|---|---|
| | 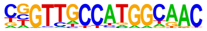 | | | | | | | | | | |
| 5 | 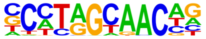 | Rfx5(HTH)/GM12878-Rfx5-ChIP-Seq(GSE31477)/Homer | 1e-28 | -6.480e+01 | 0.0000 | 76.0 | 12.71% | 2561.2 | 2.66% | motif file (matrix) | svg |
| 6 | 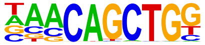 | Tcf21(bHLH)/ArterySmoothMuscle-Tcf21-ChIP-Seq(GSE61369)/Homer | 1e-23 | -5.457e+01 | 0.0000 | 136.0 | 22.74% | 8544.2 | 8.87% | motif file (matrix) | svg |
| 7 | 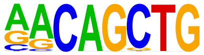 | MyoG(bHLH)/C2C12-MyoG-ChIP-Seq(GSE36024)/Homer | 1e-21 | -4.870e+01 | 0.0000 | 137.0 | 22.91% | 9229.7 | 9.59% | motif file (matrix) | svg |
| 8 | 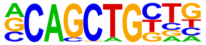 | Tcf12(bHLH)/GM12878-Tcf12-ChIP-Seq(GSE32465)/Homer | 1e-19 | -4.474e+01 | 0.0000 | 127.0 | 21.24% | 8565.7 | 8.90% | motif file (matrix) | svg |
| 9 | 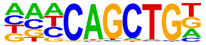 | Ap4(bHLH)/AML-Tfap4-ChIP-Seq(GSE45738)/Homer | 1e-19 | -4.439e+01 | 0.0000 | 149.0 | 24.92% | 11048.3 | 11.47% | motif file (matrix) | svg |
| 10 | 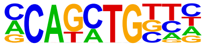 | Twist2(bHLH)/Myoblast-Twist2.Ty1-ChIP-Seq(GSE127998)/Homer | 1e-18 | -4.182e+01 | 0.0000 | 190.0 | 31.77% | 16341.0 | 16.97% | motif file (matrix) | svg |
| 11 | | Myf5(bHLH)/GM-Myf5-ChIP-Seq(GSE24852)/Homer | 1e-17 | -4.116e+01 | 0.0000 | 101.0 | 16.89% | 6209.8 | 6.45% | motif file (matrix) | svg |

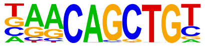

| | | | | | | | | | | | |
|---|---|---|---|---|---|---|---|---|---|---|---|
| 12 | 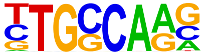 | NF1-halfsite(CTF)/LNCaP-NF1-ChIP-Seq(Unpublished)/Homer | 1e-15 | -3.529e+01 | 0.0000 | 187.0 | 31.27% | 17008.0 | 17.66% | motif file (matrix) | svg |
| 13 | 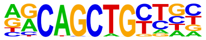 | MyoD(bHLH)/Myotube-MyoD-ChIP-Seq(GSE21614)/Homer | 1e-15 | -3.466e+01 | 0.0000 | 99.0 | 16.56% | 6646.2 | 6.90% | motif file (matrix) | svg |
| 14 | 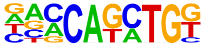 | BHLHA15(bHLH)/NIH3T3-BHLHB8.HA-ChIP-Seq(GSE119782)/Homer | 1e-15 | -3.456e+01 | 0.0000 | 152.0 | 25.42% | 12722.8 | 13.21% | motif file (matrix) | svg |
| 15 | 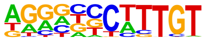 | Sox9(HMG)/Limb-SOX9-ChIP-Seq(GSE73225)/Homer | 1e-14 | -3.397e+01 | 0.0000 | 110.0 | 18.39% | 7911.4 | 8.22% | motif file (matrix) | svg |
| 16 | 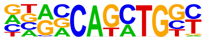 | Atoh1(bHLH)/Cerebellum-Atoh1-ChIP-Seq(GSE22111)/Homer | 1e-14 | -3.360e+01 | 0.0000 | 126.0 | 21.07% | 9766.5 | 10.14% | motif file (matrix) | svg |
| 17 | 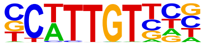 | Sox10(HMG)/SciaticNerve-Sox3-ChIP-Seq(GSE35132)/Homer | 1e-14 | -3.293e+01 | 0.0000 | 183.0 | 30.60% | 16898.1 | 17.55% | motif file (matrix) | svg |
| 18 | | Rfx6(HTH)/Min6b1-Rfx6.HA-ChIP-Seq(GSE62844)/Homer | 1e-13 | -3.166e+01 | 0.0000 | 116.0 | 19.40% | 8872.8 | 9.21% | motif file (matrix) | svg |

| | | | | | | | | | | | |
|---|---|---|---|---|---|---|---|---|---|---|---|
| | 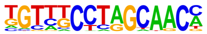 | | | | | | | | | | |
| 19 | 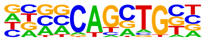 | Ascl1(bHLH)/NeuralTubes-Ascl1-ChIP-Seq(GSE55840)/Homer | 1e-13 | -2.996e+01 | 0.0000 | 158.0 | 26.42% | 14205.8 | 14.75% | motif file (matrix) | svg |
| 20 | 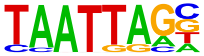 | Lhx2(Homeobox)/HFSC-Lhx2-ChIP-Seq(GSE48068)/Homer | 1e-12 | -2.890e+01 | 0.0000 | 136.0 | 22.74% | 11640.4 | 12.09% | motif file (matrix) | svg |
| 21 | 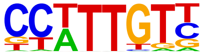 | Sox3(HMG)/NPC-Sox3-ChIP-Seq(GSE33059)/Homer | 1e-12 | -2.783e+01 | 0.0000 | 190.0 | 31.77% | 18791.6 | 19.52% | motif file (matrix) | svg |
| 22 | 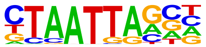 | Gsx2(Homeobox)/LGE-Gsx2.Flag-ChIP-Seq(GSE162589)/Homer | 1e-11 | -2.755e+01 | 0.0000 | 161.0 | 26.92% | 15017.0 | 15.60% | motif file (matrix) | svg |
| 23 | 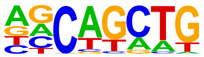 | SCL(bHLH)/HPC7-Scl-ChIP-Seq(GSE13511)/Homer | 1e-11 | -2.698e+01 | 0.0000 | 353.0 | 59.03% | 43122.0 | 44.78% | motif file (matrix) | svg |
| 24 | 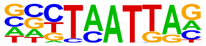 | Emx2(Homeobox)/Cortex-Emx2-ChIP-Seq(GSE183130)/Homer | 1e-11 | -2.586e+01 | 0.0000 | 139.0 | 23.24% | 12517.9 | 13.00% | motif file (matrix) | svg |
| 25 | | LHX9(Homeobox)/Hct116-LHX9.V5-ChIP-Seq(GSE116822)/Homer | 1e-10 | -2.462e+01 | 0.0000 | 159.0 | 26.59% | 15317.1 | 15.91% | motif file (matrix) | svg |

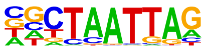

| | | | | | | | | | | | |
|---|---|---|---|---|---|---|---|---|---|---|---|
| 26 | 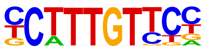 | Sox4(HMG)/proB-Sox4-ChIP-Seq(GSE50066)/Homer | 1e-10 | -2.420e+01 | 0.0000 | 105.0 | 17.56% | 8639.9 | 8.97% | motif file (matrix) | svg |
| 27 | 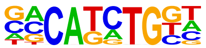 | TCF4(bHLH)/SHSY5Y-TCF4-ChIP-Seq(GSE96915)/Homer | 1e-10 | -2.378e+01 | 0.0000 | 149.0 | 24.92% | 14180.1 | 14.73% | motif file (matrix) | svg |
| 28 | 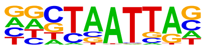 | En1(Homeobox)/SUM149-EN1-ChIP-Seq(GSE120957)/Homer | 1e-10 | -2.352e+01 | 0.0000 | 194.0 | 32.44% | 20271.2 | 21.05% | motif file (matrix) | svg |
| 29 | 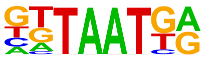 | Nkx6.1(Homeobox)/Islet-Nkx6.1-ChIP-Seq(GSE40975)/Homer | 1e-10 | -2.342e+01 | 0.0000 | 254.0 | 42.47% | 28871.4 | 29.98% | motif file (matrix) | svg |
| 30 | 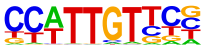 | SOX1(HMG)/NPC-SOX1-ChIP-Seq(GSE138215)/Homer | 1e-10 | -2.321e+01 | 0.0000 | 223.0 | 37.29% | 24423.4 | 25.36% | motif file (matrix) | svg |
| 31 | 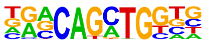 | Atoh7(bHLH)/Retina-Atoh7-CutnRun(GSE156756)/Homer | 1e-10 | -2.312e+01 | 0.0000 | 88.0 | 14.72% | 6824.4 | 7.09% | motif file (matrix) | svg |
| 32 | | NeuroG2(bHLH)/Fibroblast-NeuroG2-ChIP-Seq(GSE75910)/Homer | 1e-10 | -2.310e+01 | 0.0000 | 149.0 | 24.92% | 14314.3 | 14.87% | motif file (matrix) | svg |

| | | | | | | | | | | | |
|---|---|---|---|---|---|---|---|---|---|---|---|
| | 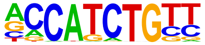 | | | | | | | | | | |
| 33 | 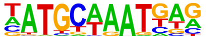 | Oct6(POU,Homeobox)/NPC-Pou3f1-ChIP-Seq(GSE35496)/Homer | 1e-9 | -2.296e+01 | 0.0000 | 65.0 | 10.87% | 4338.8 | 4.51% | motif file (matrix) | svg |
| 34 | 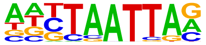 | Lhx3(Homeobox)/Neuron-Lhx3-ChIP-Seq(GSE31456)/Homer | 1e-9 | -2.234e+01 | 0.0000 | 182.0 | 30.43% | 18892.0 | 19.62% | motif file (matrix) | svg |
| 35 | 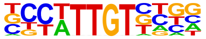 | Sox21(HMG)/ESC-SOX21-ChIP-Seq(GSE110505)/Homer | 1e-8 | -2.045e+01 | 0.0000 | 181.0 | 30.27% | 19200.0 | 19.94% | motif file (matrix) | svg |
| 36 | 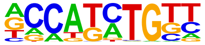 | Olig2(bHLH)/Neuron-Olig2-ChIP-Seq(GSE30882)/Homer | 1e-8 | -1.924e+01 | 0.0000 | 175.0 | 29.26% | 18670.5 | 19.39% | motif file (matrix) | svg |
| 37 | 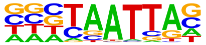 | DLX2(Homeobox)/BasalGanglia-Dlx2-ChIP-seq(GSE124936)/Homer | 1e-8 | -1.879e+01 | 0.0000 | 163.0 | 27.26% | 17131.9 | 17.79% | motif file (matrix) | svg |
| 38 | 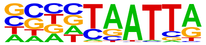 | DLX1(Homeobox)/BasalGanglia-Dlx1-ChIP-seq(GSE124936)/Homer | 1e-8 | -1.877e+01 | 0.0000 | 149.0 | 24.92% | 15241.0 | 15.83% | motif file (matrix) | svg |
| 39 | | Ascl2(bHLH)/ESC-Ascl2-ChIP-Seq(GSE97712)/Homer | 1e-7 | -1.745e+01 | 0.0000 | 116.0 | 19.40% | 11192.9 | 11.62% | motif file (matrix) | svg |

| | | | | | | | | | | |
|---|---|---|---|---|---|---|---|---|---|---|
| | 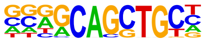 | | | | | | | | | |
| 40 | Lhx1(Homeobox)/EmbryoCarcinoma-Lhx1-ChIP-Seq(GSE70957)/Homer 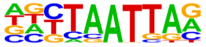 | 1e-7 | -1.681e+01 | 0.0000 | 129.0 | 21.57% | 13027.6 | 13.53% | motif file (matrix) | svg |
| 41 | Isl1(Homeobox)/Neuron-Isl1-ChIP-Seq(GSE31456)/Homer 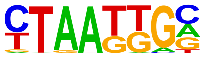 | 1e-7 | -1.636e+01 | 0.0000 | 182.0 | 30.43% | 20416.3 | 21.20% | motif file (matrix) | svg |
| 42 | Brn1(POU,Homeobox)/NPC-Brn1-ChIP-Seq(GSE35496)/Homer 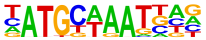 | 1e-6 | -1.597e+01 | 0.0000 | 47.0 | 7.86% | 3236.4 | 3.36% | motif file (matrix) | svg |
| 43 | Sox6(HMG)/Myotubes-Sox6-ChIP-Seq(GSE32627)/Homer 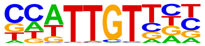 | 1e-6 | -1.583e+01 | 0.0000 | 159.0 | 26.59% | 17338.2 | 18.01% | motif file (matrix) | svg |
| 44 | NF1(CTF)/LNCAP-NF1-ChIP-Seq(Unpublished)/Homer 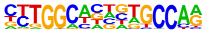 | 1e-6 | -1.544e+01 | 0.0000 | 46.0 | 7.69% | 3191.1 | 3.31% | motif file (matrix) | svg |
| 45 | STAT4(Stat)/CD4-Stat4-ChIP-Seq(GSE22104)/Homer 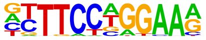 | 1e-6 | -1.511e+01 | 0.0000 | 100.0 | 16.72% | 9629.5 | 10.00% | motif file (matrix) | svg |
| 46 | E2A(bHLH)/proBcell-E2A-ChIP-Seq(GSE21978)/Homer | 1e-6 | -1.484e+01 | 0.0000 | 143.0 | 23.91% | 15398.1 | 15.99% | motif file (matrix) | svg |

| | | | | | | | | | | | |
|---|---|---|---|---|---|---|---|---|---|---|---|
| | 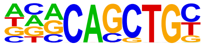 | | | | | | | | | | |
| 47 | 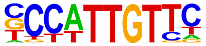 | Sox2(HMG)/mES-Sox2-ChIP-Seq(GSE11431)/Homer | 1e-6 | -1.463e+01 | 0.0000 | 95.0 | 15.89% | 9092.4 | 9.44% | motif file (matrix) | svg |
| 48 | 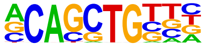 | Ptf1a(bHLH)/Panc1-Ptf1a-ChIP-Seq(GSE47459)/Homer | 1e-6 | -1.433e+01 | 0.0000 | 220.0 | 36.79% | 26556.5 | 27.58% | motif file (matrix) | svg |
| 49 | 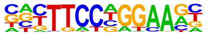 | Stat3+il21(Stat)/CD4-Stat3-ChIP-Seq(GSE19198)/Homer | 1e-6 | -1.403e+01 | 0.0000 | 71.0 | 11.87% | 6232.7 | 6.47% | motif file (matrix) | svg |
| 50 | 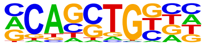 | HEB(bHLH)/mES-Heb-ChIP-Seq(GSE53233)/Homer | 1e-5 | -1.351e+01 | 0.0000 | 168.0 | 28.09% | 19270.1 | 20.01% | motif file (matrix) | svg |
| 51 | 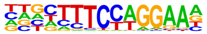 | Bcl6(Zf)/Liver-Bcl6-ChIP-Seq(GSE31578)/Homer | 1e-5 | -1.336e+01 | 0.0000 | 122.0 | 20.40% | 12932.3 | 13.43% | motif file (matrix) | svg |
| 52 | 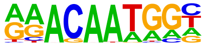 | Sox15(HMG)/CPA-Sox15-ChIP-Seq(GSE62909)/Homer | 1e-5 | -1.329e+01 | 0.0000 | 115.0 | 19.23% | 12009.9 | 12.47% | motif file (matrix) | svg |
| 53 | | NFATC2(RHD)/Islets-NFATC2-ChIP-Seq(GSE158496)/Homer | 1e-5 | -1.327e+01 | 0.0000 | 190.0 | 31.77% | 22511.3 | 23.38% | motif file (matrix) | svg |

| | | | | | | | | | | | |
|---|---|---|---|---|---|---|---|---|---|---|---|
| | 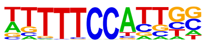 | | | | | | | | | | |
| 54 | 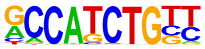 | NeuroD1(bHLH)/Islet-NeuroD1-ChIP-Seq(GSE30298)/Homer | 1e-5 | -1.277e+01 | 0.0000 | 77.0 | 12.88% | 7203.5 | 7.48% | motif file (matrix) | svg |
| 55 | 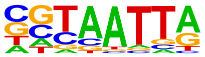 | DLX5(Homeobox)/BasalGanglia-Dlx5-ChIP-seq(GSE124936)/Homer | 1e-5 | -1.223e+01 | 0.0000 | 89.0 | 14.88% | 8844.3 | 9.19% | motif file (matrix) | svg |
| 56 | 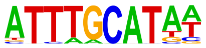 | Oct4(POU,Homeobox)/mES-Oct4-ChIP-Seq(GSE11431)/Homer | 1e-5 | -1.220e+01 | 0.0000 | 60.0 | 10.03% | 5232.7 | 5.43% | motif file (matrix) | svg |
| 57 | 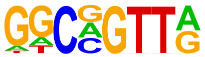 | MYB(HTH)/ERMYB-Myb-ChIPSeq(GSE22095)/Homer | 1e-5 | -1.215e+01 | 0.0000 | 139.0 | 23.24% | 15602.6 | 16.20% | motif file (matrix) | svg |
| 58 | 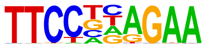 | STAT6(Stat)/Macrophage-Stat6-ChIP-Seq(GSE38377)/Homer | 1e-4 | -1.111e+01 | 0.0001 | 61.0 | 10.20% | 5552.0 | 5.77% | motif file (matrix) | svg |
| 59 | 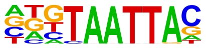 | Dlx3(Homeobox)/Kerainocytes-Dlx3-ChIP-Seq(GSE89884)/Homer | 1e-4 | -1.097e+01 | 0.0001 | 76.0 | 12.71% | 7458.0 | 7.75% | motif file (matrix) | svg |
| 60 | | STAT6(Stat)/CD4-Stat6-ChIP-Seq(GSE22104)/Homer | 1e-4 | -1.070e+01 | 0.0002 | 61.0 | 10.20% | 5630.8 | 5.85% | motif file (matrix) | svg |

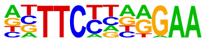

| | | | | | | | | | | | |
|---|---|---|---|---|---|---|---|---|---|---|---|
| 61 | 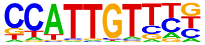 | Sox17(HMG)/Endoderm-Sox17-ChIP-Seq(GSE61475)/Homer | 1e-4 | -1.043e+01 | 0.0002 | 73.0 | 12.21% | 7194.4 | 7.47% | motif file (matrix) | svg |
| 62 | 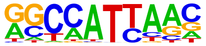 | Nanog(Homeobox)/mES-Nanog-ChIP-Seq(GSE11724)/Homer | 1e-4 | -9.895e+00 | 0.0004 | 321.0 | 53.68% | 43962.7 | 45.66% | motif file (matrix) | svg |
| 63 | 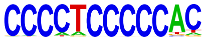 | Zfp281(Zf)/ES-Zfp281-ChIP-Seq(GSE81042)/Homer | 1e-4 | -9.724e+00 | 0.0004 | 18.0 | 3.01% | 979.6 | 1.02% | motif file (matrix) | svg |
| 64 | 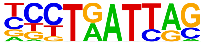 | Lhx6/Neurons-Lhx6-ChIP-seq(GSE85704)/Homer | 1e-4 | -9.653e+00 | 0.0005 | 116.0 | 19.40% | 13168.8 | 13.68% | motif file (matrix) | svg |
| 65 | 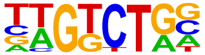 | Smad3(MAD)/NPC-Smad3-ChIP-Seq(GSE36673)/Homer | 1e-4 | -9.488e+00 | 0.0006 | 209.0 | 34.95% | 26746.2 | 27.78% | motif file (matrix) | svg |
| 66 | 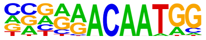 | Sox7(HMG)/ESC-Sox7-ChIP-Seq(GSE133899)/Homer | 1e-4 | -9.297e+00 | 0.0007 | 34.0 | 5.69% | 2673.7 | 2.78% | motif file (matrix) | svg |
| 67 | | Tlx?(NR)/NPC-H3K4me1-ChIP-Seq(GSE16256)/Homer | 1e-3 | -8.682e+00 | 0.0012 | 41.0 | 6.86% | 3580.9 | 3.72% | motif file (matrix) | svg |

| | | | | | | | | | | | |
|---|---|---|---|---|---|---|---|---|---|---|---|
| | 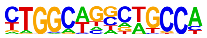 | | | | | | | | | | |
| 68 | 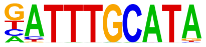 | Oct11(POU,Homeobox)/NCIH1048-POU2F3-ChIP-seq(GSE115123)/Homer | 1e-3 | -8.334e+00 | 0.0017 | 39.0 | 6.52% | 3403.9 | 3.54% | motif file (matrix) | svg |
| 69 | 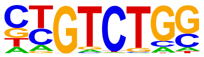 | Smad2(MAD)/ES-SMAD2-ChIP-Seq(GSE29422)/Homer | 1e-3 | -7.515e+00 | 0.0037 | 110.0 | 18.39% | 13051.2 | 13.55% | motif file (matrix) | svg |
| 70 | 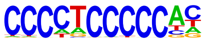 | ZNF148(Zf)/MDAMB231-ZNF148-ChIP-Seq(GSE147020)/Homer | 1e-3 | -7.076e+00 | 0.0057 | 39.0 | 6.52% | 3638.2 | 3.78% | motif file (matrix) | svg |
| 71 | 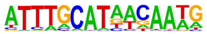 | OCT4-SOX2-TCF-NANOG(POU,Homeobox,HMG)/mES-Oct4-ChIP-Seq(GSE11431)/Homer | 1e-2 | -6.895e+00 | 0.0067 | 25.0 | 4.18% | 2008.8 | 2.09% | motif file (matrix) | svg |
| 72 | 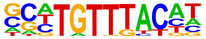 | FOXK2(Forkhead)/U2OS-FOXK2-ChIP-Seq(E-MTAB-2204)/Homer | 1e-2 | -6.582e+00 | 0.0091 | 57.0 | 9.53% | 6061.7 | 6.30% | motif file (matrix) | svg |
| 73 | 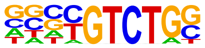 | Smad4(MAD)/ESC-SMAD4-ChIP-Seq(GSE29422)/Homer | 1e-2 | -6.388e+00 | 0.0109 | 107.0 | 17.89% | 13055.2 | 13.56% | motif file (matrix) | svg |
| 74 | | ZBTB18(Zf)/HEK293-ZBTB18.GFP-ChIP-Seq(GSE58341)/Homer | 1e-2 | -6.114e+00 | 0.0141 | 45.0 | 7.53% | 4611.0 | 4.79% | motif file (matrix) | svg |

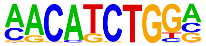

| | | | | | | | | | | | |
|---|---|---|---|---|---|---|---|---|---|---|---|
| 75 | 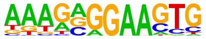 | SpiB(ETS)/OCILY3-SPIB-ChIP-Seq(GSE56857)/Homer | 1e-2 | -5.717e+00 | 0.0207 | 25.0 | 4.18% | 2197.8 | 2.28% | motif file (matrix) | svg |
| 76 | 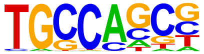 | HIC1(Zf)/Treg-ZBTB29-ChIP-Seq(GSE99889)/Homer | 1e-2 | -5.363e+00 | 0.0291 | 153.0 | 25.59% | 20298.8 | 21.08% | motif file (matrix) | svg |
| 77 | 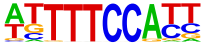 | NFAT(RHD)/Jurkat-NFATC1-ChIP-Seq(Jolma_et_al.)/Homer | 1e-2 | -5.350e+00 | 0.0291 | 78.0 | 13.04% | 9339.5 | 9.70% | motif file (matrix) | svg |
| 78 | 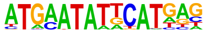 | OCT:OCT(POU,Homeobox)/NPC-Brn1-ChIP-Seq(GSE35496)/Homer | 1e-2 | -5.323e+00 | 0.0295 | 3.0 | 0.50% | 54.8 | 0.06% | motif file (matrix) | svg |
| 79 | 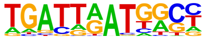 | Hoxb4(Homeobox)/ES-Hoxb4-ChIP-Seq(GSE34014)/Homer | 1e-2 | -5.259e+00 | 0.0311 | 19.0 | 3.18% | 1570.4 | 1.63% | motif file (matrix) | svg |
| 80 | 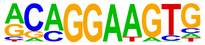 | ETS1(ETS)/Jurkat-ETS1-ChIP-Seq(GSE17954)/Homer | 1e-2 | -4.927e+00 | 0.0428 | 73.0 | 12.21% | 8797.8 | 9.14% | motif file (matrix) | svg |
| 81 | | AMYB(HTH)/Testes-AMYB-ChIP-Seq(GSE44588)/Homer | 1e-2 | -4.904e+00 | 0.0432 | 108.0 | 18.06% | 13853.5 | 14.39% | motif file (matrix) | svg |

| | | | | | | | | | | | |
|---|---|---|---|---|---|---|---|---|---|---|---|
| | 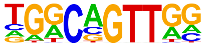 | | | | | | | | | | |
| 82 | 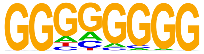 | Maz(Zf)/HepG2-Maz-ChIP-Seq(GSE31477)/Homer | 1e-2 | -4.872e+00 | 0.0441 | 65.0 | 10.87% | 7693.3 | 7.99% | motif file (matrix) | svg |
| 83 | 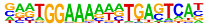 | NFAT:AP1(RHD,bZIP)/Jurkat-NFATC1-ChIP-Seq(Jolma_et_al.)/Homer | 1e-2 | -4.655e+00 | 0.0541 | 19.0 | 3.18% | 1668.1 | 1.73% | motif file (matrix) | svg |
| 84 | 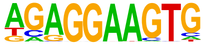 | PU.1(ETS)/ThioMac-PU.1-ChIP-Seq(GSE21512)/Homer | 1e-2 | -4.624e+00 | 0.0551 | 39.0 | 6.52% | 4217.7 | 4.38% | motif file (matrix) | svg |

### Motifs found for: age at diagnosis

### Homer Known Motif Enrichment Results (homerpos11)

[Homer *de novo* Motif Results](#)
[Gene Ontology Enrichment Results](#)
[Known Motif Enrichment Results (txt file)](#)
Total Target Sequences = 350, Total Background Sequences = 97581

| Rank | Motif | Name | P-value | log P-pvalue | q-value (Benjamini) | # Target Sequences with Motif | % of Targets Sequences with Motif | # Background Sequences with Motif | % of Background Sequences with Motif | Motif File | SVG |
|---|---|---|---|---|---|---|---|---|---|---|---|
| 1 | 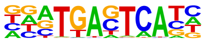 | Fra1(bZIP)/BT549-Fra1-ChIP-Seq(GSE46166)/Homer | 1e-48 | -1.122e+02 | 0.0000 | 103.0 | 29.43% | 4869.3 | 4.99% | motif file (matrix) | svg |
| 2 | 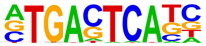 | AP-1(bZIP)/ThioMac-PU.1-ChIP-Seq(GSE21512)/Homer | 1e-47 | -1.085e+02 | 0.0000 | 118.0 | 33.71% | 6923.7 | 7.10% | motif file (matrix) | svg |
| 3 | 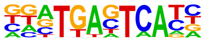 | Fos(bZIP)/TSC-Fos-ChIP-Seq(GSE110950)/Homer | 1e-46 | -1.074e+02 | 0.0000 | 104.0 | 29.71% | 5253.6 | 5.39% | motif file (matrix) | svg |
| 4 | | BATF(bZIP)/Th17-BATF-ChIP-Seq(GSE39756)/Homer | 1e-46 | -1.071e+02 | 0.0000 | 109.0 | 31.14% | 5869.3 | 6.02% | motif file (matrix) | svg |

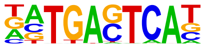

| | | | | | | | | | | | |
|---|---|---|---|---|---|---|---|---|---|---|---|
| 5 | 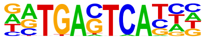 | Atf3(bZIP)/GBM-ATF3-ChIP-Seq(GSE33912)/Homer | 1e-45 | -1.047e+02 | 0.0000 | 109.0 | 31.14% | 6022.1 | 6.18% | motif file (matrix) | svg |
| 6 | 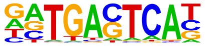 | JunB(bZIP)/DendriticCells-Junb-ChIP-Seq(GSE36099)/Homer | 1e-44 | -1.021e+02 | 0.0000 | 98.0 | 28.00% | 4862.4 | 4.99% | motif file (matrix) | svg |
| 7 | 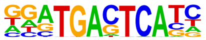 | Fra2(bZIP)/Striatum-Fra2-ChIP-Seq(GSE43429)/Homer | 1e-40 | -9.292e+01 | 0.0000 | 87.0 | 24.86% | 4137.8 | 4.24% | motif file (matrix) | svg |
| 8 | 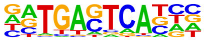 | Fosl2(bZIP)/3T3L1-Fosl2-ChIP-Seq(GSE56872)/Homer | 1e-36 | -8.450e+01 | 0.0000 | 69.0 | 19.71% | 2697.0 | 2.77% | motif file (matrix) | svg |
| 9 | 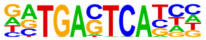 | Jun-AP1(bZIP)/K562-cJun-ChIP-Seq(GSE31477)/Homer | 1e-32 | -7.586e+01 | 0.0000 | 57.0 | 16.29% | 1960.5 | 2.01% | motif file (matrix) | svg |
| 10 | 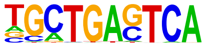 | Bach2(bZIP)/OCILy7-Bach2-ChIP-Seq(GSE44420)/Homer | 1e-20 | -4.659e+01 | 0.0000 | 39.0 | 11.14% | 1557.3 | 1.60% | motif file (matrix) | svg |
| 11 | | Sox21(HMG)/ESC-SOX21-ChIP-Seq(GSE110505)/Homer | 1e-16 | -3.894e+01 | 0.0000 | 134.0 | 38.29% | 18280.8 | 18.75% | motif file (matrix) | svg |

| | | | | | | | | | | | |
|---|---|---|---|---|---|---|---|---|---|---|---|
| | 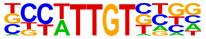 | | | | | | | | | | |
| 12 | 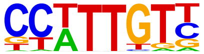 | Sox3(HMG)/NPC-Sox3-ChIP-Seq(GSE33059)/Homer | 1e-16 | -3.807e+01 | 0.0000 | 134.0 | 38.29% | 18470.6 | 18.94% | motif file (matrix) | svg |
| 13 | 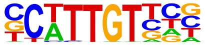 | Sox10(HMG)/SciaticNerve-Sox3-ChIP-Seq(GSE35132)/Homer | 1e-14 | -3.366e+01 | 0.0000 | 122.0 | 34.86% | 16873.0 | 17.30% | motif file (matrix) | svg |
| 14 | 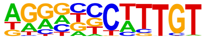 | Sox9(HMG)/Limb-SOX9-ChIP-Seq(GSE73225)/Homer | 1e-11 | -2.654e+01 | 0.0000 | 70.0 | 20.00% | 7969.8 | 8.17% | motif file (matrix) | svg |
| 15 | 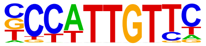 | Sox2(HMG)/mES-Sox2-ChIP-Seq(GSE11431)/Homer | 1e-11 | -2.567e+01 | 0.0000 | 73.0 | 20.86% | 8682.4 | 8.90% | motif file (matrix) | svg |
| 16 | 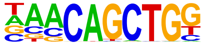 | Tcf21(bHLH)/ArterySmoothMuscle-Tcf21-ChIP-Seq(GSE61369)/Homer | 1e-11 | -2.558e+01 | 0.0000 | 74.0 | 21.14% | 8887.5 | 9.11% | motif file (matrix) | svg |
| 17 | 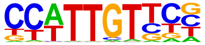 | SOX1(HMG)/NPC-SOX1-ChIP-Seq(GSE138215)/Homer | 1e-10 | -2.516e+01 | 0.0000 | 140.0 | 40.00% | 23160.0 | 23.75% | motif file (matrix) | svg |
| 18 | | BHLHA15(bHLH)/NIH3T3-BHLHB8.HA-ChIP-Seq(GSE119782)/Homer | 1e-10 | -2.369e+01 | 0.0000 | 96.0 | 27.43% | 13711.2 | 14.06% | motif file (matrix) | svg |

| | | | | | | | | | | | |
|---|---|---|---|---|---|---|---|---|---|---|---|
| | 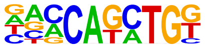 | | | | | | | | | | |
| 19 | 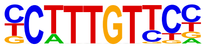 | Sox4(HMG)/proB-Sox4-ChIP-Seq(GSE50066)/Homer | 1e-10 | -2.339e+01 | 0.0000 | 69.0 | 19.71% | 8362.7 | 8.58% | motif file (matrix) | svg |
| 20 | 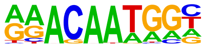 | Sox15(HMG)/CPA-Sox15-ChIP-Seq(GSE62909)/Homer | 1e-8 | -2.059e+01 | 0.0000 | 82.0 | 23.43% | 11554.5 | 11.85% | motif file (matrix) | svg |
| 21 | 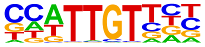 | Sox6(HMG)/Myotubes-Sox6-ChIP-Seq(GSE32627)/Homer | 1e-8 | -1.905e+01 | 0.0000 | 105.0 | 30.00% | 16969.7 | 17.40% | motif file (matrix) | svg |
| 22 | 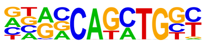 | Atoh1(bHLH)/Cerebellum-Atoh1-ChIP-Seq(GSE22111)/Homer | 1e-8 | -1.878e+01 | 0.0000 | 74.0 | 21.14% | 10352.1 | 10.62% | motif file (matrix) | svg |
| 23 | 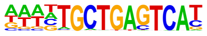 | Bach1(bZIP)/K562-Bach1-ChIP-Seq(GSE31477)/Homer | 1e-8 | -1.849e+01 | 0.0000 | 13.0 | 3.71% | 431.6 | 0.44% | motif file (matrix) | svg |
| 24 | 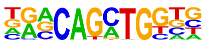 | Atoh7(bHLH)/Retina-Atoh7-CutnRun(GSE156756)/Homer | 1e-7 | -1.792e+01 | 0.0000 | 58.0 | 16.57% | 7372.7 | 7.56% | motif file (matrix) | svg |
| 25 | | Fli1(ETS)/CD8-FLI-ChIP-Seq(GSE20898)/Homer | 1e-7 | -1.759e+01 | 0.0000 | 66.0 | 18.86% | 9018.9 | 9.25% | motif file (matrix) | svg |

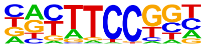

| | | | | | | | | | | | |
|---|---|---|---|---|---|---|---|---|---|---|---|
| 26 | 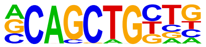 | Tcf12(bHLH)/GM12878-Tcf12-ChIP-Seq(GSE32465)/Homer | 1e-7 | -1.754e+01 | 0.0000 | 65.0 | 18.57% | 8830.4 | 9.06% | motif file (matrix) | svg |
| 27 | 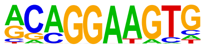 | ETS1(ETS)/Jurkat-ETS1-ChIP-Seq(GSE17954)/Homer | 1e-7 | -1.721e+01 | 0.0000 | 65.0 | 18.57% | 8907.0 | 9.13% | motif file (matrix) | svg |
| 28 | 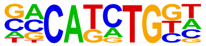 | TCF4(bHLH)/SHSY5Y-TCF4-ChIP-Seq(GSE96915)/Homer | 1e-7 | -1.660e+01 | 0.0000 | 95.0 | 27.14% | 15494.3 | 15.89% | motif file (matrix) | svg |
| 29 | 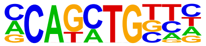 | Twist2(bHLH)/Myoblast-Twist2.Ty1-ChIP-Seq(GSE127998)/Homer | 1e-6 | -1.581e+01 | 0.0000 | 103.0 | 29.43% | 17582.3 | 18.03% | motif file (matrix) | svg |
| 30 | 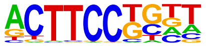 | Elf4(ETS)/BMDM-Elf4-ChIP-Seq(GSE88699)/Homer | 1e-6 | -1.555e+01 | 0.0000 | 62.0 | 17.71% | 8710.5 | 8.93% | motif file (matrix) | svg |
| 31 | 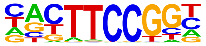 | Elk1(ETS)/Hela-Elk1-ChIP-Seq(GSE31477)/Homer | 1e-6 | -1.544e+01 | 0.0000 | 34.0 | 9.71% | 3485.4 | 3.57% | motif file (matrix) | svg |
| 32 | | MafK(bZIP)/C2C12-MafK-ChIP-Seq(GSE36030)/Homer | 1e-6 | -1.518e+01 | 0.0000 | 23.0 | 6.57% | 1807.0 | 1.85% | motif file (matrix) | svg |

| | | | | | | | | | | | |
|---|---|---|---|---|---|---|---|---|---|---|---|
| | 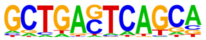 | | | | | | | | | | |
| 33 | 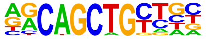 | MyoD(bHLH)/Myotube-MyoD-ChIP-Seq(GSE21614)/Homer | 1e-6 | -1.512e+01 | 0.0000 | 52.0 | 14.86% | 6823.7 | 7.00% | motif file (matrix) | svg |
| 34 | 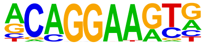 | ERG(ETS)/VCaP-ERG-ChIP-Seq(GSE14097)/Homer | 1e-6 | -1.497e+01 | 0.0000 | 89.0 | 25.43% | 14680.3 | 15.06% | motif file (matrix) | svg |
| 35 | 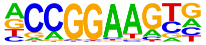 | ETV4(ETS)/HepG2-ETV4-ChIP-Seq(ENCODE)/Homer | 1e-6 | -1.495e+01 | 0.0000 | 62.0 | 17.71% | 8863.1 | 9.09% | motif file (matrix) | svg |
| 36 | 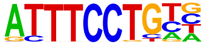 | EWS:ERG-fusion(ETS)/CADO_ES1-EWS:ERG-ChIP-Seq(SRA014231)/Homer | 1e-6 | -1.487e+01 | 0.0000 | 53.0 | 15.14% | 7074.0 | 7.25% | motif file (matrix) | svg |
| 37 | 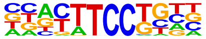 | Etv2(ETS)/ES-ER71-ChIP-Seq(GSE59402)/Homer | 1e-6 | -1.478e+01 | 0.0000 | 59.0 | 16.86% | 8295.8 | 8.51% | motif file (matrix) | svg |
| 38 | 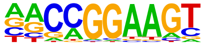 | GABPA(ETS)/Jurkat-GABPa-ChIP-Seq(GSE17954)/Homer | 1e-6 | -1.472e+01 | 0.0000 | 54.0 | 15.43% | 7307.4 | 7.49% | motif file (matrix) | svg |
| 39 | | NFE2L2(bZIP)/HepG2-NFE2L2-ChIP-Seq(Encode)/Homer | 1e-6 | -1.467e+01 | 0.0000 | 11.0 | 3.14% | 412.4 | 0.42% | motif file (matrix) | svg |

| | | | | | | | | | | | |
|---|---|---|---|---|---|---|---|---|---|---|---|
| | 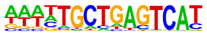 | | | | | | | | | | |
| 40 | 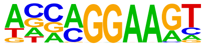 | EHF(ETS)/LoVo-EHF-ChIP-Seq(GSE49402)/Homer | 1e-6 | -1.422e+01 | 0.0000 | 76.0 | 21.71% | 12041.2 | 12.35% | motif file (matrix) | svg |
| 41 | 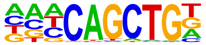 | Ap4(bHLH)/AML-Tfap4-ChIP-Seq(GSE45738)/Homer | 1e-6 | -1.414e+01 | 0.0000 | 74.0 | 21.14% | 11630.0 | 11.93% | motif file (matrix) | svg |
| 42 | 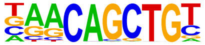 | Myf5(bHLH)/GM-Myf5-ChIP-Seq(GSE24852)/Homer | 1e-6 | -1.411e+01 | 0.0000 | 49.0 | 14.00% | 6468.4 | 6.63% | motif file (matrix) | svg |
| 43 | 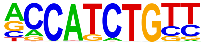 | NeuroG2(bHLH)/Fibroblast-NeuroG2-ChIP-Seq(GSE75910)/Homer | 1e-5 | -1.360e+01 | 0.0000 | 90.0 | 25.71% | 15380.3 | 15.77% | motif file (matrix) | svg |
| 44 | 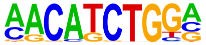 | ZBTB18(Zf)/HEK293-ZBTB18.GFP-ChIP-Seq(GSE58341)/Homer | 1e-5 | -1.352e+01 | 0.0000 | 40.0 | 11.43% | 4882.5 | 5.01% | motif file (matrix) | svg |
| 45 | 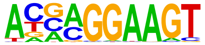 | ELF5(ETS)/T47D-ELF5-ChIP-Seq(GSE30407)/Homer | 1e-5 | -1.340e+01 | 0.0000 | 48.0 | 13.71% | 6437.3 | 6.60% | motif file (matrix) | svg |
| 46 | | NF-E2(bZIP)/K562-NFE2-ChIP-Seq(GSE31477)/Homer | 1e-5 | -1.320e+01 | 0.0000 | 11.0 | 3.14% | 480.1 | 0.49% | motif file (matrix) | svg |

| | | | | | | | | | | | |
|---|---|---|---|---|---|---|---|---|---|---|---|
| | 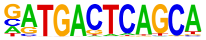 | | | | | | | | | | |
| 47 | 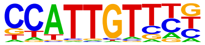 | Sox17(HMG)/Endoderm-Sox17-ChIP-Seq(GSE61475)/Homer | 1e-5 | -1.287e+01 | 0.0000 | 53.0 | 15.14% | 7567.4 | 7.76% | motif file (matrix) | svg |
| 48 | 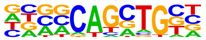 | Ascl1(bHLH)/NeuralTubes-Ascl1-ChIP-Seq(GSE55840)/Homer | 1e-5 | -1.285e+01 | 0.0000 | 88.0 | 25.14% | 15197.8 | 15.59% | motif file (matrix) | svg |
| 49 | 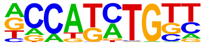 | Olig2(bHLH)/Neuron-Olig2-ChIP-Seq(GSE30882)/Homer | 1e-5 | -1.280e+01 | 0.0000 | 107.0 | 30.57% | 19680.3 | 20.18% | motif file (matrix) | svg |
| 50 | 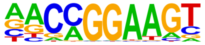 | ETV1(ETS)/GIST48-ETV1-ChIP-Seq(GSE22441)/Homer | 1e-5 | -1.261e+01 | 0.0000 | 72.0 | 20.57% | 11681.7 | 11.98% | motif file (matrix) | svg |
| 51 | 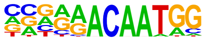 | Sox7(HMG)/ESC-Sox7-ChIP-Seq(GSE133899)/Homer | 1e-5 | -1.223e+01 | 0.0000 | 27.0 | 7.71% | 2809.1 | 2.88% | motif file (matrix) | svg |
| 52 | 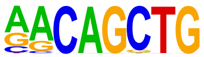 | MyoG(bHLH)/C2C12-MyoG-ChIP-Seq(GSE36024)/Homer | 1e-4 | -1.091e+01 | 0.0002 | 61.0 | 17.43% | 9832.4 | 10.08% | motif file (matrix) | svg |
| 53 | | PU.1(ETS)/ThioMac-PU.1-ChIP-Seq(GSE21512)/Homer | 1e-4 | -1.061e+01 | 0.0002 | 34.0 | 9.71% | 4368.6 | 4.48% | motif file (matrix) | svg |

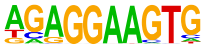

| | | | | | | | | | | | |
|---|---|---|---|---|---|---|---|---|---|---|---|
| 54 | 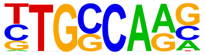 | NF1-halfsite(CTF)/LNCaP-NF1-ChIP-Seq(Unpublished)/Homer | 1e-4 | -1.048e+01 | 0.0002 | 93.0 | 26.57% | 17331.2 | 17.77% | motif file (matrix) | svg |
| 55 | 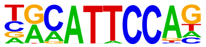 | TEAD3(TEA)/HepG2-TEAD3-ChIP-Seq(Encode)/Homer | 1e-4 | -9.786e+00 | 0.0005 | 70.0 | 20.00% | 12248.9 | 12.56% | motif file (matrix) | svg |
| 56 | 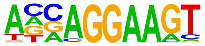 | ELF3(ETS)/PDAC-ELF3-ChIP-Seq(GSE64557)/Homer | 1e-4 | -9.705e+00 | 0.0005 | 44.0 | 12.57% | 6588.7 | 6.76% | motif file (matrix) | svg |
| 57 | 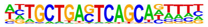 | MafB(bZIP)/BMM-Mafb-ChIP-Seq(GSE75722)/Homer | 1e-4 | -9.676e+00 | 0.0005 | 30.0 | 8.57% | 3814.0 | 3.91% | motif file (matrix) | svg |
| 58 | 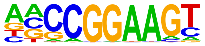 | ELF1(ETS)/Jurkat-ELF1-ChIP-Seq(SRA014231)/Homer | 1e-4 | -9.623e+00 | 0.0005 | 27.0 | 7.71% | 3268.8 | 3.35% | motif file (matrix) | svg |
| 59 | 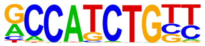 | NeuroD1(bHLH)/Islet-NeuroD1-ChIP-Seq(GSE30298)/Homer | 1e-4 | -9.598e+00 | 0.0005 | 50.0 | 14.29% | 7885.0 | 8.09% | motif file (matrix) | svg |
| 60 | | EWS:FLI1-fusion(ETS)/SK_N_MC-EWS:FLI1-ChIP-Seq(SRA014231)/Homer | 1e-4 | -9.462e+00 | 0.0006 | 35.0 | 10.00% | 4829.4 | 4.95% | motif file (matrix) | svg |

| | | | | | | | | | | | |
|---|---|---|---|---|---|---|---|---|---|---|---|
| | 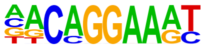 | | | | | | | | | | |
| 61 | 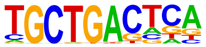 | MafA(bZIP)/Islet-MafA-ChIP-Seq(GSE30298)/Homer | 1e-3 | -8.750e+00 | 0.0012 | 44.0 | 12.57% | 6876.6 | 7.05% | motif file (matrix) | svg |
| 62 | 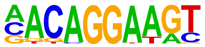 | Ets1-distal(ETS)/CD4+-PolII-ChIP-Seq(Barski_et_al.)/Homer | 1e-3 | -8.735e+00 | 0.0012 | 22.0 | 6.29% | 2536.4 | 2.60% | motif file (matrix) | svg |
| 63 | 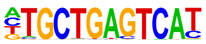 | Nrf2(bZIP)/Lymphoblast-Nrf2-ChIP-Seq(GSE37589)/Homer | 1e-3 | -8.702e+00 | 0.0012 | 8.0 | 2.29% | 420.5 | 0.43% | motif file (matrix) | svg |
| 64 | 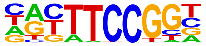 | Elk4(ETS)/Hela-Elk4-ChIP-Seq(GSE31477)/Homer | 1e-3 | -8.688e+00 | 0.0012 | 27.0 | 7.71% | 3463.4 | 3.55% | motif file (matrix) | svg |
| 65 | 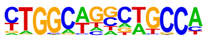 | Tlx?(NR)/NPC-H3K4me1-ChIP-Seq(GSE16256)/Homer | 1e-3 | -7.989e+00 | 0.0025 | 27.0 | 7.71% | 3622.6 | 3.72% | motif file (matrix) | svg |
| 66 | 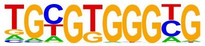 | Egr1(Zf)/K562-Egr1-ChIP-Seq(GSE32465)/Homer | 1e-3 | -7.571e+00 | 0.0037 | 29.0 | 8.29% | 4120.3 | 4.23% | motif file (matrix) | svg |
| 67 | | ETS(ETS)/Promoter/Homer | 1e-3 | -7.172e+00 | 0.0054 | 18.0 | 5.14% | 2113.0 | 2.17% | motif file (matrix) | svg |

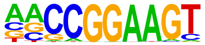

| | | | | | | | | | | | |
|---|---|---|---|---|---|---|---|---|---|---|---|
| 68 | 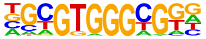 | Egr2(Zf)/Thymocytes-Egr2-ChIP-Seq(GSE34254)/Homer | 1e-3 | -6.997e+00 | 0.0063 | 10.0 | 2.86% | 822.9 | 0.84% | motif file (matrix) | svg |
| 69 | 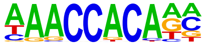 | RUNX1(Runt)/Jurkat-RUNX1-ChIP-Seq(GSE29180)/Homer | 1e-2 | -6.343e+00 | 0.0120 | 52.0 | 14.86% | 9563.4 | 9.81% | motif file (matrix) | svg |
| 70 | 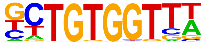 | RUNX-AML(Runt)/CD4+-PolII-ChIP-Seq(Barski_et_al.)/Homer | 1e-2 | -6.101e+00 | 0.0151 | 39.0 | 11.14% | 6713.8 | 6.89% | motif file (matrix) | svg |
| 71 | 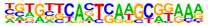 | PAX6(Paired,Homeobox)/Forebrain-Pax6-ChIP-Seq(GSE66961)/Homer | 1e-2 | -6.026e+00 | 0.0160 | 9.0 | 2.57% | 785.5 | 0.81% | motif file (matrix) | svg |
| 72 | 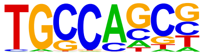 | HIC1(Zf)/Treg-ZBTB29-ChIP-Seq(GSE99889)/Homer | 1e-2 | -5.160e+00 | 0.0376 | 97.0 | 27.71% | 21309.0 | 21.85% | motif file (matrix) | svg |
| 73 | 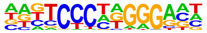 | EBF2(EBF)/BrownAdipose-EBF2-ChIP-Seq(GSE97114)/Homer | 1e-2 | -4.772e+00 | 0.0547 | 42.0 | 12.00% | 7979.3 | 8.18% | motif file (matrix) | svg |
| 74 | | EBF(EBF)/proBcell-EBF-ChIP-Seq(GSE21978)/Homer | 1e-2 | -4.650e+00 | 0.0610 | 12.0 | 3.43% | 1515.3 | 1.55% | motif file (matrix) | svg |

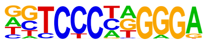

### Motifs found for: survival time

### Homer Known Motif Enrichment Results (homerpos12)

Homer *de novo* Motif Results
Gene Ontology Enrichment Results
Known Motif Enrichment Results (txt file)
Total Target Sequences = 298, Total Background Sequences = 97897

| Rank | Motif | Name | P-value | log P-pvalue | q-value (Benjamini) | # Target Sequences with Motif | % of Targets Sequences with Motif | # Background Sequences with Motif | % of Background Sequences with Motif | Motif File | SVG |
|---|---|---|---|---|---|---|---|---|---|---|---|
| 1 | 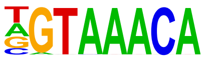 | Foxo3(Forkhead)/U2OS-Foxo3-ChIP-Seq(E-MTAB-2701)/Homer | 1e-7 | -1.716e+01 | 0.0000 | 53.0 | 17.79% | 7822.0 | 7.99% | motif file (matrix) | svg |
| 2 | 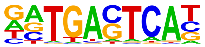 | JunB(bZIP)/DendriticCells-Junb-ChIP-Seq(GSE36099)/Homer | 1e-7 | -1.646e+01 | 0.0000 | 35.0 | 11.74% | 4149.2 | 4.24% | motif file (matrix) | svg |
| 3 | 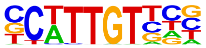 | Sox10(HMG)/SciaticNerve-Sox3-ChIP-Seq(GSE35132)/Homer | 1e-7 | -1.639e+01 | 0.0000 | 88.0 | 29.53% | 16706.1 | 17.06% | motif file (matrix) | svg |
| 4 | | Fra2(bZIP)/Striatum-Fra2-ChIP-Seq(GSE43429)/Homer | 1e-7 | -1.632e+01 | 0.0000 | 31.0 | 10.40% | 3405.9 | 3.48% | motif file (matrix) | svg |

| | | | | | | | | | | | |
|---|---|---|---|---|---|---|---|---|---|---|---|
| | 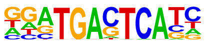 | | | | | | | | | | |
| 5 | 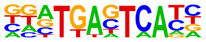 | Fos(bZIP)/TSC-Fos-ChIP-Seq(GSE110950)/ Homer | 1e-7 | -1.621e+01 | 0.0000 | 37.0 | 12.42% | 4593.9 | 4.69% | motif file (matrix) | svg |
| 6 | 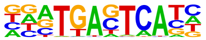 | Fra1(bZIP)/BT549-Fra1-ChIP-Seq(GSE46166)/ Homer | 1e-6 | -1.578e+01 | 0.0000 | 35.0 | 11.74% | 4271.2 | 4.36% | motif file (matrix) | svg |
| 7 | 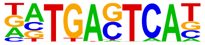 | BATF(bZIP)/Th17-BATF-ChIP-Seq(GSE39756)/ Homer | 1e-6 | -1.575e+01 | 0.0000 | 39.0 | 13.09% | 5092.7 | 5.20% | motif file (matrix) | svg |
| 8 | 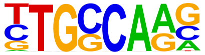 | NF1-halfsite(CTF)/LNCaP-NF1-ChIP-Seq(Unpublished)/Homer | 1e-6 | -1.488e+01 | 0.0000 | 84.0 | 28.19% | 16205.4 | 16.55% | motif file (matrix) | svg |
| 9 | 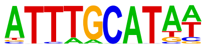 | Oct4(POU,Homeobox)/mES-Oct4-ChIP-Seq(GSE11431)/Homer | 1e-6 | -1.460e+01 | 0.0000 | 40.0 | 13.42% | 5546.7 | 5.66% | motif file (matrix) | svg |
| 10 | 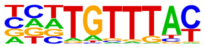 | FOXK1(Forkhead)/HEK293-FOXK1-ChIP-Seq(GSE51673)/Homer | 1e-6 | -1.422e+01 | 0.0000 | 63.0 | 21.14% | 11019.9 | 11.25% | motif file (matrix) | svg |
| 11 | | Foxf1(Forkhead)/Lung-Foxf1-ChIP-Seq(GSE77951)/Homer | 1e-6 | -1.404e+01 | 0.0000 | 61.0 | 20.47% | 10583.8 | 10.81% | motif file (matrix) | svg |

| | | | | | | | | | | |
|---|---|---|---|---|---|---|---|---|---|---|
| | 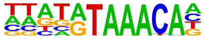 | | | | | | | | | |
| 12 | AP-1(bZIP)/ThioMac-PU.1-ChIP-Seq(GSE21512)/Homer 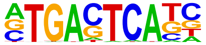 | 1e-5 | -1.365e+01 | 0.0000 | 40.0 | 13.42% | 5761.5 | 5.88% | motif file (matrix) | svg |
| 13 | Sox9(HMG)/Limb-SOX9-ChIP-Seq(GSE73225)/Homer 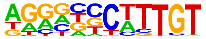 | 1e-5 | -1.335e+01 | 0.0001 | 47.0 | 15.77% | 7423.5 | 7.58% | motif file (matrix) | svg |
| 14 | Brn1(POU,Homeobox)/NPC-Brn1-ChIP-Seq(GSE35496)/Homer 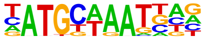 | 1e-5 | -1.321e+01 | 0.0001 | 29.0 | 9.73% | 3544.1 | 3.62% | motif file (matrix) | svg |
| 15 | Sox3(HMG)/NPC-Sox3-ChIP-Seq(GSE33059)/Homer 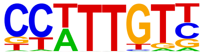 | 1e-5 | -1.212e+01 | 0.0002 | 88.0 | 29.53% | 18437.7 | 18.83% | motif file (matrix) | svg |
| 16 | Fosl2(bZIP)/3T3L1-Fosl2-ChIP-Seq(GSE56872)/Homer 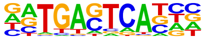 | 1e-5 | -1.166e+01 | 0.0003 | 20.0 | 6.71% | 2088.3 | 2.13% | motif file (matrix) | svg |
| 17 | Zic3(Zf)/mES-Zic3-ChIP-Seq(GSE37889)/Homer 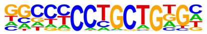 | 1e-4 | -1.135e+01 | 0.0003 | 32.0 | 10.74% | 4551.0 | 4.65% | motif file (matrix) | svg |
| 18 | Sox6(HMG)/Myotubes-Sox6-ChIP-Seq(GSE32627)/Homer | 1e-4 | -1.110e+01 | 0.0004 | 83.0 | 27.85% | 17520.4 | 17.89% | motif file (matrix) | svg |

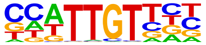

| | | | | | | | | | | | |
|---|---|---|---|---|---|---|---|---|---|---|---|
| 19 | 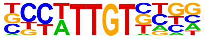 | Sox21(HMG)/ESC-SOX21-ChIP-Seq(GSE110505)/Homer | 1e-4 | -1.109e+01 | 0.0004 | 87.0 | 29.19% | 18639.7 | 19.04% | motif file (matrix) | svg |
| 20 | 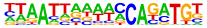 | Pitx1:Ebox(Homeobox,bHLH)/Hindlimb-Pitx1-ChIP-Seq(GSE41591)/Homer | 1e-4 | -1.092e+01 | 0.0004 | 19.0 | 6.38% | 2016.3 | 2.06% | motif file (matrix) | svg |
| 21 | 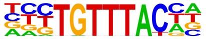 | FOXP1(Forkhead)/H9-FOXP1-ChIP-Seq(GSE31006)/Homer | 1e-4 | -1.077e+01 | 0.0005 | 30.0 | 10.07% | 4252.8 | 4.34% | motif file (matrix) | svg |
| 22 | 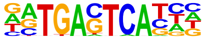 | Atf3(bZIP)/GBM-ATF3-ChIP-Seq(GSE33912)/Homer | 1e-4 | -1.046e+01 | 0.0006 | 34.0 | 11.41% | 5215.8 | 5.33% | motif file (matrix) | svg |
| 23 | 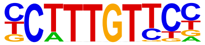 | Sox4(HMG)/proB-Sox4-ChIP-Seq(GSE50066)/Homer | 1e-4 | -9.877e+00 | 0.0011 | 46.0 | 15.44% | 8249.0 | 8.42% | motif file (matrix) | svg |
| 24 | 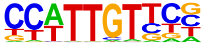 | SOX1(HMG)/NPC-SOX1-ChIP-Seq(GSE138215)/Homer | 1e-4 | -9.822e+00 | 0.0011 | 103.0 | 34.56% | 23900.2 | 24.41% | motif file (matrix) | svg |
| 25 | | Oct11(POU,Homeobox)/NCIH1048-POU2F3-ChIP-seq(GSE115123)/Homer | 1e-4 | -9.725e+00 | 0.0011 | 26.0 | 8.72% | 3630.5 | 3.71% | motif file (matrix) | svg |

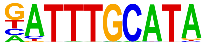

| | | | | | | | | | | | |
|---|---|---|---|---|---|---|---|---|---|---|---|
| 26 | 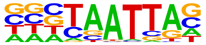 | DLX2(Homeobox)/BasalGanglia-Dlx2-ChIP-seq(GSE124936)/Homer | 1e-4 | -9.610e+00 | 0.0012 | 81.0 | 27.18% | 17687.4 | 18.06% | motif file (matrix) | svg |
| 27 | 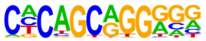 | Zic2(Zf)/ESC-Zic2-ChIP-Seq(SRP197560)/Homer | 1e-4 | -9.532e+00 | 0.0013 | 25.0 | 8.39% | 3460.7 | 3.53% | motif file (matrix) | svg |
| 28 | 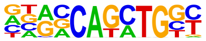 | Atoh1(bHLH)/Cerebellum-Atoh1-ChIP-Seq(GSE22111)/Homer | 1e-4 | -9.413e+00 | 0.0014 | 48.0 | 16.11% | 8916.6 | 9.11% | motif file (matrix) | svg |
| 29 | 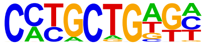 | Zic(Zf)/Cerebellum-ZIC1.2-ChIP-Seq(GSE60731)/Homer | 1e-4 | -9.374e+00 | 0.0014 | 38.0 | 12.75% | 6460.3 | 6.60% | motif file (matrix) | svg |
| 30 | 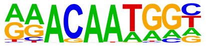 | Sox15(HMG)/CPA-Sox15-ChIP-Seq(GSE62909)/Homer | 1e-4 | -9.351e+00 | 0.0014 | 58.0 | 19.46% | 11529.3 | 11.78% | motif file (matrix) | svg |
| 31 | 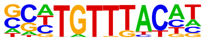 | FOXK2(Forkhead)/U2OS-FOXK2-ChIP-Seq(E-MTAB-2204)/Homer | 1e-3 | -9.101e+00 | 0.0017 | 37.0 | 12.42% | 6306.8 | 6.44% | motif file (matrix) | svg |
| 32 | | | 1e-3 | -9.099e+00 | 0.0017 | 18.0 | 6.04% | 2121.0 | 2.17% | motif file (matrix) | svg |

| | | | | | | | | | | | |
|---|---|---|---|---|---|---|---|---|---|---|---|
| | 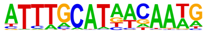 | OCT4-SOX2-TCF-NANOG(POU,Homeobox,HMG)/mES-Oct4-ChIP-Seq(GSE11431)/Homer | | | | | | | | | |
| 33 | 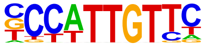 | Sox2(HMG)/mES-Sox2-ChIP-Seq(GSE11431)/Homer | 1e-3 | -9.019e+00 | 0.0017 | 47.0 | 15.77% | 8810.6 | 9.00% | motif file (matrix) | svg |
| 34 | 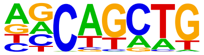 | SCL(bHLH)/HPC7-Scl-ChIP-Seq(GSE13511)/Homer | 1e-3 | -8.990e+00 | 0.0017 | 155.0 | 52.01% | 40451.7 | 41.31% | motif file (matrix) | svg |
| 35 | 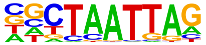 | LHX9(Homeobox)/Hct116-LHX9.V5-ChIP-Seq(GSE116822)/Homer | 1e-3 | -8.903e+00 | 0.0018 | 72.0 | 24.16% | 15532.0 | 15.86% | motif file (matrix) | svg |
| 36 | 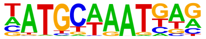 | Oct6(POU,Homeobox)/NPC-Pou3f1-ChIP-Seq(GSE35496)/Homer | 1e-3 | -8.860e+00 | 0.0019 | 30.0 | 10.07% | 4734.0 | 4.83% | motif file (matrix) | svg |
| 37 | 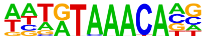 | FoxL2(Forkhead)/Ovary-FoxL2-ChIP-Seq(GSE60858)/Homer | 1e-3 | -8.823e+00 | 0.0019 | 50.0 | 16.78% | 9656.5 | 9.86% | motif file (matrix) | svg |
| 38 | 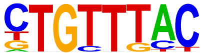 | Foxo1(Forkhead)/RAW-Foxo1-ChIP-Seq(Fan_et_al.)/Homer | 1e-3 | -8.806e+00 | 0.0019 | 85.0 | 28.52% | 19250.7 | 19.66% | motif file (matrix) | svg |
| 39 | | TEAD1(TEAD)/HepG2-TEAD1-ChIP-Seq(Encode)/Homer | 1e-3 | -8.730e+00 | 0.0020 | 51.0 | 17.11% | 9953.2 | 10.17% | motif file (matrix) | svg |

| | | | | | | | | | | | |
|---|---|---|---|---|---|---|---|---|---|---|---|
| | 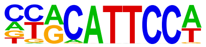 | | | | | | | | | | |
| 40 | 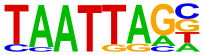 | Lhx2(Homeobox)/HFSC-Lhx2-ChIP-Seq(GSE48068)/Homer | 1e-3 | -8.665e+00 | 0.0020 | 59.0 | 19.80% | 12096.7 | 12.35% | motif file (matrix) | svg |
| 41 | 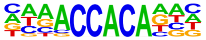 | RUNX2(Runt)/PCa-RUNX2-ChIP-Seq(GSE33889)/Homer | 1e-3 | -8.083e+00 | 0.0036 | 40.0 | 13.42% | 7392.9 | 7.55% | motif file (matrix) | svg |
| 42 | 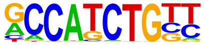 | NeuroD1(bHLH)/Islet-NeuroD1-ChIP-Seq(GSE30298)/Homer | 1e-3 | -8.039e+00 | 0.0036 | 36.0 | 12.08% | 6414.8 | 6.55% | motif file (matrix) | svg |
| 43 | 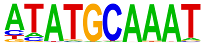 | Oct2(POU,Homeobox)/Bcell-Oct2-ChIP-Seq(GSE21512)/Homer | 1e-3 | -7.804e+00 | 0.0045 | 24.0 | 8.05% | 3656.4 | 3.73% | motif file (matrix) | svg |
| 44 | 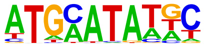 | Pit1(Homeobox)/GCrat-Pit1-ChIP-Seq(GSE58009)/Homer | 1e-3 | -7.733e+00 | 0.0047 | 55.0 | 18.46% | 11453.4 | 11.70% | motif file (matrix) | svg |
| 45 | 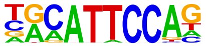 | TEAD3(TEA)/HepG2-TEAD3-ChIP-Seq(Encode)/Homer | 1e-3 | -7.624e+00 | 0.0051 | 56.0 | 18.79% | 11775.8 | 12.03% | motif file (matrix) | svg |
| 46 | | NeuroG2(bHLH)/Fibroblast-NeuroG2-ChIP-Seq(GSE75910)/Homer | 1e-3 | -7.611e+00 | 0.0051 | 61.0 | 20.47% | 13147.1 | 13.43% | motif file (matrix) | svg |

| | | | | | | | | | | | |
|---|---|---|---|---|---|---|---|---|---|---|---|
| | 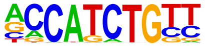 | | | | | | | | | | |
| 47 | 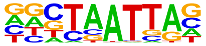 | En1(Homeobox)/SUM149-EN1-ChIP-Seq(GSE120957)/Homer | 1e-3 | -7.566e+00 | 0.0052 | 88.0 | 29.53% | 20852.8 | 21.30% | motif file (matrix) | svg |
| 48 | 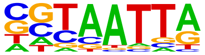 | DLX5(Homeobox)/BasalGanglia-Dlx5-ChIP-seq(GSE124936)/Homer | 1e-3 | -7.451e+00 | 0.0057 | 46.0 | 15.44% | 9187.9 | 9.38% | motif file (matrix) | svg |
| 49 | 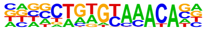 | Fox:Ebox(Forkhead,bHLH)/Panc1-Foxa2-ChIP-Seq(GSE47459)/Homer | 1e-3 | -7.445e+00 | 0.0057 | 45.0 | 15.10% | 8928.3 | 9.12% | motif file (matrix) | svg |
| 50 | 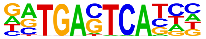 | Jun-AP1(bZIP)/K562-cJun-ChIP-Seq(GSE31477)/Homer | 1e-3 | -7.218e+00 | 0.0069 | 13.0 | 4.36% | 1482.9 | 1.51% | motif file (matrix) | svg |
| 51 | 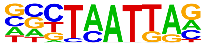 | Emx2(Homeobox)/Cortex-Emx2-ChIP-Seq(GSE183130)/Homer | 1e-3 | -7.165e+00 | 0.0072 | 58.0 | 19.46% | 12551.2 | 12.82% | motif file (matrix) | svg |
| 52 | 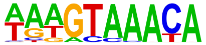 | FOXA1(Forkhead)/LNCAP-FOXA1-ChIP-Seq(GSE27824)/Homer | 1e-2 | -6.887e+00 | 0.0093 | 64.0 | 21.48% | 14370.6 | 14.68% | motif file (matrix) | svg |
| 53 | | Hoxd13(Homeobox)/ChickenMSG-Hoxd13.Flag-ChIP-Seq(GSE86088)/Homer | 1e-2 | -6.685e+00 | 0.0111 | 81.0 | 27.18% | 19371.0 | 19.78% | motif file (matrix) | svg |

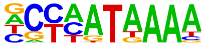

| | | | | | | | | | | |
|---|---|---|---|---|---|---|---|---|---|---|
| 54 | Foxa2(Forkhead)/Liver-Foxa2-ChIP-Seq(GSE25694)/Homer | 1e-2 | -6.670e+00 | 0.0111 | 40.0 | 13.42% | 7957.7 | 8.13% | motif file (matrix) | svg |

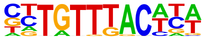

| | | | | | | | | | | |
|---|---|---|---|---|---|---|---|---|---|---|
| 55 | TCF4(bHLH)/SHSY5Y-TCF4-ChIP-Seq(GSE96915)/Homer | 1e-2 | -6.565e+00 | 0.0121 | 59.0 | 19.80% | 13151.7 | 13.43% | motif file (matrix) | svg |

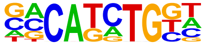

| | | | | | | | | | | |
|---|---|---|---|---|---|---|---|---|---|---|
| 56 | RUNX-AML(Runt)/CD4+-PolII-ChIP-Seq(Barski_et_al.)/Homer | 1e-2 | -6.474e+00 | 0.0130 | 33.0 | 11.07% | 6239.6 | 6.37% | motif file (matrix) | svg |

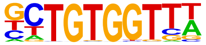

| | | | | | | | | | | |
|---|---|---|---|---|---|---|---|---|---|---|
| 57 | Sox7(HMG)/ESC-Sox7-ChIP-Seq(GSE133899)/Homer | 1e-2 | -6.322e+00 | 0.0149 | 18.0 | 6.04% | 2699.7 | 2.76% | motif file (matrix) | svg |

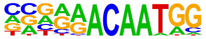

| | | | | | | | | | | |
|---|---|---|---|---|---|---|---|---|---|---|
| 58 | STAT6(Stat)/CD4-Stat6-ChIP-Seq(GSE22104)/Homer | 1e-2 | -6.245e+00 | 0.0158 | 28.0 | 9.40% | 5075.8 | 5.18% | motif file (matrix) | svg |

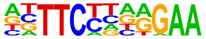

| | | | | | | | | | | |
|---|---|---|---|---|---|---|---|---|---|---|
| 59 | POU4F3(POU,Homeobox)/MEF-Pou4f3-ChIP-Seq(GSE150279)/Homer | 1e-2 | -6.168e+00 | 0.0168 | 14.0 | 4.70% | 1875.5 | 1.92% | motif file (matrix) | svg |

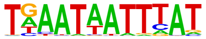

| | | | | | | | | | | |
|---|---|---|---|---|---|---|---|---|---|---|
| 60 | Twist2(bHLH)/Myoblast-Twist2.Ty1-ChIP-Seq(GSE127998)/Homer | 1e-2 | -6.150e+00 | 0.0168 | 66.0 | 22.15% | 15375.9 | 15.70% | motif file (matrix) | svg |

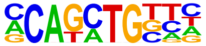

| | | | | | | | | | | | |
|---|---|---|---|---|---|---|---|---|---|---|---|
| 61 | 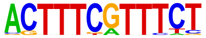 | T1ISRE(IRF)/ThioMac-Ifnb-Expression/Homer | 1e-2 | -6.049e+00 | 0.0183 | 3.0 | 1.01% | 85.9 | 0.09% | motif file (matrix) | svg |
| 62 | 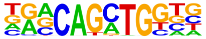 | Atoh7(bHLH)/Retina-Atoh7-CutnRun(GSE156756)/Homer | 1e-2 | -5.859e+00 | 0.0217 | 32.0 | 10.74% | 6231.4 | 6.36% | motif file (matrix) | svg |
| 63 | 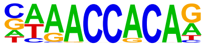 | RUNX(Runt)/HPC7-Runx1-ChIP-Seq(GSE22178)/Homer | 1e-2 | -5.662e+00 | 0.0260 | 31.0 | 10.40% | 6056.2 | 6.19% | motif file (matrix) | svg |
| 64 | 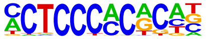 | WT1(Zf)/Kidney-WT1-ChIP-Seq(GSE90016)/Homer | 1e-2 | -5.500e+00 | 0.0302 | 25.0 | 8.39% | 4603.5 | 4.70% | motif file (matrix) | svg |
| 65 | 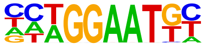 | TEAD(TEA)/Fibroblast-PU.1-ChIP-Seq(Unpublished)/Homer | 1e-2 | -5.475e+00 | 0.0304 | 36.0 | 12.08% | 7445.9 | 7.60% | motif file (matrix) | svg |
| 66 | 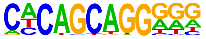 | Unknown-ESC-element(?)/mES-Nanog-ChIP-Seq(GSE11724)/Homer | 1e-2 | -5.429e+00 | 0.0314 | 24.0 | 8.05% | 4381.9 | 4.48% | motif file (matrix) | svg |
| 67 | | Gsx2(Homeobox)/LGE-Gsx2.Flag-ChIP-Seq(GSE162589)/Homer | 1e-2 | -5.402e+00 | 0.0318 | 64.0 | 21.48% | 15284.1 | 15.61% | motif file (matrix) | svg |

| | | | | | | | | | | |
|---|---|---|---|---|---|---|---|---|---|---|
| | 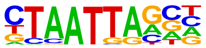 | | | | | | | | | |
| 68 | DLX1(Homeobox)/BasalGanglia-Dlx1-ChIP-seq(GSE124936)/Homer | 1e-2 | -5.296e+00 | 0.0348 | 65.0 | 21.81% | 15646.9 | 15.98% | motif file (matrix) | svg |
| | 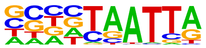 | | | | | | | | | |
| 69 | Rbpj1(?)/Panc1-Rbpj1-ChIP-Seq(GSE47459)/Homer | 1e-2 | -5.214e+00 | 0.0372 | 55.0 | 18.46% | 12827.3 | 13.10% | motif file (matrix) | svg |
| | 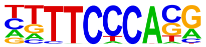 | | | | | | | | | |
| 70 | STAT6(Stat)/Macrophage-Stat6-ChIP-Seq(GSE38377)/Homer | 1e-2 | -5.133e+00 | 0.0398 | 26.0 | 8.72% | 4995.7 | 5.10% | motif file (matrix) | svg |
| | 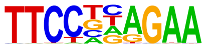 | | | | | | | | | |
| 71 | Olig2(bHLH)/Neuron-Olig2-ChIP-Seq(GSE30882)/Homer | 1e-2 | -5.127e+00 | 0.0398 | 71.0 | 23.83% | 17520.8 | 17.89% | motif file (matrix) | svg |
| | 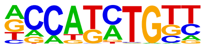 | | | | | | | | | |
| 72 | FOXA1(Forkhead)/MCF7-FOXA1-ChIP-Seq(GSE26831)/Homer | 1e-2 | -5.103e+00 | 0.0398 | 52.0 | 17.45% | 12046.4 | 12.30% | motif file (matrix) | svg |
| | 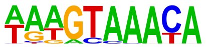 | | | | | | | | | |
| 73 | Hoxa11(Homeobox)/ChickenMSG-Hoxa11.Flag-ChIP-Seq(GSE86088)/Homer | 1e-2 | -5.096e+00 | 0.0398 | 105.0 | 35.23% | 27814.7 | 28.41% | motif file (matrix) | svg |
| | 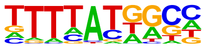 | | | | | | | | | |
| 74 | Foxa3(Forkhead)/Liver-Foxa3-ChIP-Seq(GSE77670)/Homer | 1e-2 | -5.015e+00 | 0.0424 | 18.0 | 6.04% | 3066.9 | 3.13% | motif file (matrix) | svg |

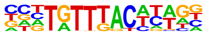

| | | | | | | | | | | | |
|---|---|---|---|---|---|---|---|---|---|---|---|
| 75 | 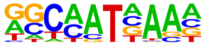 | Hoxa9(Homeobox)/ChickenMSG-Hoxa9.Flag-ChIP-Seq(GSE86088)/Homer | 1e-2 | -4.947e+00 | 0.0447 | 114.0 | 38.26% | 30746.6 | 31.40% | motif file (matrix) | svg |
| 76 | 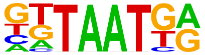 | Nkx6.1(Homeobox)/Islet-Nkx6.1-ChIP-Seq(GSE40975)/Homer | 1e-2 | -4.944e+00 | 0.0447 | 111.0 | 37.25% | 29813.0 | 30.45% | motif file (matrix) | svg |
| 77 | 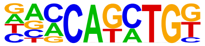 | BHLHA15(bHLH)/NIH3T3-BHLHB8.HA-ChIP-Seq(GSE119782)/Homer | 1e-2 | -4.856e+00 | 0.0477 | 51.0 | 17.11% | 11919.2 | 12.17% | motif file (matrix) | svg |
| 78 | 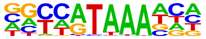 | CDX4(Homeobox)/ZebrafishEmbryos-Cdx4.Myc-ChIP-Seq(GSE48254)/Homer | 1e-2 | -4.826e+00 | 0.0485 | 45.0 | 15.10% | 10247.2 | 10.47% | motif file (matrix) | svg |
| 79 | 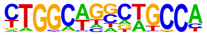 | Tlx?(NR)/NPC-H3K4me1-ChIP-Seq(GSE16256)/Homer | 1e-2 | -4.802e+00 | 0.0491 | 18.0 | 6.04% | 3134.1 | 3.20% | motif file (matrix) | svg |
| 80 | 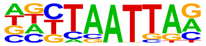 | Lhx1(Homeobox)/EmbryoCarcinoma-Lhx1-ChIP-Seq(GSE70957)/Homer | 1e-2 | -4.735e+00 | 0.0518 | 56.0 | 18.79% | 13431.0 | 13.72% | motif file (matrix) | svg |
| 81 | | NFE2L2(bZIP)/HepG2-NFE2L2-ChIP-Seq(Encode)/Homer | 1e-2 | -4.722e+00 | 0.0518 | 5.0 | 1.68% | 410.1 | 0.42% | motif file (matrix) | svg |

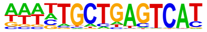

| | | | | | | | | | | | |
|---|---|---|---|---|---|---|---|---|---|---|---|
| 82 | 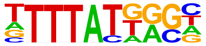 | Hoxc13(Homeobox)/EB-Hoxc13.HA-ChIP-Seq(GSE142377)/Homer | 1e-2 | -4.708e+00 | 0.0519 | 66.0 | 22.15% | 16362.0 | 16.71% | motif file (matrix) | svg |
| 83 | 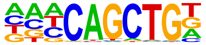 | Ap4(bHLH)/AML-Tfap4-ChIP-Seq(GSE45738)/Homer | 1e-2 | -4.644e+00 | 0.0547 | 44.0 | 14.77% | 10077.2 | 10.29% | motif file (matrix) | svg |

### Motifs found for: sex

### Homer Known Motif Enrichment Results (homerpos13)

[Homer *de novo* Motif Results]
[Gene Ontology Enrichment Results]
[Known Motif Enrichment Results (txt file)]
Total Target Sequences = 123, Total Background Sequences = 98117

| Rank | Motif | Name | P-value | log P-pvalue | q-value (Benjamini) | # Target Sequences with Motif | % of Targets Sequences with Motif | # Background Sequences with Motif | % of Background Sequences with Motif | Motif File | SVG |
|---|---|---|---|---|---|---|---|---|---|---|---|
| 1 | 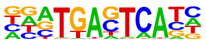 | Fra1(bZIP)/BT549-Fra1-ChIP-Seq(GSE46166)/Homer | 1e-15 | -3.517e+01 | 0.0000 | 32.0 | 26.02% | 4434.9 | 4.52% | motif file (matrix) | svg |
| 2 | 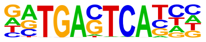 | Jun-AP1(bZIP)/K562-cJun-ChIP-Seq(GSE31477)/Homer | 1e-14 | -3.365e+01 | 0.0000 | 21.0 | 17.07% | 1644.1 | 1.68% | motif file (matrix) | svg |
| 3 | 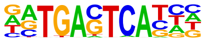 | Atf3(bZIP)/GBM-ATF3-ChIP-Seq(GSE33912)/Homer | 1e-13 | -3.188e+01 | 0.0000 | 33.0 | 26.83% | 5348.2 | 5.45% | motif file (matrix) | svg |
| 4 | | Fra2(bZIP)/Striatum-Fra2-ChIP-Seq(GSE43429)/Homer | 1e-13 | -3.182e+01 | 0.0000 | 28.0 | 22.76% | 3693.6 | 3.76% | motif file (matrix) | svg |

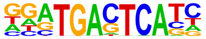

| | | | | | | | | | | | |
|---|---|---|---|---|---|---|---|---|---|---|---|
| 5 | 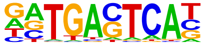 | JunB(bZIP)/DendriticCells-Junb-ChIP-Seq(GSE36099)/Homer | 1e-13 | -3.138e+01 | 0.0000 | 30.0 | 24.39% | 4404.9 | 4.49% | motif file (matrix) | svg |
| 6 | 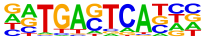 | Fosl2(bZIP)/3T3L1-Fosl2-ChIP-Seq(GSE56872)/Homer | 1e-13 | -3.127e+01 | 0.0000 | 23.0 | 18.70% | 2350.2 | 2.40% | motif file (matrix) | svg |
| 7 | 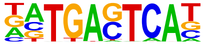 | BATF(bZIP)/Th17-BATF-ChIP-Seq(GSE39756)/Homer | 1e-13 | -3.030e+01 | 0.0000 | 32.0 | 26.02% | 5295.9 | 5.40% | motif file (matrix) | svg |
| 8 | 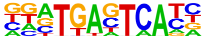 | Fos(bZIP)/TSC-Fos-ChIP-Seq(GSE110950)/Homer | 1e-12 | -2.977e+01 | 0.0000 | 30.0 | 24.39% | 4690.8 | 4.78% | motif file (matrix) | svg |
| 9 | 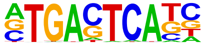 | AP-1(bZIP)/ThioMac-PU.1-ChIP-Seq(GSE21512)/Homer | 1e-11 | -2.655e+01 | 0.0000 | 32.0 | 26.02% | 6095.3 | 6.21% | motif file (matrix) | svg |
| 10 | 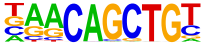 | Myf5(bHLH)/GM-Myf5-ChIP-Seq(GSE24852)/Homer | 1e-4 | -1.025e+01 | 0.0017 | 20.0 | 16.26% | 5783.9 | 5.89% | motif file (matrix) | svg |
| 11 | | Bach2(bZIP)/OCILy7-Bach2-ChIP-Seq(GSE44420)/Homer | 1e-4 | -9.850e+00 | 0.0023 | 9.0 | 7.32% | 1337.9 | 1.36% | motif file (matrix) | svg |

| | | | | | | | | | | | |
|---|---|---|---|---|---|---|---|---|---|---|---|
| | 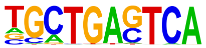 | | | | | | | | | | |
| 12 | 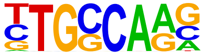 | NF1-halfsite(CTF)/LNCaP-NF1-ChIP-Seq(Unpublished)/Homer | 1e-4 | -9.739e+00 | 0.0023 | 39.0 | 31.71% | 16834.6 | 17.16% | motif file (matrix) | svg |
| 13 | 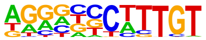 | Sox9(HMG)/Limb-SOX9-ChIP-Seq(GSE73225)/Homer | 1e-3 | -8.960e+00 | 0.0047 | 22.0 | 17.89% | 7398.7 | 7.54% | motif file (matrix) | svg |
| 14 | 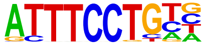 | EWS:ERG-fusion(ETS)/CADO_ES1-EWS:ERG-ChIP-Seq(SRA014231)/Homer | 1e-2 | -6.455e+00 | 0.0530 | 18.0 | 14.63% | 6623.5 | 6.75% | motif file (matrix) | svg |
| 15 | 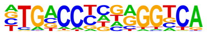 | FXR(NR),ER2/Liver-FXR-ChIP-Seq(GSE133700)/Homer | 1e-2 | -6.239e+00 | 0.0614 | 12.0 | 9.76% | 3596.7 | 3.67% | motif file (matrix) | svg |
| 16 | 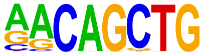 | MyoG(bHLH)/C2C12-MyoG-ChIP-Seq(GSE36024)/Homer | 1e-2 | -6.235e+00 | 0.0614 | 22.0 | 17.89% | 9059.7 | 9.23% | motif file (matrix) | svg |
| 17 | 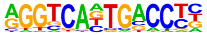 | FXR(NR),IR1/Liver-FXR-ChIP-Seq(Chong_et_al.)/Homer | 1e-2 | -6.174e+00 | 0.0614 | 11.0 | 8.94% | 3142.5 | 3.20% | motif file (matrix) | svg |
| 18 | | Etv2(ETS)/ES-ER71-ChIP-Seq(GSE59402)/Homer | 1e-2 | -6.107e+00 | 0.0614 | 20.0 | 16.26% | 7979.4 | 8.13% | motif file (matrix) | svg |

| | | | | | | | | | | |
|---|---|---|---|---|---|---|---|---|---|---|
| | 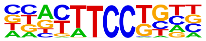 | | | | | | | | | |
| 19 | Sox10(HMG)/SciaticNerve-Sox3-ChIP-Seq(GSE35132)/Homer 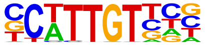 | 1e-2 | -6.036e+00 | 0.0614 | 33.0 | 26.83% | 16125.6 | 16.43% | motif file (matrix) | svg |
| 20 | Sox2(HMG)/mES-Sox2-ChIP-Seq(GSE11431)/Homer 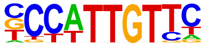 | 1e-2 | -5.791e+00 | 0.0721 | 20.0 | 16.26% | 8201.8 | 8.36% | motif file (matrix) | svg |
| 21 | ETS1(ETS)/Jurkat-ETS1-ChIP-Seq(GSE17954)/Homer 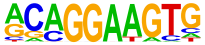 | 1e-2 | -5.651e+00 | 0.0790 | 21.0 | 17.07% | 8899.0 | 9.07% | motif file (matrix) | svg |
| 22 | Ap4(bHLH)/AML-Tfap4-ChIP-Seq(GSE45738)/Homer 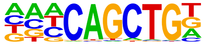 | 1e-2 | -5.625e+00 | 0.0790 | 24.0 | 19.51% | 10748.0 | 10.95% | motif file (matrix) | svg |
| 23 | ETV1(ETS)/GIST48-ETV1-ChIP-Seq(GSE22441)/Homer 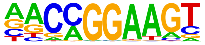 | 1e-2 | -5.286e+00 | 0.1038 | 25.0 | 20.33% | 11670.1 | 11.89% | motif file (matrix) | svg |
| 24 | ETV4(ETS)/HepG2-ETV4-ChIP-Seq(ENCODE)/Homer 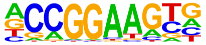 | 1e-2 | -5.224e+00 | 0.1059 | 21.0 | 17.07% | 9235.1 | 9.41% | motif file (matrix) | svg |
| 25 | E2F7(E2F)/Hela-E2F7-ChIP-Seq(GSE32673)/Homer | 1e-2 | -5.183e+00 | 0.1060 | 5.0 | 4.07% | 895.5 | 0.91% | motif file (matrix) | svg |

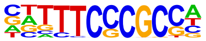

| | | | | | | | | | | | |
|---|---|---|---|---|---|---|---|---|---|---|---|
| 26 | 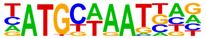 | Brn1(POU,Homeobox)/NPC-Brn1-ChIP-Seq(GSE35496)/Homer | 1e-2 | -5.012e+00 | 0.1209 | 10.0 | 8.13% | 3162.8 | 3.22% | motif file (matrix) | svg |
| 27 | 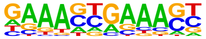 | IRF2(IRF)/Erythroblas-IRF2-ChIP-Seq(GSE36985)/Homer | 1e-2 | -4.917e+00 | 0.1280 | 5.0 | 4.07% | 955.6 | 0.97% | motif file (matrix) | svg |
| 28 | 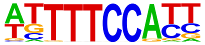 | NFAT(RHD)/Jurkat-NFATC1-ChIP-Seq(Jolma_et_al.)/Homer | 1e-2 | -4.902e+00 | 0.1280 | 20.0 | 16.26% | 8890.5 | 9.06% | motif file (matrix) | svg |
| 29 | 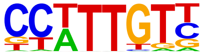 | Sox3(HMG)/NPC-Sox3-ChIP-Seq(GSE33059)/Homer | 1e-2 | -4.820e+00 | 0.1313 | 33.0 | 26.83% | 17424.2 | 17.76% | motif file (matrix) | svg |
| 30 | 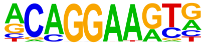 | ERG(ETS)/VCaP-ERG-ChIP-Seq(GSE14097)/Homer | 1e-2 | -4.765e+00 | 0.1341 | 28.0 | 22.76% | 14128.4 | 14.40% | motif file (matrix) | svg |
| 31 | 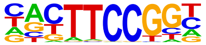 | Elk1(ETS)/Hela-Elk1-ChIP-Seq(GSE31477)/Homer | 1e-2 | -4.738e+00 | 0.1341 | 12.0 | 9.76% | 4354.4 | 4.44% | motif file (matrix) | svg |
| 32 | | Hoxc10(Homeobox)/EB-Hoxc10.iFlag-ChIP-Seq(GSE142377)/Homer | 1e-2 | -4.736e+00 | 0.1341 | 27.0 | 21.95% | 13500.8 | 13.76% | motif file (matrix) | svg |

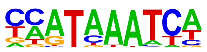

### Motifs found for: tumor initiation

### Homer Known Motif Enrichment Results (homerpos14)

Homer *de novo* Motif Results
Gene Ontology Enrichment Results
Known Motif Enrichment Results (txt file)
Total Target Sequences = 333, Total Background Sequences = 96850

| Rank | Motif | Name | P-value | log P-pvalue | q-value (Benjamini) | # Target Sequences with Motif | % of Targets Sequences with Motif | # Background Sequences with Motif | % of Background Sequences with Motif | Motif File | SVG |
|---|---|---|---|---|---|---|---|---|---|---|---|
| 1 | 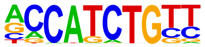 | NeuroG2(bHLH)/Fibroblast-NeuroG2-ChIP-Seq(GSE75910)/Homer | 1e-12 | -2.800e+01 | 0.0000 | 104.0 | 31.23% | 15069.8 | 15.56% | motif file (matrix) | svg |
| 2 | 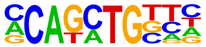 | Twist2(bHLH)/Myoblast-Twist2.Ty1-ChIP-Seq(GSE127998)/Homer | 1e-11 | -2.548e+01 | 0.0000 | 111.0 | 33.33% | 17288.2 | 17.85% | motif file (matrix) | svg |
| 3 | 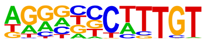 | Sox9(HMG)/Limb-SOX9-ChIP-Seq(GSE73225)/Homer | 1e-10 | -2.491e+01 | 0.0000 | 66.0 | 19.82% | 7882.8 | 8.14% | motif file (matrix) | svg |
| 4 | | Sox3(HMG)/NPC-Sox3-ChIP-Seq(GSE33059)/Homer | 1e-10 | -2.412e+01 | 0.0000 | 117.0 | 35.14% | 19069.9 | 19.69% | motif file (matrix) | svg |

| | | | | | | | | | | | |
|---|---|---|---|---|---|---|---|---|---|---|---|
| | 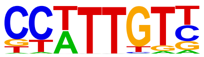 | | | | | | | | | | |
| 5 | | TCF4(bHLH)/SHSY5Y-TCF4-ChIP-Seq(GSE96915)/Homer | 1e-10 | -2.401e+01 | 0.0000 | 99.0 | 29.73% | 14960.8 | 15.45% | motif file (matrix) | svg |
| | 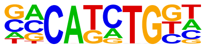 | | | | | | | | | | |
| 6 | | BHLHA15(bHLH)/NIH3T3-BHLHB8.HA-ChIP-Seq(GSE119782)/Homer | 1e-9 | -2.292e+01 | 0.0000 | 90.0 | 27.03% | 13252.7 | 13.68% | motif file (matrix) | svg |
| | 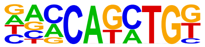 | | | | | | | | | | |
| 7 | | Olig2(bHLH)/Neuron-Olig2-ChIP-Seq(GSE30882)/Homer | 1e-9 | -2.255e+01 | 0.0000 | 117.0 | 35.14% | 19541.6 | 20.18% | motif file (matrix) | svg |
| | 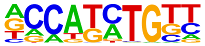 | | | | | | | | | | |
| 8 | | Sox10(HMG)/SciaticNerve-Sox3-ChIP-Seq(GSE35132)/Homer | 1e-9 | -2.205e+01 | 0.0000 | 108.0 | 32.43% | 17566.0 | 18.14% | motif file (matrix) | svg |
| | 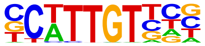 | | | | | | | | | | |
| 9 | | Atoh1(bHLH)/Cerebellum-Atoh1-ChIP-Seq(GSE22111)/Homer | 1e-9 | -2.094e+01 | 0.0000 | 72.0 | 21.62% | 9896.6 | 10.22% | motif file (matrix) | svg |
| | 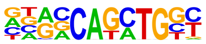 | | | | | | | | | | |
| 10 | | Atoh7(bHLH)/Retina-Atoh7-CutnRun(GSE156756)/Homer | 1e-8 | -1.875e+01 | 0.0000 | 54.0 | 16.22% | 6750.9 | 6.97% | motif file (matrix) | svg |
| | 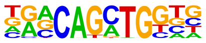 | | | | | | | | | | |
| 11 | | Tcf21(bHLH)/ArterySmoothMuscle-Tcf21-ChIP-Seq(GSE61369)/Homer | 1e-8 | -1.867e+01 | 0.0000 | 62.0 | 18.62% | 8358.9 | 8.63% | motif file (matrix) | svg |

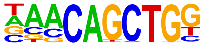

| | | | | | | | | | | | |
|---|---|---|---|---|---|---|---|---|---|---|---|
| 12 | 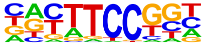 | Fli1(ETS)/CD8-FLI-ChIP-Seq(GSE20898)/Homer | 1e-8 | -1.852e+01 | 0.0000 | 64.0 | 19.22% | 8800.5 | 9.09% | motif file (matrix) | svg |
| 13 | 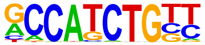 | NeuroD1(bHLH)/Islet-NeuroD1-ChIP-Seq(GSE30298)/Homer | 1e-8 | -1.846e+01 | 0.0000 | 56.0 | 16.82% | 7200.6 | 7.43% | motif file (matrix) | svg |
| 14 | 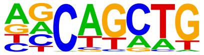 | SCL(bHLH)/HPC7-Scl-ChIP-Seq(GSE13511)/Homer | 1e-7 | -1.837e+01 | 0.0000 | 200.0 | 60.06% | 43196.5 | 44.60% | motif file (matrix) | svg |
| 15 | 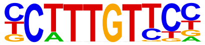 | Sox4(HMG)/proB-Sox4-ChIP-Seq(GSE50066)/Homer | 1e-7 | -1.817e+01 | 0.0000 | 63.0 | 18.92% | 8676.9 | 8.96% | motif file (matrix) | svg |
| 16 | 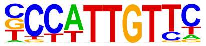 | Sox2(HMG)/mES-Sox2-ChIP-Seq(GSE11431)/Homer | 1e-7 | -1.798e+01 | 0.0000 | 65.0 | 19.52% | 9137.1 | 9.43% | motif file (matrix) | svg |
| 17 | 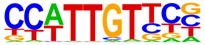 | SOX1(HMG)/NPC-SOX1-ChIP-Seq(GSE138215)/Homer | 1e-7 | -1.791e+01 | 0.0000 | 130.0 | 39.04% | 24347.1 | 25.14% | motif file (matrix) | svg |
| 18 | | Sox15(HMG)/CPA-Sox15-ChIP-Seq(GSE62909)/Homer | 1e-7 | -1.739e+01 | 0.0000 | 78.0 | 23.42% | 12093.7 | 12.49% | motif file (matrix) | svg |

| | | | | | | | | | | |
|---|---|---|---|---|---|---|---|---|---|---|
| | 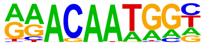 | | | | | | | | | |
| 19 | ETV4(ETS)/HepG2-ETV4-ChIP-Seq(ENCODE)/Homer 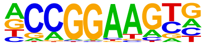 | 1e-7 | -1.714e+01 | 0.0000 | 60.0 | 18.02% | 8301.3 | 8.57% | motif file (matrix) | svg |
| 20 | Sox21(HMG)/ESC-SOX21-ChIP-Seq(GSE110505)/Homer 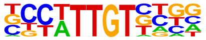 | 1e-7 | -1.672e+01 | 0.0000 | 108.0 | 32.43% | 19313.5 | 19.94% | motif file (matrix) | svg |
| 21 | Elk4(ETS)/Hela-Elk4-ChIP-Seq(GSE31477)/Homer 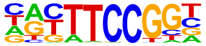 | 1e-6 | -1.600e+01 | 0.0000 | 33.0 | 9.91% | 3384.6 | 3.49% | motif file (matrix) | svg |
| 22 | Rfx2(HTH)/LoVo-RFX2-ChIP-Seq(GSE49402)/Homer 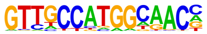 | 1e-6 | -1.521e+01 | 0.0000 | 12.0 | 3.60% | 501.0 | 0.52% | motif file (matrix) | svg |
| 23 | RFX(HTH)/K562-RFX3-ChIP-Seq(SRA012198)/Homer 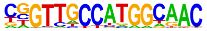 | 1e-6 | -1.461e+01 | 0.0000 | 11.0 | 3.30% | 433.5 | 0.45% | motif file (matrix) | svg |
| 24 | ERG(ETS)/VCaP-ERG-ChIP-Seq(GSE14097)/Homer 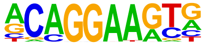 | 1e-6 | -1.452e+01 | 0.0000 | 83.0 | 24.92% | 14124.3 | 14.58% | motif file (matrix) | svg |
| 25 | X-box(HTH)/NPC-H3K4me1-ChIP-Seq(GSE16256)/Homer | 1e-5 | -1.366e+01 | 0.0000 | 14.0 | 4.20% | 807.5 | 0.83% | motif file (matrix) | svg |

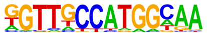

| | | | | | | | | | | | |
|---|---|---|---|---|---|---|---|---|---|---|---|
| 26 | 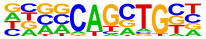 | Ascl1(bHLH)/NeuralTubes-Ascl1-ChIP-Seq(GSE55840)/Homer | 1e-5 | -1.360e+01 | 0.0000 | 81.0 | 24.32% | 13974.1 | 14.43% | [motif file (matrix)] | [svg] |
| 27 | 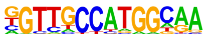 | Rfx1(HTH)/NPC-H3K4me1-ChIP-Seq(GSE16256)/Homer | 1e-5 | -1.338e+01 | 0.0000 | 19.0 | 5.71% | 1493.6 | 1.54% | [motif file (matrix)] | [svg] |
| 28 | 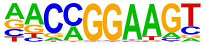 | ETV1(ETS)/GIST48-ETV1-ChIP-Seq(GSE22441)/Homer | 1e-5 | -1.278e+01 | 0.0000 | 68.0 | 20.42% | 11253.7 | 11.62% | [motif file (matrix)] | [svg] |
| 29 | 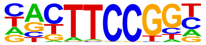 | Elk1(ETS)/Hela-Elk1-ChIP-Seq(GSE31477)/Homer | 1e-5 | -1.272e+01 | 0.0000 | 30.0 | 9.01% | 3387.4 | 3.50% | [motif file (matrix)] | [svg] |
| 30 | 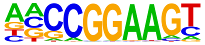 | ELF1(ETS)/Jurkat-ELF1-ChIP-Seq(SRA014231)/Homer | 1e-5 | -1.195e+01 | 0.0001 | 28.0 | 8.41% | 3161.1 | 3.26% | [motif file (matrix)] | [svg] |
| 31 | 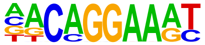 | EWS:FLI1-fusion(ETS)/SK_N_MC-EWS:FLI1-ChIP-Seq(SRA014231)/Homer | 1e-5 | -1.160e+01 | 0.0001 | 36.0 | 10.81% | 4737.9 | 4.89% | [motif file (matrix)] | [svg] |
| 32 | | ZBTB18(Zf)/HEK293-ZBTB18.GFP-ChIP-Seq(GSE58341)/Homer | 1e-4 | -1.092e+01 | 0.0003 | 35.0 | 10.51% | 4693.9 | 4.85% | [motif file (matrix)] | [svg] |

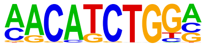

| | | | | | | | | | | | |
|---|---|---|---|---|---|---|---|---|---|---|---|
| 33 | 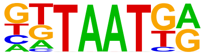 | Nkx6.1(Homeobox)/Islet-Nkx6.1-ChIP-Seq(GSE40975)/Homer | 1e-4 | -9.725e+00 | 0.0009 | 138.0 | 41.44% | 30308.5 | 31.29% | motif file (matrix) | svg |
| 34 | 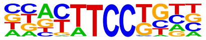 | Etv2(ETS)/ES-ER71-ChIP-Seq(GSE59402)/Homer | 1e-4 | -9.508e+00 | 0.0010 | 49.0 | 14.71% | 8052.5 | 8.31% | motif file (matrix) | svg |
| 35 | 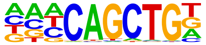 | Ap4(bHLH)/AML-Tfap4-ChIP-Seq(GSE45738)/Homer | 1e-4 | -9.469e+00 | 0.0010 | 61.0 | 18.32% | 10812.9 | 11.16% | motif file (matrix) | svg |
| 36 | 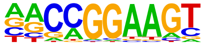 | GABPA(ETS)/Jurkat-GABPa-ChIP-Seq(GSE17954)/Homer | 1e-4 | -9.402e+00 | 0.0011 | 44.0 | 13.21% | 6981.2 | 7.21% | motif file (matrix) | svg |
| 37 | 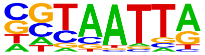 | DLX5(Homeobox)/BasalGanglia-Dlx5-ChIP-seq(GSE124936)/Homer | 1e-3 | -8.418e+00 | 0.0028 | 51.0 | 15.32% | 8890.3 | 9.18% | motif file (matrix) | svg |
| 38 | 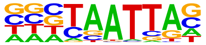 | DLX2(Homeobox)/BasalGanglia-Dlx2-ChIP-seq(GSE124936)/Homer | 1e-3 | -8.412e+00 | 0.0028 | 87.0 | 26.13% | 17643.7 | 18.22% | motif file (matrix) | svg |
| 39 | | Ets1-distal(ETS)/CD4+-PolII-ChIP-Seq(Barski_et_al.)/Homer | 1e-3 | -8.309e+00 | 0.0030 | 21.0 | 6.31% | 2546.8 | 2.63% | motif file (matrix) | svg |

| | | | | | | | | | | | |
|---|---|---|---|---|---|---|---|---|---|---|---|
| | 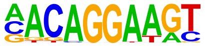 | | | | | | | | | | |
| 40 | 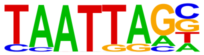 | Lhx2(Homeobox)/HFSC-Lhx2-ChIP-Seq(GSE48068)/Homer | 1e-3 | -8.079e+00 | 0.0037 | 63.0 | 18.92% | 11862.0 | 12.25% | motif file (matrix) | svg |
| 41 | 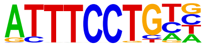 | EWS:ERG-fusion(ETS)/CADO_ES1-EWS:ERG-ChIP-Seq(SRA014231)/Homer | 1e-3 | -8.078e+00 | 0.0037 | 43.0 | 12.91% | 7192.8 | 7.43% | motif file (matrix) | svg |
| 42 | 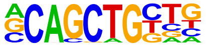 | Tcf12(bHLH)/GM12878-Tcf12-ChIP-Seq(GSE32465)/Homer | 1e-3 | -7.986e+00 | 0.0038 | 47.0 | 14.11% | 8132.9 | 8.40% | motif file (matrix) | svg |
| 43 | 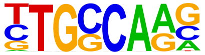 | NF1-halfsite(CTF)/LNCaP-NF1-ChIP-Seq(Unpublished)/Homer | 1e-3 | -7.958e+00 | 0.0038 | 81.0 | 24.32% | 16362.7 | 16.89% | motif file (matrix) | svg |
| 44 | 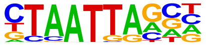 | Gsx2(Homeobox)/LGE-Gsx2.Flag-ChIP-Seq(GSE162589)/Homer | 1e-3 | -7.812e+00 | 0.0043 | 76.0 | 22.82% | 15182.1 | 15.68% | motif file (matrix) | svg |
| 45 | 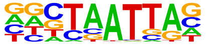 | En1(Homeobox)/SUM149-EN1-ChIP-Seq(GSE120957)/Homer | 1e-3 | -7.550e+00 | 0.0055 | 98.0 | 29.43% | 20953.9 | 21.64% | motif file (matrix) | svg |
| 46 | | ETS1(ETS)/Jurkat-ETS1-ChIP-Seq(GSE17954)/Homer | 1e-3 | -7.532e+00 | 0.0055 | 48.0 | 14.41% | 8536.6 | 8.81% | motif file (matrix) | svg |

| | | | | | | | | | | | |
|---|---|---|---|---|---|---|---|---|---|---|---|
| | 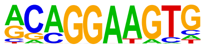 | | | | | | | | | | |
| 47 | 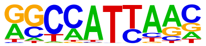 | Nanog(Homeobox)/mES-Nanog-ChIP-Seq(GSE11724)/Homer | 1e-3 | -7.389e+00 | 0.0062 | 184.0 | 55.26% | 44806.6 | 46.26% | motif file (matrix) | svg |
| 48 | 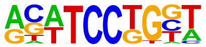 | SPDEF(ETS)/VCaP-SPDEF-ChIP-Seq(SRA014231)/Homer | 1e-3 | -7.375e+00 | 0.0062 | 48.0 | 14.41% | 8598.4 | 8.88% | motif file (matrix) | svg |
| 49 | 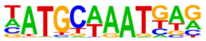 | Oct6(POU,Homeobox)/NPC-Pou3f1-ChIP-Seq(GSE35496)/Homer | 1e-3 | -7.185e+00 | 0.0073 | 31.0 | 9.31% | 4841.7 | 5.00% | motif file (matrix) | svg |
| 50 | 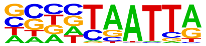 | DLX1(Homeobox)/BasalGanglia-Dlx1-ChIP-seq(GSE124936)/Homer | 1e-3 | -7.051e+00 | 0.0082 | 76.0 | 22.82% | 15582.2 | 16.09% | motif file (matrix) | svg |
| 51 | 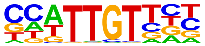 | Sox6(HMG)/Myotubes-Sox6-ChIP-Seq(GSE32627)/Homer | 1e-3 | -6.998e+00 | 0.0085 | 85.0 | 25.53% | 17908.3 | 18.49% | motif file (matrix) | svg |
| 52 | 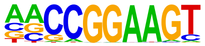 | ETS(ETS)/Promoter/Homer | 1e-3 | -6.950e+00 | 0.0087 | 17.0 | 5.11% | 2064.6 | 2.13% | motif file (matrix) | svg |
| 53 | | LHX9(Homeobox)/Hct116-LHX9.V5-ChIP-Seq(GSE116822)/Homer | 1e-3 | -6.914e+00 | 0.0088 | 76.0 | 22.82% | 15657.8 | 16.17% | motif file (matrix) | svg |

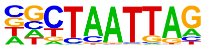

| | | | | | | | | | | | |
|---|---|---|---|---|---|---|---|---|---|---|---|
| 54 | 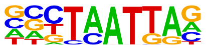 | Emx2(Homeobox)/Cortex-Emx2-ChIP-Seq(GSE183130)/Homer | 1e-2 | -6.847e+00 | 0.0093 | 64.0 | 19.22% | 12687.2 | 13.10% | motif file (matrix) | svg |
| 55 | 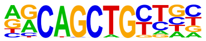 | MyoD(bHLH)/Myotube-MyoD-ChIP-Seq(GSE21614)/Homer | 1e-2 | -6.806e+00 | 0.0095 | 37.0 | 11.11% | 6289.5 | 6.49% | motif file (matrix) | svg |
| 56 | 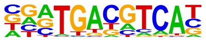 | Atf2(bZIP)/3T3L1-Atf2-ChIP-Seq(GSE56872)/Homer | 1e-2 | -6.601e+00 | 0.0115 | 21.0 | 6.31% | 2915.4 | 3.01% | motif file (matrix) | svg |
| 57 | 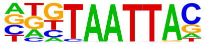 | Dlx3(Homeobox)/Kerainocytes-Dlx3-ChIP-Seq(GSE89884)/Homer | 1e-2 | -6.471e+00 | 0.0128 | 42.0 | 12.61% | 7564.7 | 7.81% | motif file (matrix) | svg |
| 58 | 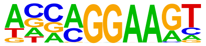 | EHF(ETS)/LoVo-EHF-ChIP-Seq(GSE49402)/Homer | 1e-2 | -6.333e+00 | 0.0145 | 60.0 | 18.02% | 11960.3 | 12.35% | motif file (matrix) | svg |
| 59 | 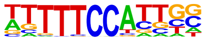 | NFATC2(RHD)/Islets-NFATC2-ChIP-Seq(GSE158496)/Homer | 1e-2 | -6.080e+00 | 0.0183 | 97.0 | 29.13% | 21625.3 | 22.33% | motif file (matrix) | svg |
| 60 | | NF1(CTF)/LNCAP-NF1-ChIP-Seq(Unpublished)/Homer | 1e-2 | -6.050e+00 | 0.0186 | 21.0 | 6.31% | 3054.0 | 3.15% | motif file (matrix) | svg |

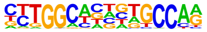

| | | | | | | | | | | | |
|---|---|---|---|---|---|---|---|---|---|---|---|
| 61 | 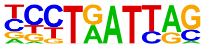 | Lhx6/Neurons-Lhx6-ChIP-seq(GSE85704)/Homer | 1e-2 | -6.033e+00 | 0.0186 | 65.0 | 19.52% | 13369.8 | 13.80% | motif file (matrix) | svg |
| 62 | 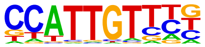 | Sox17(HMG)/Endoderm-Sox17-ChIP-Seq(GSE61475)/Homer | 1e-2 | -5.921e+00 | 0.0204 | 42.0 | 12.61% | 7795.2 | 8.05% | motif file (matrix) | svg |
| 63 | 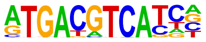 | JunD(bZIP)/K562-JunD-ChIP-Seq/Homer | 1e-2 | -5.739e+00 | 0.0241 | 7.0 | 2.10% | 549.7 | 0.57% | motif file (matrix) | svg |
| 64 | 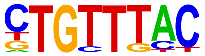 | Foxo1(Forkhead)/RAW-Foxo1-ChIP-Seq(Fan_et_al.)/Homer | 1e-2 | -5.626e+00 | 0.0266 | 91.0 | 27.33% | 20349.3 | 21.01% | motif file (matrix) | svg |
| 65 | 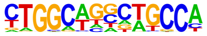 | Tlx?(NR)/NPC-H3K4me1-ChIP-Seq(GSE16256)/Homer | 1e-2 | -5.549e+00 | 0.0283 | 22.0 | 6.61% | 3403.5 | 3.51% | motif file (matrix) | svg |
| 66 | 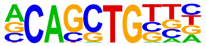 | Ptf1a(bHLH)/Panc1-Ptf1a-ChIP-Seq(GSE47459)/Homer | 1e-2 | -5.408e+00 | 0.0320 | 114.0 | 34.23% | 26696.2 | 27.56% | motif file (matrix) | svg |
| 67 | | PU.1(ETS)/ThioMac-PU.1-ChIP-Seq(GSE21512)/Homer | 1e-2 | -5.294e+00 | 0.0354 | 25.0 | 7.51% | 4135.4 | 4.27% | motif file (matrix) | svg |

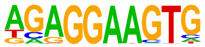

| | | | | | | | | | | | |
|---|---|---|---|---|---|---|---|---|---|---|---|
| 68 | 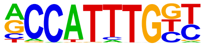 | Mesp1(bHLH)/ESC-Mesp1-ChIP-Seq(GSE165102)/Homer | 1e-2 | -5.279e+00 | 0.0354 | 30.0 | 9.01% | 5265.7 | 5.44% | motif file (matrix) | svg |
| 69 | 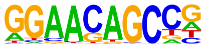 | ZNF341(Zf)/EBV-ZNF341-ChIP-Seq(GSE113194)/Homer | 1e-2 | -5.244e+00 | 0.0361 | 31.0 | 9.31% | 5508.5 | 5.69% | motif file (matrix) | svg |
| 70 | 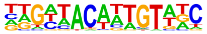 | DMRT6(DM)/Testis-DMRT6-ChIP-Seq(GSE60440)/Homer | 1e-2 | -5.183e+00 | 0.0378 | 14.0 | 4.20% | 1856.6 | 1.92% | motif file (matrix) | svg |
| 71 | 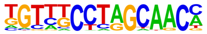 | Rfx6(HTH)/Min6b1-Rfx6.HA-ChIP-Seq(GSE62844)/Homer | 1e-2 | -5.107e+00 | 0.0403 | 43.0 | 12.91% | 8413.7 | 8.69% | motif file (matrix) | svg |
| 72 | 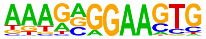 | SpiB(ETS)/OCILY3-SPIB-ChIP-Seq(GSE56857)/Homer | 1e-2 | -4.938e+00 | 0.0470 | 15.0 | 4.50% | 2112.4 | 2.18% | motif file (matrix) | svg |
| 73 | 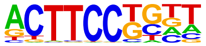 | Elf4(ETS)/BMDM-Elf4-ChIP-Seq(GSE88699)/Homer | 1e-2 | -4.717e+00 | 0.0578 | 42.0 | 12.61% | 8367.6 | 8.64% | motif file (matrix) | svg |
| 74 | | Myf5(bHLH)/GM-Myf5-ChIP-Seq(GSE24852)/Homer | 1e-2 | -4.671e+00 | 0.0597 | 32.0 | 9.61% | 5980.2 | 6.17% | motif file (matrix) | svg |

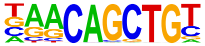

| | | | | | | | | | | |
|---|---|---|---|---|---|---|---|---|---|---|
| 75 | Rfx5(HTH)/GM12878-Rfx5-ChIP-Seq(GSE31477)/Homer | 1e-2 | -4.668e+00 | 0.0597 | 17.0 | 5.11% | 2597.3 | 2.68% | motif file (matrix) | svg |

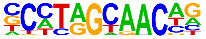

### Motifs found for: mouse survival hazard (Cox)

### Homer Known Motif Enrichment Results (homerpos15)

Homer *de novo* Motif Results
Gene Ontology Enrichment Results
Known Motif Enrichment Results (txt file)
Total Target Sequences = 462, Total Background Sequences = 95596

| Rank | Motif | Name | P-value | log P-pvalue | q-value (Benjamini) | # Target Sequences with Motif | % of Targets Sequences with Motif | # Background Sequences with Motif | % of Background Sequences with Motif | Motif File | SVG |
|---|---|---|---|---|---|---|---|---|---|---|---|
| 1 | 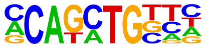 | Twist2(bHLH)/Myoblast-Twist2.Ty1-ChIP-Seq(GSE127998)/Homer | 1e-31 | -7.192e+01 | 0.0000 | 200.0 | 43.29% | 18469.2 | 19.32% | motif file (matrix) | svg |
| 2 | 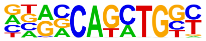 | Atoh1(bHLH)/Cerebellum-Atoh1-ChIP-Seq(GSE22111)/Homer | 1e-29 | -6.841e+01 | 0.0000 | 146.0 | 31.60% | 11063.5 | 11.57% | motif file (matrix) | svg |
| 3 | 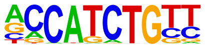 | NeuroG2(bHLH)/Fibroblast-NeuroG2-ChIP-Seq(GSE75910)/Homer | 1e-28 | -6.537e+01 | 0.0000 | 179.0 | 38.74% | 16120.7 | 16.86% | motif file (matrix) | svg |
| 4 | | BHLHA15(bHLH)/NIH3T3-BHLHB8.HA-ChIP-Seq(GSE119782)/Homer | 1e-28 | -6.508e+01 | 0.0000 | 167.0 | 36.15% | 14378.5 | 15.04% | motif file (matrix) | svg |

| | | | | | | | | | | | |
|---|---|---|---|---|---|---|---|---|---|---|---|
| | 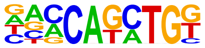 | | | | | | | | | | |
| 5 | 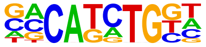 | TCF4(bHLH)/SHSY5Y-TCF4-ChIP-Seq(GSE96915)/Homer | 1e-27 | -6.330e+01 | 0.0000 | 177.0 | 38.31% | 16093.2 | 16.83% | motif file (matrix) | svg |
| 6 | 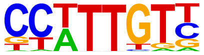 | Sox3(HMG)/NPC-Sox3-ChIP-Seq(GSE33059)/Homer | 1e-26 | -6.089e+01 | 0.0000 | 185.0 | 40.04% | 17658.9 | 18.47% | motif file (matrix) | svg |
| 7 | 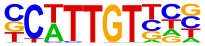 | Sox10(HMG)/SciaticNerve-Sox3-ChIP-Seq(GSE35132)/Homer | 1e-23 | -5.467e+01 | 0.0000 | 172.0 | 37.23% | 16543.4 | 17.30% | motif file (matrix) | svg |
| 8 | 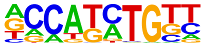 | Olig2(bHLH)/Neuron-Olig2-ChIP-Seq(GSE30882)/Homer | 1e-23 | -5.320e+01 | 0.0000 | 191.0 | 41.34% | 19796.5 | 20.71% | motif file (matrix) | svg |
| 9 | 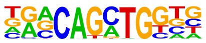 | Atoh7(bHLH)/Retina-Atoh7-CutnRun(GSE156756)/Homer | 1e-23 | -5.307e+01 | 0.0000 | 108.0 | 23.38% | 7727.0 | 8.08% | motif file (matrix) | svg |
| 10 | 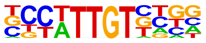 | Sox21(HMG)/ESC-SOX21-ChIP-Seq(GSE110505)/Homer | 1e-22 | -5.263e+01 | 0.0000 | 179.0 | 38.74% | 17953.5 | 18.78% | motif file (matrix) | svg |
| 11 | | NeuroD1(bHLH)/Islet-NeuroD1-ChIP-Seq(GSE30298)/Homer | 1e-22 | -5.259e+01 | 0.0000 | 110.0 | 23.81% | 8028.7 | 8.40% | motif file (matrix) | svg |

| | | | | | | | | | | | |
|---|---|---|---|---|---|---|---|---|---|---|---|
| | 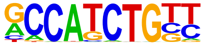 | | | | | | | | | | |
| 12 | | Tcf21(bHLH)/ArterySmoothMuscle-Tcf21-ChIP-Seq(GSE61369)/Homer | 1e-22 | -5.161e+01 | 0.0000 | 120.0 | 25.97% | 9434.5 | 9.87% | motif file (matrix) | svg |
| | 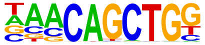 | | | | | | | | | | |
| 13 | | SOX1(HMG)/NPC-SOX1-ChIP-Seq(GSE138215)/Homer | 1e-22 | -5.094e+01 | 0.0000 | 206.0 | 44.59% | 22688.5 | 23.73% | motif file (matrix) | svg |
| | 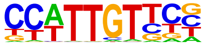 | | | | | | | | | | |
| 14 | | Sox2(HMG)/mES-Sox2-ChIP-Seq(GSE11431)/Homer | 1e-20 | -4.724e+01 | 0.0000 | 109.0 | 23.59% | 8474.2 | 8.86% | motif file (matrix) | svg |
| | 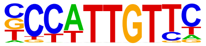 | | | | | | | | | | |
| 15 | | Sox9(HMG)/Limb-SOX9-ChIP-Seq(GSE73225)/Homer | 1e-20 | -4.715e+01 | 0.0000 | 104.0 | 22.51% | 7835.2 | 8.20% | motif file (matrix) | svg |
| | 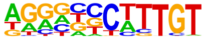 | | | | | | | | | | |
| 16 | | Sox4(HMG)/proB-Sox4-ChIP-Seq(GSE50066)/Homer | 1e-19 | -4.570e+01 | 0.0000 | 106.0 | 22.94% | 8255.8 | 8.64% | motif file (matrix) | svg |
| | 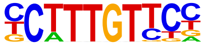 | | | | | | | | | | |
| 17 | | Ascl1(bHLH)/NeuralTubes-Ascl1-ChIP-Seq(GSE55840)/Homer | 1e-17 | -4.095e+01 | 0.0000 | 156.0 | 33.77% | 16216.3 | 16.96% | motif file (matrix) | svg |
| | 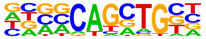 | | | | | | | | | | |
| 18 | | Sox6(HMG)/Myotubes-Sox6-ChIP-Seq(GSE32627)/Homer | 1e-16 | -3.795e+01 | 0.0000 | 154.0 | 33.33% | 16427.3 | 17.18% | motif file (matrix) | svg |

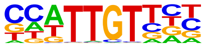

| | | | | | | | | | | | |
|---|---|---|---|---|---|---|---|---|---|---|---|
| 19 | 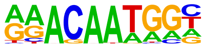 | Sox15(HMG)/CPA-Sox15-ChIP-Seq(GSE62909)/Homer | 1e-14 | -3.313e+01 | 0.0000 | 115.0 | 24.89% | 11217.7 | 11.73% | motif file (matrix) | svg |
| 20 | 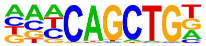 | Ap4(bHLH)/AML-Tfap4-ChIP-Seq(GSE45738)/Homer | 1e-14 | -3.312e+01 | 0.0000 | 121.0 | 26.19% | 12119.8 | 12.68% | motif file (matrix) | svg |
| 21 | 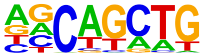 | SCL(bHLH)/HPC7-Scl-ChIP-Seq(GSE13511)/Homer | 1e-13 | -3.129e+01 | 0.0000 | 302.0 | 65.37% | 45753.9 | 47.86% | motif file (matrix) | svg |
| 22 | 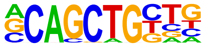 | Tcf12(bHLH)/GM12878-Tcf12-ChIP-Seq(GSE32465)/Homer | 1e-12 | -2.842e+01 | 0.0000 | 99.0 | 21.43% | 9612.1 | 10.05% | motif file (matrix) | svg |
| 23 | 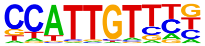 | Sox17(HMG)/Endoderm-Sox17-ChIP-Seq(GSE61475)/Homer | 1e-11 | -2.625e+01 | 0.0000 | 79.0 | 17.10% | 7085.1 | 7.41% | motif file (matrix) | svg |
| 24 | 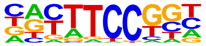 | Fli1(ETS)/CD8-FLI-ChIP-Seq(GSE20898)/Homer | 1e-11 | -2.557e+01 | 0.0000 | 92.0 | 19.91% | 9053.9 | 9.47% | motif file (matrix) | svg |
| 25 | | MyoD(bHLH)/Myotube-MyoD-ChIP-Seq(GSE21614)/Homer | 1e-9 | -2.274e+01 | 0.0000 | 79.0 | 17.10% | 7626.8 | 7.98% | motif file (matrix) | svg |

| | | | | | | | | | | | |
|---|---|---|---|---|---|---|---|---|---|---|---|
| | 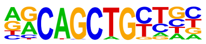 | | | | | | | | | | |
| 26 | 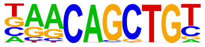 | Myf5(bHLH)/GM-Myf5-ChIP-Seq(GSE24852)/ Homer | 1e-9 | -2.198e+01 | 0.0000 | 72.0 | 15.58% | 6746.3 | 7.06% | motif file (matrix) | svg |
| 27 | 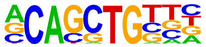 | Ptf1a(bHLH)/Panc1-Ptf1a-ChIP-Seq(GSE47459)/Homer | 1e-9 | -2.117e+01 | 0.0000 | 204.0 | 44.16% | 29276.9 | 30.62% | motif file (matrix) | svg |
| 28 | 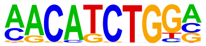 | ZBTB18(Zf)/HEK293-ZBTB18.GFP-ChIP-Seq(GSE58341)/Homer | 1e-9 | -2.077e+01 | 0.0000 | 59.0 | 12.77% | 5126.0 | 5.36% | motif file (matrix) | svg |
| 29 | 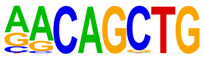 | MyoG(bHLH)/C2C12-MyoG-ChIP-Seq(GSE36024)/Homer | 1e-8 | -2.037e+01 | 0.0000 | 93.0 | 20.13% | 10162.6 | 10.63% | motif file (matrix) | svg |
| 30 | 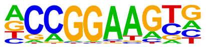 | ETV4(ETS)/HepG2-ETV4-ChIP-Seq(ENCODE)/Homer | 1e-7 | -1.823e+01 | 0.0000 | 82.0 | 17.75% | 8895.7 | 9.30% | motif file (matrix) | svg |
| 31 | 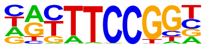 | Elk4(ETS)/Hela-Elk4-ChIP-Seq(GSE31477)/ Homer | 1e-7 | -1.722e+01 | 0.0000 | 44.0 | 9.52% | 3618.3 | 3.78% | motif file (matrix) | svg |
| 32 | | Sox7(HMG)/ESC-Sox7-ChIP-Seq(GSE133899)/ Homer | 1e-7 | -1.698e+01 | 0.0000 | 36.0 | 7.79% | 2642.5 | 2.76% | motif file (matrix) | svg |

| | | | | | | | | | | | |
|---|---|---|---|---|---|---|---|---|---|---|---|
| | 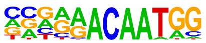 | | | | | | | | | | |
| 33 | 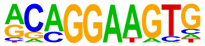 | ETS1(ETS)/Jurkat-ETS1-ChIP-Seq(GSE17954)/Homer | 1e-7 | -1.669e+01 | 0.0000 | 79.0 | 17.10% | 8748.2 | 9.15% | motif file (matrix) | svg |
| 34 | 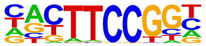 | Elk1(ETS)/Hela-Elk1-ChIP-Seq(GSE31477)/Homer | 1e-7 | -1.626e+01 | 0.0000 | 43.0 | 9.31% | 3612.9 | 3.78% | motif file (matrix) | svg |
| 35 | 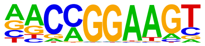 | ETV1(ETS)/GIST48-ETV1-ChIP-Seq(GSE22441)/Homer | 1e-6 | -1.498e+01 | 0.0000 | 95.0 | 20.56% | 11720.5 | 12.26% | motif file (matrix) | svg |
| 36 | 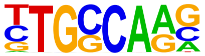 | NF1-halfsite(CTF)/LNCaP-NF1-ChIP-Seq(Unpublished)/Homer | 1e-6 | -1.414e+01 | 0.0000 | 130.0 | 28.14% | 17991.4 | 18.82% | motif file (matrix) | svg |
| 37 | 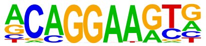 | ERG(ETS)/VCaP-ERG-ChIP-Seq(GSE14097)/Homer | 1e-5 | -1.231e+01 | 0.0001 | 107.0 | 23.16% | 14538.0 | 15.21% | motif file (matrix) | svg |
| 38 | 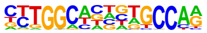 | NF1(CTF)/LNCAP-NF1-ChIP-Seq(Unpublished)/Homer | 1e-5 | -1.223e+01 | 0.0001 | 39.0 | 8.44% | 3658.1 | 3.83% | motif file (matrix) | svg |
| 39 | | | 1e-5 | -1.200e+01 | 0.0001 | 23.0 | 4.98% | 1617.9 | 1.69% | motif file (matrix) | svg |

| | | | | | | | | | | | |
|---|---|---|---|---|---|---|---|---|---|---|---|
| | 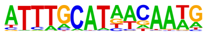 | OCT4-SOX2-TCF-NANOG(POU,Homeobox,HMG)/mES-Oct4-ChIP-Seq(GSE11431)/Homer | | | | | | | | | |
| 40 | 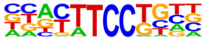 | Etv2(ETS)/ES-ER71-ChIP-Seq(GSE59402)/Homer | 1e-5 | -1.196e+01 | 0.0001 | 68.0 | 14.72% | 8092.8 | 8.46% | motif file (matrix) | svg |
| 41 | 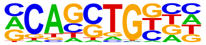 | HEB(bHLH)/mES-Heb-ChIP-Seq(GSE53233)/Homer | 1e-4 | -1.124e+01 | 0.0002 | 144.0 | 31.17% | 21592.2 | 22.58% | motif file (matrix) | svg |
| 42 | 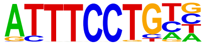 | EWS:ERG-fusion(ETS)/CADO_ES1-EWS:ERG-ChIP-Seq(SRA014231)/Homer | 1e-4 | -1.088e+01 | 0.0002 | 57.0 | 12.34% | 6606.3 | 6.91% | motif file (matrix) | svg |
| 43 | 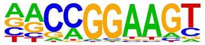 | GABPA(ETS)/Jurkat-GABPa-ChIP-Seq(GSE17954)/Homer | 1e-4 | -1.077e+01 | 0.0002 | 61.0 | 13.20% | 7266.9 | 7.60% | motif file (matrix) | svg |
| 44 | 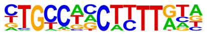 | ZNF7(Zf)/HepG2-ZNF7.Flag-ChIP-Seq(Encode)/Homer | 1e-4 | -1.060e+01 | 0.0003 | 48.0 | 10.39% | 5276.6 | 5.52% | motif file (matrix) | svg |
| 45 | 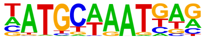 | Oct6(POU,Homeobox)/NPC-Pou3f1-ChIP-Seq(GSE35496)/Homer | 1e-4 | -1.014e+01 | 0.0004 | 39.0 | 8.44% | 4022.2 | 4.21% | motif file (matrix) | svg |
| 46 | | EWS:FLI1-fusion(ETS)/SK_N_MC-EWS:FLI1-ChIP-Seq(SRA014231)/Homer | 1e-4 | -1.007e+01 | 0.0004 | 44.0 | 9.52% | 4780.4 | 5.00% | motif file (matrix) | svg |

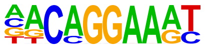

| | | | | | | | | | | |
|---|---|---|---|---|---|---|---|---|---|---|
| 47 | Brn1(POU,Homeobox)/NPC-Brn1-ChIP-Seq(GSE35496)/Homer 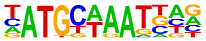 | 1e-4 | -9.881e+00 | 0.0005 | 31.0 | 6.71% | 2923.7 | 3.06% | motif file (matrix) | svg |
| 48 | Ets1-distal(ETS)/CD4+-PolII-ChIP-Seq(Barski_et_al.)/Homer 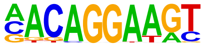 | 1e-3 | -9.109e+00 | 0.0011 | 26.0 | 5.63% | 2356.1 | 2.46% | motif file (matrix) | svg |
| 49 | ELF1(ETS)/Jurkat-ELF1-ChIP-Seq(SRA014231)/Homer 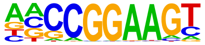 | 1e-3 | -8.938e+00 | 0.0013 | 34.0 | 7.36% | 3517.2 | 3.68% | motif file (matrix) | svg |
| 50 | Ascl2(bHLH)/ESC-Ascl2-ChIP-Seq(GSE97712)/Homer 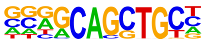 | 1e-3 | -8.917e+00 | 0.0013 | 90.0 | 19.48% | 12726.3 | 13.31% | motif file (matrix) | svg |
| 51 | Rfx1(HTH)/NPC-H3K4me1-ChIP-Seq(GSE16256)/Homer 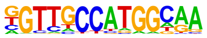 | 1e-3 | -8.133e+00 | 0.0027 | 19.0 | 4.11% | 1566.9 | 1.64% | motif file (matrix) | svg |
| 52 | Elf4(ETS)/BMDM-Elf4-ChIP-Seq(GSE88699)/Homer 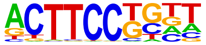 | 1e-3 | -8.117e+00 | 0.0027 | 64.0 | 13.85% | 8508.1 | 8.90% | motif file (matrix) | svg |
| 53 | SPDEF(ETS)/VCaP-SPDEF-ChIP-Seq(SRA014231)/Homer | 1e-3 | -7.840e+00 | 0.0035 | 65.0 | 14.07% | 8768.1 | 9.17% | motif file (matrix) | svg |

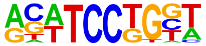

| | | | | | | | | | | | |
|---|---|---|---|---|---|---|---|---|---|---|---|
| 54 | 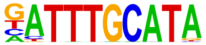 | Oct11(POU,Homeobox)/NCIH1048-POU2F3-ChIP-seq(GSE115123)/Homer | 1e-3 | -7.605e+00 | 0.0044 | 29.0 | 6.28% | 3039.5 | 3.18% | motif file (matrix) | svg |
| 55 | 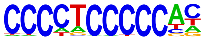 | ZNF148(Zf)/MDAMB231-ZNF148-ChIP-Seq(GSE147020)/Homer | 1e-3 | -7.552e+00 | 0.0045 | 36.0 | 7.79% | 4108.6 | 4.30% | motif file (matrix) | svg |
| 56 | 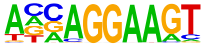 | ELF3(ETS)/PDAC-ELF3-ChIP-Seq(GSE64557)/Homer | 1e-3 | -7.390e+00 | 0.0052 | 50.0 | 10.82% | 6392.0 | 6.69% | motif file (matrix) | svg |
| 57 | 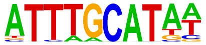 | Oct4(POU,Homeobox)/mES-Oct4-ChIP-Seq(GSE11431)/Homer | 1e-3 | -7.203e+00 | 0.0062 | 39.0 | 8.44% | 4662.5 | 4.88% | motif file (matrix) | svg |
| 58 | 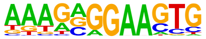 | SpiB(ETS)/OCILY3-SPIB-ChIP-Seq(GSE56857)/Homer | 1e-3 | -7.086e+00 | 0.0068 | 22.0 | 4.76% | 2124.6 | 2.22% | motif file (matrix) | svg |
| 59 | 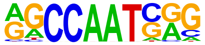 | NFY(CCAAT)/Promoter/Homer | 1e-2 | -6.895e+00 | 0.0081 | 53.0 | 11.47% | 7044.4 | 7.37% | motif file (matrix) | svg |
| 60 | | Nanog(Homeobox)/mES-Nanog-ChIP-Seq(GSE11724)/Homer | 1e-2 | -6.817e+00 | 0.0086 | 238.0 | 51.52% | 42357.0 | 44.30% | motif file (matrix) | svg |

| | | | | | | | | | | | |
|---|---|---|---|---|---|---|---|---|---|---|---|
| | 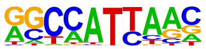 | | | | | | | | | | |
| 61 | 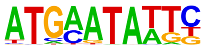 | Brn2(POU,Homeobox)/NPC-Brn2-ChIP-Seq(GSE35496)/Homer | 1e-2 | -6.779e+00 | 0.0088 | 12.0 | 2.60% | 856.4 | 0.90% | motif file (matrix) | svg |
| 62 | 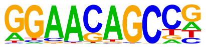 | ZNF341(Zf)/EBV-ZNF341-ChIP-Seq(GSE113194)/Homer | 1e-2 | -6.663e+00 | 0.0097 | 47.0 | 10.17% | 6113.9 | 6.39% | motif file (matrix) | svg |
| 63 | 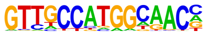 | Rfx2(HTH)/LoVo-RFX2-ChIP-Seq(GSE49402)/Homer | 1e-2 | -6.614e+00 | 0.0100 | 9.0 | 1.95% | 533.2 | 0.56% | motif file (matrix) | svg |
| 64 | 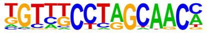 | Rfx6(HTH)/Min6b1-Rfx6.HA-ChIP-Seq(GSE62844)/Homer | 1e-2 | -6.595e+00 | 0.0101 | 65.0 | 14.07% | 9209.0 | 9.63% | motif file (matrix) | svg |
| 65 | 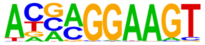 | ELF5(ETS)/T47D-ELF5-ChIP-Seq(GSE30407)/Homer | 1e-2 | -6.568e+00 | 0.0102 | 47.0 | 10.17% | 6142.8 | 6.43% | motif file (matrix) | svg |
| 66 | 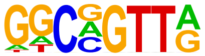 | MYB(HTH)/ERMYB-Myb-ChIPSeq(GSE22095)/Homer | 1e-2 | -6.481e+00 | 0.0110 | 97.0 | 21.00% | 15006.7 | 15.70% | motif file (matrix) | svg |
| 67 | | Foxo1(Forkhead)/RAW-Foxo1-ChIP-Seq(Fan_et_al.)/Homer | 1e-2 | -6.466e+00 | 0.0110 | 119.0 | 25.76% | 19113.3 | 19.99% | motif file (matrix) | svg |

| | | | | | | | | | | | |
|---|---|---|---|---|---|---|---|---|---|---|---|
| | 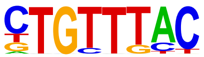 | | | | | | | | | | |
| 68 | 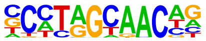 | Rfx5(HTH)/GM12878-Rfx5-ChIP-Seq(GSE31477)/Homer | 1e-2 | -6.299e+00 | 0.0128 | 25.0 | 5.41% | 2706.5 | 2.83% | motif file (matrix) | svg |
| 69 | 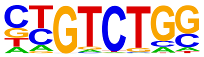 | Smad2(MAD)/ES-SMAD2-ChIP-Seq(GSE29422)/Homer | 1e-2 | -6.249e+00 | 0.0132 | 92.0 | 19.91% | 14198.3 | 14.85% | motif file (matrix) | svg |
| 70 | 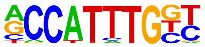 | Mesp1(bHLH)/ESC-Mesp1-ChIP-Seq(GSE165102)/Homer | 1e-2 | -5.887e+00 | 0.0187 | 40.0 | 8.66% | 5188.6 | 5.43% | motif file (matrix) | svg |
| 71 | 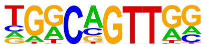 | AMYB(HTH)/Testes-AMYB-ChIP-Seq(GSE44588)/Homer | 1e-2 | -5.804e+00 | 0.0200 | 87.0 | 18.83% | 13496.0 | 14.12% | motif file (matrix) | svg |
| 72 | 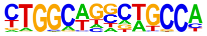 | Tlx?(NR)/NPC-H3K4me1-ChIP-Seq(GSE16256)/Homer | 1e-2 | -5.767e+00 | 0.0205 | 32.0 | 6.93% | 3918.4 | 4.10% | motif file (matrix) | svg |
| 73 | 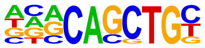 | E2A(bHLH)/proBcell-E2A-ChIP-Seq(GSE21978)/Homer | 1e-2 | -5.681e+00 | 0.0221 | 107.0 | 23.16% | 17279.0 | 18.07% | motif file (matrix) | svg |
| 74 | | EHF(ETS)/LoVo-EHF-ChIP-Seq(GSE49402)/Homer | 1e-2 | -5.670e+00 | 0.0221 | 77.0 | 16.67% | 11733.1 | 12.27% | motif file (matrix) | svg |

| | | | | | | | | | | | |
|---|---|---|---|---|---|---|---|---|---|---|---|
| | 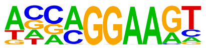 | | | | | | | | | | |
| 75 | 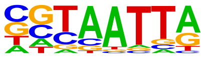 | DLX5(Homeobox)/BasalGanglia-Dlx5-ChIP-seq(GSE124936)/Homer | 1e-2 | -5.520e+00 | 0.0252 | 50.0 | 10.82% | 7004.5 | 7.33% | motif file (matrix) | svg |
| 76 | 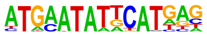 | OCT:OCT(POU,Homeobox)/NPC-Brn1-ChIP-Seq(GSE35496)/Homer | 1e-2 | -5.505e+00 | 0.0253 | 3.0 | 0.65% | 65.9 | 0.07% | motif file (matrix) | svg |
| 77 | 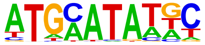 | Pit1(Homeobox)/GCrat-Pit1-ChIP-Seq(GSE58009)/Homer | 1e-2 | -5.301e+00 | 0.0306 | 66.0 | 14.29% | 9915.7 | 10.37% | motif file (matrix) | svg |
| 78 | 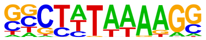 | TATA-Box(TBP)/Promoter/Homer | 1e-2 | -4.903e+00 | 0.0449 | 73.0 | 15.80% | 11373.1 | 11.90% | motif file (matrix) | svg |
| 79 | 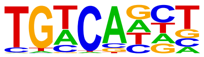 | Tgif2(Homeobox)/mES-Tgif2-ChIP-Seq(GSE55404)/Homer | 1e-2 | -4.783e+00 | 0.0500 | 187.0 | 40.48% | 33474.6 | 35.01% | motif file (matrix) | svg |

### Motifs found for: weibull alpha

### Homer Known Motif Enrichment Results (homerpos16)

Homer *de novo* Motif Results
Gene Ontology Enrichment Results
Known Motif Enrichment Results (txt file)
Total Target Sequences = 1536, Total Background Sequences = 96817

| Rank | Motif | Name | P-value | log P-pvalue | q-value (Benjamini) | # Target Sequences with Motif | % of Targets Sequences with Motif | # Background Sequences with Motif | % of Background Sequences with Motif | Motif File | SVG |
|---|---|---|---|---|---|---|---|---|---|---|---|
| 1 | 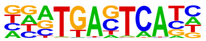 | Fra1(bZIP)/BT549-Fra1-ChIP-Seq(GSE46166)/Homer | 1e-498 | -1.147e+03 | 0.0000 | 752.0 | 48.96% | 5460.5 | 5.64% | motif file (matrix) | svg |
| 2 | 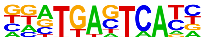 | Fos(bZIP)/TSC-Fos-ChIP-Seq(GSE110950)/Homer | 1e-494 | -1.139e+03 | 0.0000 | 771.0 | 50.20% | 5943.5 | 6.14% | motif file (matrix) | svg |
| 3 | 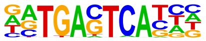 | Atf3(bZIP)/GBM-ATF3-ChIP-Seq(GSE33912)/Homer | 1e-479 | -1.105e+03 | 0.0000 | 787.0 | 51.24% | 6611.1 | 6.83% | motif file (matrix) | svg |
| 4 | | JunB(bZIP)/DendriticCells-Junb-ChIP-Seq(GSE36099)/Homer | 1e-472 | -1.087e+03 | 0.0000 | 725.0 | 47.20% | 5348.3 | 5.53% | motif file (matrix) | svg |

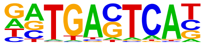

| | | | | | | | | | | | |
|---|---|---|---|---|---|---|---|---|---|---|---|
| 5 | 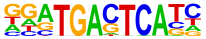 | Fra2(bZIP)/Striatum-Fra2-ChIP-Seq(GSE43429)/Homer | 1e-470 | -1.083e+03 | 0.0000 | 684.0 | 44.53% | 4535.9 | 4.69% | motif file (matrix) | svg |
| 6 | 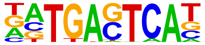 | BATF(bZIP)/Th17-BATF-ChIP-Seq(GSE39756)/Homer | 1e-466 | -1.073e+03 | 0.0000 | 772.0 | 50.26% | 6535.2 | 6.75% | motif file (matrix) | svg |
| 7 | 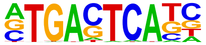 | AP-1(bZIP)/ThioMac-PU.1-ChIP-Seq(GSE21512)/Homer | 1e-448 | -1.033e+03 | 0.0000 | 791.0 | 51.50% | 7398.4 | 7.64% | motif file (matrix) | svg |
| 8 | 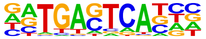 | Fosl2(bZIP)/3T3L1-Fosl2-ChIP-Seq(GSE56872)/Homer | 1e-387 | -8.917e+02 | 0.0000 | 519.0 | 33.79% | 2790.8 | 2.88% | motif file (matrix) | svg |
| 9 | 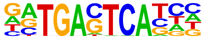 | Jun-AP1(bZIP)/K562-cJun-ChIP-Seq(GSE31477)/Homer | 1e-326 | -7.507e+02 | 0.0000 | 419.0 | 27.28% | 2009.7 | 2.08% | motif file (matrix) | svg |
| 10 | 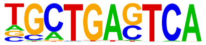 | Bach2(bZIP)/OCILy7-Bach2-ChIP-Seq(GSE44420)/Homer | 1e-153 | -3.546e+02 | 0.0000 | 243.0 | 15.82% | 1576.2 | 1.63% | motif file (matrix) | svg |
| 11 | | RUNX(Runt)/HPC7-Runx1-ChIP-Seq(GSE22178)/Homer | 1e-64 | -1.485e+02 | 0.0000 | 304.0 | 19.79% | 6420.7 | 6.63% | motif file (matrix) | svg |

| | | | | | | | | | | | |
|---|---|---|---|---|---|---|---|---|---|---|---|
| | 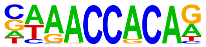 | | | | | | | | | | |
| 12 | 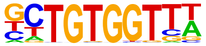 | RUNX-AML(Runt)/CD4+-PolII-ChIP-Seq(Barski_et_al.)/Homer | 1e-57 | -1.314e+02 | 0.0000 | 298.0 | 19.40% | 6728.9 | 6.95% | motif file (matrix) | svg |
| 13 | 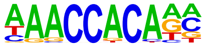 | RUNX1(Runt)/Jurkat-RUNX1-ChIP-Seq(GSE29180)/Homer | 1e-54 | -1.256e+02 | 0.0000 | 371.0 | 24.15% | 9905.8 | 10.23% | motif file (matrix) | svg |
| 14 | 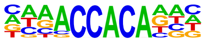 | RUNX2(Runt)/PCa-RUNX2-ChIP-Seq(GSE33889)/Homer | 1e-54 | -1.244e+02 | 0.0000 | 326.0 | 21.22% | 8076.5 | 8.34% | motif file (matrix) | svg |
| 15 | 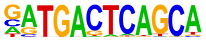 | NF-E2(bZIP)/K562-NFE2-ChIP-Seq(GSE31477)/Homer | 1e-42 | -9.775e+01 | 0.0000 | 70.0 | 4.56% | 475.0 | 0.49% | motif file (matrix) | svg |
| 16 | 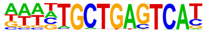 | Bach1(bZIP)/K562-Bach1-ChIP-Seq(GSE31477)/Homer | 1e-38 | -8.820e+01 | 0.0000 | 64.0 | 4.17% | 444.7 | 0.46% | motif file (matrix) | svg |
| 17 | 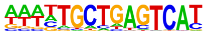 | NFE2L2(bZIP)/HepG2-NFE2L2-ChIP-Seq(Encode)/Homer | 1e-35 | -8.253e+01 | 0.0000 | 63.0 | 4.10% | 472.7 | 0.49% | motif file (matrix) | svg |
| 18 | | MafK(bZIP)/C2C12-MafK-ChIP-Seq(GSE36030)/Homer | 1e-30 | -7.035e+01 | 0.0000 | 107.0 | 6.97% | 1746.5 | 1.80% | motif file (matrix) | svg |

| Rank | Motif | Name | P-value | log P-value | q-value (Benjamini) | # Target Sequences with Motif | % of Targets Sequences with Motif | # Background Sequences with Motif | % of Background Sequences with Motif | Motif File | SVG |
|---|---|---|---|---|---|---|---|---|---|---|---|
| | 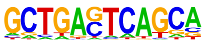 | | | | | | | | | | |
| 19 | 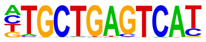 | Nrf2(bZIP)/Lymphoblast-Nrf2-ChIP-Seq(GSE37589)/Homer | 1e-28 | -6.663e+01 | 0.0000 | 51.0 | 3.32% | 386.9 | 0.40% | motif file (matrix) | svg |
| 20 | 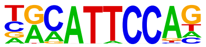 | TEAD3(TEA)/HepG2-TEAD3-ChIP-Seq(Encode)/Homer | 1e-28 | -6.585e+01 | 0.0000 | 360.0 | 23.44% | 12523.6 | 12.94% | motif file (matrix) | svg |
| 21 | 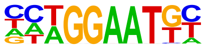 | TEAD(TEA)/Fibroblast-PU.1-ChIP-Seq(Unpublished)/Homer | 1e-26 | -6.189e+01 | 0.0000 | 253.0 | 16.47% | 7739.0 | 7.99% | motif file (matrix) | svg |
| 22 | 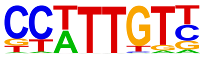 | Sox3(HMG)/NPC-Sox3-ChIP-Seq(GSE33059)/Homer | 1e-26 | -6.115e+01 | 0.0000 | 467.0 | 30.40% | 18328.5 | 18.93% | motif file (matrix) | svg |
| 23 | 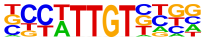 | Sox21(HMG)/ESC-SOX21-ChIP-Seq(GSE110505)/Homer | 1e-26 | -6.086e+01 | 0.0000 | 462.0 | 30.08% | 18088.4 | 18.69% | motif file (matrix) | svg |
| 24 | 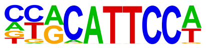 | TEAD1(TEAD)/HepG2-TEAD1-ChIP-Seq(Encode)/Homer | 1e-24 | -5.687e+01 | 0.0000 | 308.0 | 20.05% | 10604.5 | 10.96% | motif file (matrix) | svg |
| 25 | | TEAD4(TEA)/Tropoblast-Tead4-ChIP-Seq(GSE37350)/Homer | 1e-24 | -5.598e+01 | 0.0000 | 266.0 | 17.32% | 8673.6 | 8.96% | motif file (matrix) | svg |

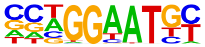

| | | | | | | | | | | | |
|---|---|---|---|---|---|---|---|---|---|---|---|
| 26 | 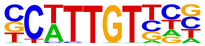 | Sox10(HMG)/SciaticNerve-Sox3-ChIP-Seq(GSE35132)/Homer | 1e-23 | -5.445e+01 | 0.0000 | 430.0 | 27.99% | 16949.0 | 17.51% | motif file (matrix) | svg |
| 27 | 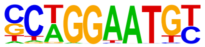 | TEAD2(TEA)/Py2T-Tead2-ChIP-Seq(GSE55709)/Homer | 1e-23 | -5.307e+01 | 0.0000 | 193.0 | 12.57% | 5563.5 | 5.75% | motif file (matrix) | svg |
| 28 | 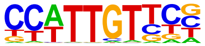 | SOX1(HMG)/NPC-SOX1-ChIP-Seq(GSE138215)/Homer | 1e-19 | -4.458e+01 | 0.0000 | 525.0 | 34.18% | 23074.8 | 23.84% | motif file (matrix) | svg |
| 29 | 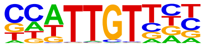 | Sox6(HMG)/Myotubes-Sox6-ChIP-Seq(GSE32627)/Homer | 1e-17 | -4.018e+01 | 0.0000 | 414.0 | 26.95% | 17436.3 | 18.01% | motif file (matrix) | svg |
| 30 | 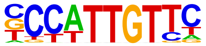 | Sox2(HMG)/mES-Sox2-ChIP-Seq(GSE11431)/Homer | 1e-14 | -3.454e+01 | 0.0000 | 234.0 | 15.23% | 8631.9 | 8.92% | motif file (matrix) | svg |
| 31 | 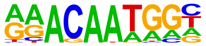 | Sox15(HMG)/CPA-Sox15-ChIP-Seq(GSE62909)/Homer | 1e-12 | -2.800e+01 | 0.0000 | 278.0 | 18.10% | 11466.2 | 11.85% | motif file (matrix) | svg |
| 32 | | MafA(bZIP)/Islet-MafA-ChIP-Seq(GSE30298)/Homer | 1e-11 | -2.719e+01 | 0.0000 | 177.0 | 11.52% | 6431.5 | 6.64% | motif file (matrix) | svg |

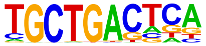

| | | | | | | | | | | | |
|---|---|---|---|---|---|---|---|---|---|---|---|
| 33 | 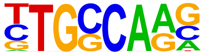 | NF1-halfsite(CTF)/LNCaP-NF1-ChIP-Seq(Unpublished)/Homer | 1e-10 | -2.459e+01 | 0.0000 | 351.0 | 22.85% | 15788.8 | 16.31% | motif file (matrix) | svg |
| 34 | 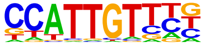 | Sox17(HMG)/Endoderm-Sox17-ChIP-Seq(GSE61475)/Homer | 1e-9 | -2.205e+01 | 0.0000 | 189.0 | 12.30% | 7473.7 | 7.72% | motif file (matrix) | svg |
| 35 | 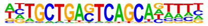 | MafB(bZIP)/BMM-Mafb-ChIP-Seq(GSE75722)/Homer | 1e-9 | -2.102e+01 | 0.0000 | 111.0 | 7.23% | 3759.4 | 3.88% | motif file (matrix) | svg |
| 36 | 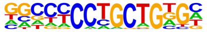 | Zic3(Zf)/mES-Zic3-ChIP-Seq(GSE37889)/Homer | 1e-7 | -1.741e+01 | 0.0000 | 104.0 | 6.77% | 3694.0 | 3.82% | motif file (matrix) | svg |
| 37 | 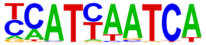 | Pdx1(Homeobox)/Islet-Pdx1-ChIP-Seq(SRA008281)/Homer | 1e-7 | -1.688e+01 | 0.0000 | 254.0 | 16.54% | 11501.5 | 11.88% | motif file (matrix) | svg |
| 38 | 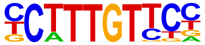 | Sox4(HMG)/proB-Sox4-ChIP-Seq(GSE50066)/Homer | 1e-7 | -1.638e+01 | 0.0000 | 190.0 | 12.37% | 8128.6 | 8.40% | motif file (matrix) | svg |
| 39 | | Brn1(POU,Homeobox)/NPC-Brn1-ChIP-Seq(GSE35496)/Homer | 1e-6 | -1.550e+01 | 0.0000 | 100.0 | 6.51% | 3652.2 | 3.77% | motif file (matrix) | svg |

| | | | | | | | | | | | |
|---|---|---|---|---|---|---|---|---|---|---|---|
| | 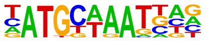 | | | | | | | | | | |
| 40 | 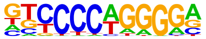 | EBF1(EBF)/Near-E2A-ChIP-Seq(GSE21512)/Homer | 1e-6 | -1.503e+01 | 0.0000 | 172.0 | 11.20% | 7341.2 | 7.58% | motif file (matrix) | svg |
| 41 | 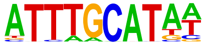 | Oct4(POU,Homeobox)/mES-Oct4-ChIP-Seq(GSE11431)/Homer | 1e-6 | -1.474e+01 | 0.0000 | 139.0 | 9.05% | 5663.0 | 5.85% | motif file (matrix) | svg |
| 42 | 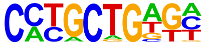 | Zic(Zf)/Cerebellum-ZIC1.2-ChIP-Seq(GSE60731)/Homer | 1e-6 | -1.417e+01 | 0.0000 | 149.0 | 9.70% | 6239.8 | 6.45% | motif file (matrix) | svg |
| 43 | 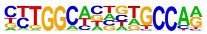 | NF1(CTF)/LNCAP-NF1-ChIP-Seq(Unpublished)/Homer | 1e-5 | -1.372e+01 | 0.0000 | 82.0 | 5.34% | 2934.2 | 3.03% | motif file (matrix) | svg |
| 44 | 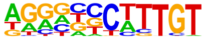 | Sox9(HMG)/Limb-SOX9-ChIP-Seq(GSE73225)/Homer | 1e-5 | -1.338e+01 | 0.0000 | 176.0 | 11.46% | 7758.4 | 8.01% | motif file (matrix) | svg |
| 45 | 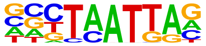 | Emx2(Homeobox)/Cortex-Emx2-ChIP-Seq(GSE183130)/Homer | 1e-5 | -1.329e+01 | 0.0000 | 279.0 | 18.16% | 13435.0 | 13.88% | motif file (matrix) | svg |
| 46 | | Ap4(bHLH)/AML-Tfap4-ChIP-Seq(GSE45738)/Homer | 1e-5 | -1.274e+01 | 0.0000 | 212.0 | 13.80% | 9797.0 | 10.12% | motif file (matrix) | svg |

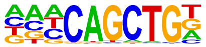

| | | | | | | | | | | | |
|---|---|---|---|---|---|---|---|---|---|---|---|
| 47 | 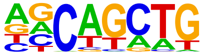 | SCL(bHLH)/HPC7-Scl-ChIP-Seq(GSE13511)/Homer | 1e-5 | -1.254e+01 | 0.0000 | 724.0 | 47.14% | 40104.8 | 41.43% | motif file (matrix) | svg |
| 48 | 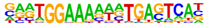 | NFAT:AP1(RHD,bZIP)/Jurkat-NFATC1-ChIP-Seq(Jolma_et_al.)/Homer | 1e-5 | -1.231e+01 | 0.0000 | 56.0 | 3.65% | 1829.6 | 1.89% | motif file (matrix) | svg |
| 49 | 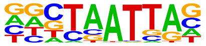 | En1(Homeobox)/SUM149-EN1-ChIP-Seq(GSE120957)/Homer | 1e-5 | -1.205e+01 | 0.0001 | 418.0 | 27.21% | 21697.1 | 22.41% | motif file (matrix) | svg |
| 50 | 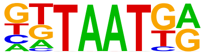 | Nkx6.1(Homeobox)/Islet-Nkx6.1-ChIP-Seq(GSE40975)/Homer | 1e-5 | -1.200e+01 | 0.0001 | 588.0 | 38.28% | 31888.5 | 32.94% | motif file (matrix) | svg |
| 51 | 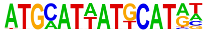 | OCT:OCT-short(POU,Homeobox)/NPC-OCT6-ChIP-Seq(GSE43916)/Homer | 1e-5 | -1.191e+01 | 0.0001 | 189.0 | 12.30% | 8663.8 | 8.95% | motif file (matrix) | svg |
| 52 | 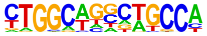 | Tlx?(NR)/NPC-H3K4me1-ChIP-Seq(GSE16256)/Homer | 1e-4 | -1.150e+01 | 0.0001 | 82.0 | 5.34% | 3119.6 | 3.22% | motif file (matrix) | svg |
| 53 | | Oct6(POU,Homeobox)/NPC-Pou3f1-ChIP-Seq(GSE35496)/Homer | 1e-4 | -1.114e+01 | 0.0001 | 119.0 | 7.75% | 5029.4 | 5.20% | motif file (matrix) | svg |

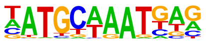

| | | | | | | | | | | | |
|---|---|---|---|---|---|---|---|---|---|---|---|
| 54 | 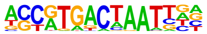 | PAX3:FKHR-fusion(Paired,Homeobox)/Rh4-PAX3:FKHR-ChIP-Seq(GSE19063)/Homer | 1e-4 | -1.106e+01 | 0.0001 | 58.0 | 3.78% | 2006.5 | 2.07% | motif file (matrix) | svg |
| 55 | 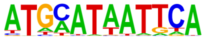 | Pit1+1bp(Homeobox)/GCrat-Pit1-ChIP-Seq(GSE58009)/Homer | 1e-4 | -1.046e+01 | 0.0002 | 102.0 | 6.64% | 4228.3 | 4.37% | motif file (matrix) | svg |
| 56 | 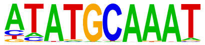 | Oct2(POU,Homeobox)/Bcell-Oct2-ChIP-Seq(GSE21512)/Homer | 1e-4 | -9.774e+00 | 0.0005 | 91.0 | 5.92% | 3739.4 | 3.86% | motif file (matrix) | svg |
| 57 | 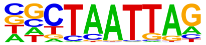 | LHX9(Homeobox)/Hct116-LHX9.V5-ChIP-Seq(GSE116822)/Homer | 1e-4 | -9.768e+00 | 0.0005 | 318.0 | 20.70% | 16347.5 | 16.89% | motif file (matrix) | svg |
| 58 | 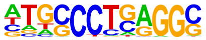 | AP-2alpha(AP2)/Hela-AP2alpha-ChIP-Seq(GSE31477)/Homer | 1e-4 | -9.372e+00 | 0.0007 | 120.0 | 7.81% | 5301.9 | 5.48% | motif file (matrix) | svg |
| 59 | 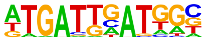 | HOXA3(Homeobox)/mEmbryo-Hoxa3-ChIP-Seq(E-MTAB-8607)/Homer | 1e-3 | -9.175e+00 | 0.0008 | 46.0 | 2.99% | 1584.6 | 1.64% | motif file (matrix) | svg |
| 60 | | NFIL3(bZIP)/HepG2-NFIL3-ChIP-Seq(Encode)/Homer | 1e-3 | -9.080e+00 | 0.0009 | 146.0 | 9.51% | 6743.6 | 6.97% | motif file (matrix) | svg |

| | | | | | | | | | | | |
|---|---|---|---|---|---|---|---|---|---|---|---|
| | 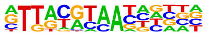 | | | | | | | | | | |
| 61 | | bZIP:IRF(bZIP,IRF)/Th17-BatF-ChIP-Seq(GSE39756)/Homer | 1e-3 | -8.974e+00 | 0.0010 | 115.0 | 7.49% | 5088.4 | 5.26% | motif file (matrix) | svg |
| | 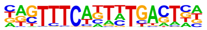 | | | | | | | | | | |
| 62 | | Oct11(POU,Homeobox)/NCIH1048-POU2F3-ChIP-seq(GSE115123)/Homer | 1e-3 | -8.671e+00 | 0.0013 | 88.0 | 5.73% | 3708.9 | 3.83% | motif file (matrix) | svg |
| | 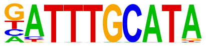 | | | | | | | | | | |
| 63 | | Unknown-ESC-element(?)/mES-Nanog-ChIP-Seq(GSE11724)/Homer | 1e-3 | -8.511e+00 | 0.0015 | 98.0 | 6.38% | 4249.1 | 4.39% | motif file (matrix) | svg |
| | 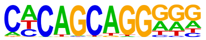 | | | | | | | | | | |
| 64 | | Gli3(Zf)/E11.5-Gli3-ChIP-Chip(GSE151646)/Homer | 1e-3 | -8.428e+00 | 0.0016 | 73.0 | 4.75% | 2966.1 | 3.06% | motif file (matrix) | svg |
| | 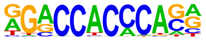 | | | | | | | | | | |
| 65 | | AP-2gamma(AP2)/MCF7-TFAP2C-ChIP-Seq(GSE21234)/Homer | 1e-3 | -8.242e+00 | 0.0019 | 148.0 | 9.64% | 6985.5 | 7.22% | motif file (matrix) | svg |
| | 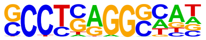 | | | | | | | | | | |
| 66 | | PBX2(Homeobox)/K562-PBX2-ChIP-Seq(Encode)/Homer | 1e-3 | -7.824e+00 | 0.0029 | 193.0 | 12.57% | 9576.9 | 9.89% | motif file (matrix) | svg |
| | 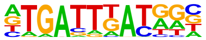 | | | | | | | | | | |
| 67 | | HLF(bZIP)/HSC-HLF.Flag-ChIP-Seq(GSE69817)/Homer | 1e-3 | -7.789e+00 | 0.0029 | 190.0 | 12.37% | 9413.3 | 9.72% | motif file (matrix) | svg |

| | | | | | | | | | | | |
|---|---|---|---|---|---|---|---|---|---|---|---|
| 68 | 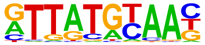 | OCT4-SOX2-TCF-NANOG(POU,Homeobox,HMG)/mES-Oct4-ChIP-Seq(GSE11431)/Homer | 1e-3 | -7.743e+00 | 0.0030 | 55.0 | 3.58% | 2132.8 | 2.20% | motif file (matrix) | svg |
| 69 | 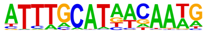 | Sox7(HMG)/ESC-Sox7-ChIP-Seq(GSE133899)/Homer | 1e-3 | -7.652e+00 | 0.0032 | 64.0 | 4.17% | 2591.6 | 2.68% | motif file (matrix) | svg |
| 70 | 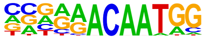 | Six1(Homeobox)/Myoblast-Six1-ChIP-Chip(GSE20150)/Homer | 1e-3 | -7.597e+00 | 0.0034 | 57.0 | 3.71% | 2245.8 | 2.32% | motif file (matrix) | svg |
| 71 | 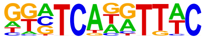 | Hoxc10(Homeobox)/EB-Hoxc10.iFlag-ChIP-Seq(GSE142377)/Homer | 1e-3 | -7.387e+00 | 0.0041 | 293.0 | 19.08% | 15447.6 | 15.96% | motif file (matrix) | svg |
| 72 | 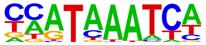 | Oct4:Sox17(POU,Homeobox,HMG)/F9-Sox17-ChIP-Seq(GSE44553)/Homer | 1e-3 | -7.232e+00 | 0.0047 | 39.0 | 2.54% | 1400.9 | 1.45% | motif file (matrix) | svg |
| 73 | 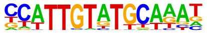 | Lhx2(Homeobox)/HFSC-Lhx2-ChIP-Seq(GSE48068)/Homer | 1e-3 | -7.141e+00 | 0.0051 | 238.0 | 15.49% | 12301.7 | 12.71% | motif file (matrix) | svg |
| 74 | 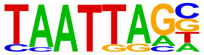 | NFkB-p65-Rel(RHD)/ThioMac-LPS-Expression(GSE23622)/Homer | 1e-2 | -6.889e+00 | 0.0065 | 17.0 | 1.11% | 445.9 | 0.46% | motif file (matrix) | svg |

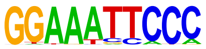

| | | | | | | | | | | | |
|---|---|---|---|---|---|---|---|---|---|---|---|
| 75 | 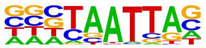 | DLX2(Homeobox)/BasalGanglia-Dlx2-ChIP-seq(GSE124936)/Homer | 1e-2 | -6.850e+00 | 0.0067 | 337.0 | 21.94% | 18188.9 | 18.79% | motif file (matrix) | svg |
| 76 | 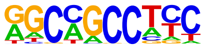 | ZNF91(Zf)/HEK-ZNF91.HA-ChIP-Seq(GSE162571)/Homer | 1e-2 | -6.822e+00 | 0.0068 | 115.0 | 7.49% | 5403.3 | 5.58% | motif file (matrix) | svg |
| 77 | 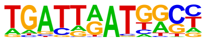 | Hoxb4(Homeobox)/ES-Hoxb4-ChIP-Seq(GSE34014)/Homer | 1e-2 | -6.738e+00 | 0.0073 | 51.0 | 3.32% | 2029.1 | 2.10% | motif file (matrix) | svg |
| 78 | 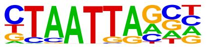 | Gsx2(Homeobox)/LGE-Gsx2.Flag-ChIP-Seq(GSE162589)/Homer | 1e-2 | -6.699e+00 | 0.0075 | 296.0 | 19.27% | 15803.2 | 16.33% | motif file (matrix) | svg |
| 79 | 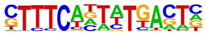 | IRF:BATF(IRF:bZIP)/pDC-Irf8-ChIP-Seq(GSE66899)/Homer | 1e-2 | -6.660e+00 | 0.0077 | 36.0 | 2.34% | 1302.9 | 1.35% | motif file (matrix) | svg |
| 80 | 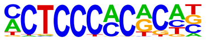 | WT1(Zf)/Kidney-WT1-ChIP-Seq(GSE90016)/Homer | 1e-2 | -6.396e+00 | 0.0098 | 85.0 | 5.53% | 3841.0 | 3.97% | motif file (matrix) | svg |
| 81 | | PAX5(Paired,Homeobox),condensed/GM12878-PAX5-ChIP-Seq(GSE32465)/Homer | 1e-2 | -6.252e+00 | 0.0112 | 25.0 | 1.63% | 820.2 | 0.85% | motif file (matrix) | svg |

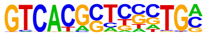

| | | | | | | | | | | | |
|---|---|---|---|---|---|---|---|---|---|---|---|
| 82 | 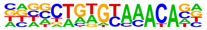 | Fox:Ebox(Forkhead,bHLH)/Panc1-Foxa2-ChIP-Seq(GSE47459)/Homer | 1e-2 | -6.218e+00 | 0.0115 | 181.0 | 11.78% | 9227.9 | 9.53% | motif file (matrix) | svg |
| 83 | 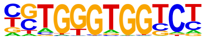 | Gli2(Zf)/GM2-Gli2-ChIP-Chip(GSE112702)/Homer | 1e-2 | -6.013e+00 | 0.0139 | 48.0 | 3.12% | 1951.7 | 2.02% | motif file (matrix) | svg |
| 84 | 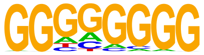 | Maz(Zf)/HepG2-Maz-ChIP-Seq(GSE31477)/Homer | 1e-2 | -5.979e+00 | 0.0142 | 153.0 | 9.96% | 7681.7 | 7.94% | motif file (matrix) | svg |
| 85 | 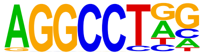 | ZFX(Zf)/mES-Zfx-ChIP-Seq(GSE11431)/Homer | 1e-2 | -5.911e+00 | 0.0150 | 186.0 | 12.11% | 9584.3 | 9.90% | motif file (matrix) | svg |
| 86 | 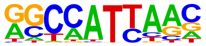 | Nanog(Homeobox)/mES-Nanog-ChIP-Seq(GSE11724)/Homer | 1e-2 | -5.867e+00 | 0.0155 | 789.0 | 51.37% | 46276.7 | 47.81% | motif file (matrix) | svg |
| 87 | 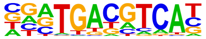 | Atf2(bZIP)/3T3L1-Atf2-ChIP-Seq(GSE56872)/Homer | 1e-2 | -5.816e+00 | 0.0162 | 73.0 | 4.75% | 3279.1 | 3.39% | motif file (matrix) | svg |
| 88 | | Stat3+il21(Stat)/CD4-Stat3-ChIP-Seq(GSE19198)/Homer | 1e-2 | -5.801e+00 | 0.0162 | 117.0 | 7.62% | 5689.9 | 5.88% | motif file (matrix) | svg |

| Rank | Motif | Name | P-value | log P-value | q-value (Benjamini) | # Target Sequences with Motif | % of Targets Sequences with Motif | # Background Sequences with Motif | % of Background Sequences with Motif | Motif File | SVG |
|---|---|---|---|---|---|---|---|---|---|---|---|
| | 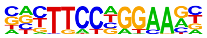 | | | | | | | | | | |
| 89 | 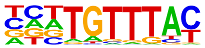 | FOXK1(Forkhead)/HEK293-FOXK1-ChIP-Seq(GSE51673)/Homer | 1e-2 | -5.767e+00 | 0.0166 | 209.0 | 13.61% | 10950.7 | 11.31% | motif file (matrix) | svg |
| 90 | 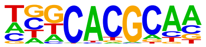 | Arnt:Ahr(bHLH)/MCF7-Arnt-ChIP-Seq(Lo_et_al.)/Homer | 1e-2 | -5.667e+00 | 0.0181 | 91.0 | 5.92% | 4275.3 | 4.42% | motif file (matrix) | svg |
| 91 | 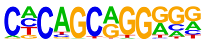 | Zic2(Zf)/ESC-Zic2-ChIP-Seq(SRP197560)/Homer | 1e-2 | -5.612e+00 | 0.0189 | 62.0 | 4.04% | 2722.1 | 2.81% | motif file (matrix) | svg |
| 92 | 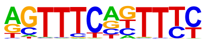 | IRF3(IRF)/BMDM-Irf3-ChIP-Seq(GSE67343)/Homer | 1e-2 | -5.518e+00 | 0.0206 | 73.0 | 4.75% | 3321.7 | 3.43% | motif file (matrix) | svg |
| 93 | 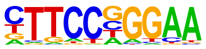 | Stat3(Stat)/mES-Stat3-ChIP-Seq(GSE11431)/Homer | 1e-2 | -5.342e+00 | 0.0243 | 82.0 | 5.34% | 3835.9 | 3.96% | motif file (matrix) | svg |
| 94 | 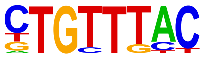 | Foxo1(Forkhead)/RAW-Foxo1-ChIP-Seq(Fan_et_al.)/Homer | 1e-2 | -5.268e+00 | 0.0259 | 343.0 | 22.33% | 19032.8 | 19.66% | motif file (matrix) | svg |
| 95 | | STAT4(Stat)/CD4-Stat4-ChIP-Seq(GSE22104)/Homer | 1e-2 | -5.207e+00 | 0.0272 | 172.0 | 11.20% | 8943.7 | 9.24% | motif file (matrix) | svg |

| | | | | | | | | | | | |
|---|---|---|---|---|---|---|---|---|---|---|---|
| | 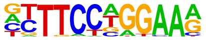 | | | | | | | | | | |
| 96 | 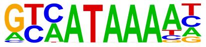 | Cdx2(Homeobox)/mES-Cdx2-ChIP-Seq(GSE14586)/Homer | 1e-2 | -5.013e+00 | 0.0327 | 163.0 | 10.61% | 8470.0 | 8.75% | motif file (matrix) | svg |
| 97 | 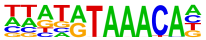 | Foxf1(Forkhead)/Lung-Foxf1-ChIP-Seq(GSE77951)/Homer | 1e-2 | -4.841e+00 | 0.0384 | 191.0 | 12.43% | 10141.8 | 10.48% | motif file (matrix) | svg |
| 98 | 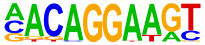 | Ets1-distal(ETS)/CD4+-PolII-ChIP-Seq(Barski_et_al.)/Homer | 1e-2 | -4.796e+00 | 0.0398 | 52.0 | 3.39% | 2299.2 | 2.38% | motif file (matrix) | svg |
| 99 | 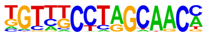 | Rfx6(HTH)/Min6b1-Rfx6.HA-ChIP-Seq(GSE62844)/Homer | 1e-2 | -4.783e+00 | 0.0399 | 154.0 | 10.03% | 8007.0 | 8.27% | motif file (matrix) | svg |
| 100 | 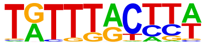 | FOXM1(Forkhead)/MCF7-FOXM1-ChIP-Seq(GSE72977)/Homer | 1e-2 | -4.748e+00 | 0.0409 | 213.0 | 13.87% | 11457.4 | 11.84% | motif file (matrix) | svg |
| 101 | 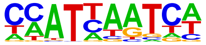 | Hoxc6(Homeobox)/EB-Hoxc6.iFlag-ChIP-Seq(GSE142377)/Homer | 1e-2 | -4.732e+00 | 0.0412 | 619.0 | 40.30% | 36131.3 | 37.33% | motif file (matrix) | svg |
| 102 | | FOXP1(Forkhead)/H9-FOXP1-ChIP-Seq(GSE31006)/Homer | 1e-2 | -4.730e+00 | 0.0412 | 85.0 | 5.53% | 4102.4 | 4.24% | motif file (matrix) | svg |

| | | | | | | | | | | | |
|---|---|---|---|---|---|---|---|---|---|---|---|
| | 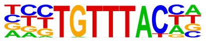 | | | | | | | | | | |
| 103 | 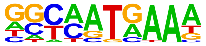 | Hoxd10(Homeobox)/ChickenMSG-Hoxd10.Flag-ChIP-Seq(GSE86088)/Homer | 1e-2 | -4.715e+00 | 0.0412 | 245.0 | 15.95% | 13358.1 | 13.80% | motif file (matrix) | svg |
| 104 | 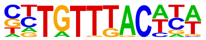 | Foxa2(Forkhead)/Liver-Foxa2-ChIP-Seq(GSE25694)/Homer | 1e-2 | -4.700e+00 | 0.0413 | 151.0 | 9.83% | 7853.3 | 8.11% | motif file (matrix) | svg |
