## Supplementary material for "A Multi-Omic Phenobank Reveals Axes of Glioblastoma Growth, Invasion, and Therapeutic Vulnerability": supp table 7

### Motifs found for: abnormal blood vessels

### Homer *de novo* Motif Results (homerpos1/)

Non-redundant Motif File of Results
Known Motif Enrichment Results
Gene Ontology Enrichment Results
If Homer is having trouble matching a motif to a known motif, try copy/pasting the matrix file into STAMP
More information on motif finding results: HOMER | Description of Results | Tips
Total target sequences = 244
Total background sequences = 96171
* - possible false positive

| Rank | Motif | P-value | log P-pvalue | % of Targets | % of Background | STD(Bg STD) | Best Match/Details | Motif File |
|---|---|---|---|---|---|---|---|---|
| 1 | 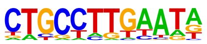 | 1e-14 | -3.421e+01 | 17.62% | 4.15% | 54.1bp (67.3bp) | PB0178.1_Sox8_2/ Jaspar(0.630) More Information \| Similar Motifs Found | motif file (matrix) |
| 2 | 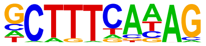 | 1e-14 | -3.363e+01 | 18.44% | 4.62% | 56.2bp (63.7bp) | TATA-Box(TBP)/ Promoter/Homer(0.794) More Information \| Similar Motifs Found | motif file (matrix) |
| 3 | 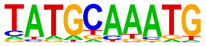 | 1e-12 | -2.963e+01 | 21.72% | 6.99% | 51.2bp (67.1bp) | POU2F3/MA0627.3/ Jaspar(0.932) More Information \| Similar Motifs Found | motif file (matrix) |
| 4 * | | 1e-11 | -2.717e+01 | 31.97% | 14.22% | 58.3bp (73.1bp) | Tcf12(bHLH)/ GM12878-Tcf12-ChIP-Seq(GSE32465)/ | motif file (matrix) |

| | | | | | | | | |
|---|---|---|---|---|---|---|---|---|
| | 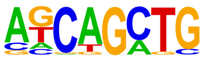 | | | | | | Homer(0.933)<br>More Information \|<br>Similar Motifs Found | |
| 5 * | 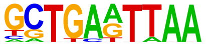 | 1e-11 | -2.562e+01 | 7.79% | 0.98% | 43.2bp (64.5bp) | LMX1B/MA0703.3/<br>Jaspar(0.725)<br>More Information \|<br>Similar Motifs Found | motif file (matrix) |
| 6 * | 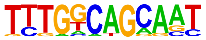 | 1e-10 | -2.355e+01 | 4.92% | 0.33% | 63.2bp (69.7bp) | PB0041.1_Mafb_1/<br>Jaspar(0.708)<br>More Information \|<br>Similar Motifs Found | motif file (matrix) |
| 7 * | 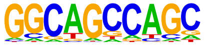 | 1e-10 | -2.329e+01 | 21.72% | 8.27% | 56.0bp (73.1bp) | ZNF91(Zf)/HEK-ZNF91.HA-ChIP-Seq(GSE162571)/<br>Homer(0.730)<br>More Information \|<br>Similar Motifs Found | motif file (matrix) |
| 8 * | 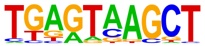 | 1e-9 | -2.290e+01 | 11.48% | 2.64% | 50.6bp (67.3bp) | MafK(bZIP)/C2C12-MafK-ChIP-Seq(GSE36030)/<br>Homer(0.710)<br>More Information \|<br>Similar Motifs Found | motif file (matrix) |
| 9 * | 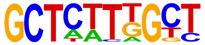 | 1e-9 | -2.239e+01 | 9.02% | 1.64% | 61.0bp (71.4bp) | POL010.1_DCE_S_III/<br>Jaspar(0.663)<br>More Information \|<br>Similar Motifs Found | motif file (matrix) |
| 10 * | 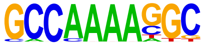 | 1e-9 | -2.221e+01 | 5.33% | 0.47% | 61.5bp (62.5bp) | PB0191.1_Tcfap2c_2/<br>Jaspar(0.723)<br>More Information \|<br>Similar Motifs Found | motif file (matrix) |
| 11 * | | 1e-9 | -2.215e+01 | 4.92% | 0.37% | 33.8bp (60.5bp) | Stat5b/MA1625.2/<br>Jaspar(0.706) | motif file (matrix) |

| | | | | | | | | |
|---|---|---|---|---|---|---|---|---|
| | 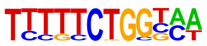 | | | | | | More Information \| Similar Motifs Found | |
| 12 * | 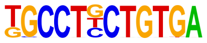 | 1e-9 | -2.116e+01 | 2.87% | 0.07% | 62.3bp (75.2bp) | Zic2/MA1629.2/ Jaspar(0.822) More Information \| Similar Motifs Found | motif file (matrix) |
| 13 * | 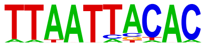 | 1e-8 | -2.072e+01 | 5.74% | 0.64% | 61.0bp (62.2bp) | mix-a/MA0621.2/ Jaspar(0.811) More Information \| Similar Motifs Found | motif file (matrix) |
| 14 * | 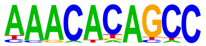 | 1e-8 | -1.996e+01 | 7.38% | 1.23% | 48.1bp (68.4bp) | RUNX-AML(Runt)/ CD4+-PolII-ChIP-Seq(Barski_et_al.)/ Homer(0.713) More Information \| Similar Motifs Found | motif file (matrix) |
| 15 * | 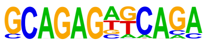 | 1e-8 | -1.857e+01 | 2.87% | 0.10% | 49.9bp (64.3bp) | Erra(NR)/HepG2-Erra-ChIP-Seq(GSE31477)/ Homer(0.715) More Information \| Similar Motifs Found | motif file (matrix) |
| 16 * | 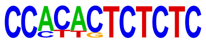 | 1e-7 | -1.829e+01 | 2.05% | 0.03% | 12.3bp (46.5bp) | PRDM1/MA0508.4/ Jaspar(0.646) More Information \| Similar Motifs Found | motif file (matrix) |
| 17 * | 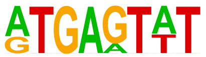 | 1e-6 | -1.559e+01 | 11.48% | 3.72% | 57.9bp (63.2bp) | MSANTD3/MA1523.2/ Jaspar(0.745) More Information \| Similar Motifs Found | motif file (matrix) |
| 18 * | | 1e-5 | -1.379e+01 | 8.20% | 2.27% | 54.5bp (58.8bp) | PB0192.1_Tcfap2e_2/ Jaspar(0.715) | motif file (matrix) |

| | | | | | | | | |
|---|---|---|---|---|---|---|---|---|
| | 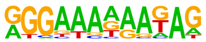 | | | | | | More Information \| Similar Motifs Found | |
| 19 * | 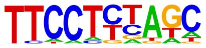 | 1e-5 | -1.290e+01 | 9.02% | 2.85% | 58.4bp (63.6bp) | IKZF2/MA2326.1/ Jaspar(0.760) More Information \| Similar Motifs Found | motif file (matrix) |

### Motifs found for: defined tumor border

### Homer *de novo* Motif Results (homerpos2/)

Non-redundant Motif File of Results
Known Motif Enrichment Results
Gene Ontology Enrichment Results
If Homer is having trouble matching a motif to a known motif, try copy/pasting the matrix file into STAMP
More information on motif finding results: HOMER | Description of Results | Tips
Total target sequences = 86
Total background sequences = 98295
* - possible false positive

| Rank | Motif | P-value | log P-pvalue | % of Targets | % of Background | STD(Bg STD) | Best Match/Details | Motif File |
|---|---|---|---|---|---|---|---|---|
| 1 * | 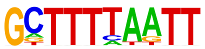 | 1e-9 | -2.103e+01 | 12.79% | 0.95% | 50.1bp (61.2bp) | MYNN(Zf)/HEK293-MYNN.eGFP-ChIP-Seq(Encode)/Homer(0.746) More Information \| Similar Motifs Found | motif file (matrix) |
| 2 * | 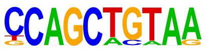 | 1e-8 | -2.002e+01 | 9.30% | 0.39% | 57.2bp (59.1bp) | Tcf21(bHLH)/ArterySmoothMuscle-Tcf21-ChIP-Seq(GSE61369)/Homer(0.821) More Information \| Similar Motifs Found | motif file (matrix) |
| 3 * | 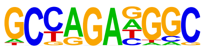 | 1e-7 | -1.732e+01 | 13.95% | 1.71% | 52.3bp (62.7bp) | PB0191.1_Tcfap2c_2/Jaspar(0.725) More Information \| Similar Motifs Found | motif file (matrix) |
| 4 * | | 1e-7 | -1.630e+01 | 6.98% | 0.24% | 53.8bp (59.1bp) | Arid5a/MA0602.2/Jaspar(0.638) | motif file (matrix) |

| | | | | | | | | |
|---|---|---|---|---|---|---|---|---|
| | 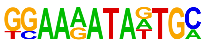 | | | | | | More Information \| Similar Motifs Found | |
| 5 * | 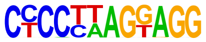 | 1e-7 | -1.612e+01 | 4.65% | 0.05% | 42.2bp (57.9bp) | GLIS3(Zf)/Thyroid-Glis3.GFP-ChIP-Seq(GSE103297)/Homer(0.709) More Information \| Similar Motifs Found | motif file (matrix) |
| 6 * | 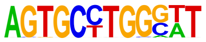 | 1e-6 | -1.528e+01 | 6.98% | 0.29% | 59.3bp (62.7bp) | ZSCAN21/MA2336.1/Jaspar(0.721) More Information \| Similar Motifs Found | motif file (matrix) |
| 7 * | 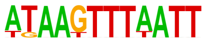 | 1e-6 | -1.500e+01 | 4.65% | 0.06% | 64.4bp (63.2bp) | PH0075.1_Hoxd10/Jaspar(0.723) More Information \| Similar Motifs Found | motif file (matrix) |
| 8 * | 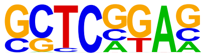 | 1e-4 | -9.747e+00 | 22.09% | 8.21% | 55.9bp (63.5bp) | POL013.1_MED-1/Jaspar(0.683) More Information \| Similar Motifs Found | motif file (matrix) |
| 9 * | 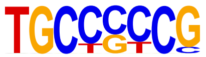 | 1e-2 | -6.726e+00 | 5.81% | 0.91% | 57.8bp (63.8bp) | HIC2/MA0738.2/Jaspar(0.735) More Information \| Similar Motifs Found | motif file (matrix) |
| 10 * | 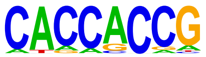 | 1e-2 | -5.565e+00 | 13.95% | 5.77% | 54.8bp (64.0bp) | ZBTB7C/MA0695.2/Jaspar(0.762) More Information \| Similar Motifs Found | motif file (matrix) |
| 11 * | | 1e-1 | -4.329e+00 | 2.33% | 0.20% | 1.5bp (104.4bp) | ZNF93/MA1721.2/Jaspar(0.730) | motif file (matrix) |

More Information |
Similar Motifs Found

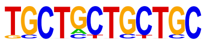

### Motifs found for: diffuse tumor border

### Homer *de novo* Motif Results (homerpos3/)

Non-redundant Motif File of Results
Known Motif Enrichment Results
Gene Ontology Enrichment Results
If Homer is having trouble matching a motif to a known motif, try copy/pasting the matrix file into STAMP
More information on motif finding results: HOMER | Description of Results | Tips
Total target sequences = 709
Total background sequences = 96194
* - possible false positive

| Rank | Motif | P-value | log P-pvalue | % of Targets | % of Background | STD(Bg STD) | Best Match/Details | Motif File |
|---|---|---|---|---|---|---|---|---|
| 1 | 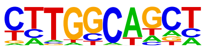 | 1e-27 | -6.360e+01 | 45.28% | 26.05% | 55.1bp (68.3bp) | NF1-halfsite(CTF)/ LNCaP-NF1-ChIP-Seq(Unpublished)/ Homer(0.884) More Information \| Similar Motifs Found | motif file (matrix) |
| 2 | 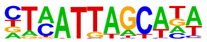 | 1e-21 | -5.065e+01 | 7.76% | 1.48% | 51.2bp (64.0bp) | POU3F2/MA0787.1/ Jaspar(0.870) More Information \| Similar Motifs Found | motif file (matrix) |
| 3 | 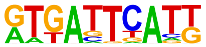 | 1e-19 | -4.531e+01 | 8.18% | 1.86% | 54.2bp (64.4bp) | FOSL2::JUNB/ MA1138.2/ Jaspar(0.860) More Information \| Similar Motifs Found | motif file (matrix) |
| 4 | | 1e-18 | -4.157e+01 | 30.89% | 17.33% | 58.3bp (72.9bp) | Olig2/MA1997.2/ Jaspar(0.948) | motif file (matrix) |

| | | | | | | | | |
|---|---|---|---|---|---|---|---|---|
| | 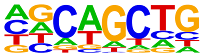 | | | | | | More Information \| Similar Motifs Found | |
| 5 | 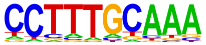 | 1e-17 | -4.110e+01 | 27.64% | 14.85% | 55.6bp (65.5bp) | Oct11(POU,Homeobox)/NCIH1048-POU2F3-ChIP-seq(GSE115123)/Homer(0.718) More Information \| Similar Motifs Found | motif file (matrix) |
| 6 | 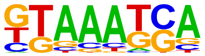 | 1e-17 | -3.989e+01 | 43.02% | 27.88% | 57.0bp (64.8bp) | Hoxd9(Homeobox)/EB-Hoxd9.HA-ChIP-Seq(GSE142377)/Homer(0.771) More Information \| Similar Motifs Found | motif file (matrix) |
| 7 | 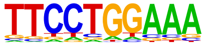 | 1e-17 | -3.917e+01 | 29.34% | 16.45% | 57.0bp (65.7bp) | Bcl6(Zf)/Liver-Bcl6-ChIP-Seq(GSE31578)/Homer(0.942) More Information \| Similar Motifs Found | motif file (matrix) |
| 8 | 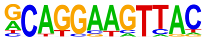 | 1e-13 | -3.151e+01 | 2.26% | 0.14% | 53.6bp (64.5bp) | ETV1/MA0761.3/Jaspar(0.830) More Information \| Similar Motifs Found | motif file (matrix) |
| 9 | 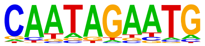 | 1e-13 | -3.067e+01 | 11.71% | 4.69% | 57.6bp (64.3bp) | HOXB13/MA0901.3/Jaspar(0.691) More Information \| Similar Motifs Found | motif file (matrix) |
| 10 | 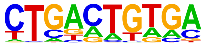 | 1e-13 | -3.005e+01 | 25.67% | 14.95% | 55.4bp (65.7bp) | PB0170.1_Sox17_2/Jaspar(0.637) More Information \| Similar Motifs Found | motif file (matrix) |
| 11 | | 1e-12 | -2.847e+01 | 1.83% | 0.10% | 57.1bp (62.2bp) | ZNF24/MA1124.1/Jaspar(0.639) | motif file (matrix) |

| | | | | | | | | |
|---|---|---|---|---|---|---|---|---|
| | 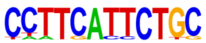 | | | | | | More Information \| Similar Motifs Found | |
| 12 | 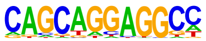 | 1e-12 | -2.843e+01 | 4.51% | 0.92% | 54.2bp (70.7bp) | Zic3(Zf)/mES-Zic3-ChIP-Seq(GSE37889)/Homer(0.818) More Information \| Similar Motifs Found | motif file (matrix) |
| 13 | 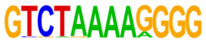 | 1e-12 | -2.789e+01 | 1.27% | 0.03% | 69.5bp (48.1bp) | TATA-Box(TBP)/Promoter/Homer(0.681) More Information \| Similar Motifs Found | motif file (matrix) |
| 14 * | 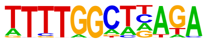 | 1e-11 | -2.738e+01 | 3.95% | 0.73% | 54.0bp (65.3bp) | NFIX/MA0671.2/Jaspar(0.683) More Information \| Similar Motifs Found | motif file (matrix) |
| 15 * | 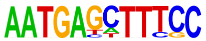 | 1e-11 | -2.692e+01 | 1.13% | 0.02% | 55.0bp (64.2bp) | PH0152.1_Pou6f1_2/Jaspar(0.717) More Information \| Similar Motifs Found | motif file (matrix) |
| 16 * | 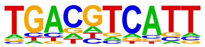 | 1e-11 | -2.620e+01 | 11.99% | 5.30% | 50.2bp (65.3bp) | JDP2/MA0656.2/Jaspar(0.915) More Information \| Similar Motifs Found | motif file (matrix) |
| 17 * | 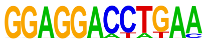 | 1e-10 | -2.466e+01 | 1.13% | 0.03% | 49.2bp (79.3bp) | Znf263(Zf)/K562-Znf263-ChIP-Seq(GSE31477)/Homer(0.723) More Information \| Similar Motifs Found | motif file (matrix) |
| 18 * | | 1e-10 | -2.460e+01 | 2.26% | 0.23% | 61.1bp (67.7bp) | Brn1(POU,Homeobox)/NPC-Brn1-ChIP- | motif file (matrix) |

| | | | | | | | | |
|---|---|---|---|---|---|---|---|---|
| | 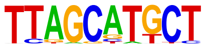 | | | | | | Seq(GSE35496)/<br>Homer(0.732)<br>More Information \|<br>Similar Motifs Found | |
| 19 * | 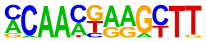 | 1e-10 | -2.456e+01 | 2.96% | 0.44% | 46.8bp (63.9bp) | Nr2e3/MA0164.2/<br>Jaspar(0.662)<br>More Information \|<br>Similar Motifs Found | motif file (matrix) |
| 20 * | 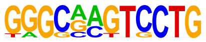 | 1e-10 | -2.328e+01 | 1.27% | 0.05% | 41.3bp (58.3bp) | Ikzf3/MA1992.2/<br>Jaspar(0.657)<br>More Information \|<br>Similar Motifs Found | motif file (matrix) |
| 21 * | 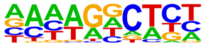 | 1e-9 | -2.159e+01 | 5.36% | 1.64% | 58.0bp (66.5bp) | PB0203.1_Zfp691_2/<br>Jaspar(0.599)<br>More Information \|<br>Similar Motifs Found | motif file (matrix) |
| 22 * | 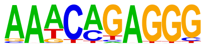 | 1e-9 | -2.121e+01 | 4.65% | 1.29% | 58.3bp (63.0bp) | NeuroG2(bHLH)/<br>Fibroblast-NeuroG2-<br>ChIP-Seq(GSE75910)/<br>Homer(0.692)<br>More Information \|<br>Similar Motifs Found | motif file (matrix) |
| 23 * | 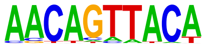 | 1e-8 | -2.001e+01 | 3.81% | 0.94% | 56.0bp (62.9bp) | DMRTA2/MA1478.2/<br>Jaspar(0.721)<br>More Information \|<br>Similar Motifs Found | motif file (matrix) |
| 24 * | 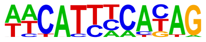 | 1e-8 | -1.984e+01 | 4.23% | 1.15% | 51.8bp (60.1bp) | TEAD4/MA0809.3/<br>Jaspar(0.710)<br>More Information \|<br>Similar Motifs Found | motif file (matrix) |
| 25 * | | 1e-8 | -1.878e+01 | 11.00% | 5.48% | 54.8bp (61.4bp) | MEF2A/MA0052.5/<br>Jaspar(0.830) | motif file (matrix) |

| | | | | | | | | |
|---|---|---|---|---|---|---|---|---|
| | 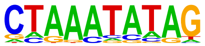 | | | | | | More Information \| Similar Motifs Found | |
| 26 * | 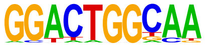 | 1e-7 | -1.648e+01 | 3.39% | 0.91% | 61.0bp (63.9bp) | Hand1/MA2123.1/ Jaspar(0.689) More Information \| Similar Motifs Found | motif file (matrix) |
| 27 * | 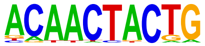 | 1e-7 | -1.619e+01 | 2.82% | 0.66% | 54.2bp (59.8bp) | OSR1/MA1542.2/ Jaspar(0.652) More Information \| Similar Motifs Found | motif file (matrix) |
| 28 * | 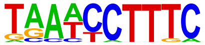 | 1e-6 | -1.550e+01 | 1.55% | 0.19% | 67.0bp (62.3bp) | ZNF184/MA2120.1/ Jaspar(0.698) More Information \| Similar Motifs Found | motif file (matrix) |
| 29 * | 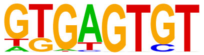 | 1e-5 | -1.220e+01 | 9.73% | 5.52% | 51.3bp (59.5bp) | MSANTD3/MA1523.2/ Jaspar(0.888) More Information \| Similar Motifs Found | motif file (matrix) |
| 30 * | 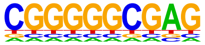 | 1e-3 | -8.690e+00 | 9.87% | 6.30% | 57.9bp (57.5bp) | KLF16/MA0741.1/ Jaspar(0.765) More Information \| Similar Motifs Found | motif file (matrix) |

### Motifs found for: gliomatosis cerebri

### Homer *de novo* Motif Results (homerpos4/)

Non-redundant Motif File of Results
Known Motif Enrichment Results
Gene Ontology Enrichment Results
If Homer is having trouble matching a motif to a known motif, try copy/pasting the matrix file into STAMP
More information on motif finding results: HOMER | Description of Results | Tips
Total target sequences = 55
Total background sequences = 97546
* - possible false positive

| Rank | Motif | P-value | log P-pvalue | % of Targets | % of Background | STD(Bg STD) | Best Match/Details | Motif File |
|---|---|---|---|---|---|---|---|---|
| 1 * | 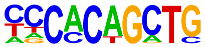 | 1e-10 | -2.475e+01 | 20.00% | 1.08% | 61.7bp (67.4bp) | Ptf1A/MA1619.2/ Jaspar(0.849) More Information \| Similar Motifs Found | motif file (matrix) |
| 2 * | 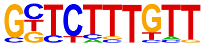 | 1e-9 | -2.297e+01 | 27.27% | 3.15% | 55.2bp (63.5bp) | PB0071.1_Sox4_1/ Jaspar(0.867) More Information \| Similar Motifs Found | motif file (matrix) |
| 3 * | 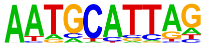 | 1e-9 | -2.247e+01 | 30.91% | 4.51% | 45.3bp (66.4bp) | PAX3/MA0780.1/ Jaspar(0.697) More Information \| Similar Motifs Found | motif file (matrix) |
| 4 * | | 1e-9 | -2.110e+01 | 50.91% | 15.23% | 55.0bp (67.4bp) | Nfatc1/MA0624.3/ Jaspar(0.913) | motif file (matrix) |

| | | | | | | | | |
|---|---|---|---|---|---|---|---|---|
| | 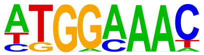  | | | | | | More Information \| Similar Motifs Found | |
| 5 * | 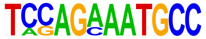  | 1e-8 | -2.035e+01 | 7.27% | 0.03% | 23.0bp (63.5bp) | ZBTB26/MA1579.2/ Jaspar(0.672) More Information \| Similar Motifs Found | motif file (matrix) |
| 6 * | 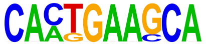  | 1e-8 | -1.984e+01 | 10.91% | 0.21% | 53.1bp (63.2bp) | OSR2/MA1646.2/ Jaspar(0.667) More Information \| Similar Motifs Found | motif file (matrix) |
| 7 * | 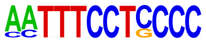  | 1e-8 | -1.879e+01 | 7.27% | 0.04% | 41.4bp (57.0bp) | ZNF263/MA0528.3/ Jaspar(0.769) More Information \| Similar Motifs Found | motif file (matrix) |
| 8 * | 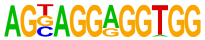  | 1e-7 | -1.642e+01 | 7.27% | 0.07% | 56.3bp (64.8bp) | ETS:RUNX(ETS,Runt)/ Jurkat-RUNX1-ChIP-Seq(GSE17954)/ Homer(0.760) More Information \| Similar Motifs Found | motif file (matrix) |
| 9 * | 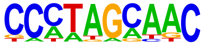  | 1e-7 | -1.637e+01 | 20.00% | 2.43% | 54.5bp (65.7bp) | PB0055.1_Rfx4_1/ Jaspar(0.914) More Information \| Similar Motifs Found | motif file (matrix) |
| 10 * | 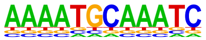  | 1e-5 | -1.373e+01 | 5.45% | 0.03% | 46.0bp (62.8bp) | Oct2(POU,Homeobox)/ Bcell-Oct2-ChIP-Seq(GSE21512)/ Homer(0.814) More Information \| Similar Motifs Found | motif file (matrix) |
| 11 * | | 1e-5 | -1.213e+01 | 5.45% | 0.06% | 57.9bp (64.1bp) | LHX2/MA0700.3/ Jaspar(0.782) | motif file (matrix) |

More Information | Similar Motifs Found

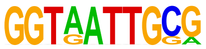

### Motifs found for: gray matter invasion

### Homer *de novo* Motif Results (homerpos5/)

Non-redundant Motif File of Results
Known Motif Enrichment Results
Gene Ontology Enrichment Results
If Homer is having trouble matching a motif to a known motif, try copy/pasting the matrix file into STAMP
More information on motif finding results: HOMER | Description of Results | Tips
Total target sequences = 66
Total background sequences = 98076
* - possible false positive

| Rank | Motif | P-value | log P-pvalue | % of Targets | % of Background | STD(Bg STD) | Best Match/Details | Motif File |
|---|---|---|---|---|---|---|---|---|
| 1 | 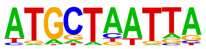 | 1e-12 | -2.857e+01 | 27.27% | 2.90% | 54.7bp (63.3bp) | Brn1(POU,Homeobox)/NPC-Brn1-ChIP-Seq(GSE35496)/Homer(0.871) More Information \| Similar Motifs Found | motif file (matrix) |
| 2 * | 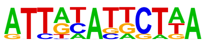 | 1e-11 | -2.574e+01 | 16.67% | 0.81% | 45.0bp (61.0bp) | Oct4:Sox17(POU,Homeobox,HMG)/F9-Sox17-ChIP-Seq(GSE44553)/Homer(0.723) More Information \| Similar Motifs Found | motif file (matrix) |
| 3 * | 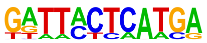 | 1e-10 | -2.453e+01 | 10.61% | 0.16% | 45.7bp (63.8bp) | Brn2(POU,Homeobox)/NPC-Brn2-ChIP-Seq(GSE35496)/Homer(0.674) More Information \| Similar Motifs Found | motif file (matrix) |
| 4 * | | 1e-10 | -2.403e+01 | 10.61% | 0.18% | 47.5bp (64.8bp) | Foxj2/MA0614.1/Jaspar(0.638) | motif file (matrix) |

| | | | | | | | | |
|---|---|---|---|---|---|---|---|---|
| | 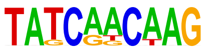 | | | | | | More Information \| Similar Motifs Found | |
| 5 * | 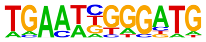 | 1e-10 | -2.398e+01 | 22.73% | 2.38% | 43.8bp (63.7bp) | PB0170.1_Sox17_2/ Jaspar(0.615) More Information \| Similar Motifs Found | motif file (matrix) |
| 6 * | 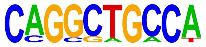 | 1e-9 | -2.268e+01 | 15.15% | 0.80% | 51.7bp (65.1bp) | Tlx?(NR)/NPC-H3K4me1-ChIP-Seq(GSE16256)/ Homer(0.719) More Information \| Similar Motifs Found | motif file (matrix) |
| 7 * | 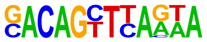 | 1e-9 | -2.135e+01 | 9.09% | 0.14% | 44.9bp (57.6bp) | POL009.1_DCE_S_II/ Jaspar(0.669) More Information \| Similar Motifs Found | motif file (matrix) |
| 8 * | 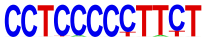 | 1e-9 | -2.085e+01 | 10.61% | 0.28% | 59.0bp (60.4bp) | SP5/MA1965.2/ Jaspar(0.728) More Information \| Similar Motifs Found | motif file (matrix) |
| 9 * | 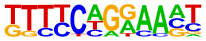 | 1e-8 | -1.993e+01 | 22.73% | 3.21% | 52.3bp (61.9bp) | STAT1(Stat)/HelaS3-STAT1-ChIP-Seq(GSE12782)/ Homer(0.829) More Information \| Similar Motifs Found | motif file (matrix) |
| 10 * | 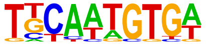 | 1e-8 | -1.881e+01 | 25.76% | 4.68% | 52.7bp (62.6bp) | SOX13/MA1120.2/ Jaspar(0.685) More Information \| Similar Motifs Found | motif file (matrix) |
| 11 * | | 1e-8 | -1.868e+01 | 7.58% | 0.10% | 28.5bp (62.1bp) | Erra(NR)/HepG2-Erra-ChIP-Seq(GSE31477)/ | motif file (matrix) |

| | | | | | | | | |
|---|---|---|---|---|---|---|---|---|
| | 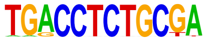 | | | | | | Homer(0.759)<br>More Information \|<br>Similar Motifs Found | |
| 12 * | 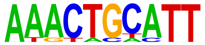 | 1e-7 | -1.828e+01 | 13.64% | 0.93% | 51.2bp (64.5bp) | PB0146.1_Mafk_2/<br>Jaspar(0.670)<br>More Information \|<br>Similar Motifs Found | motif file (matrix) |

### Motifs found for: perineuronal satellitosis

### Homer *de novo* Motif Results (homerpos6/)

Non-redundant Motif File of Results
Known Motif Enrichment Results
Gene Ontology Enrichment Results
If Homer is having trouble matching a motif to a known motif, try copy/pasting the matrix file into STAMP
More information on motif finding results: HOMER | Description of Results | Tips
Total target sequences = 188
Total background sequences = 97553
* - possible false positive

| Rank | Motif | P-value | log P-pvalue | % of Targets | % of Background | STD(Bg STD) | Best Match/Details | Motif File |
|---|---|---|---|---|---|---|---|---|
| 1 | 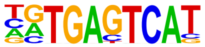 | 1e-26 | -6.182e+01 | 19.15% | 1.61% | 53.6bp (61.7bp) | FOSL2::JUN/ MA1130.2/ Jaspar(0.976) More Information \| Similar Motifs Found | motif file (matrix) |
| 2 | 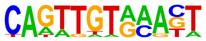 | 1e-13 | -3.200e+01 | 7.98% | 0.44% | 40.3bp (61.7bp) | MYB/MA0100.4/ Jaspar(0.730) More Information \| Similar Motifs Found | motif file (matrix) |
| 3 | 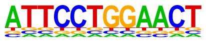 | 1e-13 | -3.061e+01 | 2.66% | 0.00% | 36.0bp (75.6bp) | STAT1/MA0137.4/ Jaspar(0.809) More Information \| Similar Motifs Found | motif file (matrix) |
| 4 | | 1e-12 | -2.932e+01 | 15.43% | 2.85% | 57.7bp (62.9bp) | Plagl1/MA1615.2/ Jaspar(0.837) | motif file (matrix) |

| | | | | | | | | |
|---|---|---|---|---|---|---|---|---|
| | 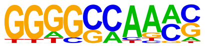 | | | | | | More Information \| Similar Motifs Found | |
| 5 | 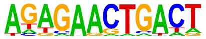 | 1e-12 | -2.765e+01 | 8.51% | 0.72% | 45.7bp (62.1bp) | ZNF528(Zf)/HEK293-ZNF528.GFP-ChIP-Seq(GSE58341)/Homer(0.656) More Information \| Similar Motifs Found | motif file (matrix) |
| 6 * | 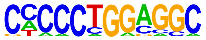 | 1e-11 | -2.545e+01 | 5.32% | 0.20% | 59.8bp (63.1bp) | BORIS(Zf)/K562-CTCFL-ChIP-Seq(GSE32465)/Homer(0.732) More Information \| Similar Motifs Found | motif file (matrix) |
| 7 * | 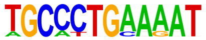 | 1e-10 | -2.489e+01 | 3.19% | 0.03% | 45.0bp (43.4bp) | THAP1/MA0597.3/Jaspar(0.706) More Information \| Similar Motifs Found | motif file (matrix) |
| 8 * | 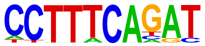 | 1e-10 | -2.480e+01 | 8.51% | 0.87% | 50.9bp (59.6bp) | LEF1(HMG)/H1-LEF1-ChIP-Seq(GSE64758)/Homer(0.701) More Information \| Similar Motifs Found | motif file (matrix) |
| 9 * | 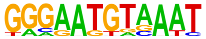 | 1e-10 | -2.390e+01 | 5.32% | 0.24% | 53.4bp (60.9bp) | TEAD2/MA1121.2/Jaspar(0.733) More Information \| Similar Motifs Found | motif file (matrix) |
| 10 * | 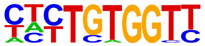 | 1e-10 | -2.342e+01 | 13.83% | 2.92% | 58.7bp (60.7bp) | Runx1/MA0002.3/Jaspar(0.935) More Information \| Similar Motifs Found | motif file (matrix) |
| 11 * | | 1e-10 | -2.331e+01 | 7.45% | 0.68% | 63.2bp (63.0bp) | PB0183.1_Sry_2/Jaspar(0.758) | motif file (matrix) |

| | | | | | | | | |
|---|---|---|---|---|---|---|---|---|
| | 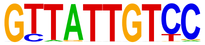 | | | | | | More Information \| Similar Motifs Found | |
| 12 * | 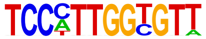 | 1e-9 | -2.292e+01 | 2.66% | 0.02% | 59.8bp (57.0bp) | Ebf2/MA1604.2/ Jaspar(0.695) More Information \| Similar Motifs Found | motif file (matrix) |
| 13 * | 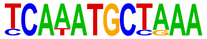 | 1e-9 | -2.210e+01 | 3.19% | 0.04% | 45.0bp (63.2bp) | Brn1(POU,Homeobox)/ NPC-Brn1-ChIP-Seq(GSE35496)/ Homer(0.651) More Information \| Similar Motifs Found | motif file (matrix) |
| 14 * | 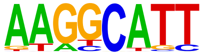 | 1e-9 | -2.180e+01 | 28.19% | 11.46% | 50.8bp (64.7bp) | ZNF528(Zf)/HEK293-ZNF528.GFP-ChIP-Seq(GSE58341)/ Homer(0.737) More Information \| Similar Motifs Found | motif file (matrix) |
| 15 * | 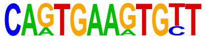 | 1e-9 | -2.167e+01 | 2.66% | 0.02% | 43.6bp (48.3bp) | NKX2-4/MA2003.2/ Jaspar(0.685) More Information \| Similar Motifs Found | motif file (matrix) |
| 16 * | 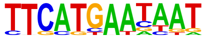 | 1e-9 | -2.112e+01 | 3.72% | 0.09% | 38.3bp (65.6bp) | PH0150.1_Pou4f3/ Jaspar(0.716) More Information \| Similar Motifs Found | motif file (matrix) |
| 17 * | 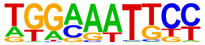 | 1e-9 | -2.104e+01 | 10.11% | 1.69% | 45.1bp (65.6bp) | NFkB-p65-Rel(RHD)/ ThioMac-LPS-Expression(GSE23622)/ Homer(0.824) More Information \| Similar Motifs Found | motif file (matrix) |
| 18 * | | 1e-8 | -2.048e+01 | 5.85% | 0.45% | 55.7bp (61.9bp) | ZNF35/MA2333.1/ Jaspar(0.670) | motif file (matrix) |

| | | | | | | | | |
|---|---|---|---|---|---|---|---|---|
| | 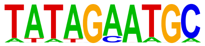 | | | | | | More Information \| Similar Motifs Found | |
| 19 * | 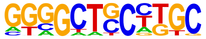 | 1e-8 | -2.027e+01 | 5.85% | 0.46% | 48.4bp (66.9bp) | ZNF165(Zf)/WHIM12-ZNF165-ChIP-Seq(GSE65937)/Homer(0.632) More Information \| Similar Motifs Found | motif file (matrix) |
| 20 * | 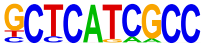 | 1e-8 | -1.969e+01 | 4.26% | 0.18% | 54.1bp (68.3bp) | ZNF669(Zf)/HEK293-ZNF669.GFP-ChIP-Seq(GSE58341)/Homer(0.690) More Information \| Similar Motifs Found | motif file (matrix) |
| 21 * | 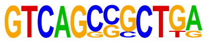 | 1e-8 | -1.864e+01 | 2.13% | 0.01% | 43.6bp (34.0bp) | Mafb/MA0117.3/Jaspar(0.648) More Information \| Similar Motifs Found | motif file (matrix) |
| 22 * | 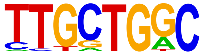 | 1e-6 | -1.599e+01 | 14.36% | 4.49% | 55.8bp (65.7bp) | PB0041.1_Mafb_1/Jaspar(0.783) More Information \| Similar Motifs Found | motif file (matrix) |
| 23 * | 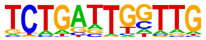 | 1e-6 | -1.427e+01 | 12.77% | 4.00% | 47.4bp (63.5bp) | NFYA/MA0060.4/Jaspar(0.726) More Information \| Similar Motifs Found | motif file (matrix) |
| 24 * | 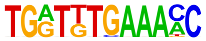 | 1e-5 | -1.320e+01 | 1.06% | 0.00% | 13.5bp (0.0bp) | CHR(?)/Hela-CellCycle-Expression/Homer(0.772) More Information \| Similar Motifs Found | motif file (matrix) |
| 25 * | | 1e-4 | -1.028e+01 | 15.43% | 6.83% | 59.1bp (62.3bp) | ZNF317/MA1593.2/Jaspar(0.750) | motif file (matrix) |

More Information |
Similar Motifs Found

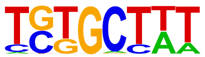

26 * 

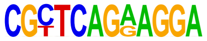

1e-2

-5.560e+00

0.53%

0.00%

0.0bp (22.6bp)

PB0205.1_Zic1_2/
Jaspar(0.667)
More Information |
Similar Motifs Found

motif file (matrix)

### Motifs found for: perivascular invasion

### Homer *de novo* Motif Results (homerpos7/)

Non-redundant Motif File of Results
Known Motif Enrichment Results
Gene Ontology Enrichment Results
If Homer is having trouble matching a motif to a known motif, try copy/pasting the matrix file into STAMP
More information on motif finding results: HOMER | Description of Results | Tips
Total target sequences = 190
Total background sequences = 98355
* - possible false positive

| Rank | Motif | P-value | log P-pvalue | % of Targets | % of Background | STD(Bg STD) | Best Match/Details | Motif File |
|---|---|---|---|---|---|---|---|---|
| 1 | 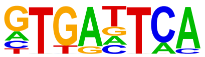 | 1e-17 | -4.126e+01 | 35.26% | 11.04% | 57.4bp (63.8bp) | Fos(bZIP)/TSC-Fos-ChIP-Seq(GSE110950)/Homer(0.935)<br>More Information \| Similar Motifs Found | motif file (matrix) |
| 2 | 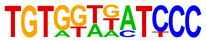 | 1e-12 | -2.813e+01 | 4.74% | 0.10% | 63.3bp (59.2bp) | RUNX2(Runt)/PCa-RUNX2-ChIP-Seq(GSE33889)/Homer(0.658)<br>More Information \| Similar Motifs Found | motif file (matrix) |
| 3 * | 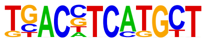 | 1e-11 | -2.719e+01 | 4.74% | 0.11% | 44.0bp (69.6bp) | ATF2/MA1632.2/Jaspar(0.795)<br>More Information \| Similar Motifs Found | motif file (matrix) |
| 4 * | | 1e-11 | -2.592e+01 | 3.16% | 0.02% | 45.7bp (48.1bp) | PH0121.1_Obox1/Jaspar(0.595) | motif file (matrix) |

| | | | | | | | | |
|---|---|---|---|---|---|---|---|---|
| | 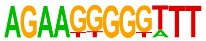 | | | | | | More Information \| Similar Motifs Found | |
| 5 * | 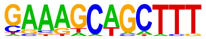 | 1e-10 | -2.528e+01 | 7.89% | 0.70% | 51.1bp (63.4bp) | SCL(bHLH)/HPC7-Scl-ChIP-Seq(GSE13511)/Homer(0.624) More Information \| Similar Motifs Found | motif file (matrix) |
| 6 * | 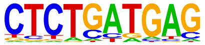 | 1e-10 | -2.435e+01 | 20.00% | 5.84% | 46.2bp (63.7bp) | POU6F1/MA1549.2/Jaspar(0.668) More Information \| Similar Motifs Found | motif file (matrix) |
| 7 * | 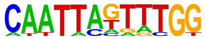 | 1e-9 | -2.204e+01 | 6.32% | 0.49% | 45.8bp (59.4bp) | LHX2/MA0700.3/Jaspar(0.740) More Information \| Similar Motifs Found | motif file (matrix) |
| 8 * | 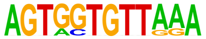 | 1e-9 | -2.166e+01 | 2.63% | 0.02% | 19.1bp (40.0bp) | Eomes(T-box)/H9-Eomes-ChIP-Seq(GSE26097)/Homer(0.646) More Information \| Similar Motifs Found | motif file (matrix) |
| 9 * | 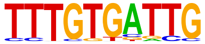 | 1e-9 | -2.146e+01 | 8.42% | 1.09% | 53.5bp (64.9bp) | PB0119.1_Foxa2_2/Jaspar(0.708) More Information \| Similar Motifs Found | motif file (matrix) |
| 10 * | 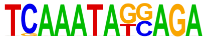 | 1e-8 | -2.044e+01 | 2.63% | 0.02% | 57.7bp (34.0bp) | PB0106.1_Arid5a_2/Jaspar(0.639) More Information \| Similar Motifs Found | motif file (matrix) |
| 11 * | | 1e-8 | -2.041e+01 | 3.16% | 0.05% | 27.6bp (59.7bp) | ESRRB/MA0141.4/Jaspar(0.681) | motif file (matrix) |

| | | | | | | | | |
|---|---|---|---|---|---|---|---|---|
| | 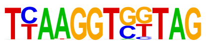 | | | | | | More Information \| Similar Motifs Found | |
| 12 * | 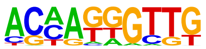 | 1e-8 | -1.943e+01 | 18.95% | 6.34% | 61.0bp (63.3bp) | SOX9/MA0077.2/ Jaspar(0.669) More Information \| Similar Motifs Found | motif file (matrix) |
| 13 * | 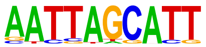 | 1e-8 | -1.868e+01 | 7.89% | 1.15% | 58.4bp (63.6bp) | Brn1(POU,Homeobox)/ NPC-Brn1-ChIP-Seq(GSE35496)/ Homer(0.831) More Information \| Similar Motifs Found | motif file (matrix) |
| 14 * | 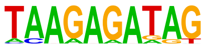 | 1e-8 | -1.845e+01 | 6.32% | 0.68% | 54.9bp (58.6bp) | PRDM1/MA0508.4/ Jaspar(0.703) More Information \| Similar Motifs Found | motif file (matrix) |
| 15 * | 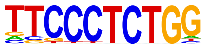 | 1e-7 | -1.830e+01 | 7.89% | 1.18% | 64.1bp (62.0bp) | NEUROG2/MA1642.2/ Jaspar(0.719) More Information \| Similar Motifs Found | motif file (matrix) |
| 16 * | 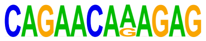 | 1e-6 | -1.553e+01 | 2.11% | 0.02% | 53.6bp (74.3bp) | Sox11/MA0869.3/ Jaspar(0.790) More Information \| Similar Motifs Found | motif file (matrix) |
| 17 * | 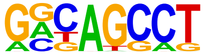 | 1e-6 | -1.392e+01 | 18.42% | 7.58% | 57.7bp (64.7bp) | POL010.1_DCE_S_III/ Jaspar(0.740) More Information \| Similar Motifs Found | motif file (matrix) |

### Motifs found for: solid tumor

### Homer *de novo* Motif Results (homerpos8/)

Non-redundant Motif File of Results
Known Motif Enrichment Results
Gene Ontology Enrichment Results
If Homer is having trouble matching a motif to a known motif, try copy/pasting the matrix file into STAMP
More information on motif finding results: HOMER | Description of Results | Tips
Total target sequences = 209
Total background sequences = 96813
* - possible false positive

| Rank | Motif | P-value | log P-pvalue | % of Targets | % of Background | STD(Bg STD) | Best Match/Details | Motif File |
|---|---|---|---|---|---|---|---|---|
| 1 | 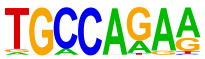 | 1e-12 | -2.948e+01 | 25.84% | 8.60% | 59.2bp (69.4bp) | NFIX/MA0671.2/ Jaspar(0.827) More Information \| Similar Motifs Found | motif file (matrix) |
| 2 * | 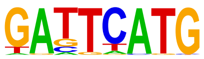 | 1e-11 | -2.724e+01 | 46.89% | 24.41% | 52.2bp (67.0bp) | MEIS1/MA1639.2/ Jaspar(0.836) More Information \| Similar Motifs Found | motif file (matrix) |
| 3 * | 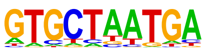 | 1e-11 | -2.563e+01 | 23.44% | 8.01% | 53.8bp (67.2bp) | MGA::EVX1/ MA1960.2/ Jaspar(0.789) More Information \| Similar Motifs Found | motif file (matrix) |
| 4 * | | 1e-10 | -2.497e+01 | 2.87% | 0.02% | 56.7bp (76.6bp) | PH0107.1_Msx2/ Jaspar(0.732) | motif file (matrix) |

| | | | | | | | | |
|---|---|---|---|---|---|---|---|---|
| | 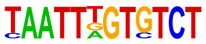 | | | | | | More Information \| Similar Motifs Found | |
| 5 * | 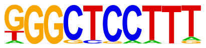 | 1e-10 | -2.405e+01 | 6.70% | 0.58% | 56.3bp (65.1bp) | PB0137.1_Irf3_2/ Jaspar(0.661) More Information \| Similar Motifs Found | motif file (matrix) |
| 6 * | 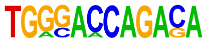 | 1e-10 | -2.398e+01 | 2.87% | 0.03% | 39.6bp (49.9bp) | Smad2(MAD)/ES-SMAD2-ChIP-Seq(GSE29422)/ Homer(0.692) More Information \| Similar Motifs Found | motif file (matrix) |
| 7 * | 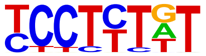 | 1e-10 | -2.380e+01 | 38.28% | 18.84% | 56.9bp (64.3bp) | PB0166.1_Sox12_2/ Jaspar(0.776) More Information \| Similar Motifs Found | motif file (matrix) |
| 8 * | 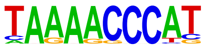 | 1e-10 | -2.320e+01 | 7.18% | 0.74% | 63.5bp (61.4bp) | PB0098.1_Zfp410_1/ Jaspar(0.622) More Information \| Similar Motifs Found | motif file (matrix) |
| 9 * | 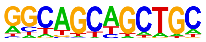 | 1e-9 | -2.228e+01 | 14.35% | 3.65% | 46.9bp (78.4bp) | MyoD(bHLH)/ Myotube-MyoD-ChIP-Seq(GSE21614)/ Homer(0.851) More Information \| Similar Motifs Found | motif file (matrix) |
| 10 * | 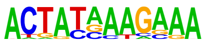 | 1e-9 | -2.186e+01 | 5.26% | 0.35% | 45.0bp (63.0bp) | PB0019.1_Foxl1_1/ Jaspar(0.633) More Information \| Similar Motifs Found | motif file (matrix) |
| 11 * | | 1e-9 | -2.157e+01 | 2.87% | 0.04% | 32.0bp (66.0bp) | Sox11/MA0869.3/ Jaspar(0.670) | motif file (matrix) |

| | | | | | | | | |
|---|---|---|---|---|---|---|---|---|
| | 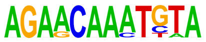 | | | | | | More Information \| Similar Motifs Found | |
| 12 * | 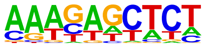 | 1e-8 | -1.879e+01 | 8.61% | 1.55% | 49.9bp (68.3bp) | POU6F2/MA0793.2/ Jaspar(0.630) More Information \| Similar Motifs Found | motif file (matrix) |
| 13 * | 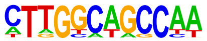 | 1e-8 | -1.846e+01 | 4.31% | 0.28% | 53.7bp (67.4bp) | NFIC/MA0161.3/ Jaspar(0.727) More Information \| Similar Motifs Found | motif file (matrix) |
| 14 * | 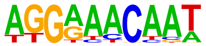 | 1e-7 | -1.775e+01 | 10.05% | 2.28% | 59.5bp (65.7bp) | SRY/MA0084.2/ Jaspar(0.777) More Information \| Similar Motifs Found | motif file (matrix) |
| 15 * | 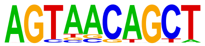 | 1e-7 | -1.721e+01 | 6.22% | 0.84% | 48.5bp (67.2bp) | FERD3L/MA1485.1/ Jaspar(0.761) More Information \| Similar Motifs Found | motif file (matrix) |
| 16 * | 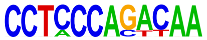 | 1e-7 | -1.695e+01 | 1.91% | 0.02% | 34.4bp (85.7bp) | SP5/MA1965.2/ Jaspar(0.729) More Information \| Similar Motifs Found | motif file (matrix) |
| 17 * | 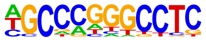 | 1e-6 | -1.595e+01 | 6.22% | 0.93% | 43.1bp (70.2bp) | Zfx/MA0146.3/ Jaspar(0.809) More Information \| Similar Motifs Found | motif file (matrix) |
| 18 * | | 1e-5 | -1.369e+01 | 2.39% | 0.08% | 53.2bp (67.2bp) | TFAP2A/MA0810.2/ Jaspar(0.696) | motif file (matrix) |

| | | | | | | | | |
|---|---|---|---|---|---|---|---|---|
| | 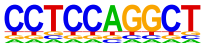 | | | | | | More Information \| Similar Motifs Found | |
| 19 * | 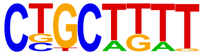 | 1e-5 | -1.289e+01 | 32.06% | 18.66% | 58.9bp (69.1bp) | ZNF317/MA1593.2/ Jaspar(0.746) More Information \| Similar Motifs Found | motif file (matrix) |
| 20 * | 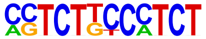 | 1e-5 | -1.172e+01 | 2.39% | 0.13% | 47.1bp (60.4bp) | ZNF257/MA1710.2/ Jaspar(0.719) More Information \| Similar Motifs Found | motif file (matrix) |
| 21 * | 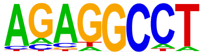 | 1e-4 | -1.048e+01 | 18.18% | 9.06% | 54.8bp (68.9bp) | ZFX(Zf)/mES-Zfx-ChIP-Seq(GSE11431)/ Homer(0.865) More Information \| Similar Motifs Found | motif file (matrix) |

### Motifs found for: subarachnoid invasion

### Homer *de novo* Motif Results (homerpos9/)

Non-redundant Motif File of Results
Known Motif Enrichment Results
Gene Ontology Enrichment Results
If Homer is having trouble matching a motif to a known motif, try copy/pasting the matrix file into STAMP
More information on motif finding results: HOMER | Description of Results | Tips
Total target sequences = 914
Total background sequences = 95666
* - possible false positive

| Rank | Motif | P-value | log P-pvalue | % of Targets | % of Background | STD(Bg STD) | Best Match/Details | Motif File |
|---|---|---|---|---|---|---|---|---|
| 1 | 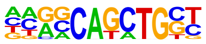 | 1e-46 | -1.060e+02 | 25.71% | 9.34% | 54.6bp (68.1bp) | Atoh1(bHLH)/ Cerebellum-Atoh1-ChIP-Seq(GSE22111)/ Homer(0.975) More Information \| Similar Motifs Found | motif file (matrix) |
| 2 | 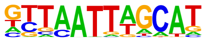 | 1e-35 | -8.277e+01 | 13.13% | 3.33% | 55.4bp (67.8bp) | Oct6(POU,Homeobox)/ NPC-Pou3f1-ChIP-Seq(GSE35496)/ Homer(0.904) More Information \| Similar Motifs Found | motif file (matrix) |
| 3 | 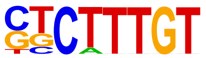 | 1e-34 | -7.838e+01 | 31.73% | 15.44% | 55.2bp (66.7bp) | SOX10/MA0442.3/ Jaspar(0.958) More Information \| Similar Motifs Found | motif file (matrix) |
| 4 | | 1e-23 | -5.368e+01 | 4.70% | 0.61% | 61.2bp (63.9bp) | Rfx1(HTH)/NPC-H3K4me1-ChIP-Seq(GSE16256)/ | motif file (matrix) |

| | | | | | | | | |
|---|---|---|---|---|---|---|---|---|
| | 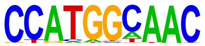 | | | | | | Homer(0.907) More Information \| Similar Motifs Found | |
| 5 | 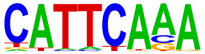 | 1e-19 | -4.456e+01 | 35.12% | 21.89% | 56.6bp (66.3bp) | LIN54/MA0619.2/ Jaspar(0.882) More Information \| Similar Motifs Found | motif file (matrix) |
| 6 | 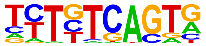 | 1e-18 | -4.340e+01 | 26.91% | 15.29% | 54.9bp (64.5bp) | MEIS1/MA0498.3/ Jaspar(0.729) More Information \| Similar Motifs Found | motif file (matrix) |
| 7 | 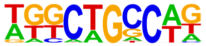 | 1e-18 | -4.230e+01 | 21.12% | 10.94% | 54.9bp (69.7bp) | ZNF549/MA1728.2/ Jaspar(0.716) More Information \| Similar Motifs Found | motif file (matrix) |
| 8 | 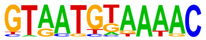 | 1e-13 | -3.108e+01 | 1.97% | 0.16% | 54.2bp (60.5bp) | Nfat5/MA0606.3/ Jaspar(0.711) More Information \| Similar Motifs Found | motif file (matrix) |
| 9 | 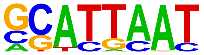 | 1e-13 | -3.028e+01 | 18.38% | 10.21% | 58.1bp (68.0bp) | Arid3b/MA0601.2/ Jaspar(0.849) More Information \| Similar Motifs Found | motif file (matrix) |
| 10 | 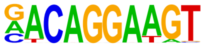 | 1e-12 | -2.933e+01 | 5.14% | 1.43% | 54.9bp (63.7bp) | Ets1-distal(ETS)/CD4+-PolII-ChIP-Seq(Barski_et_al.)/ Homer(0.979) More Information \| Similar Motifs Found | motif file (matrix) |
| 11 | | 1e-12 | -2.923e+01 | 5.14% | 1.44% | 54.3bp (67.7bp) | ZKSCAN3/MA1973.2/ Jaspar(0.717) | motif file (matrix) |

| | | | | | | | | |
|---|---|---|---|---|---|---|---|---|
| | 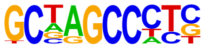 | | | | | | More Information \| Similar Motifs Found | |
| 12 * | 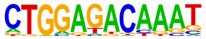 | 1e-11 | -2.678e+01 | 3.17% | 0.62% | 56.3bp (61.2bp) | ZBTB26/MA1579.2/ Jaspar(0.634) More Information \| Similar Motifs Found | motif file (matrix) |
| 13 * | 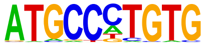 | 1e-10 | -2.500e+01 | 3.28% | 0.71% | 57.1bp (68.4bp) | THAP1/MA0597.3/ Jaspar(0.730) More Information \| Similar Motifs Found | motif file (matrix) |
| 14 * | 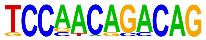 | 1e-10 | -2.444e+01 | 0.88% | 0.02% | 59.7bp (58.4bp) | AMYB(HTH)/Testes-AMYB-ChIP-Seq(GSE44588)/ Homer(0.658) More Information \| Similar Motifs Found | motif file (matrix) |
| 15 * | 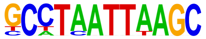 | 1e-10 | -2.441e+01 | 0.77% | 0.01% | 56.3bp (66.1bp) | ARGFX/MA1463.2/ Jaspar(0.784) More Information \| Similar Motifs Found | motif file (matrix) |
| 16 * | 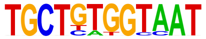 | 1e-9 | -2.300e+01 | 0.98% | 0.04% | 46.9bp (59.6bp) | RUNX-AML(Runt)/ CD4+-PolII-ChIP-Seq(Barski_et_al.)/ Homer(0.689) More Information \| Similar Motifs Found | motif file (matrix) |
| 17 * | 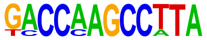 | 1e-9 | -2.288e+01 | 0.66% | 0.01% | 58.8bp (35.6bp) | FEZF2/MA2341.1/ Jaspar(0.709) More Information \| Similar Motifs Found | motif file (matrix) |
| 18 * | | 1e-8 | -1.936e+01 | 12.47% | 7.06% | 52.7bp (57.9bp) | ZNF148/MA1653.2/ Jaspar(0.757) | motif file (matrix) |

| | | | | | | | | |
|---|---|---|---|---|---|---|---|---|
| | 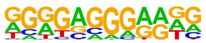 | | | | | | More Information \| Similar Motifs Found | |
| 19 * | 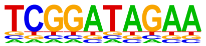 | 1e-6 | -1.575e+01 | 0.33% | 0.00% | 24.7bp (0.0bp) | ZNF35/MA2333.1/ Jaspar(0.605) More Information \| Similar Motifs Found | motif file (matrix) |
| 20 * | 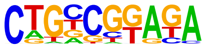 | 1e-5 | -1.237e+01 | 3.94% | 1.69% | 57.7bp (61.6bp) | ERF::NHLH1/ MA1938.2/ Jaspar(0.597) More Information \| Similar Motifs Found | motif file (matrix) |

### Motifs found for: white matter invasion

### Homer *de novo* Motif Results (homerpos10/)

Non-redundant Motif File of Results
Known Motif Enrichment Results
Gene Ontology Enrichment Results
If Homer is having trouble matching a motif to a known motif, try copy/pasting the matrix file into STAMP
More information on motif finding results: HOMER | Description of Results | Tips
Total target sequences = 598
Total background sequences = 96320
* - possible false positive

| Rank | Motif | P-value | log P-pvalue | % of Targets | % of Background | STD(Bg STD) | Best Match/Details | Motif File |
|---|---|---|---|---|---|---|---|---|
| 1 | 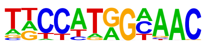 | 1e-50 | -1.174e+02 | 13.21% | 1.34% | 53.8bp (63.2bp) | X-box(HTH)/NPC-H3K4me1-ChIP-Seq(GSE16256)/Homer(0.897) More Information \| Similar Motifs Found | motif file (matrix) |
| 2 | 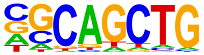 | 1e-24 | -5.639e+01 | 25.25% | 10.34% | 55.2bp (70.9bp) | MYOG/MA0500.3/Jaspar(0.970) More Information \| Similar Motifs Found | motif file (matrix) |
| 3 | 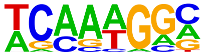 | 1e-22 | -5.223e+01 | 49.16% | 29.75% | 54.1bp (65.2bp) | Sox3(HMG)/NPC-Sox3-ChIP-Seq(GSE33059)/Homer(0.834) More Information \| Similar Motifs Found | motif file (matrix) |
| 4 | | 1e-20 | -4.692e+01 | 14.55% | 4.58% | 54.3bp (65.1bp) | POU6F1/MA1549.2/Jaspar(0.836) | motif file (matrix) |

| | | | | | | | | |
|---|---|---|---|---|---|---|---|---|
| | 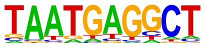 | | | | | | More Information \| Similar Motifs Found | |
| 5 | 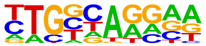 | 1e-19 | -4.482e+01 | 25.59% | 11.90% | 56.3bp (66.4bp) | NF1-halfsite(CTF)/ LNCaP-NF1-ChIP-Seq(Unpublished)/ Homer(0.854) More Information \| Similar Motifs Found | motif file (matrix) |
| 6 | 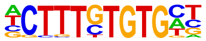 | 1e-16 | -3.749e+01 | 6.35% | 1.14% | 49.7bp (63.3bp) | PB0208.1_Zscan4_2/ Jaspar(0.788) More Information \| Similar Motifs Found | motif file (matrix) |
| 7 | 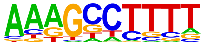 | 1e-15 | -3.550e+01 | 45.48% | 29.76% | 55.6bp (66.0bp) | MYNN(Zf)/HEK293-MYNN.eGFP-ChIP-Seq(Encode)/ Homer(0.600) More Information \| Similar Motifs Found | motif file (matrix) |
| 8 | 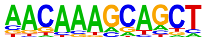 | 1e-15 | -3.467e+01 | 18.23% | 8.02% | 53.3bp (65.9bp) | Sox11/MA0869.3/ Jaspar(0.746) More Information \| Similar Motifs Found | motif file (matrix) |
| 9 | 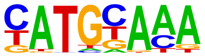 | 1e-13 | -3.217e+01 | 19.40% | 9.15% | 50.1bp (64.6bp) | POU5F1/MA1115.2/ Jaspar(0.944) More Information \| Similar Motifs Found | motif file (matrix) |
| 10 | 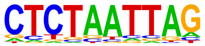 | 1e-13 | -3.028e+01 | 12.88% | 5.01% | 49.5bp (63.4bp) | LHX9(Homeobox)/ Hct116-LHX9.V5-ChIP-Seq(GSE116822)/ Homer(0.872) More Information \| Similar Motifs Found | motif file (matrix) |
| 11 | | 1e-13 | -3.014e+01 | 13.04% | 5.13% | 51.9bp (62.5bp) | MZF1/MA0056.3/ Jaspar(0.822) | motif file (matrix) |

| | | | | | | | | |
|---|---|---|---|---|---|---|---|---|
| | 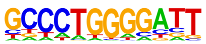  | | | | | | More Information \| Similar Motifs Found | |
| 12 | 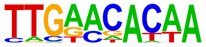  | 1e-12 | -2.899e+01 | 6.19% | 1.43% | 54.5bp (65.1bp) | Oct4:Sox17(POU,Homeobox,HMG)/F9-Sox17-ChIP-Seq(GSE44553)/Homer(0.706) More Information \| Similar Motifs Found | motif file (matrix) |
| 13 * | 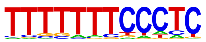  | 1e-11 | -2.741e+01 | 10.87% | 4.05% | 56.4bp (64.7bp) | ZNF384/MA1125.2/Jaspar(0.768) More Information \| Similar Motifs Found | motif file (matrix) |
| 14 * | 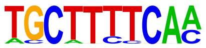  | 1e-11 | -2.732e+01 | 5.02% | 1.00% | 60.2bp (66.5bp) | PB0022.1_Gata5_1/Jaspar(0.639) More Information \| Similar Motifs Found | motif file (matrix) |
| 15 * | 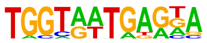  | 1e-11 | -2.650e+01 | 4.52% | 0.83% | 53.6bp (60.6bp) | Hoxa10(Homeobox)/ChickenMSG-Hoxa10.Flag-ChIP-Seq(GSE86088)/Homer(0.689) More Information \| Similar Motifs Found | motif file (matrix) |
| 16 * | 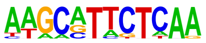  | 1e-10 | -2.421e+01 | 2.17% | 0.16% | 58.0bp (63.7bp) | PRDM1/MA0508.4/Jaspar(0.613) More Information \| Similar Motifs Found | motif file (matrix) |
| 17 * | 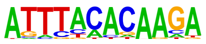  | 1e-10 | -2.347e+01 | 4.35% | 0.88% | 54.0bp (63.6bp) | CEBPD/MA0836.3/Jaspar(0.695) More Information \| Similar Motifs Found | motif file (matrix) |
| 18 * | | 1e-9 | -2.227e+01 | 2.84% | 0.37% | 47.7bp (61.4bp) | SOX15/MA1152.2/Jaspar(0.792) | motif file (matrix) |

| | | | | | | | | |
|---|---|---|---|---|---|---|---|---|
| | 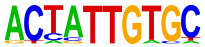 | | | | | | More Information \| Similar Motifs Found | |
| 19 * | 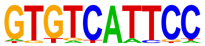 | 1e-9 | -2.086e+01 | 3.18% | 0.52% | 43.2bp (64.9bp) | PH0164.1_Six4/ Jaspar(0.756) More Information \| Similar Motifs Found | motif file (matrix) |
| 20 * | 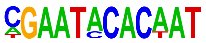 | 1e-9 | -2.075e+01 | 0.67% | 0.00% | 48.9bp (40.5bp) | Foxh1(Forkhead)/hESC-FOXH1-ChIP-Seq(GSE29422)/ Homer(0.798) More Information \| Similar Motifs Found | motif file (matrix) |
| 21 * | 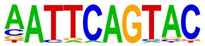 | 1e-8 | -1.994e+01 | 3.34% | 0.62% | 52.6bp (64.3bp) | PB0068.1_Sox1_1/ Jaspar(0.748) More Information \| Similar Motifs Found | motif file (matrix) |
| 22 * | 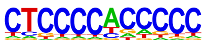 | 1e-8 | -1.963e+01 | 15.55% | 8.23% | 56.8bp (57.9bp) | ZNF281/MA1630.3/ Jaspar(0.843) More Information \| Similar Motifs Found | motif file (matrix) |
| 23 * | 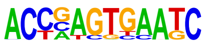 | 1e-8 | -1.914e+01 | 1.34% | 0.06% | 53.0bp (57.7bp) | DUX(Homeobox)/ C2C12-Dux-ChIP-Seq(GSE87279)/ Homer(0.586) More Information \| Similar Motifs Found | motif file (matrix) |
| 24 * | 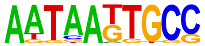 | 1e-6 | -1.601e+01 | 4.18% | 1.19% | 60.8bp (60.8bp) | MF0010.1_Homeobox_ class/Jaspar(0.774) More Information \| Similar Motifs Found | motif file (matrix) |
| 25 * | | 1e-6 | -1.500e+01 | 1.84% | 0.24% | 52.7bp (59.8bp) | PGR/MA2327.1/ Jaspar(0.729) | motif file (matrix) |

| | | | | | | | | |
|---|---|---|---|---|---|---|---|---|
| | 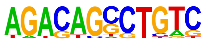 | | | | | | More Information \| Similar Motifs Found | |
| 26 * | 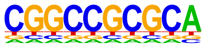 | 1e-6 | -1.398e+01 | 0.67% | 0.01% | 40.4bp (53.9bp) | ZBTB14/MA1650.2/Jaspar(0.791) More Information \| Similar Motifs Found | motif file (matrix) |

### Motifs found for: age at diagnosis

### Homer *de novo* Motif Results (homerpos11/)

Non-redundant Motif File of Results
Known Motif Enrichment Results
Gene Ontology Enrichment Results
If Homer is having trouble matching a motif to a known motif, try copy/pasting the matrix file into STAMP
More information on motif finding results: HOMER | Description of Results | Tips
Total target sequences = 350
Total background sequences = 97496
* - possible false positive

| Rank | Motif | P-value | log P-pvalue | % of Targets | % of Background | STD(Bg STD) | Best Match/Details | Motif File |
|---|---|---|---|---|---|---|---|---|
| 1 | 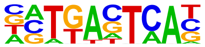 | 1e-51 | -1.192e+02 | 32.00% | 5.65% | 51.3bp (64.0bp) | Fra1(bZIP)/BT549-Fra1-ChIP-Seq(GSE46166)/Homer(0.973) More Information \| Similar Motifs Found | motif file (matrix) |
| 2 | 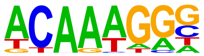 | 1e-20 | -4.799e+01 | 53.14% | 28.81% | 52.8bp (64.2bp) | Sox3(HMG)/NPC-Sox3-ChIP-Seq(GSE33059)/Homer(0.947) More Information \| Similar Motifs Found | motif file (matrix) |
| 3 | 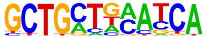 | 1e-14 | -3.401e+01 | 7.14% | 0.86% | 53.4bp (69.9bp) | MAF/MA1520.2/Jaspar(0.708) More Information \| Similar Motifs Found | motif file (matrix) |
| 4 | | 1e-14 | -3.228e+01 | 4.57% | 0.28% | 55.7bp (58.9bp) | Sox5/MA0087.3/Jaspar(0.693) | motif file (matrix) |

| | | | | | | | | |
|---|---|---|---|---|---|---|---|---|
| | 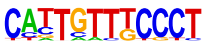 | | | | | | More Information \| Similar Motifs Found | |
| 5 | 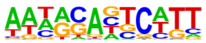 | 1e-13 | -3.106e+01 | 35.14% | 18.15% | 55.3bp (63.4bp) | PB0028.1_Hbp1_1/ Jaspar(0.599) More Information \| Similar Motifs Found | motif file (matrix) |
| 6 | 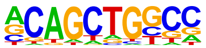 | 1e-12 | -2.872e+01 | 24.00% | 10.46% | 53.9bp (67.8bp) | Tcf21(bHLH)/ ArterySmoothMuscle-Tcf21-ChIP-Seq(GSE61369)/ Homer(0.915) More Information \| Similar Motifs Found | motif file (matrix) |
| 7 | 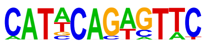 | 1e-12 | -2.859e+01 | 2.86% | 0.08% | 51.7bp (59.6bp) | Dmrt1/MA1603.2/ Jaspar(0.606) More Information \| Similar Motifs Found | motif file (matrix) |
| 8 | 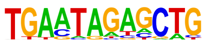 | 1e-12 | -2.791e+01 | 7.43% | 1.24% | 47.6bp (64.9bp) | PB0178.1_Sox8_2/ Jaspar(0.586) More Information \| Similar Motifs Found | motif file (matrix) |
| 9 * | 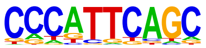 | 1e-11 | -2.615e+01 | 12.86% | 3.92% | 56.9bp (64.6bp) | PB0170.1_Sox17_2/ Jaspar(0.694) More Information \| Similar Motifs Found | motif file (matrix) |
| 10 * | 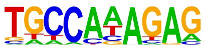 | 1e-10 | -2.433e+01 | 17.14% | 6.71% | 49.1bp (64.1bp) | ZNF768/MA1731.2/ Jaspar(0.696) More Information \| Similar Motifs Found | motif file (matrix) |
| 11 * | | 1e-10 | -2.404e+01 | 9.14% | 2.24% | 49.6bp (63.5bp) | ELF3/MA0640.3/ Jaspar(0.875) | motif file (matrix) |

| | | | | | | | | |
|---|---|---|---|---|---|---|---|---|
| | 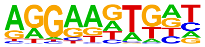 | | | | | | More Information \| Similar Motifs Found | |
| 12 * | 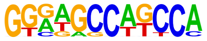 | 1e-9 | -2.242e+01 | 3.71% | 0.32% | 45.8bp (66.0bp) | Tlx?(NR)/NPC-H3K4me1-ChIP-Seq(GSE16256)/Homer(0.789) More Information \| Similar Motifs Found | motif file (matrix) |
| 13 * | 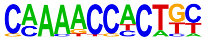 | 1e-8 | -1.985e+01 | 3.43% | 0.32% | 61.9bp (64.2bp) | RUNX1(Runt)/Jurkat-RUNX1-ChIP-Seq(GSE29180)/Homer(0.724) More Information \| Similar Motifs Found | motif file (matrix) |
| 14 * | 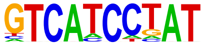 | 1e-8 | -1.942e+01 | 2.86% | 0.20% | 39.5bp (57.8bp) | PB0098.1_Zfp410_1/Jaspar(0.702) More Information \| Similar Motifs Found | motif file (matrix) |
| 15 * | 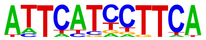 | 1e-8 | -1.884e+01 | 6.29% | 1.38% | 51.0bp (62.7bp) | Pit1+1bp(Homeobox)/GCrat-Pit1-ChIP-Seq(GSE58009)/Homer(0.676) More Information \| Similar Motifs Found | motif file (matrix) |
| 16 * | 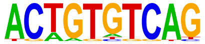 | 1e-7 | -1.818e+01 | 6.00% | 1.31% | 52.7bp (65.1bp) | MEIS1/MA0498.3/Jaspar(0.709) More Information \| Similar Motifs Found | motif file (matrix) |
| 17 * | 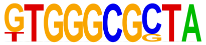 | 1e-7 | -1.695e+01 | 1.14% | 0.01% | 49.5bp (30.0bp) | Zfp961/MA2126.1/Jaspar(0.740) More Information \| Similar Motifs Found | motif file (matrix) |
| 18 * | | 1e-6 | -1.607e+01 | 2.86% | 0.29% | 35.2bp (65.6bp) | Vdr/MA0693.4/Jaspar(0.729) | motif file (matrix) |

| | | | | | | | | |
|---|---|---|---|---|---|---|---|---|
| | 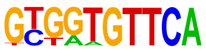 | | | | | | More Information \| Similar Motifs Found | |
| 19 * | 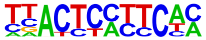 | 1e-6 | -1.534e+01 | 8.00% | 2.60% | 52.9bp (64.1bp) | PB0203.1_Zfp691_2/ Jaspar(0.649) More Information \| Similar Motifs Found | motif file (matrix) |
| 20 * | 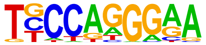 | 1e-6 | -1.465e+01 | 19.43% | 10.46% | 54.4bp (64.3bp) | EBF(EBF)/proBcell-EBF-ChIP-Seq(GSE21978)/ Homer(0.753) More Information \| Similar Motifs Found | motif file (matrix) |
| 21 * | 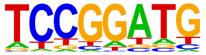 | 1e-5 | -1.215e+01 | 15.71% | 8.38% | 52.6bp (63.2bp) | PB0077.1_Spdef_1/ Jaspar(0.869) More Information \| Similar Motifs Found | motif file (matrix) |

#### Motifs found for: survival time

### Homer *de novo* Motif Results (homerpos12/)

Non-redundant Motif File of Results
Known Motif Enrichment Results
Gene Ontology Enrichment Results
If Homer is having trouble matching a motif to a known motif, try copy/pasting the matrix file into STAMP
More information on motif finding results: HOMER | Description of Results | Tips
Total target sequences = 298
Total background sequences = 97964
* - possible false positive

| Rank | Motif | P-value | log P-pvalue | % of Targets | % of Background | STD(Bg STD) | Best Match/Details | Motif File |
|---|---|---|---|---|---|---|---|---|
| 1 | 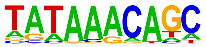 | 1e-13 | -3.015e+01 | 17.79% | 5.56% | 58.5bp (61.4bp) | Foxf1(Forkhead)/Lung-Foxf1-ChIP-Seq(GSE77951)/Homer(0.875) More Information \| Similar Motifs Found | motif file (matrix) |
| 2 * | 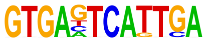 | 1e-11 | -2.652e+01 | 2.01% | 0.01% | 54.0bp (54.3bp) | FOS/MA0476.2/Jaspar(0.791) More Information \| Similar Motifs Found | motif file (matrix) |
| 3 * | 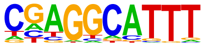 | 1e-11 | -2.580e+01 | 18.79% | 6.85% | 57.1bp (61.2bp) | PB0149.1_Myb_2/Jaspar(0.656) More Information \| Similar Motifs Found | motif file (matrix) |
| 4 * | | 1e-11 | -2.563e+01 | 3.02% | 0.08% | 44.2bp (65.4bp) | NFIA/MA0670.2/Jaspar(0.671) | motif file (matrix) |

| | | | | | | | | |
|---|---|---|---|---|---|---|---|---|
| | 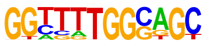 | | | | | | More Information \| Similar Motifs Found | |
| 5 * | 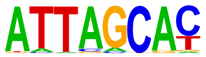 | 1e-11 | -2.543e+01 | 16.78% | 5.70% | 53.6bp (62.7bp) | MGA::EVX1/ MA1960.2/ Jaspar(0.821) More Information \| Similar Motifs Found | motif file (matrix) |
| 6 * | 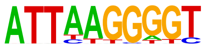 | 1e-11 | -2.539e+01 | 4.36% | 0.29% | 55.4bp (56.5bp) | ZNF410(Zf)/CD34-ZNF410-ChIP-Seq(GSE154960)/ Homer(0.685) More Information \| Similar Motifs Found | motif file (matrix) |
| 7 * | 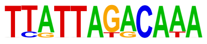 | 1e-10 | -2.410e+01 | 2.01% | 0.02% | 52.8bp (65.8bp) | SATB1/MA1963.2/ Jaspar(0.753) More Information \| Similar Motifs Found | motif file (matrix) |
| 8 * | 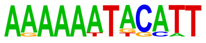 | 1e-10 | -2.395e+01 | 3.69% | 0.20% | 44.1bp (56.0bp) | ZNF384/MA1125.2/ Jaspar(0.725) More Information \| Similar Motifs Found | motif file (matrix) |
| 9 * | 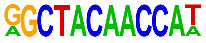 | 1e-10 | -2.361e+01 | 1.34% | 0.00% | 15.0bp (14.8bp) | ZNF418/MA1980.1/ Jaspar(0.707) More Information \| Similar Motifs Found | motif file (matrix) |
| 10 * | 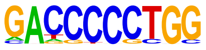 | 1e-10 | -2.323e+01 | 3.69% | 0.21% | 61.9bp (56.7bp) | PB0025.1_Glis2_1/ Jaspar(0.713) More Information \| Similar Motifs Found | motif file (matrix) |
| 11 * | | 1e-10 | -2.317e+01 | 4.03% | 0.28% | 65.3bp (63.8bp) | CEBPD/MA0836.3/ Jaspar(0.790) | motif file (matrix) |

| | | | | | | | | |
|---|---|---|---|---|---|---|---|---|
| | 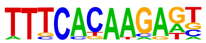 | | | | | | More Information \| Similar Motifs Found | |
| 12 * | 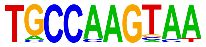 | 1e-9 | -2.257e+01 | 4.36% | 0.37% | 48.8bp (61.6bp) | NFIC/MA0161.3/ Jaspar(0.862) More Information \| Similar Motifs Found | motif file (matrix) |
| 13 * | 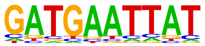 | 1e-9 | -2.186e+01 | 10.74% | 2.90% | 52.7bp (60.2bp) | POU4F3/MA0791.2/ Jaspar(0.764) More Information \| Similar Motifs Found | motif file (matrix) |
| 14 * | 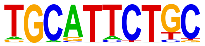 | 1e-9 | -2.185e+01 | 5.70% | 0.77% | 52.2bp (65.0bp) | OCT:OCT-short(POU,Homeobox)/ NPC-OCT6-ChIP-Seq(GSE43916)/ Homer(0.680) More Information \| Similar Motifs Found | motif file (matrix) |
| 15 * | 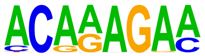 | 1e-9 | -2.180e+01 | 33.22% | 18.15% | 52.2bp (60.9bp) | SOX10/MA0442.3/ Jaspar(0.846) More Information \| Similar Motifs Found | motif file (matrix) |
| 16 * | 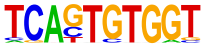 | 1e-9 | -2.150e+01 | 6.04% | 0.90% | 58.9bp (58.2bp) | RUNX2(Runt)/PCa-RUNX2-ChIP-Seq(GSE33889)/ Homer(0.770) More Information \| Similar Motifs Found | motif file (matrix) |
| 17 * | 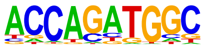 | 1e-9 | -2.079e+01 | 16.78% | 6.52% | 55.3bp (61.5bp) | NeuroD1(bHLH)/Islet-NeuroD1-ChIP-Seq(GSE30298)/ Homer(0.896) More Information \| Similar Motifs Found | motif file (matrix) |
| 18 * | | 1e-8 | -2.026e+01 | 5.70% | 0.86% | 57.5bp (61.7bp) | Brn2(POU,Homeobox)/ NPC-Brn2-ChIP- | motif file (matrix) |

| | | | | | | | | |
|---|---|---|---|---|---|---|---|---|
| | 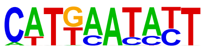 | | | | | | Seq(GSE35496)/ Homer(0.811) More Information \| Similar Motifs Found | |
| 19 * | 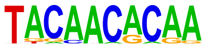 | 1e-8 | -1.960e+01 | 5.03% | 0.67% | 52.8bp (61.6bp) | ZIM3/MA1709.2/ Jaspar(0.617) More Information \| Similar Motifs Found | motif file (matrix) |
| 20 * | 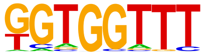 | 1e-8 | -1.892e+01 | 23.49% | 11.57% | 57.6bp (61.6bp) | RUNX1(Runt)/Jurkat-RUNX1-ChIP-Seq(GSE29180)/ Homer(0.886) More Information \| Similar Motifs Found | motif file (matrix) |
| 21 * | 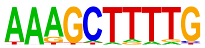 | 1e-8 | -1.887e+01 | 6.38% | 1.20% | 55.7bp (62.7bp) | Nr2e3/MA0164.2/ Jaspar(0.771) More Information \| Similar Motifs Found | motif file (matrix) |
| 22 * | 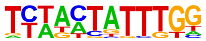 | 1e-7 | -1.820e+01 | 2.35% | 0.09% | 37.8bp (58.4bp) | Mesp1(bHLH)/ESC-Mesp1-ChIP-Seq(GSE165102)/ Homer(0.660) More Information \| Similar Motifs Found | motif file (matrix) |
| 23 * | 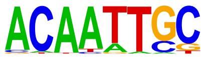 | 1e-7 | -1.766e+01 | 11.41% | 3.84% | 63.6bp (65.1bp) | PB0165.1_Sox11_2/ Jaspar(0.812) More Information \| Similar Motifs Found | motif file (matrix) |
| 24 * | 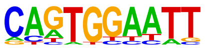 | 1e-7 | -1.633e+01 | 13.76% | 5.51% | 52.8bp (62.3bp) | TEAD3(TEA)/HepG2-TEAD3-ChIP-Seq(Encode)/ Homer(0.738) More Information \| Similar Motifs Found | motif file (matrix) |
| 25 * | | 1e-5 | -1.247e+01 | 3.02% | 0.40% | 60.4bp (58.8bp) | PB0119.1_Foxa2_2/ Jaspar(0.739) | motif file (matrix) |

[More Information](#) |
[Similar Motifs Found](#)

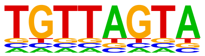

### Motifs found for: sex

### Homer *de novo* Motif Results (homerpos13/)

Non-redundant Motif File of Results
Known Motif Enrichment Results
Gene Ontology Enrichment Results
If Homer is having trouble matching a motif to a known motif, try copy/pasting the matrix file into STAMP
More information on motif finding results: HOMER | Description of Results | Tips
Total target sequences = 123
Total background sequences = 98265
* - possible false positive

| Rank | Motif | P-value | log P-pvalue | % of Targets | % of Background | STD(Bg STD) | Best Match/Details | Motif File |
|---|---|---|---|---|---|---|---|---|
| 1 | 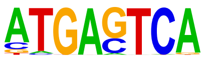  | 1e-16 | -3.864e+01 | 28.46% | 4.94% | 54.4bp (62.6bp) | Jun/MA0489.3/ Jaspar(0.993) More Information \| Similar Motifs Found | motif file (matrix) |
| 2 | 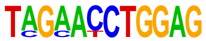  | 1e-15 | -3.605e+01 | 4.88% | 0.01% | 50.5bp (51.3bp) | Prdm15/MA1616.2/ Jaspar(0.701) More Information \| Similar Motifs Found | motif file (matrix) |
| 3 | 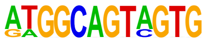  | 1e-15 | -3.481e+01 | 4.07% | 0.00% | 43.0bp (28.0bp) | PB0149.1_Myb_2/ Jaspar(0.635) More Information \| Similar Motifs Found | motif file (matrix) |
| 4 | | 1e-13 | -3.147e+01 | 8.13% | 0.17% | 48.7bp (59.2bp) | CEBP(bZIP)/ThioMac-CEBPb-ChIP-Seq(GSE21512)/ | motif file (matrix) |

| | | | | | | | | |
|---|---|---|---|---|---|---|---|---|
| | 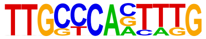 | | | | | | Homer(0.682) More Information \| Similar Motifs Found | |
| 5 | 1e-13 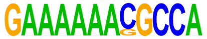 | | -3.135e+01 | 4.07% | 0.00% | 33.6bp (56.7bp) | PB0192.1_Tcfap2e_2/ Jaspar(0.670) More Information \| Similar Motifs Found | motif file (matrix) |
| 6 | 1e-13 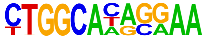 | | -3.061e+01 | 6.50% | 0.07% | 64.6bp (63.4bp) | HIC1(Zf)/Treg-ZBTB29-ChIP-Seq(GSE99889)/ Homer(0.727) More Information \| Similar Motifs Found | motif file (matrix) |
| 7 | 1e-13 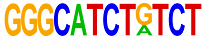 | | -2.996e+01 | 3.25% | 0.00% | 59.6bp (1.6bp) | MYOD1/MA0499.3/ Jaspar(0.693) More Information \| Similar Motifs Found | motif file (matrix) |
| 8 | 1e-12 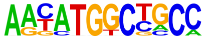 | | -2.983e+01 | 6.50% | 0.08% | 53.4bp (62.7bp) | ZFP42/MA1651.2/ Jaspar(0.910) More Information \| Similar Motifs Found | motif file (matrix) |
| 9 * | 1e-11 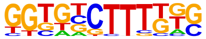 | | -2.742e+01 | 24.39% | 5.25% | 53.1bp (59.8bp) | SOX10/MA0442.3/ Jaspar(0.680) More Information \| Similar Motifs Found | motif file (matrix) |
| 10 * | 1e-11 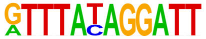 | | -2.718e+01 | 3.25% | 0.00% | 76.9bp (28.6bp) | Hoxa13/MA0650.4/ Jaspar(0.681) More Information \| Similar Motifs Found | motif file (matrix) |
| 11 * | 1e-11 | | -2.585e+01 | 4.88% | 0.03% | 45.8bp (50.9bp) | INSM1/MA0155.1/ Jaspar(0.606) | motif file (matrix) |

| | | | | | | | | |
|---|---|---|---|---|---|---|---|---|
| | 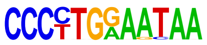 | | | | | | More Information \| Similar Motifs Found | |
| 12 * | 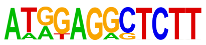 | 1e-10 | -2.485e+01 | 4.88% | 0.04% | 72.4bp (59.4bp) | LRF(Zf)/Erythroblasts-ZBTB7A-ChIP-Seq(GSE74977)/Homer(0.585) More Information \| Similar Motifs Found | motif file (matrix) |
| 13 * | 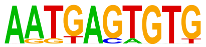 | 1e-10 | -2.441e+01 | 8.94% | 0.47% | 45.1bp (58.9bp) | MSANTD3/MA1523.2/Jaspar(0.698) More Information \| Similar Motifs Found | motif file (matrix) |
| 14 * | 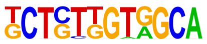 | 1e-10 | -2.336e+01 | 7.32% | 0.27% | 60.2bp (62.6bp) | ZBTB9(Zf)/HEK293T-ZBTB9-ChIP-Seq(GSE251687)/Homer(0.611) More Information \| Similar Motifs Found | motif file (matrix) |
| 15 * | 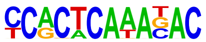 | 1e-9 | -2.268e+01 | 5.69% | 0.11% | 63.6bp (55.6bp) | NKX2-2/MA1645.2/Jaspar(0.754) More Information \| Similar Motifs Found | motif file (matrix) |
| 16 * | 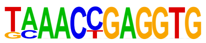 | 1e-9 | -2.218e+01 | 3.25% | 0.01% | 45.3bp (65.2bp) | Tbx5(T-box)/HL1-Tbx5.biotin-ChIP-Seq(GSE21529)/Homer(0.616) More Information \| Similar Motifs Found | motif file (matrix) |
| 17 * | 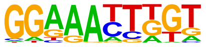 | 1e-9 | -2.175e+01 | 14.63% | 2.23% | 48.9bp (59.3bp) | NFATC3/MA0625.3/Jaspar(0.768) More Information \| Similar Motifs Found | motif file (matrix) |
| 18 * | | 1e-9 | -2.118e+01 | 8.13% | 0.48% | 44.3bp (57.3bp) | KLF15/MA1513.2/Jaspar(0.745) | motif file (matrix) |

| | | | | | | | | |
|---|---|---|---|---|---|---|---|---|
| | 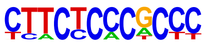 | | | | | | More Information \| Similar Motifs Found | |
| 19 * | 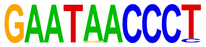 | 1e-8 | -2.058e+01 | 4.88% | 0.08% | 46.1bp (55.8bp) | ZNF766/MA2098.1/ Jaspar(0.702) More Information \| Similar Motifs Found | motif file (matrix) |
| 20 * | 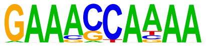 | 1e-8 | -2.037e+01 | 8.94% | 0.70% | 57.5bp (64.4bp) | Bcl11B/MA1989.2/ Jaspar(0.780) More Information \| Similar Motifs Found | motif file (matrix) |
| 21 * | 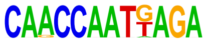 | 1e-8 | -1.864e+01 | 3.25% | 0.02% | 65.1bp (68.6bp) | PH0005.1_Barhl1/ Jaspar(0.753) More Information \| Similar Motifs Found | motif file (matrix) |
| 22 * | 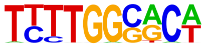 | 1e-7 | -1.751e+01 | 9.76% | 1.16% | 61.4bp (61.6bp) | NFIX/MA0671.2/ Jaspar(0.738) More Information \| Similar Motifs Found | motif file (matrix) |
| 23 * | 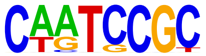 | 1e-2 | -6.712e+00 | 2.44% | 0.17% | 19.3bp (62.5bp) | HINFP(Zf)/K562-HINFP.eGFP-ChIP-Seq(Encode)/ Homer(0.674) More Information \| Similar Motifs Found | motif file (matrix) |
| 24 * | 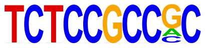 | 1e-1 | -3.826e+00 | 1.63% | 0.18% | 54.8bp (65.4bp) | SP1/MA0079.5/ Jaspar(0.755) More Information \| Similar Motifs Found | motif file (matrix) |

### Motifs found for: tumor initiation

### Homer *de novo* Motif Results (homerpos14/)

[Non-redundant Motif File of Results](#)
[Known Motif Enrichment Results](#)
[Gene Ontology Enrichment Results](#)
If Homer is having trouble matching a motif to a known motif, try copy/pasting the matrix file into STAMP
More information on motif finding results: HOMER | Description of Results | Tips
Total target sequences = 333
Total background sequences = 96869
* - possible false positive

| Rank | Motif | P-value | log P-pvalue | % of Targets | % of Background | STD(Bg STD) | Best Match/Details | Motif File |
|---|---|---|---|---|---|---|---|---|
| 1 | 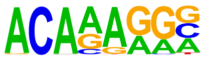 | 1e-19 | -4.483e+01 | 37.84% | 16.77% | 57.1bp (65.5bp) | SOX4/MA0867.3/ Jaspar(0.885) More Information \| Similar Motifs Found | motif file (matrix) |
| 2 | 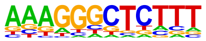 | 1e-14 | -3.440e+01 | 49.25% | 28.50% | 52.3bp (62.0bp) | PB0099.1_Zfp691_1/ Jaspar(0.595) More Information \| Similar Motifs Found | motif file (matrix) |
| 3 | 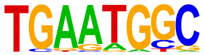 | 1e-13 | -3.008e+01 | 18.02% | 6.15% | 60.6bp (67.2bp) | PB0178.1_Sox8_2/ Jaspar(0.760) More Information \| Similar Motifs Found | motif file (matrix) |
| 4 | | 1e-13 | -3.004e+01 | 7.81% | 1.19% | 58.9bp (62.9bp) | PB0170.1_Sox17_2/ Jaspar(0.605) | motif file (matrix) |

| | | | | | | | | |
|---|---|---|---|---|---|---|---|---|
| | 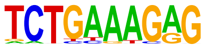 | | | | | | More Information \| Similar Motifs Found | |
| 5 | 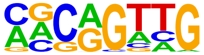 | 1e-13 | -2.999e+01 | 27.33% | 12.24% | 53.7bp (65.4bp) | MYB/MA0100.4/ Jaspar(0.850) More Information \| Similar Motifs Found | motif file (matrix) |
| 6 | 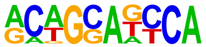 | 1e-12 | -2.895e+01 | 18.62% | 6.68% | 56.5bp (66.4bp) | Nanog/MA2339.1/ Jaspar(0.676) More Information \| Similar Motifs Found | motif file (matrix) |
| 7 * | 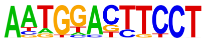 | 1e-10 | -2.440e+01 | 4.50% | 0.42% | 53.9bp (64.8bp) | Elf5/MA0136.4/ Jaspar(0.741) More Information \| Similar Motifs Found | motif file (matrix) |
| 8 * | 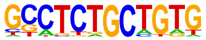 | 1e-10 | -2.403e+01 | 5.11% | 0.60% | 64.2bp (64.9bp) | ZNF317/MA1593.2/ Jaspar(0.867) More Information \| Similar Motifs Found | motif file (matrix) |
| 9 * | 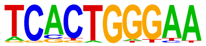 | 1e-10 | -2.376e+01 | 6.61% | 1.11% | 54.4bp (62.3bp) | RBPJ/MA1116.2/ Jaspar(0.801) More Information \| Similar Motifs Found | motif file (matrix) |
| 10 * | 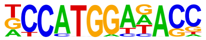 | 1e-10 | -2.373e+01 | 4.20% | 0.37% | 58.6bp (63.0bp) | Rfx1(HTH)/NPC-H3K4me1-ChIP-Seq(GSE16256)/ Homer(0.812) More Information \| Similar Motifs Found | motif file (matrix) |
| 11 * | | 1e-10 | -2.337e+01 | 1.80% | 0.02% | 52.8bp (78.7bp) | POU5F1/MA1115.2/ Jaspar(0.775) | motif file (matrix) |

| | | | | | | | | |
|---|---|---|---|---|---|---|---|---|
| | 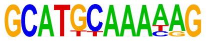 | | | | | | More Information \| Similar Motifs Found | |
| 12 * | 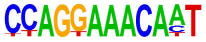 | 1e-10 | -2.332e+01 | 2.10% | 0.04% | 66.8bp (65.3bp) | FOXP2/MA0593.2/ Jaspar(0.715) More Information \| Similar Motifs Found | motif file (matrix) |
| 13 * | 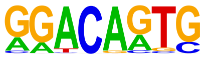 | 1e-9 | -2.271e+01 | 12.91% | 4.25% | 50.8bp (65.7bp) | Sox6/MA0515.1/ Jaspar(0.802) More Information \| Similar Motifs Found | motif file (matrix) |
| 14 * | 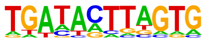 | 1e-9 | -2.251e+01 | 11.41% | 3.44% | 51.4bp (61.6bp) | PH0039.1_Mnx1/ Jaspar(0.625) More Information \| Similar Motifs Found | motif file (matrix) |
| 15 * | 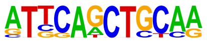 | 1e-9 | -2.238e+01 | 3.90% | 0.34% | 47.5bp (68.5bp) | MYOG/MA0500.3/ Jaspar(0.739) More Information \| Similar Motifs Found | motif file (matrix) |
| 16 * | 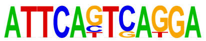 | 1e-9 | -2.191e+01 | 1.80% | 0.02% | 59.3bp (54.5bp) | PB0028.1_Hbp1_1/ Jaspar(0.720) More Information \| Similar Motifs Found | motif file (matrix) |
| 17 * | 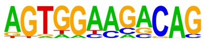 | 1e-9 | -2.105e+01 | 4.20% | 0.45% | 44.4bp (64.8bp) | Tbox:Smad(T-box,MAD)/ESCd5-Smad2_3-ChIP-Seq(GSE29422)/ Homer(0.687) More Information \| Similar Motifs Found | motif file (matrix) |
| 18 * | | 1e-8 | -1.933e+01 | 3.90% | 0.43% | 52.9bp (60.5bp) | ZIM3/MA1709.2/ Jaspar(0.658) | motif file (matrix) |

| | | | | | | | | |
|---|---|---|---|---|---|---|---|---|
| | 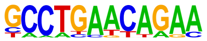  | | | | | | More Information \| Similar Motifs Found | |
| 19 * | 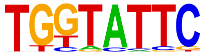  | 1e-7 | -1.802e+01 | 18.62% | 8.78% | 60.2bp (62.0bp) | PB0119.1_Foxa2_2/ Jaspar(0.702) More Information \| Similar Motifs Found | motif file (matrix) |
| 20 * | 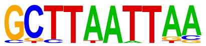  | 1e-7 | -1.800e+01 | 3.60% | 0.40% | 52.9bp (63.0bp) | LMX1B/MA0703.3/ Jaspar(0.890) More Information \| Similar Motifs Found | motif file (matrix) |
| 21 * | 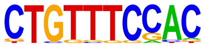  | 1e-7 | -1.793e+01 | 6.61% | 1.53% | 55.5bp (64.3bp) | Foxo1(Forkhead)/RAW-Foxo1-ChIP-Seq(Fan_et_al.)/ Homer(0.759) More Information \| Similar Motifs Found | motif file (matrix) |
| 22 * | 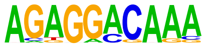  | 1e-7 | -1.782e+01 | 6.01% | 1.27% | 54.8bp (62.4bp) | ZNF675(Zf)/HEK293-ZNF675.GFP-ChIP-Seq(GSE58341)/ Homer(0.731) More Information \| Similar Motifs Found | motif file (matrix) |
| 23 * | 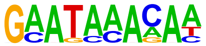  | 1e-7 | -1.743e+01 | 24.02% | 12.93% | 58.3bp (65.4bp) | Foxq1/MA0040.2/ Jaspar(0.804) More Information \| Similar Motifs Found | motif file (matrix) |
| 24 * | 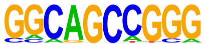  | 1e-6 | -1.581e+01 | 4.20% | 0.69% | 63.4bp (67.4bp) | Zic3/MA0697.3/ Jaspar(0.742) More Information \| Similar Motifs Found | motif file (matrix) |
| 25 * | | 1e-6 | -1.474e+01 | 3.30% | 0.44% | 57.0bp (66.4bp) | ETV1/MA0761.3/ Jaspar(0.817) | motif file (matrix) |

| | | | | | | | | |
|---|---|---|---|---|---|---|---|---|
| | 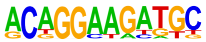 | | | | | | More Information \| Similar Motifs Found | |
| 26 * | 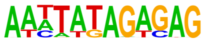 | 1e-6 | -1.417e+01 | 2.40% | 0.21% | 50.7bp (65.5bp) | ZNF768/MA1731.2/ Jaspar(0.598) More Information \| Similar Motifs Found | motif file (matrix) |
| 27 * | 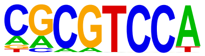 | 1e-5 | -1.364e+01 | 5.41% | 1.37% | 56.4bp (61.7bp) | Foxn1/MA1684.1/ Jaspar(0.791) More Information \| Similar Motifs Found | motif file (matrix) |
| 28 * | 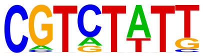 | 1e-5 | -1.321e+01 | 6.31% | 1.87% | 55.3bp (62.1bp) | BARHL1/MA0877.4/ Jaspar(0.665) More Information \| Similar Motifs Found | motif file (matrix) |
| 29 * | 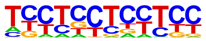 | 1e-3 | -7.660e+00 | 8.11% | 3.99% | 50.3bp (58.1bp) | ZNF148/MA1653.2/ Jaspar(0.636) More Information \| Similar Motifs Found | motif file (matrix) |

### Motifs found for: mouse survival hazard (Cox)

### Homer *de novo* Motif Results (homerpos15/)

Non-redundant Motif File of Results
Known Motif Enrichment Results
Gene Ontology Enrichment Results
If Homer is having trouble matching a motif to a known motif, try copy/pasting the matrix file into STAMP
More information on motif finding results: HOMER | Description of Results | Tips
Total target sequences = 462
Total background sequences = 95584
* - possible false positive

| Rank | Motif | P-value | log P-pvalue | % of Targets | % of Background | STD(Bg STD) | Best Match/Details | Motif File |
|---|---|---|---|---|---|---|---|---|
| 1 | 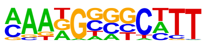 | 1e-33 | -7.692e+01 | 21.00% | 4.86% | 49.5bp (65.6bp) | Sox9(HMG)/Limb-SOX9-ChIP-Seq(GSE73225)/Homer(0.639) More Information \| Similar Motifs Found | motif file (matrix) |
| 2 | 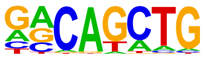 | 1e-30 | -6.969e+01 | 43.07% | 19.47% | 54.0bp (71.4bp) | Twist2(bHLH)/Myoblast-Twist2.Ty1-ChIP-Seq(GSE127998)/Homer(0.959) More Information \| Similar Motifs Found | motif file (matrix) |
| 3 | 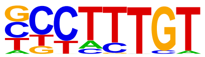 | 1e-29 | -6.743e+01 | 41.34% | 18.49% | 57.7bp (67.7bp) | SOX4/MA0867.3/Jaspar(0.945) More Information \| Similar Motifs Found | motif file (matrix) |
| 4 | | 1e-22 | -5.210e+01 | 30.95% | 13.14% | 55.9bp (66.9bp) | Hoxd10(Homeobox)/ChickenMSG-Hoxd10.Flag-ChIP- | motif file (matrix) |

| | | | | | | | | |
|---|---|---|---|---|---|---|---|---|
| | 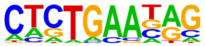 | | | | | | Seq(GSE86088)/ Homer(0.648) More Information \| Similar Motifs Found | |
| 5 | 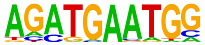 | 1e-14 | -3.316e+01 | 9.74% | 2.39% | 56.5bp (66.1bp) | PB0178.1_Sox8_2/ Jaspar(0.767) More Information \| Similar Motifs Found | motif file (matrix) |
| 6 | 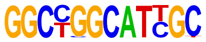 | 1e-13 | -3.138e+01 | 1.52% | 0.01% | 38.9bp (83.7bp) | PB0029.1_Hic1_1/ Jaspar(0.727) More Information \| Similar Motifs Found | motif file (matrix) |
| 7 | 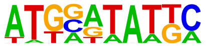 | 1e-12 | -2.814e+01 | 6.71% | 1.34% | 54.2bp (67.5bp) | Pit1(Homeobox)/GCrat-Pit1-ChIP-Seq(GSE58009)/ Homer(0.887) More Information \| Similar Motifs Found | motif file (matrix) |
| 8 * | 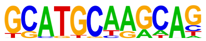 | 1e-10 | -2.505e+01 | 2.38% | 0.12% | 56.4bp (68.8bp) | POU5F1/MA1115.2/ Jaspar(0.682) More Information \| Similar Motifs Found | motif file (matrix) |
| 9 * | 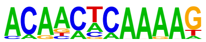 | 1e-10 | -2.501e+01 | 3.90% | 0.46% | 57.8bp (60.5bp) | MYNN(Zf)/HEK293-MYNN.eGFP-ChIP-Seq(Encode)/ Homer(0.644) More Information \| Similar Motifs Found | motif file (matrix) |
| 10 * | 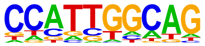 | 1e-10 | -2.501e+01 | 33.77% | 20.38% | 53.7bp (68.2bp) | NFIA/MA0670.2/ Jaspar(0.756) More Information \| Similar Motifs Found | motif file (matrix) |
| 11 * | | 1e-10 | -2.448e+01 | 3.25% | 0.30% | 52.4bp (70.4bp) | Elk4(ETS)/Hela-Elk4-ChIP-Seq(GSE31477)/ | motif file (matrix) |

| | | | | | | | | |
|---|---|---|---|---|---|---|---|---|
| | 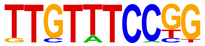 | | | | | | Homer(0.765) More Information \| Similar Motifs Found | |
| 12 * | 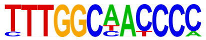 | 1e-10 | -2.426e+01 | 1.30% | 0.01% | 61.8bp (30.7bp) | NFIA/MA0670.2/ Jaspar(0.708) More Information \| Similar Motifs Found | motif file (matrix) |
| 13 * | 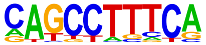 | 1e-10 | -2.403e+01 | 13.64% | 5.46% | 56.8bp (64.5bp) | FEZF2/MA2341.1/ Jaspar(0.689) More Information \| Similar Motifs Found | motif file (matrix) |
| 14 * | 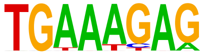 | 1e-9 | -2.274e+01 | 17.75% | 8.43% | 48.7bp (67.8bp) | IRF4(IRF)/GM12878-IRF4-ChIP-Seq(GSE32465)/ Homer(0.674) More Information \| Similar Motifs Found | motif file (matrix) |
| 15 * | 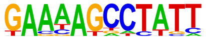 | 1e-9 | -2.269e+01 | 3.46% | 0.40% | 56.8bp (62.3bp) | ZNF341/MA1655.2/ Jaspar(0.707) More Information \| Similar Motifs Found | motif file (matrix) |
| 16 * | 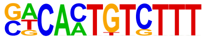 | 1e-9 | -2.257e+01 | 1.95% | 0.08% | 57.6bp (62.6bp) | Sox6/MA0515.1/ Jaspar(0.616) More Information \| Similar Motifs Found | motif file (matrix) |
| 17 * | 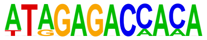 | 1e-9 | -2.242e+01 | 1.30% | 0.02% | 49.1bp (67.2bp) | Prdm14/MA1998.2/ Jaspar(0.819) More Information \| Similar Motifs Found | motif file (matrix) |
| 18 * | | 1e-9 | -2.221e+01 | 1.52% | 0.03% | 54.0bp (57.7bp) | ZNF467(Zf)/HEK293-ZNF467.GFP-ChIP- | motif file (matrix) |

| | | | | | | | | |
|---|---|---|---|---|---|---|---|---|
| | 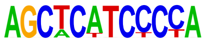 | | | | | | Seq(GSE58341)/ Homer(0.703) More Information \| Similar Motifs Found | |
| 19 * | 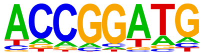 | 1e-9 | -2.148e+01 | 15.37% | 6.99% | 58.3bp (63.2bp) | ETS2/MA1484.2/ Jaspar(0.912) More Information \| Similar Motifs Found | motif file (matrix) |
| 20 * | 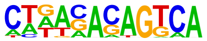 | 1e-8 | -2.025e+01 | 5.84% | 1.44% | 48.4bp (65.1bp) | Nr2e1/MA0676.1/ Jaspar(0.666) More Information \| Similar Motifs Found | motif file (matrix) |
| 21 * | 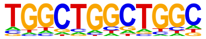 | 1e-8 | -1.966e+01 | 14.29% | 6.55% | 58.7bp (67.3bp) | Tlx?(NR)/NPC-H3K4me1-ChIP-Seq(GSE16256)/ Homer(0.719) More Information \| Similar Motifs Found | motif file (matrix) |
| 22 * | 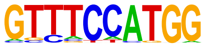 | 1e-8 | -1.934e+01 | 4.76% | 1.01% | 57.8bp (65.5bp) | Rfx1(HTH)/NPC-H3K4me1-ChIP-Seq(GSE16256)/ Homer(0.837) More Information \| Similar Motifs Found | motif file (matrix) |
| 23 * | 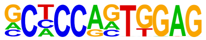 | 1e-7 | -1.788e+01 | 1.73% | 0.09% | 57.5bp (53.6bp) | Ebf4/MA2122.1/ Jaspar(0.617) More Information \| Similar Motifs Found | motif file (matrix) |
| 24 * | 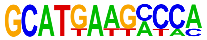 | 1e-7 | -1.705e+01 | 1.52% | 0.07% | 53.7bp (63.3bp) | PB0178.1_Sox8_2/ Jaspar(0.595) More Information \| Similar Motifs Found | motif file (matrix) |
| 25 * | | 1e-6 | -1.443e+01 | 3.46% | 0.74% | 63.1bp (53.3bp) | ZNF281/MA1630.3/ Jaspar(0.891) | motif file (matrix) |

| | | | | | | | | |
|---|---|---|---|---|---|---|---|---|
| | 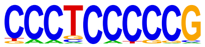 | | | | | | [More Information](#) \| [Similar Motifs Found](#) | |
| 26 * | 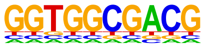 | 1e-5 | -1.365e+01 | 0.65% | 0.00% | 5.0bp (26.5bp) | ZNF417/MA1727.2/ Jaspar(0.673) [More Information](#) \| [Similar Motifs Found](#) | [motif file (matrix)](#) |
| 27 * | 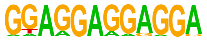 | 1e-3 | -7.178e+00 | 1.95% | 0.51% | 39.9bp (56.7bp) | ZFP14/MA1972.1/ Jaspar(0.661) [More Information](#) \| [Similar Motifs Found](#) | [motif file (matrix)](#) |

### Motifs found for: weibull alpha

### Homer *de novo* Motif Results (homerpos16/)

Non-redundant Motif File of Results
Known Motif Enrichment Results
Gene Ontology Enrichment Results
If Homer is having trouble matching a motif to a known motif, try copy/pasting the matrix file into STAMP
More information on motif finding results: HOMER | Description of Results | Tips
Total target sequences = 1536
Total background sequences = 96866
* - possible false positive

| Rank | Motif | P-value | log P-pvalue | % of Targets | % of Background | STD(Bg STD) | Best Match/Details | Motif File |
|---|---|---|---|---|---|---|---|---|
| 1 | 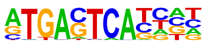 | 1e-506 | -1.166e+03 | 47.14% | 4.90% | 51.4bp (64.6bp) | Atf3(bZIP)/GBM-ATF3-ChIP-Seq(GSE33912)/Homer(0.992) More Information \| Similar Motifs Found | motif file (matrix) |
| 2 | 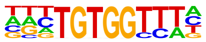 | 1e-64 | -1.489e+02 | 15.62% | 4.32% | 52.8bp (63.1bp) | Runx1/MA0002.3/Jaspar(0.976) More Information \| Similar Motifs Found | motif file (matrix) |
| 3 | 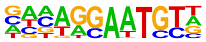 | 1e-35 | -8.233e+01 | 16.60% | 7.06% | 53.4bp (62.4bp) | TEAD2/MA1121.2/Jaspar(0.885) More Information \| Similar Motifs Found | motif file (matrix) |
| 4 | | 1e-25 | -5.864e+01 | 42.58% | 29.86% | 54.4bp (62.6bp) | SOX15/MA1152.2/Jaspar(0.954) | motif file (matrix) |

| | | | | | | | | |
|---|---|---|---|---|---|---|---|---|
| | 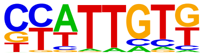 | | | | | | More Information \| Similar Motifs Found | |
| 5 | 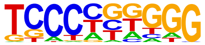 | 1e-18 | -4.189e+01 | 24.15% | 15.47% | 54.4bp (60.8bp) | Ebf4/MA2122.1/ Jaspar(0.820) More Information \| Similar Motifs Found | motif file (matrix) |
| 6 | 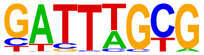 | 1e-17 | -3.927e+01 | 14.71% | 8.15% | 54.0bp (63.3bp) | CRX(Homeobox)/ Retina-Crx-ChIP-Seq(GSE20012)/ Homer(0.688) More Information \| Similar Motifs Found | motif file (matrix) |
| 7 | 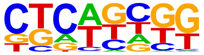 | 1e-15 | -3.637e+01 | 14.39% | 8.12% | 58.4bp (63.4bp) | NHLH1/MA0048.3/ Jaspar(0.618) More Information \| Similar Motifs Found | motif file (matrix) |
| 8 | 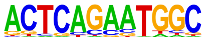 | 1e-15 | -3.522e+01 | 1.89% | 0.26% | 56.2bp (63.0bp) | Yy1/MA0095.4/ Jaspar(0.662) More Information \| Similar Motifs Found | motif file (matrix) |
| 9 | 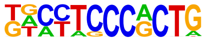 | 1e-13 | -3.060e+01 | 1.76% | 0.27% | 48.6bp (62.8bp) | E2F6(E2F)/Hela-E2F6-ChIP-Seq(GSE31477)/ Homer(0.737) More Information \| Similar Motifs Found | motif file (matrix) |
| 10 | 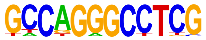 | 1e-12 | -2.848e+01 | 0.65% | 0.02% | 60.3bp (45.6bp) | Zfx/MA0146.3/ Jaspar(0.761) More Information \| Similar Motifs Found | motif file (matrix) |
| 11 * | | 1e-11 | -2.729e+01 | 8.66% | 4.50% | 53.9bp (61.9bp) | HIC1(Zf)/Treg-ZBTB29-ChIP- | motif file (matrix) |

| | | | | | | | | |
|---|---|---|---|---|---|---|---|---|
| | 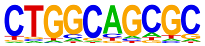 | | | | | | Seq(GSE99889)/ Homer(0.736) More Information \| Similar Motifs Found | |
| 12 * | 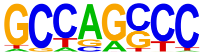 | 1e-11 | -2.565e+01 | 14.13% | 8.83% | 59.4bp (61.4bp) | ZNF711(Zf)/SHSY5Y-ZNF711-ChIP-Seq(GSE20673)/ Homer(0.716) More Information \| Similar Motifs Found | motif file (matrix) |
| 13 * | 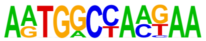 | 1e-10 | -2.474e+01 | 0.85% | 0.06% | 48.4bp (62.5bp) | POL004.1_CCAAT-box/ Jaspar(0.637) More Information \| Similar Motifs Found | motif file (matrix) |
| 14 * | 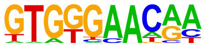 | 1e-10 | -2.468e+01 | 3.45% | 1.19% | 51.2bp (62.0bp) | Rbpj1(?)/Panc1-Rbpj1-ChIP-Seq(GSE47459)/ Homer(0.844) More Information \| Similar Motifs Found | motif file (matrix) |
| 15 * | 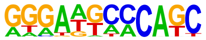 | 1e-10 | -2.334e+01 | 3.32% | 1.16% | 52.3bp (62.3bp) | Zfp809/MA2125.1/ Jaspar(0.678) More Information \| Similar Motifs Found | motif file (matrix) |
| 16 * | 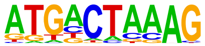 | 1e-9 | -2.284e+01 | 9.11% | 5.16% | 57.6bp (60.9bp) | FOSL1::JUND/ MA1142.2/ Jaspar(0.689) More Information \| Similar Motifs Found | motif file (matrix) |
| 17 * | 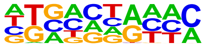 | 1e-9 | -2.269e+01 | 4.88% | 2.15% | 51.7bp (62.5bp) | Pit1(Homeobox)/GCrat-Pit1-ChIP-Seq(GSE58009)/ Homer(0.519) More Information \| Similar Motifs Found | motif file (matrix) |
| 18 * | | 1e-6 | -1.589e+01 | 0.65% | 0.07% | 55.4bp (54.8bp) | BACH1/MA1633.2/ Jaspar(0.576) | motif file (matrix) |

| | | | | | | | | |
|---|---|---|---|---|---|---|---|---|
| | 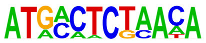 | | | | | | More Information \| Similar Motifs Found | |
| 19 * | 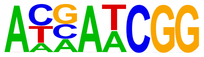 | 1e-6 | -1.383e+01 | 2.21% | 0.86% | 55.6bp (59.2bp) | OVOL2/MA1545.2/ Jaspar(0.668) More Information \| Similar Motifs Found | motif file (matrix) |
| 20 * | 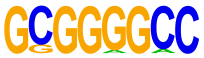 | 1e-5 | -1.274e+01 | 2.67% | 1.19% | 54.1bp (52.7bp) | Sp1(Zf)/Promoter/ Homer(0.812) More Information \| Similar Motifs Found | motif file (matrix) |
| 21 * | 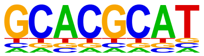 | 1e-4 | -1.096e+01 | 0.72% | 0.14% | 62.1bp (61.7bp) | Arnt:Ahr(bHLH)/ MCF7-Arnt-ChIP-Seq(Lo_et_al.)/ Homer(0.832) More Information \| Similar Motifs Found | motif file (matrix) |
| 22 * | 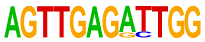 | 1e-3 | -8.991e+00 | 0.13% | 0.00% | 41.1bp (40.2bp) | RUNX3/MA0684.3/ Jaspar(0.649) More Information \| Similar Motifs Found | motif file (matrix) |
| 23 * | 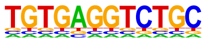 | 1e-1 | -4.151e+00 | 0.07% | 0.00% | 27.0bp (0.0bp) | Hand1/MA2123.1/ Jaspar(0.673) More Information \| Similar Motifs Found | motif file (matrix) |
| 24 * | 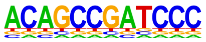 | 1e-1 | -4.151e+00 | 0.07% | 0.00% | 19.0bp (0.0bp) | ZNF317/MA1593.2/ Jaspar(0.689) More Information \| Similar Motifs Found | motif file (matrix) |
